# Functional and molecular early enteric biomarkers for Parkinson’s disease in mice and men

**DOI:** 10.1101/2020.06.06.136556

**Authors:** Manuela Gries, Anne Christmann, Steven Schulte, Maximilian Weyland, Stephanie Rommel, Monika Martin, Marko Baller, Ralph Röth, Stefanie Schmitteckert, Marcus Unger, Yang Liu, Frederik Sommer, Timo Mühlhaus, Michael Schroda, Jean-Pierre Timmermans, Isabel Pintelon, Gudrun A. Rappold, Markus Britschgi, Hilal Lashuel, Michael D. Menger, Matthias W. Laschke, Beate Niesler, Karl-Herbert Schäfer

## Abstract

Parkinson’s disease (PD) usually has a late clinical onset. The lack of early biomarkers for the disease represents a major challenge for developing timely treatment interventions. Here, we use an α-synuclein-overexpressing transgenic (Th-1-SNCA-A30P) mouse model of PD to identify appropriate candidate markers in the gut for early stages of PD before hallmark symptoms begin to manifest. A30P mice did not show alterations in gait parameters at 2 months of age, and these mice were therefore defined as pre-symptomatic A30P mice (psA30P). We discovered early functional motility changes in the gut and early molecular dysregulations in the myenteric plexus of psA30P mice by comparative protein and miRNA profiling and cell culture experiments. We found that the proteins neurofilament light chain, vesicle-associated membrane protein 2 and calbindin 2, together with the miRNAs that regulate them, are potential biomarkers of early PD that may facilitate timely treatment and/or prevention of PD in men.

## Introduction

In Parkinson’s disease (PD), the loss of nigrostriatal dopaminergic neurons causes the hallmark motor symptoms of muscle rigidity, tremor at rest, and bradykinesia^1,2,3^. In recent years, early non-motor symptoms of PD have gained attention; these include sleep disturbance, pain, depression, and constipation, and can occur more than 15 years before clinical motor symptoms manifest^4,5,6^. Histopathological examination of postmortem PD brains has revealed intraneuronal accumulations of misfolded α-synuclein, termed Lewy bodies^7,8^. These Lewy bodies disrupt protein ubiquitination, cytoskeleton reorganization, synaptic function and vesicle release, and increase oxidative stress levels to promote PD pathogenesis^9^. Lewy bodies are present not only in the central nervous system (CNS) but also in the enteric nervous system (ENS) of the intestine, so it is not only the brain but also the gut that is involved in PD pathogenesis^10,11^. In 2002, Del Tredici et al. postulated that neurotoxic α-synucleins may originate in the gut and migrate to the brain via the vagal nerve, the gut-brain-axis, causing neuronal cell death and inflammation in the substantia nigra^12,13,14,15,16^. This theory was recently supported by studies in rat models^17,18,19^, which confirmed that α-synuclein transport can be purged by vagal nerve resection^20^. Previous research has shown that crosstalk between the gut and the brain during PD pathogenesis is mainly influenced by intestinal dysbiosis^21,22,23,24^. This suggests that the gastrointestinal (GI) motility deficits associated with PD, such as constipation, are caused by alterations in the microbial composition of the gut, which disrupt gut homeostasis and increase inflammation and permeability of the gut^25,26,27^. These findings indicate that PD originates in the gut, and that a dysregulated gut microbiome and gut-brain-axis may contribute to PD symptoms. However, changes in the ENS during PD onset have not been well-described. To better understand the role of the gut in PD, a more detailed investigation of the early phase of PD within the gut is essential. The current study aimed to assess early pathological changes associated with PD in the ENS to identify biomarkers of pre-clinical PD. In recent years, miRNAs have emerged as important regulators of gene expression, cell differentiation, cell maturation, apoptosis, and immune responses^28,29^. They are present in circulating body fluids^30^ as well as gut mucosal tissues, so they can be investigated by colonoscopy^31^. Abnormal miRNA expression has been linked to many human diseases, and miRNAs have been identified as diagnostic and prognostic biomarkers for several disorders, including PD^32,33,34,35,36^. However, no PD-specific miRNAs have been identified in the gut for the pre-clinical diagnosis of PD so far^37,38,39^. To address this, we studied functional and molecular changes in the ENS of A30P mice before onset of PD symptoms to identify gut-related biomarkers for early stages of PD.

## Results

### Gait analysis showed 2-month-old A30P mice are pre-symptomatic (psA30P mice)

Gait analysis has been used to show onset of obvious PD manifestations in mice^40^. We analyzed ten gait parameters of A30P and wild type (WT) mice using the CatWalk XT system. Animal footprints were automatically captured by the video camera of the system and were further categorized into individual paws like right forepaw (RF), right hindpaw (RH), left forepaw (LF), and left hindpaw (LH) via the software (Fig. 1a). To characterize pre-symptomatic PD or phenotype-manifested PD in mice, we assessed the gait in two age groups: 2 months old and 12–13 months old. Body mass was not different between 2-month-old A30P and WT mice, but 12–13-month-old A30P mice were significantly heavier than age-matched WT mice (Supplementary Fig. 1).

**Figure 1.**
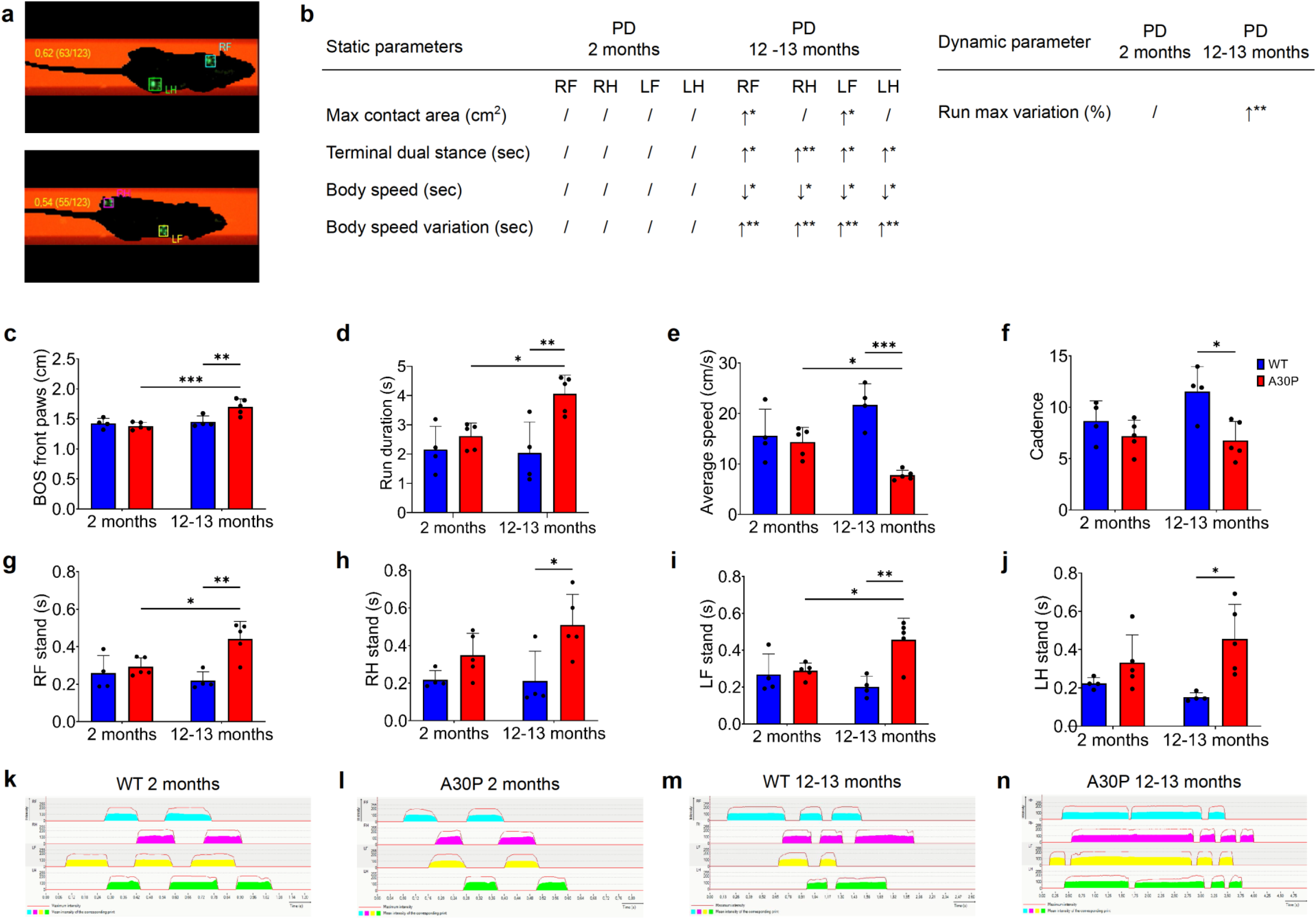
CatWalk XT analysis of different aged A30P and wild type (WT) mice. (a) Individual paws were classified automatically with the CatWalk XT software as right forepaw (RF), right hindpaw (RH), left forepaw (LF), and left hindpaw (LH). (b) Dynamic and static parameters were all significantly different in older A30P mice compared with age-matched controls, while 2-month-old A30P and WT mice did not show any motoric disturbances. (c) Base of support on front paws and run duration were significantly increased in 12–13-month-old A30P mice (d), while the average speed (e) and (f) cadence were decreased. Stands of all paws, including RF (g), RH (h), LF (i), and LH (j) was significantly augmented in 12–13-month-old A30P mice compared with WT animals. This was confirmed by corresponding 2-D maps from WT (k,m) and A30P mice (l,n). Two-month-old A30P mice showed no motoric changes, which was confirmed by 2-D maps. WT mice are shown in blue (n = 4), A30P mice in red (n = 5). 2-D maps determine the gait sequence of paw contact with the glass plate over time. * *p* ≤ 0.05, ** *p* ≤ 0.01, and *** *p* ≤ 0.001 using ANOVA.

As expected, the older A30P mice showed significant motor impairments in all tested parameters: maximum contact area, terminal dual stance, body speed, body speed variation, run maximum variation (*p* ≤ 0.05, Fig. 1b and Supplementary Fig. 2), base of support on front paws (Fig. 1c), run duration (Fig. 1d), average speed (Fig. 1e) and cadence (Fig. 1f). In addition, the older mouse group displayed significantly extended stands (*p* ≤ 0.05, Fig. 1g–j) and markedly changed footprint patterns from all paws (Fig. 1k-n). In contrast, younger A30P mice showed no significant changes in all investigated gait parameters compared with age-matched WT mice, indicating that these mice have a pre-symptomatic stage of the disease.

We also measured gut length in these mice. Gut length did not differ between A30P and WT mice in both age groups, but both intestines were significantly longer in 12–13-month-old A30P and WT mice compared with 2-month-old A30P and WT mice (Supplementary Fig. 3). Based on these findings, we defined the non-motor-impaired 2-month-old mice as the pre-symptomatic A30P mouse group (psA30P), and used these mice for further investigations in the gut.

### Altered GI motility in psA30P mice

To investigate the hypothesis that the gut is the first site of PD manifestation, we compared GI motility patterns in the small intestine (SI) and the large intestine (LI) of psA30P mice and age-matched WT animals in isolated gut segments. We measured GI movement using several parameters, including changes in amplitude (Δ amplitude), number of contractions, mean interval (Fig. 2a), and velocity of motions (Fig. 2b).

**Figure 2:**
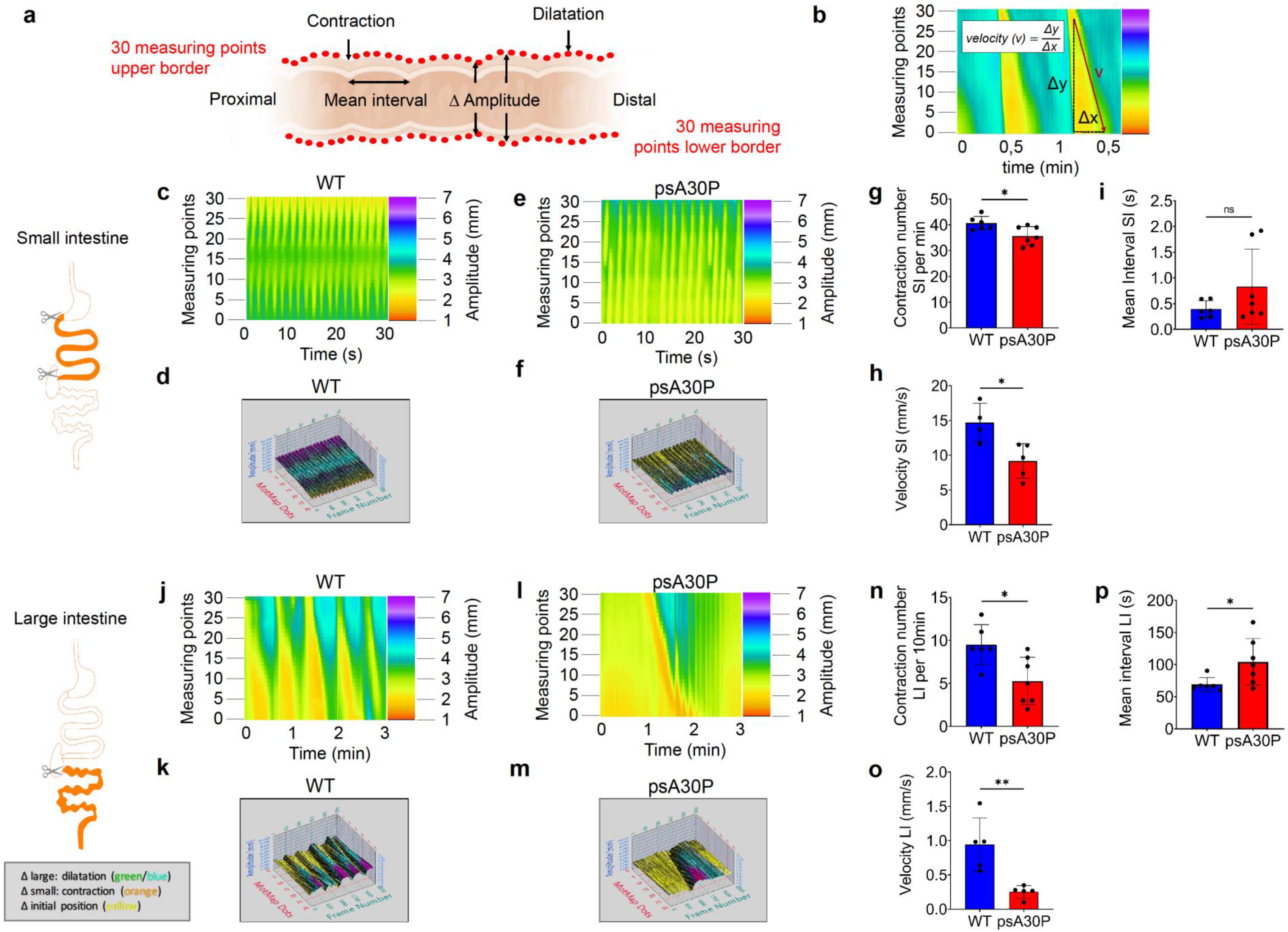
Motility patterns during *ex vivo* luminal perfusion experiments in pre-symptomatic (ps)A30P mice and wild type (WT) controls. (a) Video recordings of gut movements were analyzed with MotMap and LabView to determine alterations in amplitudes Δ, mean intervals and contraction numbers. (b) Schematic illustration of the velocity measured by LabView. Spatiotemporal maps and corresponding 3-D MotMaps showed altered small intestine (SI) motility (c–f) and large intestine (LI) motility (j–m). Less SI and LI motility was shown in psA30P mice compared with WT. The gut diameter in the heatmaps (pixel color) is indicated on the y-axis, the time on the x-axis. Contraction numbers were significantly lower in the SI (g) and LI (n) of psA30P mice compared with WT controls. Mean intervals were slightly higher with more variation in the SI of psA30P mice compared with WT (i), while in the LI of psA30P mice mean intervals were significantly prolonged compared with WT (p). Velocity was significantly decreased in the SI and LI of psA30P mice compared with WT (h,o). WT mice are shown in blue, psA30P mice in red. Calculations were performed with LabView and GraphPad Prism 8. Quantitative data are expressed as means ± SD from segments of n = 7 psA30P mice and n = 6 age-matched WT controls for mean interval and contraction numbers. For velocity determination, n = 5 psA30P mice and n = 5 WT mice were examined. * *p* ≤ 0.05, ** *p* ≤ 0.01, and *** *p* ≤ 0.001 using Student’s t test and Cohen’s d (Supplementary Table 6a).

Both SI and LI had an isochronous motility pattern in WT mice (SI: Fig. 2c,d; LI: Fig. 2j,k), whereas psA30P mice showed a discontinuous outline with extended passive interphases (SI: Fig. 2e,f; LI: Fig. 2l,m).

In the SI, more contractions were observed in WT mice (40.7 ± 2.6 contractions/min) than in psA30P mice (35.7 ± 3.6 contractions/min; *p* ≤ 0.05, Fig. 2g). Furthermore, the movements of WT mice were significantly faster (14.7 ± 2.8 mm/s) than those of psA30P mice (9.2 ± 2.2 mm/s; *p* ≤ 0.05, Fig. 2h). The mean interval was higher in psA30P mice because of extended and variable interphases of (0.50 ± 0.1 s) compared with WT mice (0.39 ± 0.7 s), but this difference was not significant (Fig. 2i).

In the LI, WT mice exhibited 9.5 ± 2.4 contractions (Fig. 2n) with an appropriate velocity of 0.9 ± 0.4 mm/s (Fig. 2o) over a 10-min period. The number (5.3 ± 2.6 contractions per 10 min; *p* ≤ 0.05, Fig. 2n) and velocity of contractions (0.3 ± 0.1 mm/s; *p* ≤ 0.01, Fig. 2o) were significantly lower in psA30P mice. In addition, the mean interval was significantly shorter in WT mice (69.1 ± 10.8 s) compared with in psA30P mice (104.4 ± 33.4 s; *p* ≤ 0.05, Fig. 2p).

The reduced gut motility in psA30P mice can be visualized in the video recordings (Supplementary Videos 1–4); these videos show retarded activity, slower contractions, and longer static phases in psA30P mice compared with WT mice (Fig. 2).

### Dysregulated protein expressions in the ENS of psA30P mice

To investigate the role of the ENS in PD pathogenesis, we dissected the myenteric plexus (MP) of the SI and the LI in psA30P and WT mice. Whole protein was isolated from the gut tissue, separated by high-performance liquid chromatography (HPLC), and further analyzed by mass spectroscopy.

The MP protein expression profile was different between psA30P and corresponding WT mice in the SI (Fig. 3a) and LI (Fig. 3b). In total, 1,044 proteins were detected by mass spectroscopy; in 7% of proteins, expression was significantly altered in the SI (Fig. 3a, Supplementary Fig. 4a, Supplementary Table 2) and almost twice as many proteins (14%) in the LI (Fig. 3b, Supplementary Fig. 4b and Supplementary Table 2), indicating a more serious manifestation of PD in the LI. Proteins with significant changes in expression were analyzed and visualized by Search Tool for the Retrieval of Interacting Genes/Proteins (STRING). Each dot represents a single protein, while clustered proteins or proteins in the direct neighborhood indicate a similar function (Fig. 3c and d; Supplementary Fig. 4).

**Figure 3:**
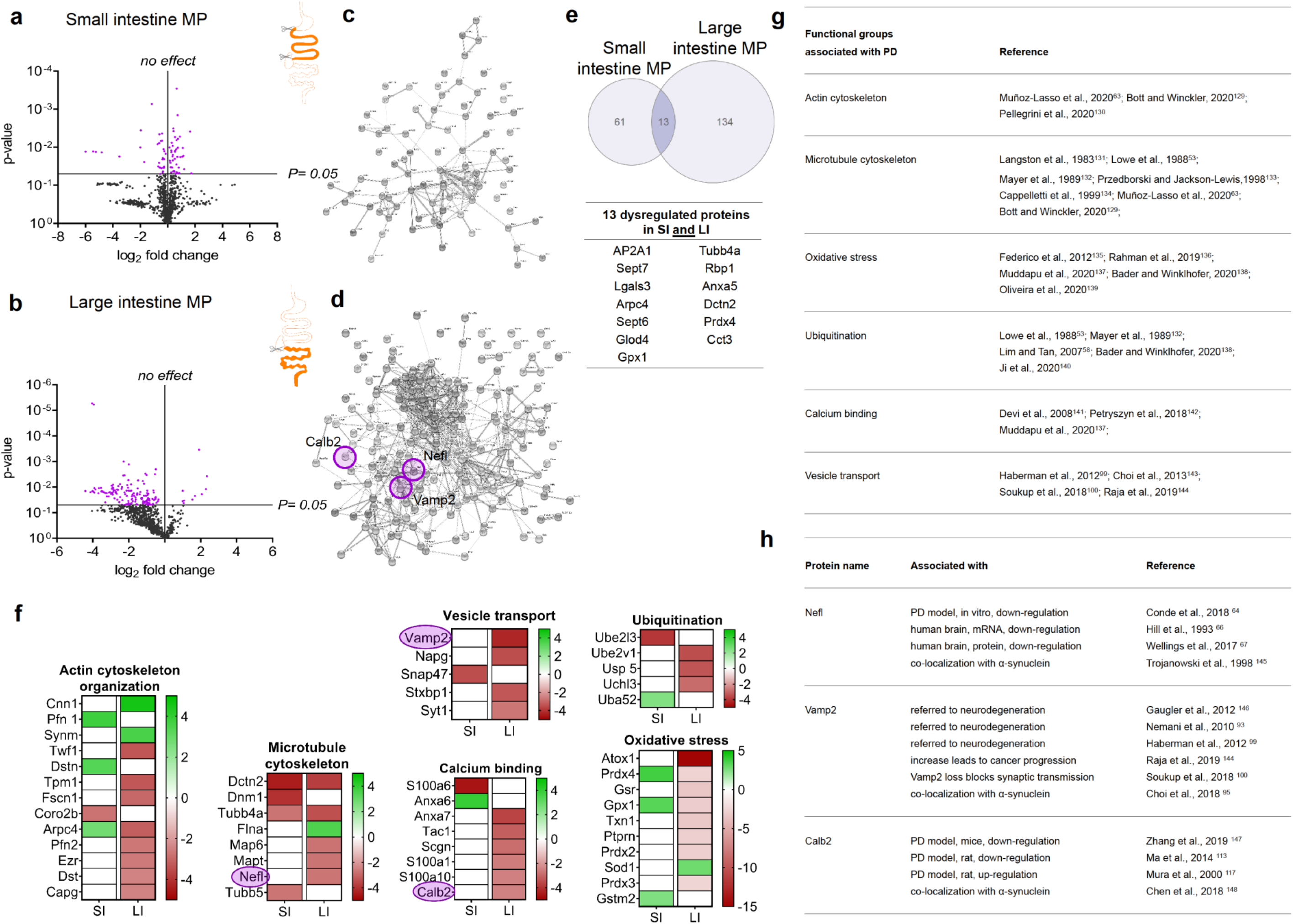
Mass spectroscopy of the myenteric plexus (MP) from pre-symptomatic (ps)A30P mice compared with wild type (WT) controls. Regulated proteins in the MP of the small intestine (SI, a) and large intestine (LI, b) were volcano plotted using GraphPad Prism 8 according to their statistical significance. *p*-Values were calculated using Student’s t test and Cohen’s d (Supplementary Table 6b). Proteins with significantly altered expression are shown in violet, proteins with non-significantly altered expression in black. Protein-protein interactions obtained from the STRING database are shown for MP samples of the SI (c) and LI (d). Neurofilament light chain (Nefl), vesicle-associated membrane protein 2 (Vamp2), and calbindin 2 (Calb2) are highlighted with violet circles. Classified functional groups of regulated proteins in the gut (actin and microtubule organization, vesicle transport, calcium binding, ubiquitination, and oxidative stress) are shown in (f). The intensity levels (percentage of normalized volumes) of the protein spots were shown on a heatmap according to their expression levels. Individual rows represent single spots and graduated scale color codes from red (decreased expression levels) to green (increased expression levels) and white (no altered expression levels). Each column represents data from n = 4 independent experiments. (e) In total, 74 proteins were affected in the SI and 147 in the LI. Among these, 13 proteins were changed significantly in both, the SI and the LI. The Venn diagram and the heatmaps were generated using GraphPad Prism 8. Here, only proteins with significantly altered expression were included. Categorized functional groups from the STRING database which have already been implicated with PD, and their corresponding references are shown in (g). (h) List of references which display an association of Nefl, Vamp2 and Calb2 with neurodegenerative diseases or PD.

We used a Venn diagram to illustrate the distribution of dysregulated proteins in the SI (74 proteins) and the LI (147 proteins); 13 proteins were differentially expressed in both tissues (Fig. 3e). Categorization into groups by the STRING database yielded i.a. six different functional clusters: actin and microtubule organization, vesicle transport, calcium binding, ubiquitination, and response to oxidative stress (Fig. 3f). Proteins in these groups are known to be involved in PD pathogenesis in mice and humans, mainly affecting the CNS (references in Fig. 3g). This functional classification emphasized that more proteins are dysregulated in the LI than in the SI. Therefore, we focused all further investigations on the LI. We selected three proteins with altered expression for detailed examinations: neurofilament light chain (Nefl), vesicle-associated membrane protein 2 (Vamp2) and calbindin 2 (Calb2). We chose these proteins because they have already been implicated in PD pathology. For example, Nefl and Calb2 expression is altered in the brain of humans with PD as well as in PD models, but nothing is known about how these proteins are affected in the gut during early stages of PD. In addition, both, Nefl and Calb2 can co-localize with α-synuclein (Fig. 3h). Several studies have shown a role for Vamp2 in neurodegeneration, but no direct link to PD pathogenesis has been described to date, although Vamp2 does bind to α-synuclein (Fig. 3h).

### Differential Nefl, Calb2, and Vamp2 expression in the ENS of psA30P mice

To evaluate the expression of Nefl, Calb2, and Vamp2 in the LI during pre-symptomatic early-onset PD, we performed immunohistochemically stains on whole-mounts of the LI from psA30P and WT mice (Fig. 4). Based on our proteomic data and because of their pivotal role in the ENS, we decided to see whether these proteins can serve as biomarkers for early PD. Preliminary assessments of the LI whole-mounts revealed a significantly smaller ganglionic area in psA30P mice than in WT mice (Supplementary Fig. 5).

**Figure 4:**
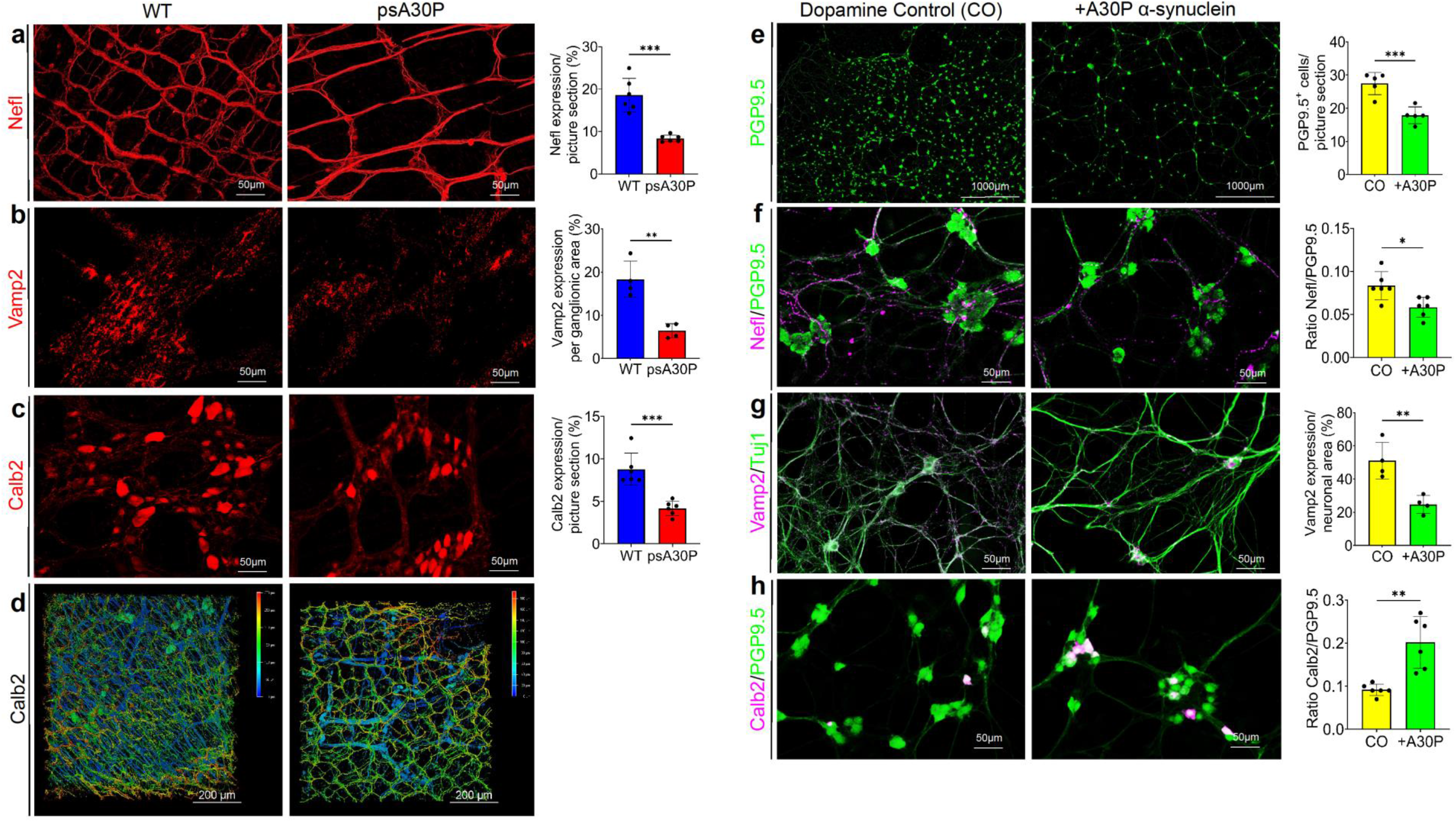
Effect of A30P α-synuclein on enteric cells. Distribution of (a) neurofilament light chain (Nefl), (b) vesicle-associated membrane protein 2 (Vamp2) and (c) calbindin 2 (Calb2), in full thickness muscle layers of the large intestine (LI) in psA30P and wild type (WT) mice. There were significantly fewer Nefl-, Vamp2-, and Calb2-positive cells in pre-symptomatic (ps)A30P mice than in WT mice. (d)Three-dimensional (3-D) images were made of full thickness intestinal walls immunostained for Calb2 and confirmed the 2-D impressions (c) of muscle layer stainings. WT mice are shown in blue, psA30P mice in red. Quantitative data were analyzed with ImageJ and are expressed as means ± SD from n = 5 independent experiments using GraphPad Prism 8. Nefl and Calb2 expressions were calculated per picture section, Vamp2 expression was analyzed per ganglionic area (class III beta tubulin (Tuj1) positive area). * *p* ≤ 0.05, ** *p* ≤ 0.01, and *** *p* ≤ 0.001 using Student’s t test and Cohen’s d (Supplementary Table 6c). 3-D images are presented using the depth-coding mode, where a depth color code corresponds to the position within the volume. The scale bar displays the spatial distribution of the color coding. The serosa side of whole-tissue samples was set at 0 μm and is depicted in blue. The top of the villi is displayed in red. Images are shown as a top view. For *in vitro* studies, myenteric plexus (MP) cells isolated from C57B6/J mice (postnatal day 2) were treated with 0.5 μM A30P α-synuclein for 5 days. This significantly reduced the number of protein gene product (PGP) 9.5-positive neurons (e). In addition, exposure to A30P α-synuclein significantly reduced the number of Nefl-positive cells in relation to total neuron number (f) and significantly reduced the expression of Vamp2 (g) compared with controls. The number of Calb2-positive cells significantly increased in relation to total neuron number (h). Nefl and Calb2 expression are indicated as a ratio to PGP9.5 and were normalized to the control. Vamp2 expression was only measured in neuronal areas. Non-treated cells (control = CO) are shown in yellow, A30P α-synuclein-treated cells are shown in green. Quantitative data were generated with ImageJ and are expressed as means ± SD from four to six independent experiments (N = 90 images per condition) using GraphPad Prism 8. * *p* ≤ 0.05, ** *p* ≤ 0.01, and *** *p* ≤ 0.001 using Student’s t and Cohen’s d (Supplementary Table 6d).

Nefl staining revealed a large number of myenteric ganglia that were evenly distributed over the underlying circular muscle layer. Nefl-positive cells were also frequently encountered in the interconnecting strands running parallel to the circular muscle layer. These fibers had a thinner and smoother morphology than those running longitudinally. In general, Nefl expression was weaker in psA30P mice (8.4 ± 0.8%) than in WT mice (18.6 ± 3.9%; *p* ≤ 0.001, Fig. 4a). Fiber density was especially reduced in the interconnecting strands. Consistently, there were significantly fewer synaptic vesicles in the muscle layers of psA30P mice (6.4 ± 1.6%) compared with in WT animals (18.3 ± 4.2%; *p* ≤ 0.01, Fig. 4b). Calb2 showed prominent immunoreactivity in the cytoplasm of neuronal soma and in axonal projections in both psA30P and WT mice. However, there were significantly more Calb2-positive cells in WT mice (8.8 ± 1.5%) than in psA30P mice (4.2 ± 0.86%; *p* ≤ 0.001, Fig. 4 c). Calb2 is expressed not only in the MP, but also in the submucosal plexus, which is located in the submucosal layer. Therefore, we imaged the whole gut in 3-D to include both plexuses, and confirmed the reduced Calb2 expression in psA30P mice (Fig. 4d).

### Differential Nefl, Calb2, and Vamp2 expression in primary ENS cells from an *in vitro* PD model

To verify the dose-dependent toxicity of the aggregated mutant A30P α-synuclein protein on primary MP cells, we performed a live-dead assay as a pilot experiment with different incubation times (Supplementary Fig. 6a). Based on the results, we chose a concentration of 0.5 μM A30P α-synuclein and an incubation time of 5 days for further experiments. To ascertain the cellular effects of A30P α-synuclein on MP cells, we isolated and cultivated MP cells and performed immunocytochemistry for protein gene product (PGP) 9.5, class III beta tubulin (Tuj1), Nefl, Calb2, and Vamp2. The total number of neurons was calculated as the number of PGP9.5-positive neurons, and was reduced after exposure to A30P α-synuclein (WT: 27.5 ± 3.4 PGP9.5-positive cells; A30P: 17.9 ± 2.5 PGP9.5-positive cells; *p* ≤ 0.001, Fig. 4e). There were also significantly fewer Nefl-positive cells after exposure to A30P α-synuclein (WT: 0.08 ± 0.02 ratio Nefl/PGP9.5; A30P: 0.06 ± 0.01 ratio Nefl/PGP9.5; *p* ≤ 0.05, Fig. 4f and Supplementary Fig. 6b). Examination of synaptic vesicles revealed significantly fewer Vamp2-positive signals in A30P α-synuclein-treated cells per neuronal area (WT: 53.7 ± 11.1%; A30P: 24.7 ± 5.4%; *p* ≤ 0.01, Fig. 4g). Conversely, we found nearly twice as many Calb2-positive neurons in α-synuclein-treated cells compared with unchallenged cultures (ratio Calb2/PGP9.5 WT: 0.09 ± 0.01; A30P: 0.2 ± 0.06, *p* ≤ 0.01, Fig. 4h and Supplementary Fig. 6c).

### Dysregulated miRNA expression in the ENS of psA30P mice

To measure miRNA expression in psA30P mice, a NanoString nCounter^®^ mouse expression assay was performed (Supplementary Table 3). Volcano plots showed that 166 miRNAs were robustly expressed (more than 100 counts) in the MP of psA30P mice and 210 miRNAs were in the mesencephalon of psA30P mice compared with WT mice (Supplementary Tables 4 and 5, Fig. 5a,b). This dysregulation was statistically significant in 45 miRNAs in the MP (Fig. 5a), and in only eight miRNAs in the mesencephalon (Fig. 5b). Three miRNAs were differentially expressed in both tissues, and the fold changes in expression of these miRNAs were distinctly higher in the LI than in the mesencephalon (Fig. 5c).

**Figure 5:**
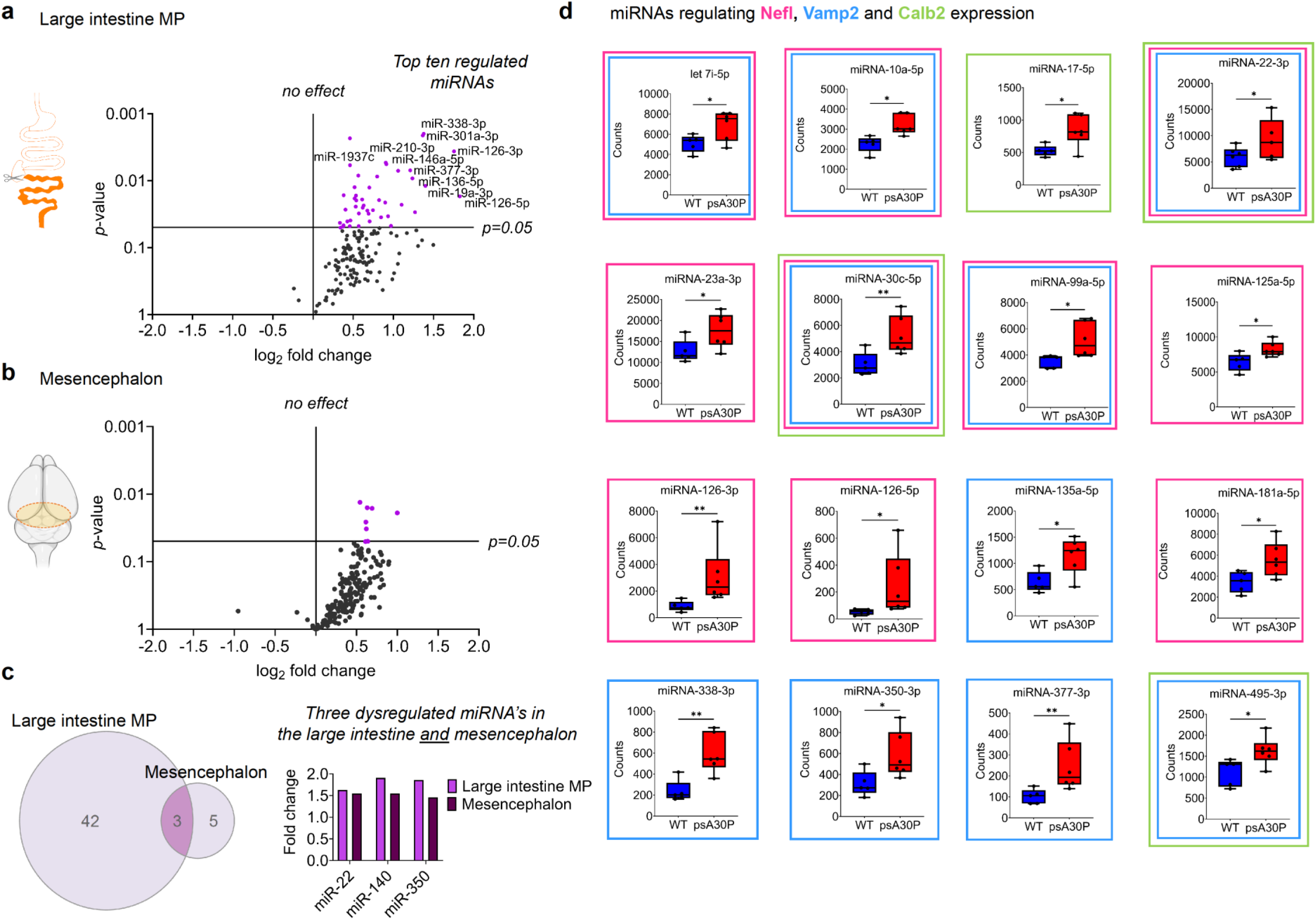
miRNA expression analysis of 2-month-old pre-symptomatic (ps)A30P mice. Volcano plots show miRNA profiling in the myenteric plexus (MP) of the large intestine (LI, a) and the mesencephalon (b) of psA30P mice compared with controls. Expression data for 578 miRNAs were generated by nCounter NanoString analysis. In total, 166 miRNAs were expressed in the MP of the LI (a) and 210 miRNAs in the mesencephalon (b) with a minimum of 100 counts. The left side represents the significance level of downregulated miRNAs and the right side the significance level of upregulated miRNAs. Violet circles indicate significantly dysregulated miRNAs; black circles indicate miRNAs whose expression was not significantly altered. The horizontal line in miRNA expression marks the threshold for the t-test *p* value (= 0.05). (c) In total, we identified 45 significantly upregulated miRNAs in the MP of the LI and eight upregulated miRNAs in the mesencephalon of psA30P mice. Among them, three are expressed in the MP of the LI and the mesencephalon, but with a higher fold change in the MP. (d) Box plots show regulated miRNAs in psA30P mice compared with wild types (WT) targeting neurofilament light chain (Nefl, pink framed boxes), vesicle-associated membrane protein 2 (Vamp2, blue framed boxes), and calbindin 2 (Calb2, green framed boxes). WT mice are shown in blue (n = 5), A30P mice in red (n = 6). Quantitative data are expressed as means ± SD using GraphPad Prism 8. * *p* ≤ 0.05, ** *p* ≤ 0.01, and *** *p* ≤ 0.001 using Student’s t test and Cohen’s d (Supplementary Table 6e).

The top ten regulated miRNAs with the highest fold change and the lowest *p*-value were miRNA-19a, 126-3p, 126-5p, 136-5p, 146a-5p, 210-3p, 301a-3p, 338-3p, 377-3p, and 1937c (Supplementary Figure 7 and Fig. 5a). Significant changes in expression were observed in miRNAs that target Nefl, Vamp2, and Calb2, supporting our idea that these proteins are potential biomarkers for early-stage PD. Detailed examination of protein-miRNA clustering using mirWALK 3.0^41^ revealed that Nefl (Fig. 5d, pink framed boxes) and Vamp2 (Fig. 5d, blue framed boxes) are both targeted by ten miRNAs, and Calb2 (Fig. 5d, green framed boxes) is clustering with four miRNAs. All miRNAs were significantly upregulated. Two miRNAs, miR-22-3p and miR-30c-5p, target our three selected proteins, Nefl, Vamp2 and Calb2.

### miRNA dysregulation is consistent in mice and men

To determine the functional impact of miRNA dysregulation on PD-associated protein networks, we performed an IPA-based network analysis using all 45 upregulated miRNAs and all 49 altered proteins from our defined functional groups (Fig. 6a). The majority of dysregulated miRNAs were involved in the same pathways as the proteins identified in our proteome analyses, such as protein ubiquitination, calcium signaling, cell and oxidative stress, synaptic transmission, and cytoskeleton assembly (Fig. 6a). These data revealed 77 protein-miRNA interactions, involving 25 proteins and 31 miRNAs, in which the proteins were downregulated and the respective miRNAs were upregulated, as well as eight protein-protein interactions.

**Figure 6:**
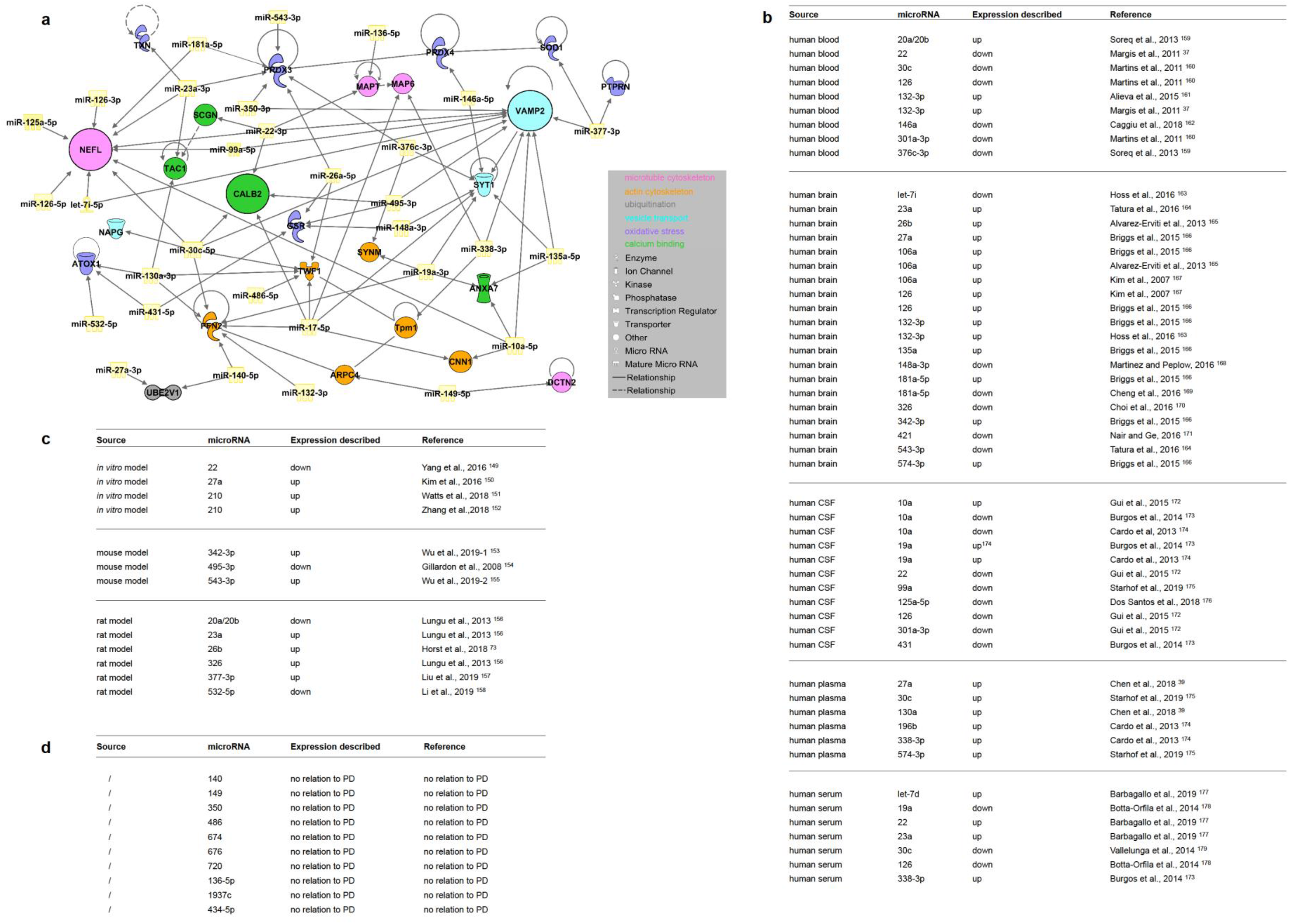
Ingenuity pathway analysis (IPA) of altered miRNAs in the myenteric plexus (MP) of the large intestine (LI) from pre-symptomatic (ps)A30P mice and their correlation to Parkinson’s disease (PD) models and patients. (a) IPA functional pathway analysis was used to predict the top transcriptional regulators from differentially expressed proteins in murine PD models. We identified 77 protein-miRNA interactions which involved 25 dysregulated proteins and 31 miRNAs and eight protein-protein interactions. These predictions are based on Ingenuity Pathway Knowledge Base (IPKB). Summary of dysregulated miRNAs in PD patients (b) and PD models (c) described in the literature, correlating with our regulated murine miRNAs in psA30P mice. (d) Ten murine miRNAs were altered in the MP of the LI in psA30P mice with no correlation to PD.

Nefl, Calb2, and Vamp2 clustered extensively with several miRNAs (Fig. 5d and 6a). Thirty-one miRNAs that were dysregulated in psA30P mice have been shown to be dysregulated in human PD tissues and body fluids (Fig. 6b), and 13 have been implicated in PD in mouse, rat, and *in vitro* models (Fig. 6c). However, these miRNAs were only investigated during clinical stages of PD and in the CNS, and not during pre-symptomatic stages and in the gut or MP like in the present study. In addition, we detected 10 significantly dysregulated miRNAs in the MP of the LI in psA30P mice, which, to best of our knowledge, have not been associated with PD to date (Fig. 6d).

## Discussion

PD is currently diagnosed based on motor impairments that are only present during progressive stages of the disease^42,43,44^. These clinical symptoms are caused by Lewy bodies aggregating in the brain and by degeneration of dopaminergic neurons in the substantia nigra. Up to 60–70% of these neurons are already lost when the first motor symptoms appear^45^. The gold-standard treatment for managing these motor symptoms in advanced PD is oral administration of the dopamine precursor levodopa (L-dopa). Although short-term oral use of L-dopa is highly effective, long-term use is associated with several complications^46^. Thus, early diagnosis of PD combined with an efficient prophylaxis or therapy that delays or even prevents PD is essential. However, early diagnosis is challenging because biopsies cannot be easily obtained from the brain to monitor the progression of the disease over time. Early pathological signs of PD have been reported in the ENS of patients, suggesting that neurodegeneration may start in the gut and spreads to the brain^10^. Based on this hypothesis, pathological analysis of the GI tract may detect PD early enough to apply preventative therapies. In addition to motor impairments, bowel disturbances have been described in PD patients before clinical symptoms develop. A major finding of the present study is decelerated GI motility in early PD, before motoric disturbances are detected. Consistent with these findings, PD patients with chronic constipation typically exhibit with infrequent bowel movements, impaired propulsive colonic motility, and prolonged colonic transit with reduced rectal contractions^47,48^. Impaired colonic transit has also been described in 4- to 7-month-old α-synuclein-overexpressing animals^49,50^, but has not yet been investigated in very young pre-symptomatic PD mice with an age of 2 months. Additionally, the LI is more affected by Lewy bodies in the myenteric neurons than the SI is^51,52^. These observations support our finding of a stronger pathological effect in the LI than in the SI.

To provide evidence that the gut and ENS are involved in PD onset, and to confirm that PD pathogenesis starts in the gut, we used the A30P mouse PD model, which allows an investigation of pre-symptomatic stages unlike in human patients. Defined pre-symptomatic (ps)A30P mice showed normal gait analysis results but had striking functional alterations in the gut. We investigated whether these functional changes correspond to molecular or morphological abnormalities in the GI tract at this stage. Changes in protein and miRNA expression in body fluid samples and post mortem brain biopsies from PD patients have improved our understanding of PD pathology. PD-related proteins are involved in several pathways including protein ubiquitination, oxidative stress, cytoskeleton development, synaptic function and vesicle release, and calcium binding^9^. Lowe et al. discovered that the majority of Lewy bodies in the brain contain ubiquitin^53^, which conjugates to other proteins to bring about protein degradation^54, 55^. In line with this, post mortem studies have shown defective proteasome activity in PD brains^56^ that involve alterations in the ubiquitin-proteasome pathway^57,58^. To the best of our knowledge, dysregulation of proteins that regulate ubiquitination has not been investigated in the ENS during early stages of PD. In this study, we found that the expression of proteins, responsible for protein folding and ubiquitination, was altered in the MP of the SI and the LI. Of note, these proteins were all involved in regulating the ubiquitin-proteasome pathway^59^.

Oxidative stress reflects the imbalance between the production and detoxification of reactive species, involving oxygen species and nitrogen species. At the cellular level, thiol redox homeostasis is maintained by the thioredoxin/peroxiredoxin (Trx/Prdx) pathway or by glutathione peroxidase (Gpx), which is part of the glutathione (Gsh) pathway. A marker for oxidative stress in PD postmortem brains is an increased level of hydroxyl radicals, which reduces expression of neuroprotective proteins like Trx1, glutathione-S-reductase (Gsr), and Gpx^60^. Moreover, reduced expression of Prdx in the brain has been linked to neurodegenerative diseases including Alzheimer’s disease and PD^61,62^. We found evidence of oxidative stress in the gut during early stages of PD in the present study. Expression of neuroprotective proteins was significantly reduced in the MP of psA30P mice. These changes coincided with significantly increased expression of several miRNAs, which are known to regulate these proteins. These findings provide the first evidence that oxidative stress markers are strikingly dysregulated in the gut during early PD, and that these proteins may represent markers of early PD.

Cytoskeletal reorganization is required for neuronal homeostasis and plasticity, and for responses to axon injury and degeneration^63^. Disrupted post-translational modification of neurofilament proteins have been linked to neurodegenerative disorders, including PD^64,65^. Downregulation of neurofilament mRNA and proteins is common in the brain of PD patients and PD animal models^66,67^. In agreement with these observations, we found significantly reduced Nefl protein expression in the ENS. In addition, we observed significant upregulation of miRNAs that regulate Nefl^41,68^. Globular and filamentous actins are highly enriched in dendritic spines and are regulated during synaptic plasticity by actin-binding proteins (ABPs)^69^. ABPs, such as profilin and the actin-related-protein 2/3 complex (Arpc4), control actin polymerization and are dysregulated in neurological brain disorders including PD^70,71,72, 73^. Here, we showed that expression of profilin and Arpc4 is markedly changed in the LI of psA30P mice, indicating neurofilament disruption in the gut during early stages of PD. Interestingly, profilin is known to cluster with several miRNAs, e.g. miR-130a-3p and 132-3p, which were strongly upregulated in our psA30P model. Consistent with these observations, profilin expression was reduced in the MP of psA30P mice. In line with our findings, miR-132-3p expression was increased in the striatum of a symptomatic rotenone-induced PD rat model^73^. We present evidence that cytoskeletal disruption also contributes to neurodegeneration in the ENS during early stages of PD.

Dynactin is, as a part of the dynactin-dynein-complex (DDC) responsible for axonal transport, the passage of cytoplasmic vesicles^74^ and it affects α-synuclein aggregation increases Lewy body formation in different brain regions^75^. In this study, we found reduced dynactin 2 expression in the MP of psA30P mice, suggesting a reduced axonal transport of motor proteins in the gut at early stages of PD. This assumption is strengthened by upregulations of the counter-rotating miRNAs, which cluster to the DDC and regulate the same pathways^76,41^.

Different tau isoforms, like Mapt, are expressed in the CNS and ENS, but their expression is not altered in the ENS of PD patients with clinical symptoms^77,78,79^. Hallmark characteristics of prion diseases like PD are reduced synapse numbers and reduced dendritic spine densities. Boese et al. described a strong enrichment of several miRNAs including miR-136-5p, which targets Mapt, during the pre-clinical phase of prion disease^80^. Relevant to this, we report for the first time that Mapt is highly downregulated in the colonic MP during the pre-symptomatic stage of PD. This downregulation correlated with an increased expression of miR-22-3p and miR-136-5p. Very recently, deletion of Map6, which interacts with Mapt^81^ and was markedly decreased in our model, was reported to cause muscle weakness and atrophy, reduced calcium release, and alterations in the microtubule network, mirroring the early motoric impairments observed in the gut of α-synuclein-overexpressing mice^82,83^. Interestingly, α-synuclein interacts with microtubules in HeLa cells^84^ and preferably with tyrosinated tubulin in human mesencephalic neurons^85^. In agreement with our *in vitro* results, A30P α-synucleins regulate the microtubule network in PD. These mutated α-synucleins can affect neuronal microtubule assembly, and their overexpression leads to microtubule dysfunction and nigral neurite degeneration^86,87^. Our data imply that A30P α-synuclein pathology also impacts the cytoskeleton of ENS neurons during early PD.

Synaptic transmission is driven by Ca^2+^-dependent vesicle soluble N-ethylmaleimide-sensitive factor attachment (SNAREs), which allow synaptic vesicles to fuse to the plasma membrane^88^. Ca^2+^-binding synaptotagmins (Syt) interact with the SNARE proteins Vamp2, synataxin1 (Stx1), and synaptosomal-associated protein 25 (Snap25) for exocytosis^89^. Native α-synuclein, which is a neuronal presynaptic protein, is involved in the trafficking of synaptic vesicles and in vesicle exocytosis^90,91^. However, aggregated α-synuclein is cytotoxic and mediates synaptic dysfunction by preventing the transport of synaptic proteins or the re-clustering of synaptic vesicles, resulting in reduced neurotransmitter release^92,93^. Misfolded α-synuclein co-localizes with Vamp2 and promotes SNARE-complex assembly^94,95^, while α-synuclein triple-knockout mice show neurological damage and reduced SNARE assembly^96^. In our study, Stx1, Syt1, and Vamp2, which are all involved in the synaptic vesicle cycle^97^, were significantly downregulated in the gut of psA30P mice. α-synuclein aggregation can cause neurotoxicity of cortical neurons through sequestration of Vamp2^95^, leading to neuronal loss and reduced synaptic transmission^98,99,100^. This may explain the gut pathology observed in our psA30P mice supported by our findings that miRNAs clustering with Vamp2 are upregulated. This theory is also supported by the results of our *in vitro* experiments. Thus, the reduced expression of synaptic vesicle proteins, together with increased expression of the miRNAs that regulate them, may contribute to neuronal cell death in pre-symptomatic PD.

In addition to its important role in synaptic transmission, calcium also plays a pivotal role in neurodegeneration^101^. Intracellular Ca^2+^-dependent pathways dramatically increase calcium levels, which can trigger apoptotic cascades. Calcium-binding proteins (CaBPs) maintain calcium homeostasis for neuronal survival^102,103^. CaBPs like calbindin-D (CalbD) and Calb2 are widely expressed in the CNS^104,105^ and ENS^106,107,108^. Different neuronal populations have variable vulnerability to degeneration. Lack of CalbD in dopaminergic neurons increases susceptibility to PD; this is further supported by the low percentage of CalbD-positive neurons in advanced stages of PD^109^. Calb2 is expressed close to the plasma membrane and calcium channels^110^, so may be involved in calcium influx, neuronal excitability, and neurotransmitter release^111,112^. Moreover, Calb2 has been associated with other neurological disorders including schizophrenia and autism^113,114,115^. Here, Calb2 positive neurons were increased after acute α- synuclein exposure suggesting a temporary higher resistance of these neurons. However, a chronic exposure reduced Calb2 neurons and implicated an increase of miR-17-5p that regulates Calb2. It has been reported, that an upregulation of miR-17-5p inhibits cell proliferation and induces apoptosis, while and inhibition induces neurite outgrowth^116^. Based on these findings, we postulate that Calb2 may initiate PD development in the gut. Our hypothesis is supported by previous findings that Calb2 expression is reduced in neurons following 6-OHDA treatment in a PD rat model^117,113^.

Markers like PINK1, Parkin, DJ1, and LRRK2 are dysregulated in postmortem brains and liquid biopsies of PD patients^118,119,120,121^. Hence, these are rather more suitable markers for later stages of the disease in the CNS. In the present study we identified a whole panel of gut-related biomarkers that may detect PD during early stages of the disease, including Nefl, Vamp2, and Calb2, and their regulating miRNAs. These proteins and miRNAs are mainly involved in cytoskeleton assembly, oxidative stress, ubiquitin-proteasome degradation, synaptic transmission, and calcium signaling, and may contribute to intestinal dysfunction during PD. In the future, the ENS may serve as a source for minimally invasive gut biopsies, which can be taken easily from human patients. Therefore, our novel biomarkers may open up the possibility of screening gut tissue to diagnose PD during its early stages, which may facilitate the timely treatment or even prevention of PD.

## Online methods

### Animals

α-synuclein-overexpressing transgenic A30P mice (Th-1-SNCA-A30P^122^), based on a genetic C57B6/J background and age- and sex-matched wild type (WT) mice (postnatal day 2, adult 2 months and 12–13 months old) were used in experiments. Two-month-old A30P mice were defined as pre-symptomatic (ps)A30P mice and were used in expression profiling experiments. Animals were housed under specific pathogen-free conditions on a 12 h light/12 h dark cycle according to German regulations. For tissue dissections, adult mice were deeply anesthetized with isoflurane (Piramal Critical Care) and sacrificed by inhalation of an overdose. Newborn mice were killed by decapitation. Animals were dissected according to the guidelines of the local ethics committee and in accordance with the animal protection laws in Rhineland-Palatinate, Germany.

### Gait analysis

Gait was assessed in adult A30P and WT mice (2 months and 12–13 months old) using the Noldus CatWalk XT system. The system consists of a 70-cm long green illuminated glass runway. Above this corridor is a lid with red LED lights to create a silhouette of the mouse running in the corridor. Below the walkway, a high-speed camera records the scattered light from the paw prints, which is digitized and analyzed by CatWalk XT software. The calibrated walkway was set to a 10 × 20 cm area. Before the behavior test, mice were weighed. Mice were allowed to walk in an unforced manner on the glass plate at least five times in each experiment after a minimum training period of five runs the day before the experimental run. Mice that failed the training were excluded from the study. Footprints were classified automatically as right forepaw, right hindpaw, left forepaw, and left hindpaw. For each experiment, correct paw labels and footfall patterns were controlled, followed by an automated analysis of wide-range parameters involving spatial and temporal characteristics. After the gait analysis, the small intestine (SI) and the large intestine (LI) of 2-month-old and 12-13-month-old A30P and WT mice were removed and the gut lengths were measured.

### GI motility recordings

Video recording with spatiotemporal mapping is a reliable technique for monitoring and displaying motility of gut segments. To evaluate GI motility in psA30P and WT mice, a luminal perfusion setup was used as previously described by Schreiber et al. 2014^123^. In brief, a 3.5-cm gut segment of the ileum and proximal LI were dissected and the mesentery was removed. Then, the tissue was fixed on Luer locks in an organ bath filled with tyrode buffer (130 mM NaCl, 24.2 mM NaHCO_3_, 11 mM glucose, 4.5 mM KCl, 2.2 mM CaCl_2_, 1.2 mM NaH_2_PO_4_, 0.6 mM MgCl_2_), gassed with carbogen to reach a stable physiological pH 7.4. The perfusion bath was kept at 37°C using a heat exchanger. Intestinal segments were luminally perfused at a flow rate of 0.2 ml/min while the height of the luminal efflux tubing relative to the water level (3 cm) was set to a corresponding luminal pressure. All preparations were allowed to equilibrate for 10 min followed by a 10 min spontaneous motility period.

GI motility was assessed by video recordings at 25 frames/s. Videos were analyzed and quantified by MotMap (www.smoothmap.org) and a custom written LabVIEW program (LabVIEW 2019, National Instruments). The video analysis software MotMap was used to set 30 imaginary dots along a 2 cm gut segment on the lower and upper border of the gut as seen in Fig. 2a. The up and down movements of these dot pairs were tracked automatically to calculate the positions to each other for every frame, representing the amplitude of gut activity (Δ). Before the analysis, the program automatically applies a low frequency baseline correction to each of the 30 sections to compensate for slow gut movements. To validate amplitude changes, all values were normalized to their initial diameter. The motility data represent dilatations as positive values vs. the average diameter and contractions as negative values. Motility was converted to high-resolution spatiotemporal heatmaps, while all values were specified in a color range using the custom written LabVIEW software. Gut contractions were counted using spatiotemporal heatmaps. The mean interval was defined as the duration of one contraction and one dilatation (Fig. 2a) and indicates the corresponding frequency. The contraction duration was calculated as the time between adjacent leading edge to leading edge zero crossings. To suppress a noise-induced false triggering, a hysteresis threshold was set at the order of 0.05 to 0.1 mm. Each of the 30 dots over the 2 cm were separately analyzed by determining the duration of all full cycles followed by averaging over all detected cycle durations. To determine the velocity of GI motility, slopes of individual contractions were measured in the LabView program by manually aligning a cursor over the wave top, as shown in Fig. 2b. The program calculates the velocity from the start and end point of the cursor. Together, mean intervals, contraction numbers, and contraction velocity provide information about peristaltic activity.

### Tissue preparation

LI fragments from psA30P and WT mice (2 months old and/or newborn) were collected in MEM-Hepes + 100 U/ml pencillin/100 μg/ml streptomycin (P/S) and the mesentery was removed. For full-thickness intestinal wall and whole-muscle layer preparations from adult mice, excrement was washed out with MEM-Hepes + 1% P/S. For muscle layer preparations, the intestine was cut open along the mesenteric line and the outer muscle layer was separated from the mucosa and kept for further experiments. Next, the myenteric plexus (MP) was isolated for whole-mount immunostaining. Brains were removed from adult mice for further experiments.

#### Isolation of the MP

Primary enteric cells, in particular MP from newborn and adult mice, were isolated as previously described^124,125^. In brief, stripped muscle layers were cut into small pieces and digested with 0.375 mg/ml Liberase (Roche) and 0.2 mg/ml DNase (Roche) in Hank’s balanced salt solution. Adult SI was incubated at 37°C for 4 h and adult LI for 4.5 h. Thereafter, purified MP networks were washed two times with 0.01 M phosphate-buffered saline (PBS), frozen in liquid nitrogen, and stored at −80°C for further experiments. Tissue from newborn C57B6/J mice was digested for 2 h and MP cells were dissociated for cell culture experiments by incubation with TrypLE (Gibco) (2 × 6 min at 37°C) and triturated into a single-cell suspension with 23G and 27G needles.

#### Isolation of the mesencephalon

The mesencephalon was also isolated from psA30P and WT mice along with the MP. The brain was separated from the skin and the skull, and placed dorsal side up. The mesencephalon was dissected using a scalpel blade. A first cut was made adjacent to the inferior colliculi, followed by a second cut of approximately 4 mm in a rostral direction. The slice was placed with the rostral side up and the mesencephalon was taken out without including the hippocampus, cortex, or cerebellum. The mesencephalon was snap-frozen with liquid nitrogen and stored at −80°C for further experiments.

#### Whole-mount preparations

Segments of full-thickness intestinal wall and the outer muscle layer of psA30P and WT mice were fixed and stretched flat on a Sylgard plate followed by fixation with 4% paraformaldehyde for 2 h at 4°C. Afterwards, samples were washed three times with PBS for 10 min. Intestinal wall samples were cut into 1 cm pieces, the muscle layers were removed, and samples were stored in 0.1% NaN_3_ in PBS at 4°C for further experiments.

### Protein isolation for mass spectroscopy of the MP

Protein profiles of purified MP from SI and LI tissue obtained from psA30P and WT mice were investigated by mass spectroscopy. Harvested MP networks were thawed and transferred into 0.1 μM PBS, containing 1:100 protease inhibitor (Roche) and nuclease mix (GE Healthcare), and were vortexed. After five freeze-thaw cycles in liquid nitrogen, samples were purified with the 2-D Clean up kit (GE Healthcare). Pelleted proteins were resuspended in 6 M urea, 2M thiourea and 25 mM NH_4_HCO_3_ at pH 8. The protein concentration was measured using the 2-D Quant kit (GE Healthcare). A total protein amount of approximately 16 μg for the colon and 25 μg for the small intestine was determined.

Mass spectrometry analysis was performed on a high-resolution LC-MS system (Eksigent nanoLC 425 coupled to a Triple-TOF 6600, AB Sciex) in information-dependent acquisition (IDA) mode. HPLC separation was performed in trap-elution mode using a Symmetry C18 column (5 μm particles, 0.18 × 20 mm, Waters) for trapping and a self-packed analytical column (75 μm × 150 mm, 3 μm particles ReproSil-PurC18-AQ) for separation. A constant flow of 300 nl/min was used and the gradient was ramped within 108 min from 2% to 33% of HPLC buffer B (buffer A: 2% acetonitrile, 0.1% formic acid; buffer B: 90% acetonitrile, 0.1% formic acid), then within 12 min to 50% buffer B, followed by washing and equilibration steps. The mass spectrometer was run in IDA mode and recorded one survey scan (250 ms, 350–1500 m/z) and fragment spectra (70 ms, 100–1400 m/z) of the 25 most intense parent ions (charge state > 2, intensity > 300 cps, exclusion for 15 s after one occurrence), giving a cycle time of 2 sec. In total, 1,044 different proteins were investigated (Supplementary Table 2). The yielded proteins were then analyzed and visualized by Search Tool for the Retrieval of Interacting Genes/Proteins (STRING) and Ingenuity Pathway Analysis (IPA) software tool (QIAGEN Inc., https://www.qiagenbioinformatics.com/products/ingenuitypathway-analysis).

### Whole-mount immunostaining

Immunostaining and clearing procedures for the full-thickness intestinal wall and muscle layer samples were performed as previously described^126^. In brief, whole-mount muscle layer and intestinal wall preparations of LI specimens from psA30P mice and corresponding WT were permeabilized for 4 h at 37°C on a shaker with permeabilization solution (0.01% NaN_3_ + 1% normal donkey serum + 1% Triton-X-100 in PBS). Then, samples were blocked in blocking solution (0.01% NaN_3_ + 10% normal donkey serum + 0.1% BSA + 1% Triton-X-100 in PBS) for 4 h at room temperature (RT) on a shaker. Subsequently, samples were incubated with primary antibodies (diluted in blocking solution, Supplementary Table 1) at 37°C for 48 h using an orbital shaker. After rinsing four times in PBST (0.05% Tween20 in PBS) for 30 min, tissues were incubated with respective secondary antibodies (1:500 in 0,01% NaN_3_ ^+^ 1% Triton-X-100 in PBS) overnight at 37°C on an orbital shaker (Supplementary Table 1). Again, samples were rinsed with PBST four times for 30 min and then counterstained with DAPI (1:1000) for 2 h at RT. Samples were washed in PBS five times for 10 min at RT. For the clearing procedure, Histodenz (Sigma) was diluted with melted N-methylacetamide (40% in PBS, Sigma) to 86% (w/v) concentration and incubated at 37°C until dissolved. Then 0.1% Triton-X-100 and 0.5% 1-thioglycerol were added to the clearing solution. After being incubated in completed clearing solution for 1–5 min, the outer muscle layers were mounted in fluorescent mounting medium. Detection and image processing were performed with CellObserver Z1 using the ApoTome technology and Axiovision software (Zeiss). Full-thickness samples were incubated in completed clearing solution overnight and imaged on a Leica SP8 confocal microscope (Leica Microsystems CMS GmbH, Germany). For imaging, samples were positioned in a glass-bottomed petri dish, submerged in the clearing solution and covered by a golden ring with nylon mesh to prevent the tissue from floating. 3-D renderings from intestinal wall samples were obtained using the 3-D Viewer of the Leica LAS X software (Leica microsystems, Wetzlar, Germany).

### *In vitro* studies

#### Aggregation of A30P α-synuclein

To create aggregated α-synuclein peptides, 50 μM of A30P α-synuclein (rPeptid) and 50 μM dopamine (Sigma) were diluted in 5 mM Tris-HCl with 100 mM NaCl at pH 7.4 using low protein binding tubes. The aggregation process was enforced by permanent shaking for 3 days at 37°C as previously described^127^.

#### Cell treatment with A30P α-synuclein

Dissociated MP cells from newborn mice were plated on extracellular matrix (ECM) (Sigma)-coated coverslips (40.000 cells/coverslip). Cells were maintained for 7 days in differentiation medium consisting of DMEM/F-12 medium, 1% bovine serum albumin (BSA), 2% B27 with retinoic acid supplement, 0.1% β-mercaptoethanol, P/S and 10 ng/μl human glial cell-derived neurotrophic factor (Immuno Tools) before exposure to A30P α-synuclein. Culture medium was replaced by medium with B27 without antioxidants, containing 0.5 μM A30P α-synuclein in Tris-HCl. After α-synuclein treatment for an additional 5 days, cells were fixed with 4% formaldehyde for 15 min at RT. To determine cell viability after α-synuclein treatment, a live-dead-assay was performed. Cells were incubated with 1 μg/ml calcein and 0.5 μg/ml propidiumiodide in PBS for 15 min at 37°C and analyzed with the CellObserver Z1 and Axiovision software (Zeiss). Total numbers of living and dead cells were calculated per picture section.

#### Immunocytochemistry

Cells were fixed with 4% formaldehyde and permeabilized with 0.1% Triton-X-100 for 10 min, followed by 30 min blocking in normal donkey serum. Primary antibody incubation (Supplementary Table 1) was performed for 1 h at RT in PBS. Samples were washed three times in PBS and further incubated with respective secondary antibodies (1:500, Supplementary Table 1) in 0.01 M PBS for 1 h at RT. Samples were washed three times in PBS and once in distilled water. Cells were then mounted on slides with fluorescent mounting medium with DAPI. Detection and image processing were performed with CellObserver Z1 using the ApoTome technology and Axiovision software (Zeiss).

### nCounter analysis

For miRNA profiling, total RNA (including small RNAs) from the MP of the LI and the mesencephalon samples from psA30P and WT mice was isolated with the miRNeasy Micro Kit (Qiagen). RNA was eluted in 10 μl (50–100 ng/μl) nuclease-free H_2_O. RNA quantity and quality were assessed using the Qubit™ 4 Fluorometer (Qubit RNA HS Kit; Invitrogen) and the NanoDrop 2000 (Thermo Fisher Scientific). Agilent Bioanalyzer 2100 (TotalRNA Nanokit) and miRNA profiling was performed using the nCounter SPRINT system (NanoString Technologies, USA, www.nanostring.com) at the nCounter Core Facility of the University of Heidelberg (Heidelberg, Germany, www.ncounter.uni-hd.de). For miRNA library preparation and subsequent probe hybridization using the mouse miRNA panel 1.5 (Supplementary Table 3), 100 ng of total RNA was applied. Using the standard nCounter miRNA assay protocol, expression profiles of 578 mouse miRNAs derived from miRbase v.22 were determined. Only miRNAs with at least 100 counts were considered as robustly expressed and included in differential expression analysis of psA30P vs. WT tissue. Data were analyzed using the nSolver software version 4.0 (NanoString Technologies). Stably expressed housekeeping genes such as beta-actin (*Actb*), beta-2-microglobulin (*B2m*), glyceraldehyde 3-phosphate dehydrogenase (*Gapdh*), and ribosomal protein L19 (*Rpl19*) were used for data normalization. To investigate a potential impact of the dysregulated miRNAs on target gene translation, the top-regulated miRNAs were subjected to target gene prediction using mirWALK 3.0 integrating the algorithms of multiple miRNA target prediction tools ^41^.

To gain further insight into the function of the most dysregulated miRNAs and their targets, data were put into biological context using the knowledge-based IPA. IPA integrates selected data sets (in our case, miRNA profiles and proteomics) with mining techniques to predict functional connections, protein networks, protein-protein interactions, and related biological functions as well as canonical signaling pathways. miRNA target filter plus miRNA/target = protein pairing analysis were used to identify co-regulated miRNA/target mRNA pairs.

### Statistical analyses

For statistical analysis, GraphPad Prism Software 8 was used. Normality of the sample populations was tested with the Shapiro-Wilk test. For normally distributed data, group differences were analyzed with the Student’s t-test and the associated effect sizes with the Cohen’s test^128^ or two-way ANOVA. Data are displayed as means ± SD and *p*-values ≤ 0.05 (*), ≤ 0.01 (**) and ≤ 0.001 (***) were considered statistically significant.

## Acknowledgements

The authors thank Tobias Fritz for technical support with the CatWalk XT System, which was performed at the Saarland University, Homburg. We thank Salam Warda for analyzing the motility videos, David Grundmann and Alexandra Conrad for the help with the myenteric plexus isolation used for the proteomics. We thank Liz Spycher for technical assistance. This work is partly supported by the Ministry of Education and Research (BMBF, OD Pfalz: 03IHS075A) and a grant from the Deutsche Forschungsgemeinschaft (DFG SCHA/11-1).

## Author contributions

Manuela Gries, Anne Christmann and Karl-Herbert Schäfer conceived the project, designed and supervised the experiments, analyzed and interpreted results and wrote the manuscript. Steven Schulte helped generating and analyzing the immunohistochemical data and with the sample preparations. Maximilian Weyland assisted with heatmap establishment and sample preparations. We thank Marko Baller for statistical support and writing the commercial LabView software. Ralph Röth, Stefanie Schmitteckert, Beate Niesler and Gudrun A. Rappold performed the nCounter expression profile studies and helped with pathway design, analyses and comparisons. Marcus Unger supported in clinical associated PD questions. Monika Martin und Stephanie Rommel assisted with sample preparation. Yang Liu and Markus Britschgi kindly provided the Th-1-SNCA-A30P mice. Michael Schroda, Frederik Sommer and Timo Mühlhaus made the mass spectroscopy experiments. Jean-Pierre Timmermans and Isabel Pintelon generated the three-dimensional images. Hilal Lashuel maintained synuclein related issues. Michael D. Menger and Matthias W. Laschke provided the CatWalk XT System. All authors discussed the results and commented on the manuscript.

## References

1. Hayes, M. W., Fung, V. S., Kimber, T. E. & O’Sullivan, J. D. Current concepts in the management of Parkinson disease. Medical Journal of Australia (2010) doi:10.5694/j.1326-5377.2010.tb03453.x.

2. Postuma, R. B. et al. MDS clinical diagnostic criteria for Parkinson’s disease. Movement Disorders (2015) doi:10.1002/mds.26424.

3. Tysnes, O. B. & Storstein, A. Epidemiology of Parkinson’s disease. Journal of Neural Transmission (2017) doi:10.1007/s00702-017-1686-y.

4. Savica, R. et al. Medical records documentation of constipation preceding Parkinson disease: A case-control study. Neurology (2009) doi:10.1212/WNL.0b013e3181c34af5.

5. Cersosimo, M. G. et al. Gastrointestinal manifestations in Parkinson’s disease: Prevalence and occurrence before motor symptoms. J. Neurol. (2013) doi:10.1007/s00415-012-6801-2.

6. Radicati, F. G. et al. Non motor symptoms in progressive supranuclear palsy: prevalence and severity. npj Park. Dis. (2017) doi:10.1038/s41531-017-0037-x.

7. Spillantini, M. G. et al. $α$-synuclein in Lewy bodies [8]. Nature (1997) doi:10.1038/42166.

8. Lees, A. J., Hardy, J. & Revesz, T. Parkinson’s disease. The Lancet (2009) doi:10.1016/S0140-6736(09)60492-X.

9. Hernandez, D. G., Reed, X. & Singleton, A. B. Genetics in Parkinson disease: Mendelian versus non-Mendelian inheritance. Journal of Neurochemistry (2016) doi:10.1111/jnc.13593.

10. Braak, H., De Vos, R. A. I., Bohl, J. & Del Tredici, K. Gastric $α$-synuclein immunoreactive inclusions in Meissner’s and Auerbach’s plexuses in cases staged for Parkinson’s disease-related brain pathology. Neurosci. Lett. (2006) doi:10.1016/j.neulet.2005.11.012.

11. Lebouvier, T. et al. Pathological lesions in colonic biopsies during Parkinson’s disease. Gut (2008) doi:10.1136/gut.2008.162503.

12. Del Tredici, K., Rüb, U., De Vos, R. A. I., Bohl, J. R. E. & Braak, H. Where does Parkinson disease pathology begin in the brain? J. Neuropathol. Exp. Neurol. (2002) doi:10.1093/jnen/61.5.413.

13. Goedert, M., Spillantini, M. G., Del Tredici, K. & Braak, H. 100 years of Lewy pathology. Nature Reviews Neurology (2013) doi:10.1038/nrneurol.2012.242.

14. Mukherjee, A., Biswas, A. & Das, S. K. Gut dysfunction in Parkinson’s disease. World Journal of Gastroenterology (2016) doi:10.3748/wjg.v22.i25.5742.

15. Perez-Pardo, P. et al. The gut-brain axis in Parkinson’s disease: Possibilities for food-based therapies. Eur. J. Pharmacol. (2017) doi:10.1016/j.ejphar.2017.05.042.

16. Santos, S. F., De Oliveira, H. L., Yamada, E. S., Neves, B. C. & Pereira, A. The gut and Parkinson’s disease - A bidirectional pathway. Frontiers in Neurology (2019) doi:10.3389/fneur.2019.00574.

17. Holmqvist, S. et al. Direct evidence of Parkinson pathology spread from the gastrointestinal tract to the brain in rats. Acta Neuropathol. (2014) doi:10.1007/s00401-014-1343-6.

18. Uemura, N. et al. Inoculation of α-synuclein preformed fibrils into the mouse gastrointestinal tract induces Lewy body-like aggregates in the brainstem via the vagus nerve. Mol. Neurodegener. (2018) doi:10.1186/s13024-018-0257-5.

19. Challis, C. et al. Gut-seeded α-synuclein fibrils promote gut dysfunction and brain pathology specifically in aged mice. Nat. Neurosci. (2020) doi:10.1038/s41593-020-0589-7.

20. Pan-Montojo, F. et al. Progression of Parkinson’s disease pathology is reproduced by intragastric administration of rotenone in mice. PLoS One (2010) doi:10.1371/journal.pone.0008762.

21. Scheperjans, F. et al. Gut microbiota are related to Parkinson’s disease and clinical phenotype. Mov. Disord. (2015) doi:10.1002/mds.26069.

22. Unger, M. M. et al. Short chain fatty acids and gut microbiota differ between patients with Parkinson’s disease and age-matched controls. Park. Relat. Disord. (2016) doi:10.1016/j.parkreldis.2016.08.019.

23. Weis, S. et al. Effect of Parkinson’s disease and related medications on the composition of the fecal bacterial microbiota. npj Park. Dis. (2019) doi:10.1038/s41531-019-0100-x.

24. Devos, D. et al. Colonic inflammation in Parkinson’s disease. Neurobiol. Dis. (2013) doi:10.1016/j.nbd.2012.09.007.

25. Endres, K. & Schäfer, K. H. Influence of Commensal Microbiota on the Enteric Nervous System and Its Role in Neurodegenerative Diseases. J. Innate Immun. (2018) doi:10.1159/000488629.

26. Dutta, S. K. et al. Parkinson’s disease: The emerging role of gut dysbiosis, antibiotics, probiotics, and fecal microbiota transplantation. Journal of Neurogastroenterology and Motility (2019) doi:10.5056/jnm19044.

27. Seguella, L., Sarnelli, G. & Esposito, G. Leaky gut, dysbiosis, and enteric glia activation: The trilogy behind the intestinal origin of Parkinson’s disease. Neural Regeneration Research (2020) doi:10.4103/1673-5374.270308.

28. Mi, S., Zhang, J., Zhang, W. & Huang, R. S. Circulating MicroRNAs as Biomarkers for Inflammatory Diseases. MicroRNA (2013) doi:10.2174/2211536611302010007.

29. Zibert, J. R. et al. MicroRNAs and potential target interactions in psoriasis. J. Dermatol. Sci. (2010) doi:10.1016/j.jdermsci.2010.03.004.

30. Mitchell, P. S. et al. Circulating microRNAs as stable blood-based markers for cancer detection. Proc. Natl. Acad. Sci. U. S. A. (2008) doi:10.1073/pnas.0804549105.

31. Gwiggner, M. et al. MicroRNA-31 and MicroRNA-155 are overexpressed in ulcerative colitis and regulate IL-13 signaling by targeting interleukin 13 receptor $α$-1. Genes (Basel). (2018) doi:10.3390/genes9020085.

32. Chen, X. et al. MicroRNAs tend to synergistically control expression of genes encoding extensively-expressed proteins in humans. PeerJ (2017) doi:10.7717/peerj.3682.

33. Liu, J. et al. Combination of plasma microRNAs with serum CA19-9 for early detection of pancreatic cancer. Int. J. Cancer (2012) doi:10.1002/ijc.26422.

34. Zhao, H. et al. A pilot study of circulating miRNAs as potential biomarkers of early stage breast cancer. PLoS One (2010) doi:10.1371/journal.pone.0013735.

35. Junn, E. & Mouradian, M. M. MicroRNAs in neurodegenerative diseases and their therapeutic potential. Pharmacology and Therapeutics (2012) doi:10.1016/j.pharmthera.2011.10.002.

36. De Guire, V. et al. Circulating miRNAs as sensitive and specific biomarkers for the diagnosis and monitoring of human diseases: Promises and challenges. Clinical Biochemistry (2013) doi:10.1016/j.clinbiochem.2013.03.015.

37. Margis, R., Margis, R. & Rieder, C. R. M. Identification of blood microRNAs associated to Parkinsonós disease. J. Biotechnol. (2011) doi:10.1016/j.jbiotec.2011.01.023.

38. Mushtaq, G. et al. miRNAs as Circulating Biomarkers for Alzheimer’s Disease and Parkinson’s Disease. Med. Chem. (Los. Angeles). (2016) doi:10.2174/1573406411666151030112140.

39. Chen, L. et al. Identification of aberrant circulating miRNAs in Parkinson’s disease plasma samples. Brain Behav. (2018) doi:10.1002/brb3.941.

40. Pistacchi, M. et al. Gait analysis and clinical correlations in early Parkinson’s disease. Funct. Neurol. (2017) doi:10.11138/FNeur/2017.32.1.028.

41. Sticht, C., De La Torre, C., Parveen, A. & Gretz, N. Mirwalk: An online resource for prediction of microrna binding sites. PLoS One (2018) doi:10.1371/journal.pone.0206239.

42. Reichmann, H. Clinical criteria for the diagnosis of Parkinson’s disease. Neurodegenerative Diseases (2010) doi:10.1159/000314478.

43. Warlop, T. et al. Temporal organization of stride duration variability as a marker of gait instability in Parkinson’s disease. J. Rehabil. Med. (2016) doi:10.2340/16501977-2158.

44. Brognara, L., Palumbo, P., Grimm, B. & Palmerini, L. Assessing Gait in Parkinson’s Disease Using Wearable Motion Sensors: A Systematic Review. Diseases (2019) doi:10.3390/diseases7010018.

45. Lang, A. E. & Lozano, A. M. Parkinson’s disease. First of two parts. N. Engl. J. Med. (1998) doi:10.1056/NEJM199810083391506.

46. Mulroy, E. & Bhatia, K. P. The Gut Microbiome: A Therapeutically Targetable Site of Peripheral Levodopa Metabolism. Mov. Disord. Clin. Pract. (2019) doi:10.1002/mdc3.12828.

47. Fasano, A., Visanji, N. P., Liu, L. W. C., Lang, A. E. & Pfeiffer, R. F. Gastrointestinal dysfunction in Parkinson’s disease. The Lancet Neurology (2015) doi:10.1016/S1474-4422(15)00007-1.

48. Knudsen, K. et al. Gastrointestinal Transit Time in Parkinson’s Disease Using a Magnetic Tracking System. J. Parkinsons. Dis. (2017) doi:10.3233/JPD-171131.

49. Drolet, R. E., Cannon, J. R., Montero, L. & Greenamyre, J. T. Chronic rotenone exposure reproduces Parkinson’s disease gastrointestinal neuropathology. Neurobiol. Dis. (2009) doi:10.1016/j.nbd.2009.06.017.

50. Kuo, Y.-M., Nwankwo, E. I., Nussbaum, R. L., Rogers, J. & Maccecchini, M. L. Translational inhibition of $α$-synuclein by Posiphen normalizes distal colon motility in transgenic Parkinson mice. Am. J. Neurodegener. Dis. (2019).

51. Pouclet, H., Lebouvier, T., Coron, E., Neunlist, M. & Derkinderen, P. Lewy pathology in gastric and duodenal biopsies in Parkinson’s Disease. Movement Disorders (2012) doi:10.1002/mds.24993.

52. Wakabayashi, K., Takahashi, H., Ohama, E. & Ikuta, F. Parkinson’s disease: an immunohistochemical study of Lewy body-containing neurons in the enteric nervous system. Acta Neuropathol. (1990) doi:10.1007/BF00294234.

53. Lowe, J. et al. Ubiquitin is a common factor in intermediate filament inclusion bodies of diverse type in man, including those of Parkinson’s disease, Pick’s disease, and Alzheimer’s disease, as well as Rosenthal fibres in cerebellar astrocytomas, cytoplasmic bodies in m. J. Pathol. (1988) doi:10.1002/path.1711550105.

54. Hershko, A., Heller, H., Elias, S. & Ciechanover, A. Components of ubiquitin-protein ligase system. Resolution, affinity purification, and role in protein breakdown. J. Biol. Chem. (1983).

55. Walden, H. & Muqit, M. M. K. Ubiquitin and Parkinson’s disease through the looking glass of genetics. Biochemical Journal (2017) doi:10.1042/BCJ20160498.

56. Furukawa, Y. et al. Brain proteasomal function in sporadic Parkinson’s disease and related disorders. Ann. Neurol. (2002) doi:10.1002/ana.10207.

57. Grünblatt, E. et al. Gene expression profiling of parkinsonian substantia nigra pars compacta; alterations in ubiquitin-proteasome, heat shock protein, iron and oxidative stress regulated proteins, cell adhesion/cellular matrix and vesicle trafficking genes. J. Neural Transm. (2004) doi:10.1007/s00702-004-0212-1.

58. Lim, K. L. & Tan, J. M. M. Role of the ubiquitin proteasome system in Parkinson’s disease. BMC Biochemistry (2007) doi:10.1186/1471-2091-8-S1-S13.

59. Lehtonen, Š., Sonninen, T. M., Wojciechowski, S., Goldsteins, G. & Koistinaho, J. Dysfunction of cellular proteostasis in Parkinson’s disease. Frontiers in Neuroscience (2019) doi:10.3389/fnins.2019.00457.

60. Sian, J. et al. Alterations in glutathione levels in Parkinson’s disease and other neurodegenerative disorders affecting basal ganglia. Ann. Neurol. (1994) doi:10.1002/ana.410360305.

61. Parmigiani, R. B. et al. HDAC6 is a specific deacetylase of peroxiredoxins and is involved in redox regulation. Proc. Natl. Acad. Sci. U. S. A. (2008) doi:10.1073/pnas.0803749105.

62. Jian, W. et al. Inhibition of HDAC6 increases acetylation of peroxiredoxin1/2 and ameliorates 6-OHDA induced dopaminergic injury. Neurosci. Lett. (2017) doi:10.1016/j.neulet.2017.08.029.

63. Muñoz-Lasso, D. C., Romá-Mateo, C., Pallardó, F. V & Gonzalez-Cabo, P. Much More Than a Scaffold: Cytoskeletal Proteins in Neurological Disorders. Cells (2020) doi:10.3390/cells9020358.

64. Conde, M. A. et al. Phospholipase D1 downregulation by α-synuclein: Implications for neurodegeneration in Parkinson’s disease. Biochim. Biophys. Acta - Mol. Cell Biol. Lipids (2018) doi:10.1016/j.bbalip.2018.03.006.

65. Shukla, V., Skuntz, S. & Pant, H. C. Deregulated Cdk5 Activity Is Involved in Inducing Alzheimer’s Disease. Archives of Medical Research (2012) doi:10.1016/j.arcmed.2012.10.015.

66. Hill, W. D., Arai, M., Cohen, J. A. & Trojanowski, J. Q. Neurofilament mRNA is reduced in Parkinson’s disease substantia nigra pars compacta neurons. J. Comp. Neurol. (1993) doi:10.1002/cne.903290304.

67. Wellings, T. P., Brichta, A. M. & Lim, R. Altered neurofilament protein expression in the lateral vestibular nucleus in Parkinson’s disease. Exp. Brain Res. (2017) doi:10.1007/s00221-017-5092-3.

68. Krämer, A., Green, J., Pollard, J. & Tugendreich, S. Causal analysis approaches in ingenuity pathway analysis. Bioinformatics (2014) doi:10.1093/bioinformatics/btt703.

69. Welch, M. D., Iwamatsu, A. & Mitchison, T. J. Actin polymerization is induced by Arp2/3 protein complex at the surface of Listeria monocytogenes. Nature (1997) doi:10.1038/385265a0.

70. Bamburg, J. R. & Bernstein, B. W. Actin dynamics and cofilin-actin rods in alzheimer disease. Cytoskeleton (2016) doi:10.1002/cm.21282.

71. Yan, Z., Kim, E., Datta, D., Lewis, D. A. & Soderling, S. H. Synaptic actin dysregulation, a convergent mechanism of mental disorders? in Journal of Neuroscience (2016). doi:10.1523/JNEUROSCI.2360-16.2016.

72. Datta, D., Arion, D., Roman, K. M., Volk, D. W. & Lewis, D. A. Altered expression of ARP2/3 complex signaling pathway genes in prefrontal layer 3 pyramidal cells in schizophrenia. Am. J. Psychiatry (2017) doi:10.1176/appi.ajp.2016.16020204.

73. Horst, C. H. et al. Signature of Aberrantly Expressed microRNAs in the Striatum of Rotenone-Induced Parkinsonian Rats. Neurochem. Res. (2018) doi:10.1007/s11064-018-2638-0.

74. Urnavicius, L. et al. The structure of the dynactin complex and its interaction with dynein. Science (80-.). (2015) doi:10.1126/science.aaa4080.

75. Shen, C. et al. Dynactin is involved in Lewy body pathology. Neuropathology (2018) doi:10.1111/neup.12512.

76. Dweep, H., Sticht, C., Pandey, P. & Gretz, N. MiRWalk - Database: Prediction of possible miRNA binding sites by ‘ walking’ the genes of three genomes. J. Biomed. Inform. (2011) doi:10.1016/j.jbi.2011.05.002.

77. Arai T, A. et al. Distinct isoforms of tau aggregated in neurons and glial cells in brains of patients with Pick’s disease, corticobasal degeneration and progressive supranuclear palsy. Acta Neuropathol. (2001) doi:10.1007/s004010000283.

78. Armstrong, R. A. & Cairns, N. J. Spatial patterns of the tau pathology in progressive supranuclear palsy. Neurol. Sci. (2013) doi:10.1007/s10072-012-1006-0.

79. Kovacs, G. G. et al. Evaluating the patterns of aging-related tau astrogliopathy unravels novel insights into brain aging and neurodegenerative diseases. J. Neuropathol. Exp. Neurol. (2017) doi:10.1093/jnen/nlx007.

80. Boese, A. S. et al. MicroRNA abundance is altered in synaptoneurosomes during prion disease. Mol. Cell. Neurosci. (2016) doi:10.1016/j.mcn.2015.12.001.

81. Liu, W. & Wang, X. Prediction of functional microRNA targets by integrative modeling of microRNA binding and target expression data. Genome Biol. (2019) doi:10.1186/s13059-019-1629-z.

82. Kuo, Y. M. et al. Extensive enteric nervous system abnormalities in mice transgenic for artificial chromosomes containing Parkinson disease-associated $α$-synuclein gene mutations precede central nervous system changes. Hum. Mol. Genet. (2010) doi:10.1093/hmg/ddq038.

83. Sébastien, M. et al. Deletion of the microtubule-associated protein 6 (MAP6) results in skeletal muscle dysfunction. Skelet. Muscle (2018) doi:10.1186/s13395-018-0176-8.

84. Zhou, R. M. et al. Molecular interaction of α-synuclein with tubulin influences on the polymerization of microtubule in vitro and structure of microtubule in cells. Mol. Biol. Rep. (2010) doi:10.1007/s11033-009-9899-2.

85. Cartelli, D. et al. α-Synuclein is a Novel Microtubule Dynamase. Sci. Rep. (2016) doi:10.1038/srep33289.

86. Lee, H. J., Khoshaghideh, F., Lee, S. & Lee, S. J. Impairment of microtubule-dependent trafficking by overexpression of α-synuclein. Eur. J. Neurosci. (2006) doi:10.1111/j.1460-9568.2006.05210.x.

87. Calogero, A. M., Mazzetti, S., Pezzoli, G. & Cappelletti, G. Neuronal microtubules and proteins linked to Parkinson’s disease: A relevant interaction? Biological Chemistry (2019) doi:10.1515/hsz-2019-0142.

88. McNew, J. A. et al. Compartmental specificity of cellular membrane fusion encoded in SNARE proteins. Nature (2000) doi:10.1038/35025000.

89. Littleton, J. T. et al. Temperature-sensitive paralytic mutations demonstrate that synaptic exocytosis requires SNARE complex assembly and disassembly. Neuron (1998) doi:10.1016/S0896-6273(00)80549-8.

90. Murphy, D. D., Rueter, S. M., Trojanowski, J. Q. & Lee, V. M. Y. Synucleins are developmentally expressed, and α-synuclein regulates the size of the presynaptic vesicular pool in primary hippocampal neurons. J. Neurosci. (2000) doi:10.1523/jneurosci.20-09-03214.2000.

91. Vargas, K. J. et al. Synucleins regulate the kinetics of synaptic vesicle endocytosis. J. Neurosci. (2014) doi:10.1523/JNEUROSCI.4787-13.2014.

92. Scott, D. A. et al. A pathologic cascade leading to synaptic dysfunction in α-synuclein-induced neurodegeneration. J. Neurosci. (2010) doi:10.1523/JNEUROSCI.1091-10.2010.

93. Nemani, V. M. et al. Increased Expression of α-Synuclein Reduces Neurotransmitter Release by Inhibiting Synaptic Vesicle Reclustering after Endocytosis. Neuron (2010) doi:10.1016/j.neuron.2009.12.023.

94. Burré, J. et al. $α$-Synuclein promotes SNARE-complex assembly in vivo and in vitro. Science (80-.). (2010) doi:10.1126/science.1195227.

95. Choi, M. G. et al. Sequestration of synaptic proteins by alphasynuclein aggregates leading to neurotoxicity is inhibited by small peptide. PLoS One (2018) doi:10.1371/journal.pone.0195339.

96. Diao, J. et al. Native α-synuclein induces clustering of synaptic-vesicle mimics via binding to phospholipids and synaptobrevin-2/VAMP2. Elife (2013) doi:10.7554/eLife.00592.

97. Szklarczyk, D. et al. STRING v11: Protein-protein association networks with increased coverage, supporting functional discovery in genome-wide experimental datasets. Nucleic Acids Res. (2019) doi:10.1093/nar/gky1131.

98. Nemani, V. M. et al. Increased Expression of $α$-Synuclein Reduces Neurotransmitter Release by Inhibiting Synaptic Vesicle Reclustering after Endocytosis. Neuron (2010) doi:10.1016/j.neuron.2009.12.023.

99. Haberman, A. et al. The synaptic vesicle SNARE neuronal synaptobrevin promotes endolysosomal degradation and prevents neurodegeneration. J. Cell Biol. (2012) doi:10.1083/jcb.201108088.

100. Soukup, S., Vanhauwaert, R. & Verstreken, P. Parkinson’s disease: convergence on synaptic homeostasis. EMBO J. (2018) doi:10.15252/embj.201898960.

101. Fairless, R., Williams, S. K. & Diem, R. Dysfunction of neuronal calcium signalling in neuroinflammation and neurodegeneration. Cell and Tissue Research (2014) doi:10.1007/s00441-013-1758-8.

102. Persechini, A., Moncrief, N. D. & Kretsinger, R. H. The EF-hand family of calcium-modulated proteins. Trends in Neurosciences (1989) doi:10.1016/0166-2236(89)90097-0.

103. Kennedy, M. B. Regulation of neuronal function by calcium. Trends Neurosci. (1989) doi:10.1016/0166-2236(89)90089-1.

104. Antal, M., Freund, T. F. & Polgár, E. Calcium-binding proteins, parvalbumin- and calbindin-D 28k-immunoreactive neurons in the rat spinal cord and dorsal root ganglia: A light and electron microscopic study. J. Comp. Neurol. (1990) doi:10.1002/cne.902950310.

105. Foo, K. S., Hellysaz, A. & Broberger, C. Expression and colocalization patterns of calbindin-D28k, calretinin and parvalbumin in the rat hypothalamic arcuate nucleus. J. Chem. Neuroanat. (2014) doi:10.1016/j.jchemneu.2014.06.008.

106. Timmermans, J. P., Adriaensen, D., Cornelissen, W. & Scheuermann, D. W. Structural organization and neuropeptide distribution in the mammalian enteric nervous system, with special attention to those components involved in mucosal reflexes. Comp. Biochem. Physiol. - A Physiol. (1997) doi:10.1016/S0300-9629(96)00314-3.

107. Furness, J. B. Types of neurons in the enteric nervous system. J. Auton. Nerv. Syst. (2000) doi:10.1016/S0165-1838(00)00127-2.

108. Quinson, N., Robbins, H., Clark, M. & Furness, J. Calbindin immunoreactivity of enteric neurons in the guinea-pig ileum. Cell Tissue Res. (2001) doi:10.1007/s004410100395.

109. Hirsch, E. C. Why are nigral catecholaminergic neurons more vulnerable than other cells in Parkinson’s disease? Ann. Neurol. (1992) doi:10.1002/ana.410320715.

110. Hack, N. J., Wride, M. C., Charters, K. M., Kater, S. B. & Parks, T. N. Developmental changes in the subcellular localization of calretinin. J. Neurosci. (2000) doi:10.1523/jneurosci.20-07-j0001.2000.

111. Pangrsic, T. et al. EF-hand protein Ca2+ buffers regulate Ca2+ influx and exocytosis in sensory hair cells. Proc. Natl. Acad. Sci. U. S. A. (2015) doi:10.1073/pnas.1416424112.

112. Christel, C. J. et al. Calretinin regulates Ca2+-dependent inactivation and facilitation of Cav2.1 Ca2+ channels through a direct interaction with the $α$12.1 subunit. J. Biol. Chem. (2012) doi:10.1074/jbc.M112.406363.

113. Ma, Y. et al. The effects of unilateral 6-OHDA lesion in medial forebrain bundle on the motor, cognitive dysfunctions and vulnerability of different striatal interneuron types in rats. Behav. Brain Res. (2014) doi:10.1016/j.bbr.2014.02.039.

114. Gao, R. & Penzes, P. Common Mechanisms of Excitatory and Inhibitory Imbalance in Schizophrenia and Autism Spectrum Disorders. Curr. Mol. Med. (2015) doi:10.2174/1566524015666150303003028.

115. Hoftman, G. D. et al. Altered cortical expression of GABA-related genes in schizophrenia: Illness progression vs developmental disturbance. Schizophr. Bull. (2015) doi:10.1093/schbul/sbt178.

116. Dai, H. et al. MiR-17 Regulates Prostate Cancer Cell Proliferation and Apoptosis Through Inhibiting JAK-STAT3 Signaling Pathway. Cancer Biother. Radiopharm. (2018) doi:10.1089/cbr.2017.2386.

117. Mura, A., Feldon, J. & Mintz, M. The expression of the calcium binding protein calretinin in the rat striatum: Effects of dopamine depletion and L-DOPA treatment. Exp. Neurol. (2000) doi:10.1006/exnr.2000.7441.

118. Abbas, N. et al. A wide variety of mutations in the parkin gene are responsible for autosomal recessive parkinsonism in Europe. Hum. Mol. Genet. (1999) doi:10.1093/hmg/8.4.567.

119. Valente, E. M. et al. Hereditary early-onset Parkinson’s disease caused by mutations in PINK1. Science (80-.). (2004) doi:10.1126/science.1096284.

120. Di Fonzo, A. et al. A frequent LRRK2 gene mutation associated with autosomal dominant Parkinson’s disease. Lancet (2005) doi:10.1016/S0140-6736(05)17829-5.

121. Bonifati, V. et al. Mutations in the DJ-1 gene associated with autosomal recessive early-onset parkinsonism. Science (80-.). (2003) doi:10.1126/science.1077209.

122. Kahle, P. J. et al. Subcellular localization of wild-type and Parkinson’s disease-associated mutant α- synuclein in human and transgenic mouse brain. J. Neurosci. (2000) doi:10.1523/jneurosci.20-17-06365.2000.

123. Schreiber, D. et al. Motility patterns of ex vivo intestine segments depend on perfusion mode. World J. Gastroenterol. (2014) doi:10.3748/wjg.v20.i48.18216.

124. Schäfer, K. H., Saffrey, M. J., Burnstock, G. & Mestres-Ventura, P. A new method for the isolation of myenteric plexus from the newborn rat gastrointestinal tract. Brain Res. Protoc. (1997) doi:10.1016/S1385-299X(96)00017-7.

125. Grundmann, D., Klotz, M., Rabe, H., Glanemann, M. & Schäfer, K. H. Isolation of high-purity myenteric plexus from adult human and mouse gastrointestinal tract. Sci. Rep. (2015) doi:10.1038/srep09226.

126. Bossolani, G. D. P. et al. Comparative analysis reveals Ce3D as optimal clearing method for in toto imaging of the mouse intestine. Neurogastroenterol. Motil. (2019) doi:10.1111/nmo.13560.

127. Follmer, C. et al. Dopamine affects the stability, hydration, and packing of protofibrils and fibrils of the wild type and variants of $α$-synuclein. Biochemistry (2007) doi:10.1021/bi061871+.

128. Cohen, J. 2.2. The Effect Size Index: d. in Statistical Power Analysis for the Behavioral Sciences (1988).

129. Bott, C. J. & Winckler, B. Intermediate filaments in developing neurons: Beyond structure. Cytoskeleton (2020) doi:10.1002/cm.21597.

130. Pellegrini, C. et al. Pathological remodelling of colonic wall following dopaminergic nigrostriatal neurodegeneration. Neurobiol. Dis. (2020) doi:10.1016/j.nbd.2020.104821.

131. Langston, J., Ballard, P., Tetrud, J. & Irwin, I. Chronic Parkinsonism in humans due to a product of meperidine-analog synthesis. Science (80-.). (1983) doi:10.1126/science.6823561.

132. Mayer, R. J., Lowe, J., Lennox, G., Doherty, F. & Landon, M. Intermediate filaments and ubiquitin: a new thread in the understanding of chronic neurodegenerative diseases. Progress in clinical and biological research (1989).

133. Przedborski, S. & Jackson-Lewis, V. Mechanisms of MPTP toxicity. Mov. Disord. (1998).

134. Cappelletti, G., Maggioni, M. G. & Maci, R. Influence of MPP+ on the state of tubulin polymerisation in NGF- differentiated PC12 cells. J. Neurosci. Res. (1999) doi:10.1002/(SICI)1097-4547(19990401)56:1<28::AID-JNR4>3.0.CO;2-2.

135. Federico, A. et al. Mitochondria, oxidative stress and neurodegeneration. J. Neurol. Sci. (2012) doi:10.1016/j.jns.2012.05.030.

136. Rahman, A. A. & Morrison, B. E. Contributions of VPS35 Mutations to Parkinson’s Disease. Neuroscience (2019) doi:10.1016/j.neuroscience.2019.01.006.

137. Muddapu, V. R., Dharshini, S. A. P., Chakravarthy, V. S. & Gromiha, M. M. Neurodegenerative Diseases – Is Metabolic Deficiency the Root Cause? Frontiers in Neuroscience (2020) doi:10.3389/fnins.2020.00213.

138. Bader, V. & Winklhofer, K. F. PINK1 and Parkin: Team players in stress-induced mitophagy. Biological Chemistry (2020) doi:10.1515/hsz-2020-0135.

139. Oliveira, L. O. D. et al. Prior exercise protects against oxidative stress and motor deficit in a rat model of Parkinson’s disease. Metab. Brain Dis. (2020) doi:10.1007/s11011-019-00507-z.

140. Ji, T. et al. Does perturbation in the mitochondrial protein folding pave the way for neurodegeneration diseases? Ageing Research Reviews (2020) doi:10.1016/j.arr.2019.100997.

141. Devi, L., Raghavendran, V., Prabhu, B. M., Avadhani, N. G. & Anandatheerthavarada, H. K. Mitochondrial import and accumulation of α-synuclein impair complex I in human dopaminergic neuronal cultures and Parkinson disease brain. J. Biol. Chem. (2008) doi:10.1074/jbc.M710012200.

142. Petryszyn, S., Parent, A. & Parent, M. The calretinin interneurons of the striatum: comparisons between rodents and primates under normal and pathological conditions. Journal of Neural Transmission (2018) doi:10.1007/s00702-017-1687-x.

143. Choi, B. K. et al. Large α-synuclein oligomers inhibit neuronal SNARE-mediated vesicle docking. Proc. Natl. Acad. Sci. U. S. A. (2013) doi:10.1073/pnas.1218424110.

144. Raja, S. A. et al. Increased expression levels of syntaxin 1a and synaptobrevin 2/vesicle-associated membrane protein-2 are associated with the progression of bladder cancer. Genet. Mol. Biol. (2019) doi:10.1590/1678-4685-gmb-2017-0339.

145. Trojanowski, J. Q. & Lee, V. M. Y. Aggregation of neurofilament and $α$-synuclein proteins in Lewy bodies: Implications for the pathogenesis of Parkinson disease and Lewy body dementia. Archives of Neurology (1998) doi:10.1001/archneur.55.2.151.

146. Gaugler, M. N. et al. Nigrostriatal overabundance of α-synuclein leads to decreased vesicle density and deficits in dopamine release that correlate with reduced motor activity. Acta Neuropathol. (2012) doi:10.1007/s00401-012-0963-y.

147. Zhang, X. M. et al. The A30P α-synuclein mutation decreases subventricular zone proliferation. Hum. Mol. Genet. (2019) doi:10.1093/hmg/ddz057.

148. Chen, Q. Q. et al. Age-dependent alpha-synuclein accumulation and aggregation in the colon of a transgenic mouse model of Parkinson’s disease. Transl. Neurodegener. (2018) doi:10.1186/s40035-018-0118-8.

149. Yang, C. P., Zhang, Z. H., Zhang, L. H. & Rui, H. C. Neuroprotective Role of MicroRNA-22 in a 6-Hydroxydopamine-Induced Cell Model of Parkinson’s Disease via Regulation of Its Target Gene TRPM7. J. Mol. Neurosci. (2016) doi:10.1007/s12031-016-0828-2.

150. Kim, J. et al. MiR-27a and miR-27b regulate autophagic clearance of damaged mitochondria by targeting PTEN-induced putative kinase 1 (PINK1). Mol. Neurodegener. (2016) doi:10.1186/s13024-016-0121-4.

151. Watts, M. E., Williams, S. M., Nithianantharajah, J. & Claudianos, C. Hypoxia-induced MicroRNA-210 targets neurodegenerative pathways. Non-coding RNA (2018) doi:10.3390/ncrna4020010.

52. Zhang, S. et al. Inhibition of BDNF production by MPP+ through up-regulation of miR-210-3p contributes to dopaminergic neuron damage in MPTP model. Neurosci. Lett. (2018) doi:10.1016/j.neulet.2017.10.014.

153. Wu, D. M. et al. Suppression of microRNA-342-3p increases glutamate transporters and prevents dopaminergic neuron loss through activating the Wnt signaling pathway via p21-activated kinase 1 in mice with Parkinson’s disease. J. Cell. Physiol. (2019) doi:10.1002/jcp.27577.

154. Gillardon, F. et al. MicroRNA and proteome expression profiling in early-symptomatic α-synuclein(A30P)-transgenic mice. Proteomics - Clin. Appl. (2008) doi:10.1002/prca.200780025.

155. Wu, X. et al. Regulatory Mechanism of miR-543-3p on GLT-1 in a Mouse Model of Parkinson’s Disease. ACS Chem. Neurosci. (2019) doi:10.1021/acschemneuro.8b00683.

156. Lungu, G., Stoica, G. & Ambrus, A. MicroRNA profiling and the role of microRNA-132 in neurodegeneration using a rat model. Neurosci. Lett. (2013) doi:10.1016/j.neulet.2013.08.001.

157. Liu, W. et al. MicroRNA Expression Profiling Screen miR-3557/324-Targeted CaMK/mTOR in the Rat Striatum of Parkinson’s Disease in Regular Aerobic Exercise. Biomed Res. Int. (2019) doi:10.1155/2019/7654798.

158. Li, J., Sun, Y. & Chen, J. Transcriptome sequencing in a 6-hydroxydopamine rat model of Parkinson’s disease. Genes Genet. Syst. (2019) doi:10.1266/ggs.18-00036.

159. Soreq, L. et al. Small RNA sequencing-microarray analyses in Parkinson leukocytes reveal deep brain stimulation-induced and splicing changes that classify brain region transcriptomes. Front. Mol. Neurosci. (2013) doi:10.3389/fnmol.2013.00010.

160. Martins, M. et al. Convergence of mirna expression profiling, $α$-synuclein interacton and GWAS in Parkinson’s disease. PLoS One (2011) doi:10.1371/journal.pone.0025443.

161. Alieva, A. K. et al. MiRNA expression is highly sensitive to a drug therapy in Parkinson’s disease. Park. Relat. Disord. (2015) doi:10.1016/j.parkreldis.2014.10.018.

162. Caggiu, E. et al. Differential expression of miRNA 155 and miRNA 146a in Parkinson’s disease patients. eNeurologicalSci (2018) doi:10.1016/j.ensci.2018.09.002.

163. Hoss, A. G., Labadorf, A., Beach, T. G., Latourelle, J. C. & Myers, R. H. microRNA profiles in Parkinson’s disease prefrontal cortex. Front. Aging Neurosci. (2016) doi:10.3389/fnagi.2016.00036.

164. Tatura, R. et al. Parkinson’s disease: SNCA-, PARK2-, and LRRK2- targeting microRNAs elevated in cingulate gyrus. Park. Relat. Disord. (2016) doi:10.1016/j.parkreldis.2016.09.028.

165. Alvarez-Erviti, L. et al. Influence of microRNA deregulation on chaperone-mediated autophagy and α-synuclein pathology in parkinson’s disease. Cell Death Dis. (2013) doi:10.1038/cddis.2013.73.

166. Briggs, C. E. et al. Midbrain dopamine neurons in Parkinson’s disease exhibit a dysregulated miRNA and target-gene network. Brain Res. (2015) doi:10.1016/j.brainres.2015.05.021.

167. Kim, J. et al. A microRNA feedback circuit in midbrain dopamine neurons. Science (80-.). (2007) doi:10.1126/science.1140481.

168. Martinez, B. & Peplow, P. V. MicroRNAs in Parkinson’s disease and emerging therapeutic targets. Neural Regeneration Research (2017) doi:10.4103/1673-5374.221147.

169. Cheng, M. et al. MicroRNA-181a suppresses parkin-mediated mitophagy and sensitizes neuroblastoma cells to mitochondrial uncoupler-induced apoptosis. Oncotarget (2016) doi:10.18632/oncotarget.9786.

170. Choi, I., Woo, J. H., Jou, I. & Joe, E. H. PINK1 deficiency decreases expression levels of mir-326, mir-330, and mir-3099 during brain development and neural stem cell differentiation. Exp. Neurobiol. (2016) doi:10.5607/en.2016.25.1.14.

171. Nair, V. D. & Ge, Y. Alterations of miRNAs reveal a dysregulated molecular regulatory network in Parkinson’s disease striatum. Neurosci. Lett. (2016) doi:10.1016/j.neulet.2016.06.061.

172. Gui, Y. X., Liu, H., Zhang, L. S., Lv, W. & Hu, X. Y. Altered microRNA profiles in cerebrospinal fluid exosome in Parkinson disease and Alzheimer disease. Oncotarget (2015) doi:10.18632/oncotarget.6158.

173. Burgos, K. et al. Profiles of extracellular miRNA in cerebrospinal fluid and serum from patients with Alzheimer’s and Parkinson’s diseases correlate with disease status and features of pathology. PLoS One (2014) doi:10.1371/journal.pone.0094839.

174. Cardo, L. F. et al. Profile of microRNAs in the plasma of Parkinson’s disease patients and healthy controls. Journal of Neurology (2013) doi:10.1007/s00415-013-6900-8.

175. Starhof, C. et al. The biomarker potential of cell-free microRNA from cerebrospinal fluid in Parkinsonian Syndromes. Mov. Disord. (2019) doi:10.1002/mds.27542.

176. dos Santos, M. C. T. et al. miRNA-based signatures in cerebrospinal fluid as potential diagnostic tools for early stage Parkinson’s disease. Oncotarget (2018) doi:10.18632/oncotarget.24736.

177. Barbagallo, C. et al. Specific Signatures of Serum miRNAs as Potential Biomarkers to Discriminate Clinically Similar Neurodegenerative and Vascular-Related Diseases. Cell. Mol. Neurobiol. (2020) doi:10.1007/s10571-019-00751-y.

178. Botta-Orfila, T. et al. Identification of blood serum micro-RNAs associated with idiopathic and LRRK2 Parkinson’s disease. J. Neurosci. Res. (2014) doi:10.1002/jnr.23377.

179. Vallelunga, A. et al. Identification of circulating microRNAs for the differential diagnosis of Parkinson’s disease and Multiple System Atrophy. Front. Cell. Neurosci. (2014) doi:10.3389/fncel.2014.00156.

