## Supplementary Information for "Functional and molecular early enteric biomarkers for Parkinson’s disease in mice and men"

#### Supplementary Figure 1

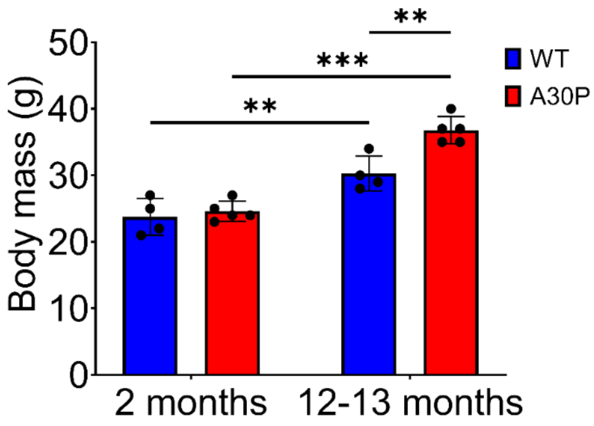


**Supplementary Figure 1. Body mass of 2-month-old and 12–13-month-old A30P and wild type (WT) mice.** The body weight of 2-month-old A30P mice did not differ from that of age-matched WT mice, while the body mass of 12–13-month-old A30P mice was significantly higher than that in age-matched WT mice. A30P mice are shown in red, WT mice in blue. Quantitative data were analyzed using GraphPad PRISM 8 and are expressed as means ± SD (WT: n = 4; A30P: n = 5 independent experiments). * $p \leq 0.05$, ** $p \leq 0.01$, and *** $p \leq 0.001$ using two-way ANOVA.

**Supplementary Figure 2**

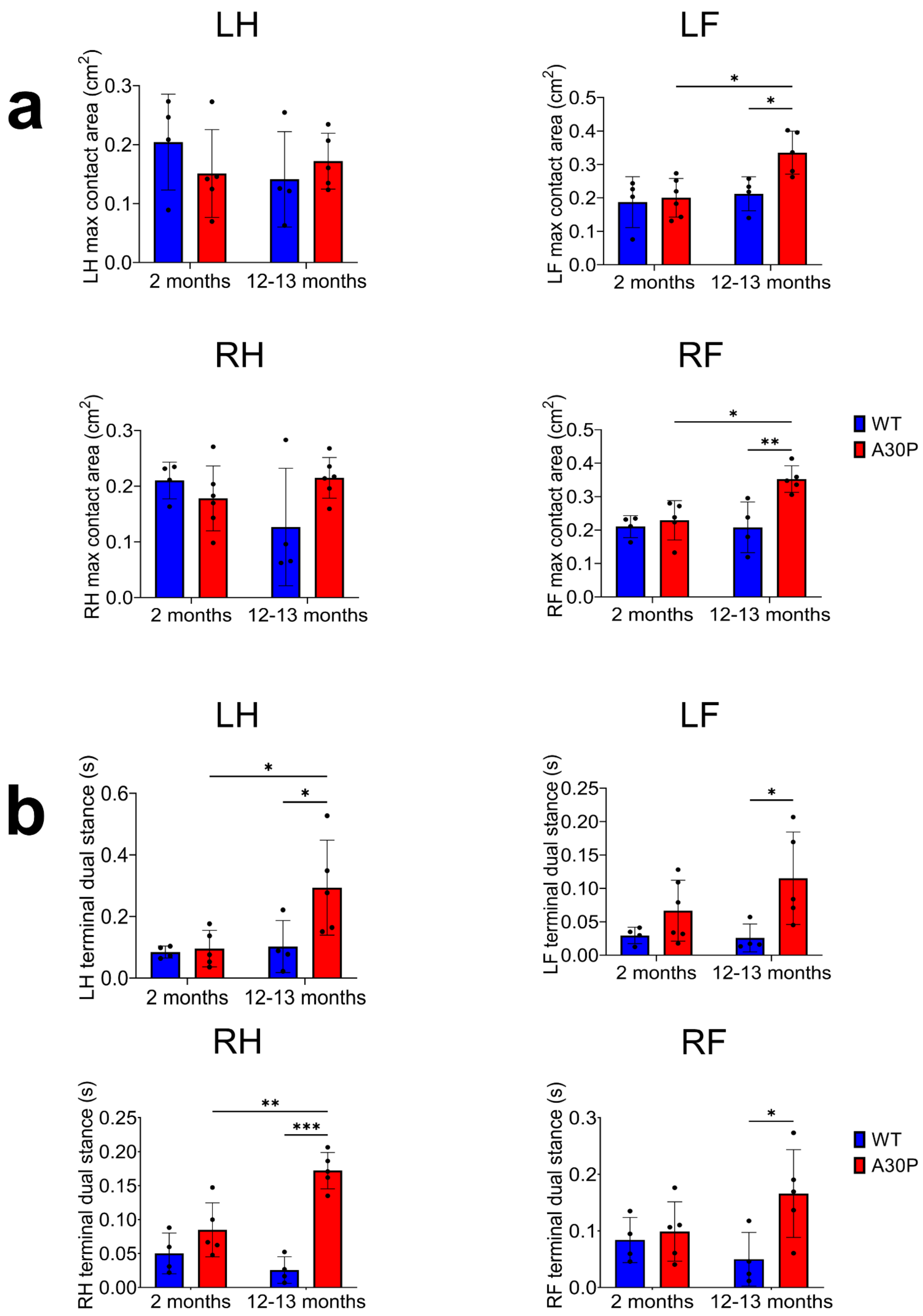

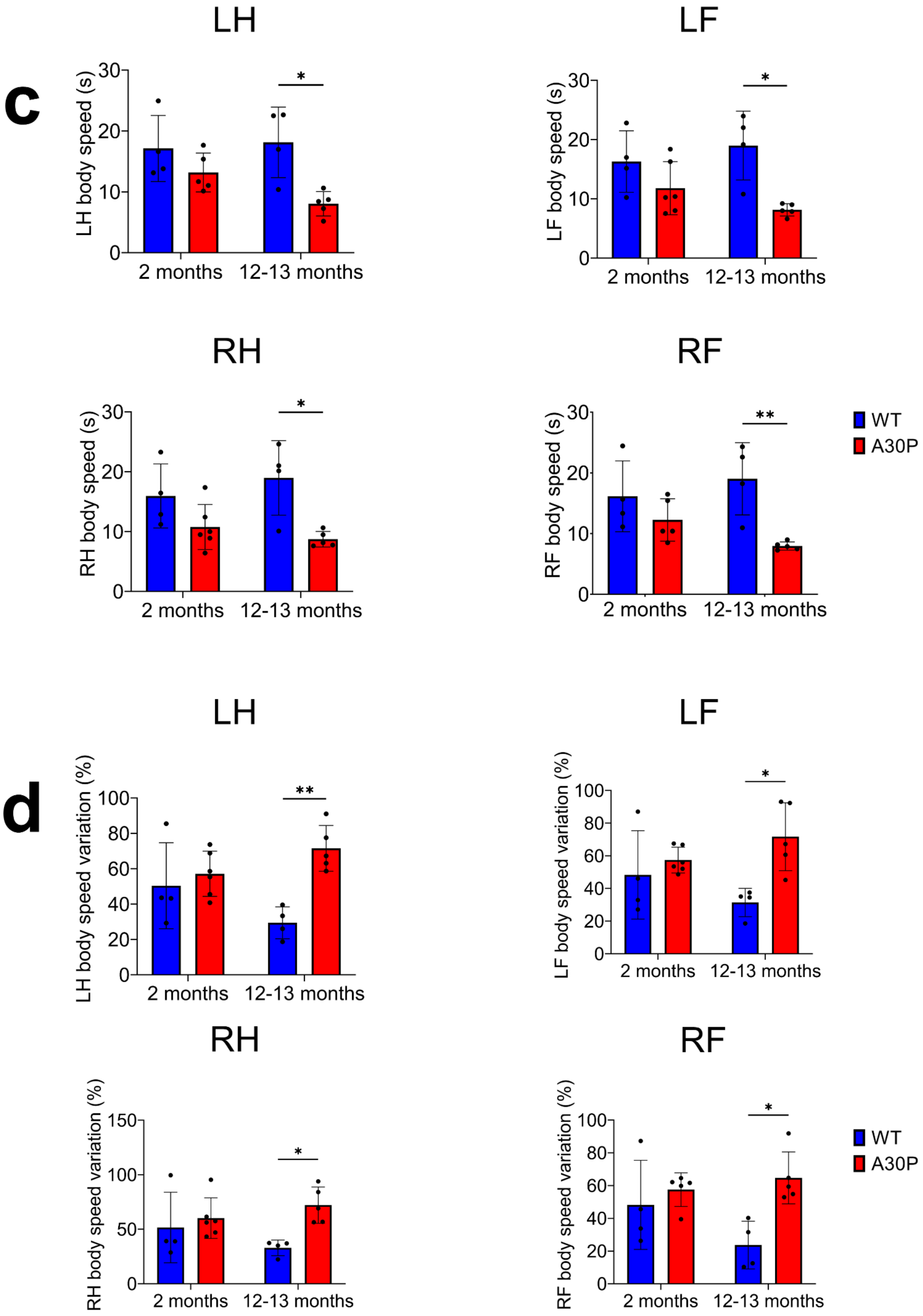
c
LH
LH body speed (s)
LF
LF body speed (s)
RH
RH body speed (s)
RF
RF body speed (s)
2 months
12-13 months
WT
A30P
d
LH
LH body speed variation (%)
LF
LF body speed variation (%)
RH
RH body speed variation (%)
RF
RF body speed variation (%)
2 months
12-13 months
WT
A30P

e

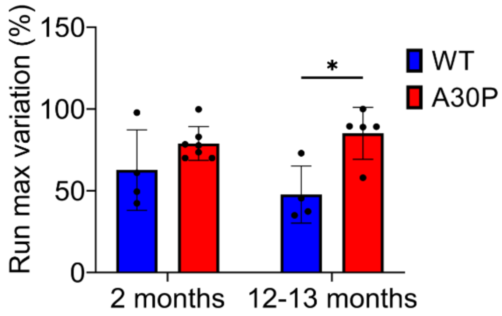


**Supplementary Figure 2. CatWalk XT analysis of 2-month-old and 12–13-month-old A30P and wild type (WT) mice.** Motoric disturbances in individual paws were significantly higher in 12–13-month-old A30P mice for all investigated parameters, including maximum contact area (a), terminal dual stance (b), body speed (c), and body speed variation (d), except for maximum contact areas (a) of hind paws. Two-month-old A30P mice did not show any motoric impairments compared with age-matched WT control mice. Maximum contact area represents the paw contact region within the glass plate. Terminal dual stance is the second step in a step cycle of a paw where the contra-lateral paw also makes contact with the glass plate. The speed of each single paw during a step cycle is defined as the body speed, while the body speed variation indicates changes of speed within placement of a single paw. (e) Run maximum variation was consistently higher in 12–13-month old A30P mice compared with age-matched WT controls. The run maximum variation designates a change of the speed within the same run; a low value shows a constant speed, while a high value indicates a large change in speed. WT mice are shown in blue, A30P mice in redA30P. Quantitative data were analyzed using GraphPad PRISM 8 and are expressed as means ± SD (WT: n = 4; A30P: n = 5 independent experiments). * $p \leq 0.05$, ** $p \leq 0.01$, and *** $p \leq 0.001$ using two-way ANOVA. LH = left hindpaw, LF = left frontpaw, RH = right hindpaw, RF = right frontpaw.

#### Supplementary Figure 3

# a

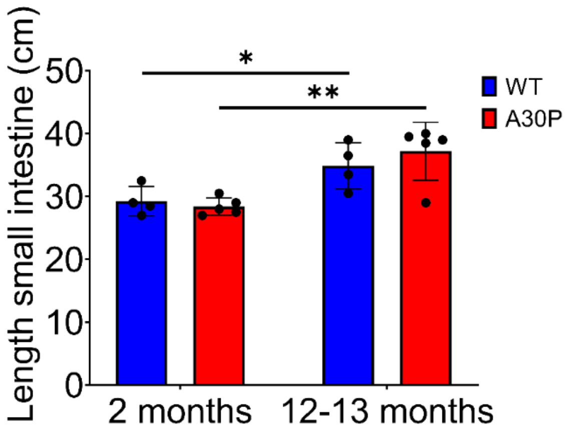


# b

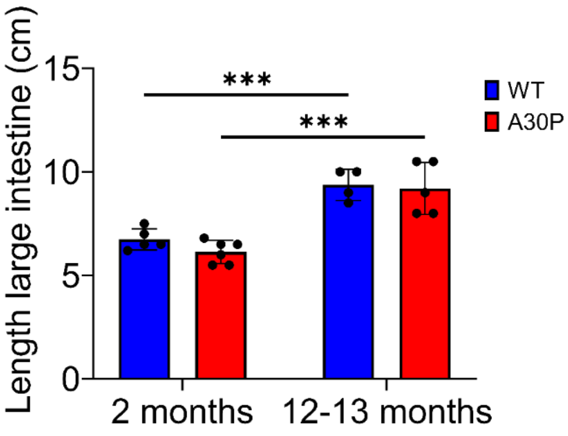


**Supplementary Figure 3. Gut length of 2-month-old and 12–13-month-old A30P and WT mice.** Lengths of small intestine (a) and large intestine (b) were measured in 2-month-old and 12–13-month-old A30P and WT mice. Gut lengths were not markedly different in A30P compared with age-matched WT controls. However, intestines were significantly longer in 12–13-month-old mice than in 2-month-old mice. A30P mice are shown in red, WT mice in blue. Quantitative data were analyzed using GraphPad PRISM 8 and are expressed as means ± SD (WT: n = 4; A30P: n = 5 independent experiments). * $p \leq 0.05$, ** $p \leq 0.01$, and *** $p \leq 0.001$ using two-way ANOVA.

**Supplementary Figure 4**

**a**

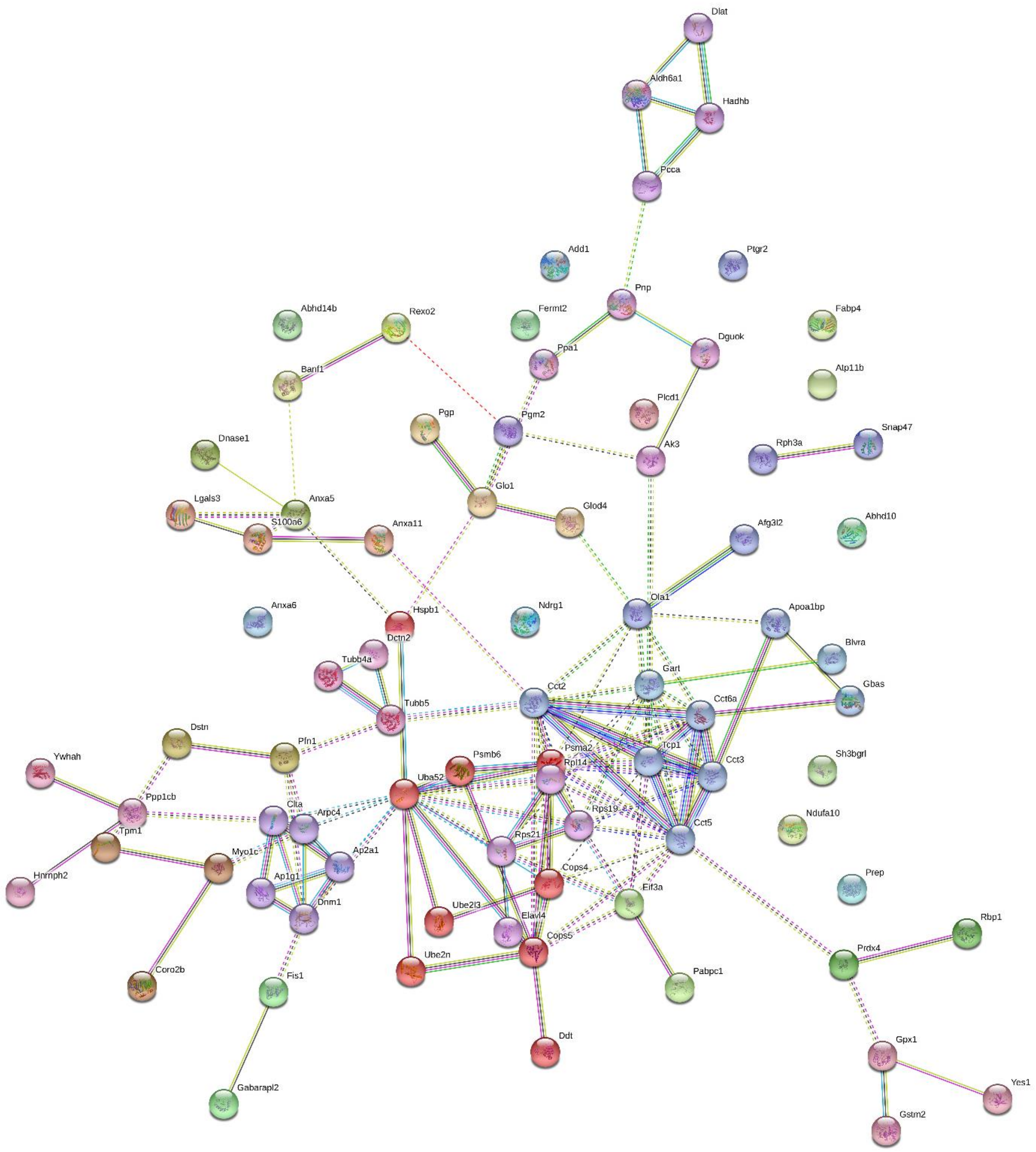

**b**

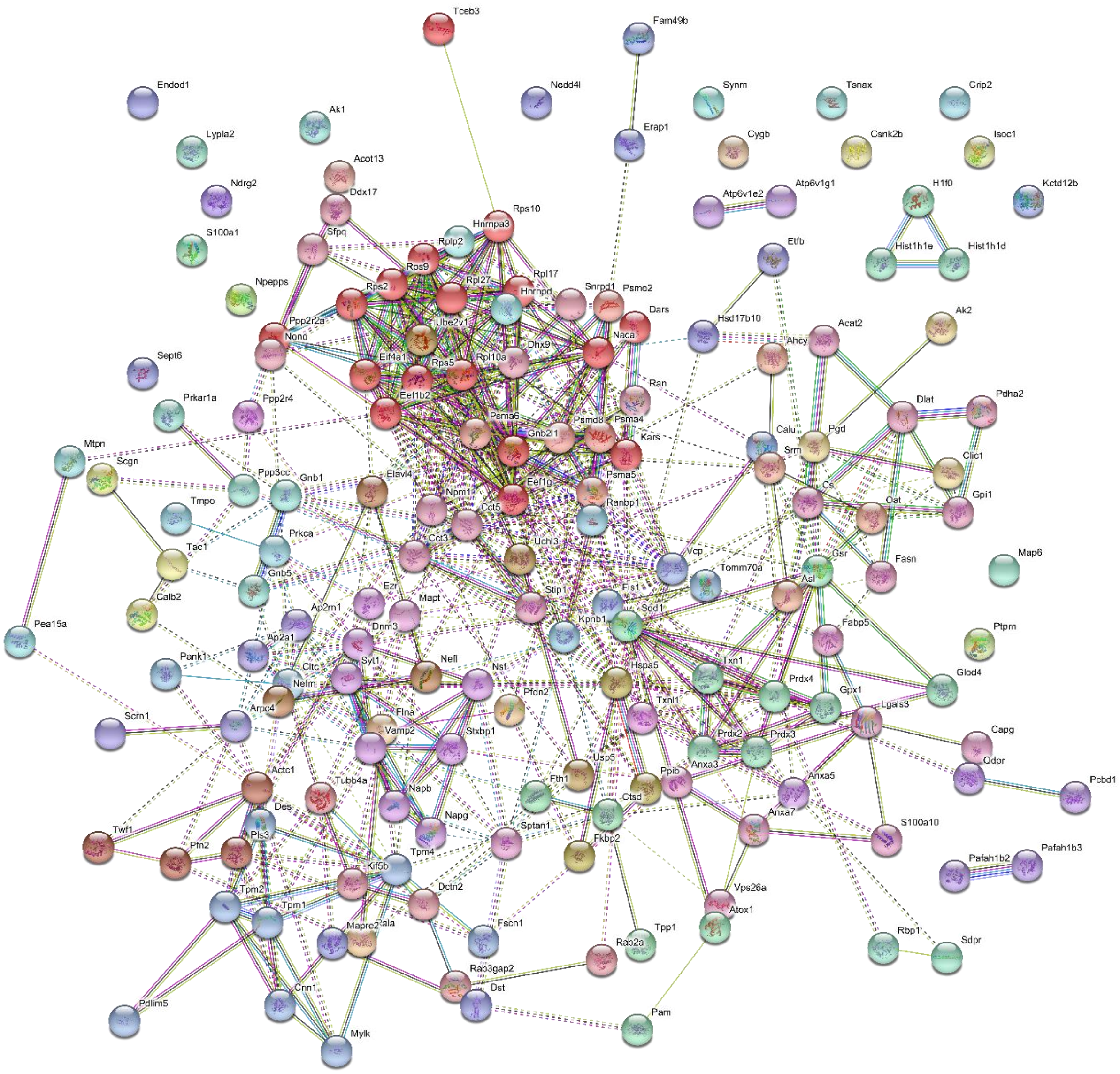


**Supplementary Figure 4. STRING network analysis of 1,044 proteins in the myenteric plexus (MP) of the small intestine (SI) and large intestine (LI).** (a) The network contains 74 proteins identified by mass spectroscopy as being differentially expressed in the SI MP of pre-symptomatic (ps)A30P mice vs. wild type (WT), while 12 proteins were not included in the network. (b) In the LI MP, 147 proteins were dysregulated in psA30P mice compared with in WT mice; here 15 proteins did not cluster within the protein network. Classified clusters are indicated by numerals. Network nodes symbolize proteins. The colored nodes are query proteins and first shell interactors. The edges represent protein-protein interactions. Colored lines between the proteins show the various associations in STRING. Data represent n = 4 independent experiments. STRING = Search Tool for the Retrieval of Interacting Genes.

#### Supplementary Figure 5

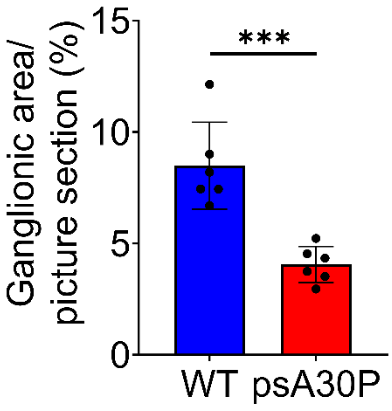


**Supplementary Figure 5. Ganglionic areas in pre-symptomatic (ps)A30P and wild type (WT) mice.** Ganglionic areas in full-thickness muscle layer preparations of the large intestine in psA30P mice were significantly reduced compared with those in WT mice. Quantitative data were analyzed using ImageJ and GraphPad PRISM 8, and are expressed as means ± SD from n = 6 independent experiments. WT are shown in blue, A30P mice in red. * $p \leq 0.05$, ** $p \leq 0.01$, and *** $p \leq 0.001$ using Student's t test.

#### Supplementary Figure 6

**a**

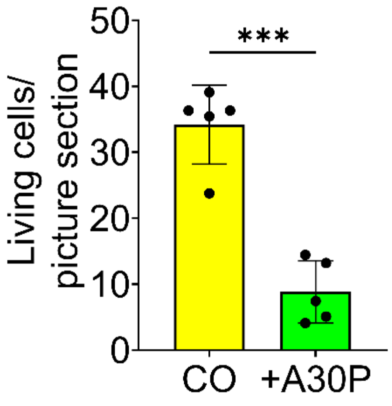

**b**

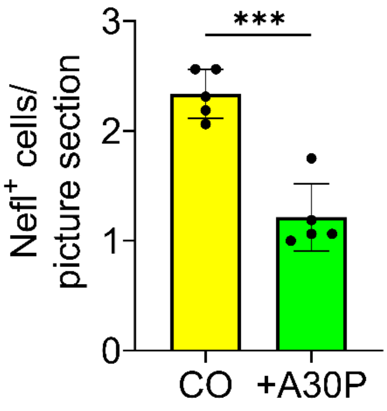


**c**

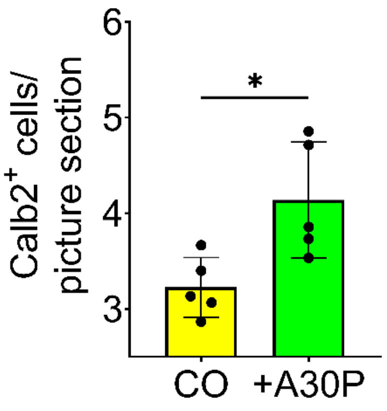


**Supplementary Figure 6. Acute treatment of enteric cells with A30P α-synuclein.** Cells isolated from the myenteric plexus of C57B6/J mice (postnatal day 2) were exposed to 0.5 μM A30P α-synuclein for 5 days *in vitro*. (a) Live-dead-assay showed that A30P α-synuclein treated cells (acute exposure) had significantly fewer viable cells than control cells did. Compared with the dopamine control (CO), acute exposure of A30P α-synuclein significantly reduced the total number of neurofilament light chain-positive cells (b) and increased the number of calbindin 2-positive cells (c). Quantitative data were analyzed using ImageJ and GraphPad PRISM 8 and are expressed as means ± SD from n = 5 independent experiments (N = 90 images per condition). WT controls are shown in yellow, A30P α-synuclein-treated cells in green. * $p \leq 0.05$, ** $p \leq 0.01$, and *** $p \leq 0.001$ using Student's t test.

Supplementary Figure 7

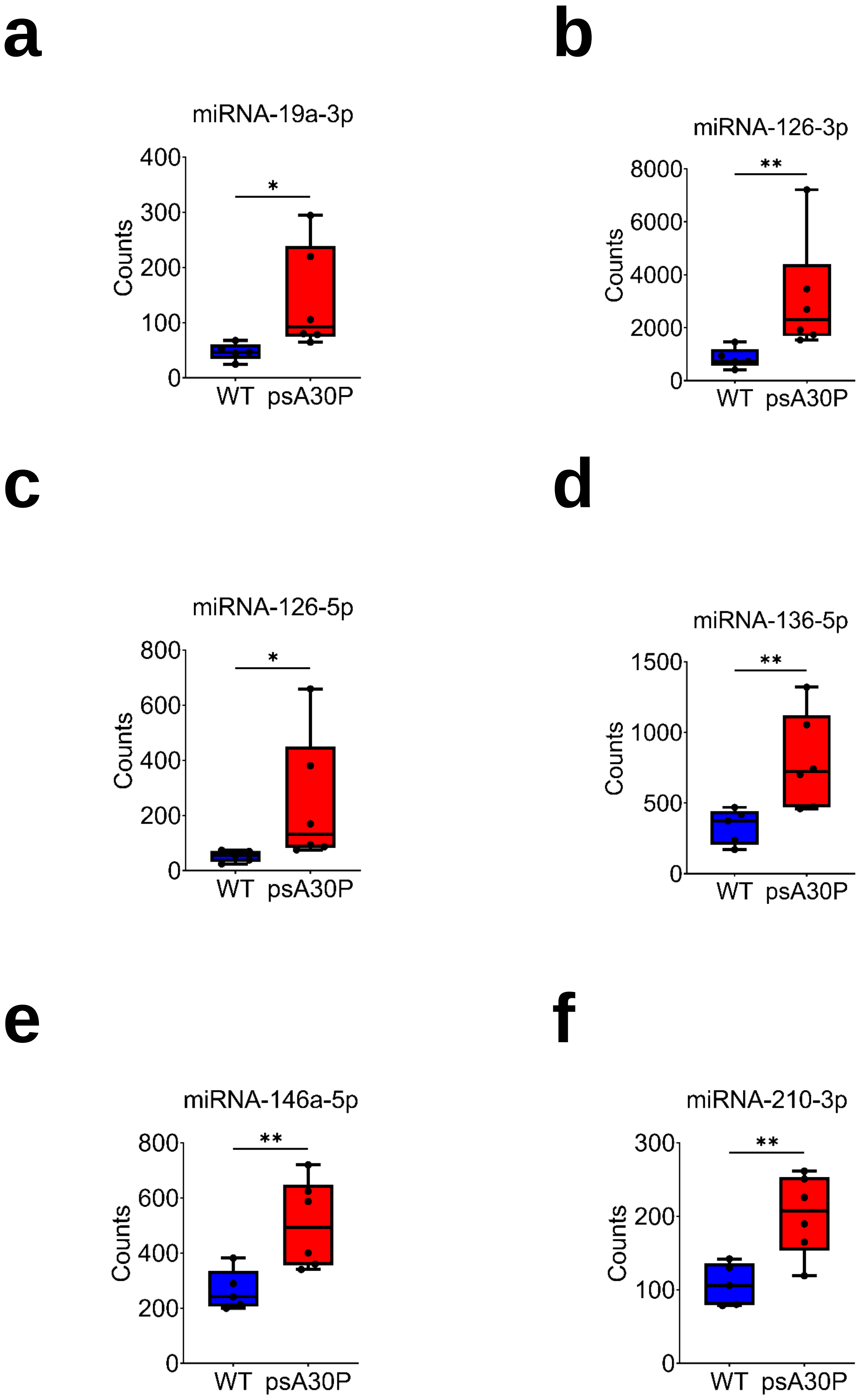

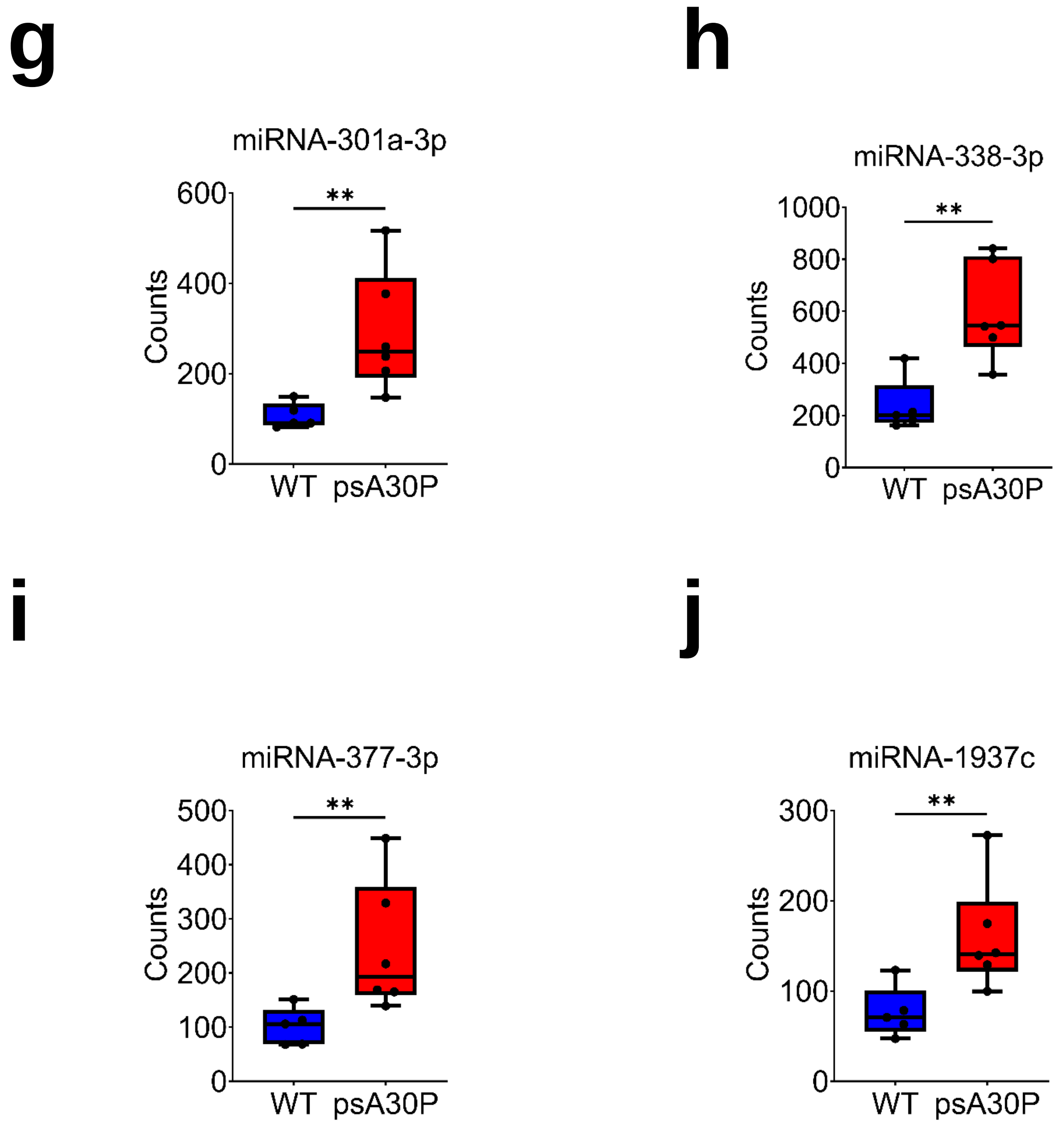


**Supplementary Figure 7. Top ten regulated miRNAs in the myenteric plexus (MP) of the large intestine (LI) in pre-symptomatic (ps)A30P mice and wild type (WT) mice.** Expression profiles of (a) miR-19a-3p, (b) miR-126-3p, (c) miR-12-5p, (d) miR-136-5p, (e) miR-146a-5p, (f) miR-210-3p, (g) miR-301a-3p, (h) miR-338-3p, (i) miR-377-3p, and (j) miR-1937c in the MP of the LI in psA30P mice compared with WT mice. All miRNAs were significantly upregulated in psA30P mice compared with WT mice. Quantitative data were analyzed using GraphPad PRISM 8 and are expressed as means ± SD. WT mice are shown in blue (n = 5), A30P mice in red (n = 6). * $p \leq 0.05$, ** $p \leq 0.01$, and *** $p \leq 0.001$ using Student's t test.

#### Supplementary Table 1

| Antibody | Dilution | Source | Host Species | Application |
|---|---|---|---|---|
| **Primary Antibodies** | | | | |
| anti-Calb2 | 1:200 | Abcam (ab702) | rabbit | IC, WMI |
| anti-Nefl | 1:500 | Santa Cruz (Biotech.sc-12980) | goat | IC, WMI |
| anti-PGP9.5 | 1:500 | Abcam (ab72910) | chicken | IC |
| anti-Vamp2 | 1:500 | Synaptic Systems (104 202) | rabbit | IC, WMI |
| anti-Tuj1 | 1:500 | BioLegend (801202) | mouse | IC, WMI |
| **Secondary Antibodies** | | | | |
| anti-goat IgG Alexa Fluor 594 | 1:500 | Thermofisher (A32758) | donkey | IC, WMI |
| anti-rabbit IgG Alexa Fluor 594 | 1:500 | Thermofisher (A32754) | donkey | IC, WMI |
| anti-mouse IgG Alexa Fluor 488 | 1:500 | Thermofisher (A-21206) | donkey | IC, WMI |
| anti-chicken IgG Alexa Fluor 488 | 1:500 | Jackson ImmunoResearch (703-545-155) | donkey | IC |

IC = immunocytochemistry; WMI= whole mount immunostaining

**Supplying companies:** Abcam, Cambridge, UK; Amersham, Synaptic Systems Gesellschaft für neurobiologische Forschung, Entwicklung und Produktion mbH; BioLegend, San Diego, USA; Santa Cruz Biotechnology, Inc., CA, USA. Invitrogen, Thermo Fisher Scientifi; DAKO, Germany; Jackson ImmunoResearch, Cambridge, UK

**Supplementary Table 1. Primary and secondary antibodies used and respective dilutions and applications.**

#### Supplementary Table 2

| | Small Intestine | | | | Large Intestine | | | |
|---|---|---|---|---|---|---|---|---|
| **Protein Name** | **WT** | **psA30P** | **Fold Change** | ***p*-value** | **WT** | **psA30P** | **Fold Change** | ***p*-value** |
| A0A087WPL5;E9QNN1;O70133;Q3UR42;A0A087WRT3 | 18514 | 25023 | 1.35 | 0.24827 | 33355 | 13192 | -2.53 | 0.05053 |
| A0A087WQE6;A0A087WNT1;P83940;A0A087WPE4 | 10986 | 10503 | -1.05 | 0.44105 | 75514 | 36184 | -2.09 | 0.18466 |
| A0A087WRU0;A0A087WQ94;A0A087WQM0;A0A087WQS0;E9Q0S6;Q9DBT6;A0A087WR29; A0A087WP40;Q8CGB6 | 62353 | 3509 | -17.77 | 0.09483 | 78597 | 25067 | -3.14 | 0.03645 |
| A0A087WRZ7;E9PWG4;P05977;P09542 | 40160 | 18740 | -2.14 | 0.43177 | 77826 | 48273 | -1.61 | 0.22143 |
| A0A087WS46;O70251;M0QWK5;M0QWH8;G3UX43;G3UZ47 | 22874 | 22634 | -1.01 | 0.89290 | 67538 | 15300 | -4.41 | 0.04620 |
| A0A087WS96;Q8BG73;A0A087WQX3;A0A087WQU5 | 9331 | 8867 | -1.05 | 0.65089 | 58475 | 5696 | -10.27 | 0.07912 |
| A0A0A0MQ80;Q3UMC0;D3Z0U5;D6RGM7;D3Z4J2;Q9D3R6 | 25411 | 26152 | 1.03 | 0.79617 | 77170 | 10610 | -7.27 | 0.01531 |
| A0A0A0MQ90;P97352 | 47131 | 41610 | -1.13 | 0.39027 | 60118 | 32038 | -1.88 | 0.33967 |
| A0A0A0MQA5;P68368;A0A087WQS4;A0A087WRB4;Q3UX10;A0A087WSB0;A0A087WSL5 | 42461 | 9804 | -4.33 | 0.22220 | 51121 | 15709 | -3.25 | 0.10029 |
| A0A0A0MQA8;P41539 | 25629 | 34766 | 1.36 | 0.27405 | 75328 | 21769 | -3.46 | 0.01974 |
| A0A0A0MQC7;A2A5Y6;P10637;B1AQW6;B1AQW5 | 44618 | 35596 | -1.25 | 0.46059 | 88328 | 49171 | -1.80 | 0.03365 |
| A0A0A0MQF6;P16858;S4R257;S4R1W1;D3YYI5;D3Z0Z9;S4R1W8;V9GXA7;V9GX06;S4R1N5; V9GXK0;S4R2G5;Q64467;REV__Q9Z2B5;REV__E9QQ30 | 1668119 | 1416277 | -1.18 | 0.17536 | 1314019 | 1363793 | 1.04 | 0.71670 |
| A0A0A0MQM0;P63242;J3QPS8;Q8BGY2 | 42465 | 59050 | 1.39 | 0.09175 | 68149 | 25780 | -2.64 | 0.08402 |
| A1BN54;Q7TPR4;D3YY95 | 491288 | 548202 | 1.12 | 0.31345 | 456807 | 500898 | 1.10 | 0.55253 |
| A2A6U3;Q80UG5;A2A6U5;A8Y5D3 | 35745 | 1956 | -18.28 | 0.25569 | 74929 | 23077 | -3.25 | 0.09469 |
| A2A7S7;Q91WQ3;F6VXZ2 | 3642 | 4549 | 1.25 | 0.48658 | 74808 | 20137 | -3.72 | 0.08654 |
| A2A9S2;A2A9R8;A2A9R6;Q8BVA9;A2A9S0;Q61701;A2A9S3;Q3UR02;Q80Y51;B1AXZ4;B1AXZ5; Q60899;B1AXZ0;Q80UJ0;B1AXZ6;Q60900;A2A9R4;H3BLG7;A2A9S1;A2A9R5 | 15741 | 9705 | -1.62 | 0.03172 | 61857 | 5477 | -11.29 | 0.05193 |
| A2AEX8;A2AEX6;P97447;A2AEX7;A2AEY1;A2AEY2;A2AEY0;A2AEX9;Q8CDC8 | 4813 | 69707 | 14.48 | 0.09704 | 79200 | 35051 | -2.26 | 0.06157 |
| A2AFI9;A2AFJ1;Q60973;F6ZLC6;F6U539 | 37482 | 7897 | -4.75 | 0.28876 | 75764 | 38699 | -1.96 | 0.21316 |
| A2AGN7;B7ZCF1;O88685;A0A087WPH7;F6Q2E3 | 9203 | 9543 | 1.04 | 0.93092 | 78079 | 22910 | -3.41 | 0.06186 |
| A2AH25;Q5FWK3 | 9421 | 12148 | 1.29 | 0.11853 | 38971 | 11872 | -3.28 | 0.25376 |
| A2AIM4;CON__Q3SX28;A2AIM5;E9PXF7;D3Z6F0;Q7TMK6 | 279172 | 483942 | 1.73 | 0.29282 | 173796 | 725901 | 4.18 | 0.01885 |
| A2AKU9 | 20500 | 25200 | 1.23 | 0.38157 | 109427 | 63673 | -1.72 | 0.40788 |

| | | | | | | | | |
|---|---|---|---|---|---|---|---|---|
| A2ALV3;Q62420;Q8BXU5;A2ALV1;F6ZL13 | 35302 | 8295 | -4.26 | 0.33914 | 74908 | 21131 | -3.54 | 0.08847 |
| A2AMW0;F7CAZ6 | 30239 | 27912 | -1.08 | 0.29485 | 80229 | 35837 | -2.24 | 0.07359 |
| A2AQ43;A2AQ44;A2AQ42;A2AQ45;A2AQ41;Q80TY0;F6VVN1;A2AQ47;A2AQ39 | 33740 | 7475 | -4.51 | 0.36033 | 75035 | 20616 | -3.64 | 0.08552 |
| A2AUD5;Q9CYZ2;Q3TAI4;F6VQ81;V9GWU5;Q8BKP1;Q3TUJ9 | 7431 | 7464 | 1.00 | 0.94902 | 58561 | 41208 | -1.42 | 0.59411 |
| A2BE93;Q9EQU5;A2BE92 | 24278 | 24357 | 1.00 | 0.98780 | 39496 | 30393 | -1.30 | 0.60509 |
| A3KGU7;P16546;A3KGU9;A3KGU4;P08032 | 1489433 | 1310457 | -1.14 | 0.45723 | 1601784 | 1256270 | -1.28 | 0.03014 |
| A3KMP2;D6RH59 | 35027 | 6077 | -5.76 | 0.30919 | 74959 | 39662 | -1.89 | 0.26587 |
| A6MDD2;Q60673;A0A087WPU7;A0A087WQF2 | 44682 | 40447 | -1.10 | 0.37941 | 77103 | 47812 | -1.61 | 0.02846 |
| A6PWS5 | 21900 | 19708 | -1.11 | 0.34081 | 77202 | 45179 | -1.71 | 0.25733 |
| A8DUK4;P02088;E9Q223;P02089;CON__Q3SX09;CON__P02070;P02104 | 327531 | 540373 | 1.65 | 0.11564 | 132571 | 145092 | 1.09 | 0.62553 |
| B0QZN5;P63044 | 70095 | 35890 | -1.95 | 0.42149 | 83626 | 20552 | -4.07 | 0.00530 |
| B1AQF4;Q9D7X3;H3BKL8;Q3V2Y9;H3BKD1 | 11388 | 7462 | -1.53 | 0.23176 | 59944 | 16514 | -3.63 | 0.12531 |
| B1AQY9;B1AQZ0;Q8CHH9;B7ZC46;E0CYM4 | 15084 | 10720 | -1.41 | 0.06672 | 78187 | 29797 | -2.62 | 0.07847 |
| B1ARA3;P61255;B1ARA5 | 36710 | 5322 | -6.90 | 0.26571 | 76391 | 22995 | -3.32 | 0.07795 |
| B1ATI9;Q3U432;Q60780 | 8550 | 6293 | -1.36 | 0.61054 | 76413 | 41195 | -1.85 | 0.24571 |
| B1ATZ0;B1ATZ1;Q3UMA3;Q99LI8;B1ATY9;F6VV02 | 36249 | 6711 | -5.40 | 0.29426 | 57001 | 35832 | -1.59 | 0.50817 |
| B1AWE0;Q6PFA2 | 15683 | 9336 | -1.68 | 0.00830 | 76762 | 24179 | -3.17 | 0.07739 |
| B1AX58 | 48517 | 55472 | 1.14 | 0.50034 | 58938 | 11276 | -5.23 | 0.05222 |
| B1AX78;Q2TPA8;B1AX77 | 34946 | 4118 | -8.49 | 0.28196 | 75665 | 40920 | -1.85 | 0.25970 |
| B1AXW5;B1AXW6;P35700;B1AXW4 | 266447 | 269241 | 1.01 | 0.88353 | 82505 | 113337 | 1.37 | 0.38434 |
| B1AYL1 | 97929 | 30353 | -3.23 | 0.10366 | 79186 | 82629 | 1.04 | 0.94085 |
| B1B1A8;Q6PDN3 | 54875 | 96039 | 1.75 | 0.16760 | 64379 | 135687 | 2.11 | 0.03342 |
| B2M1R6;P61979;H3BLL4;H3BKD0;D3Z5X4;D3YWG1;H3BK96;H3BKI8;H3BLP7;H3BJS9;Q8BT23; H3BJ43 | 139362 | 130618 | -1.07 | 0.44014 | 84839 | 84027 | -1.01 | 0.96950 |
| B7FAU9;Q8BTM8;B7FAV1;F6XC15;J3JS91;F6Z2C0;F7AVL7 | 364969 | 651919 | 1.79 | 0.16341 | 727783 | 1452739 | 2.00 | 0.01382 |
| B7ZBY7;E9PY39;Q9CZY3;B2KF55;B7ZBY6;D3Z6R2 | 18903 | 24703 | 1.31 | 0.31814 | 78423 | 12784 | -6.13 | 0.01290 |
| CON__P00761 | 256430 | 202689 | -1.27 | 0.10482 | 160998 | 117243 | -1.37 | 0.32620 |
| CON__P04264;CON__H-INV:HIT000016045 | 303739 | 243465 | -1.25 | 0.63423 | 313051 | 294915 | -1.06 | 0.86350 |
| CON__P13645;A2A513;CON__P02535-1;P02535;CON__Q7Z3Y7;CON__Q148H6;CON__Q7Z3Z0; CON__Q7Z3Y8;CON__Q99456;CON__Q2M2I5;CON__Q7Z3Y9;Q8VCW2;Q9Z320;A6BLY7 | 126149 | 118960 | -1.06 | 0.92245 | 86331 | 144279 | 1.67 | 0.35703 |
| CON__P19001;P19001 | 56764 | 52116 | -1.09 | 0.91608 | 74939 | 46874 | -1.60 | 0.35934 |
| CON__P35527 | 144806 | 100582 | -1.44 | 0.46525 | 121961 | 107990 | -1.13 | 0.80507 |

| | | | | | | | | |
|---|---|---|---|---|---|---|---|---|
| CON__P35908;CON__Q7Z794;CON__REFSEQ:XP_932229;CON__Q7RTS7;CON__Q32MB2 | 75440 | 63676 | -1.18 | 0.77582 | 72241 | 74991 | 1.04 | 0.92586 |
| D3YUT3;D3YUG3;D3Z5R8;D3Z722;Q9CZX8;S4R223 | 10658 | 6973 | -1.53 | 0.00454 | 59519 | 11730 | -5.07 | 0.09851 |
| D3YUZ8;E9Q2L2;Q91XM9;D3YWU0 | 67803 | 3612 | -18.77 | 0.08808 | 75135 | 20648 | -3.64 | 0.08454 |
| D3YVN7;Q8BFR5 | 6718 | 6090 | -1.10 | 0.59655 | 80130 | 15420 | -5.20 | 0.00997 |
| D3YVV9;E9Q1U2;Q91YE8;F6WWS1 | 4005 | 7106 | 1.77 | 0.13552 | 75105 | 26030 | -2.89 | 0.10413 |
| D3YW48;O88456;Q9D7J7 | 47677 | 44333 | -1.08 | 0.39734 | 73289 | 30461 | -2.41 | 0.12311 |
| D3YW87;D3Z576;Q8VHX6 | 35882 | 17716 | -2.03 | 0.52555 | 69051 | 75227 | 1.09 | 0.84386 |
| D3YWF6;Q7TQI3;D3Z7K0 | 36038 | 41001 | 1.14 | 0.44911 | 63926 | 21002 | -3.04 | 0.10313 |
| D3YWL1;D3YW33;P35276 | 9724 | 10582 | 1.09 | 0.39453 | 76182 | 37005 | -2.06 | 0.17641 |
| D3YXC7;H3BKL7;E9Q797;P49183 | 25139 | 11270 | -2.23 | 0.00073 | 75491 | 21080 | -3.58 | 0.08203 |
| D3YYK8;E9Q6X0;Q8R001;Q3TG90 | 8390 | 46107 | 5.50 | 0.25444 | 32855 | 15736 | -2.09 | 0.02178 |
| D3YYM6;Q91V55;P97461 | 34554 | 2629 | -13.14 | 0.26910 | 76210 | 7146 | -10.67 | 0.01557 |
| D3YYT0;P15116;P39038 | 13529 | 12473 | -1.08 | 0.79394 | 44430 | 13414 | -3.31 | 0.17053 |
| D3Z0F5;O88545;F6QK86 | 8558 | 10011 | 1.17 | 0.54782 | 56750 | 74819 | 1.32 | 0.51686 |
| D3Z1Z8;D3Z5N2;P54227 | 34377 | 32863 | -1.05 | 0.77953 | 78734 | 26553 | -2.97 | 0.06428 |
| D3Z2H9;D3YVR0 | 18222 | 49186 | 2.70 | 0.28414 | 62153 | 11259 | -5.52 | 0.07074 |
| D3Z440;Q9CZ04;D3YVI6;D3Z0S0 | 38572 | 5465 | -7.06 | 0.28502 | 55749 | 43965 | -1.27 | 0.69897 |
| D3Z4B2;Q9CWZ7 | 37395 | 6244 | -5.99 | 0.26694 | 77970 | 10752 | -7.25 | 0.01304 |
| D3Z645;Q9QZ88;D3YYD5;D3YW98 | 15232 | 13801 | -1.10 | 0.61093 | 59488 | 43302 | -1.37 | 0.60743 |
| D3Z7P3;D3Z7P4;F7B327;F6U529 | 42457 | 41334 | -1.03 | 0.86411 | 80575 | 39441 | -2.04 | 0.08804 |
| E0CXB9;Q61301;Q8BS72 | 39207 | 7277 | -5.39 | 0.24668 | 59322 | 41508 | -1.43 | 0.58018 |
| E9PUD2;Q8K1M6 | 17322 | 23392 | 1.35 | 0.07762 | 42490 | 23480 | -1.81 | 0.18357 |
| E9PUM4;Q71LX4 | 24169 | 25735 | 1.06 | 0.61181 | 26912 | 30180 | 1.12 | 0.82169 |
| E9PVC6;E9Q9E1;E9PVC5;Q6NZJ6;E9Q770;A2AMI7;Z4YKC4;A2AMI2;Q80XI3 | 35290 | 6331 | -5.57 | 0.30821 | 75398 | 40286 | -1.87 | 0.26094 |
| E9PW66;P28656 | 14447 | 13879 | -1.04 | 0.76487 | 21592 | 10877 | -1.99 | 0.16734 |
| E9PWE8;Q3TT92;Q62188;D3YUS0;D3Z567 | 597566 | 550623 | -1.09 | 0.46587 | 292927 | 342763 | 1.17 | 0.64079 |
| E9PXB7;Q8CFI0 | 33664 | 28749 | -1.17 | 0.89087 | 80200 | 21441 | -3.74 | 0.05872 |
| E9PXV3;E9Q1W0;E9Q1V9;E9Q1T1;Q6PHZ2;Q8CCM0;E9QAJ4;D6RDQ8;F6ZIE1;F6RWZ9;F7A856;<br>E9PYZ8 | 19887 | 17896 | -1.11 | 0.64158 | 62200 | 33593 | -1.85 | 0.15440 |
| E9PXX7;Q91W90 | 36059 | 6690 | -5.39 | 0.29778 | 58220 | 41306 | -1.41 | 0.60474 |
| E9PYH2;Q91V12 | 328163 | 321991 | -1.02 | 0.89768 | 107878 | 112592 | 1.04 | 0.86652 |
| E9PZ00;Q8BFQ1;K3W4L3;J3QPG5;Q61207 | 49680 | 55855 | 1.12 | 0.24403 | 65459 | 34122 | -1.92 | 0.24735 |
| E9PZF0;Q01768;Q5NC79 | 200219 | 206826 | 1.03 | 0.74294 | 68874 | 91110 | 1.32 | 0.45988 |

| | | | | | | | | |
|---|---|---|---|---|---|---|---|---|
| E9Q0B0;B7ZWM6;Q3UTJ2;B2RXQ9;B9EKP8;D3Z080;Z4YJR7;F6V513;F6ZWT0;F7BEU1;D3Z203 | 38581 | 10249 | -3.76 | 0.30177 | 63928 | 20649 | -3.10 | 0.10118 |
| E9Q0W6;F6VVW4;D3Z0Y4;F7A5C5;D3YYR9;F6W6N3;Q69ZX8 | 10526 | 11565 | 1.10 | 0.55521 | 60959 | 28898 | -2.11 | 0.28170 |
| E9Q3T0;P47955 | 19800 | 20479 | 1.03 | 0.83308 | 63286 | 12940 | -4.89 | 0.05736 |
| E9Q3X0;Q9EQK5;D3Z2N7 | 7809 | 7337 | -1.06 | 0.71848 | 58086 | 43041 | -1.35 | 0.64455 |
| E9Q452;Q8BSH3;E9Q454;P58771 | 15312 | 18046 | 1.18 | 0.05678 | 61964 | 26943 | -2.30 | 0.24136 |
| E9Q586;E9Q3M3;D3YX34;O08788;D3Z2M9;D3YYG9 | 14201 | 15266 | 1.07 | 0.71490 | 58715 | 23598 | -2.49 | 0.26849 |
| E9Q5A0;P19253;S4R281;REV__A2A8E7;REV__P52825 | 64839 | 2113 | -30.69 | 0.09055 | 76469 | 8883 | -8.61 | 0.01619 |
| E9Q5H2;E9PZF5;P97822 | 65272 | 3474 | -18.79 | 0.09259 | 56757 | 21007 | -2.70 | 0.27779 |
| E9Q616;G5E8K8 | 289932 | 232979 | -1.24 | 0.35843 | 104326 | 130443 | 1.25 | 0.64559 |
| E9Q7H5;B2RXM2;J3QNY1;A2AL13 | 121850 | 127060 | 1.04 | 0.50038 | 92879 | 60162 | -1.54 | 0.01342 |
| E9Q7L0 | 34892 | 4892 | -7.13 | 0.29442 | 76166 | 37001 | -2.06 | 0.17648 |
| E9Q7Q3;D3Z6I8 | 116002 | 119249 | 1.03 | 0.73742 | 67328 | 64776 | -1.04 | 0.90063 |
| E9Q7S0;F7BQW7;D3Z656;Q8CHC4;D3Z1M7;D3YWM9;F7BI35;D3YZB3;E9Q4P5;D3Z6E7;D3YZB4;D3YZB2;F8WHD8;Q9D2G5 | 8716 | 7515 | -1.16 | 0.39652 | 77947 | 24244 | -3.22 | 0.06644 |
| E9Q8N5;Q08EB6;F7DCH5;Q8BRT1 | 38244 | 5078 | -7.53 | 0.28568 | 35424 | 40268 | 1.14 | 0.86480 |
| E9Q912;E9Q6Q4 | 26978 | 30939 | 1.15 | 0.51351 | 43344 | 12949 | -3.35 | 0.19358 |
| E9Q9C5;P63082;D3Z3B2 | 65467 | 63802 | -1.03 | 0.96903 | 122012 | 74435 | -1.64 | 0.33922 |
| E9Q9F5;E9Q1G8;O55131;Q8C650 | 61768 | 51436 | -1.20 | 0.02055 | 99250 | 69900 | -1.42 | 0.02431 |
| E9Q9J0;E9Q5F6;Q5SX22;P62984;P62983;P0CG49;P0CG50;E9Q4P0;E9QNP0;J3QK04;D3YYZ2 | 299767 | 446117 | 1.49 | 0.04988 | 176171 | 237042 | 1.35 | 0.44558 |
| E9Q9X1;S4R1P5;Q91ZU6;A0A087WSP0;A0A087WRB8;S4R2N8;A0A087WPR7;S4R1Y6;S4R2A8;F6RCJ3;F6YKN8 | 62438 | 4694 | -13.30 | 0.10003 | 79120 | 20020 | -3.95 | 0.04831 |
| E9QA15;D3Z6I7;REV__B2RXW8;REV__Q8BSS9;REV__B8QI34;REV__B2RXQ2 | 13075 | 14713 | 1.13 | 0.68355 | 77108 | 44798 | -1.72 | 0.21207 |
| E9QAZ2;Q9CZM2;B8JKK2 | 4283 | 3358 | -1.28 | 0.04315 | 42544 | 10294 | -4.13 | 0.17831 |
| E9QJT5;P56376;Q8BMV3 | 9602 | 10526 | 1.10 | 0.43957 | 75328 | 35300 | -2.13 | 0.18352 |
| E9QK48;Q7TNG5;D3YWS2;D6RGM3 | 12706 | 18068 | 1.42 | 0.06682 | 57154 | 41371 | -1.38 | 0.63278 |
| E9QLL2;Q8BZ98;F2Z460;E0CXZ8 | 3615 | 2199 | -1.64 | 0.13001 | 78745 | 22946 | -3.43 | 0.05603 |
| E9QN08;Q80T06;P57776;Q91VK2;D3YUQ9;F6ZFU0;D3Z7N2;D3YZT9;D3YY68 | 16451 | 16927 | 1.03 | 0.73119 | 77031 | 25029 | -3.08 | 0.07692 |
| E9QN99;A0A087WPF8;A0A087WQN8;A0A087WSR2;A0A087WRJ2;Q8VCR7;A0A087WP24 | 9724 | 16252 | 1.67 | 0.01028 | 60740 | 24224 | -2.51 | 0.16353 |
| E9QNA7;Q62417;E9Q6A3;E9PYX6;D3Z5J3 | 38941 | 10354 | -3.76 | 0.35097 | 44387 | 21822 | -2.03 | 0.30786 |
| E9QNY3 | 8356 | 23079 | 2.76 | 0.28153 | 75510 | 31032 | -2.43 | 0.13704 |

| | | | | | | | | |
|---|---|---|---|---|---|---|---|---|
| E9QPE7;Q02566;A2AQP0;B8JJH5;REV__E9Q3T3;REV__L0N7N1;B1AR69;G3UW82;Q91Z83;P13542;<br>Q5SX39;P13541;Q5SX40 | 759003 | 1363327 | 1.80 | 0.36978 | 4616986 | 5779775 | 1.25 | 0.42918 |
| F2Z3Z1;Q8BGZ1 | 35038 | 40356 | 1.15 | 0.90081 | 13224 | 21012 | 1.59 | 0.63545 |
| F6RPJ9;Q9JHR7 | 35769 | 7909 | -4.52 | 0.32567 | 74986 | 41037 | -1.83 | 0.27564 |
| F6SVV1;P62082 | 7145 | 6394 | -1.12 | 0.52977 | 62608 | 16440 | -3.81 | 0.06127 |
| F6YVP7;P62270;S4R1N6 | 11062 | 10000 | -1.11 | 0.70236 | 66689 | 17118 | -3.90 | 0.05632 |
| F6YY69;F6VW30;P68254 | 99381 | 102267 | 1.03 | 0.56924 | 134636 | 55695 | -2.42 | 0.06687 |
| F6ZV59;G5E8G0;G3X9W0;Q60668;E9Q5B6;F6SHF3;Q9D3U4 | 81203 | 79497 | -1.02 | 0.82399 | 72715 | 41479 | -1.75 | 0.04866 |
| F7CDT0;F6WMC0;P61082;G5E919 | 5331 | 5418 | 1.02 | 0.82067 | 75090 | 35106 | -2.14 | 0.18696 |
| F7D432;E9PWE0;E0CYV0;P23506;F6V9F1;F6TXE3 | 43439 | 40452 | -1.07 | 0.45033 | 49723 | 18301 | -2.72 | 0.10406 |
| F8VQA4;P97467;E9Q704;F6T0D3;F6TGI4;F7D4K4 | 92587 | 89622 | -1.03 | 0.76076 | 83722 | 43086 | -1.94 | 0.03008 |
| F8WGF2;Q9Z0J4;S4R255;S4R1D5;P29477 | 93669 | 76204 | -1.23 | 0.14745 | 110267 | 167250 | 1.52 | 0.22336 |
| F8WGN6;E9Q0J5;Q9QXL2;E9QMU1;J3QNW9;B7ZNG0;Q7M6Z4 | 18362 | 14974 | -1.23 | 0.30724 | 61508 | 28352 | -2.17 | 0.26676 |
| F8WGT1;Q68FL4;F8WI65;H3BKT5;Q80SW1;E9PX77;D3YX97;H3BL31;D3YYM7 | 55058 | 32897 | -1.67 | 0.40840 | 74961 | 25810 | -2.90 | 0.10502 |
| F8WID5 | 620580 | 939093 | 1.51 | 0.38401 | 686659 | 1252153 | 1.82 | 0.11306 |
| F8WIX8;Q64523;Q6GSS7 | 28103 | 41916 | 1.49 | 0.35955 | 376921 | 488623 | 1.30 | 0.49153 |
| F8WJK8;Q99L47;E9Q1V0;E9Q1X9 | 38828 | 40713 | 1.05 | 0.69109 | 61701 | 30186 | -2.04 | 0.28014 |
| G3UVV4;P17710;D3YYR4;D3Z365;E9Q5B5;O08528;D3Z105;B4YB29;E9Q9M6;F6QD41;E9Q3Z4;<br>E9Q8S8;Q3TRM8 | 188735 | 160886 | -1.17 | 0.15817 | 189530 | 201773 | 1.06 | 0.70063 |
| G3UWG1;P62897;CON__P62894;P00015 | 44354 | 39588 | -1.12 | 0.14011 | 102339 | 88523 | -1.16 | 0.25226 |
| G3UXT7;Q8CFQ9;P56959;F6QCI0;Q8BQ46;G3UZD2;Q91VQ2 | 10058 | 10244 | 1.02 | 0.88931 | 76177 | 23099 | -3.30 | 0.08072 |
| G3UXZ5;P97371;G3UXY0;G3X9K9;G3UWN9;G3UXR1 | 51920 | 48653 | -1.07 | 0.60770 | 31920 | 18401 | -1.73 | 0.06726 |
| G3UYV7;P62858;J3QNN8 | 12218 | 10541 | -1.16 | 0.18415 | 60690 | 7928 | -7.65 | 0.06932 |
| G3UZX6;H7BWX9;D3Z794;G3UZ60;G3UWI9;G3UWX9;G3UZA7;P61957;Q9Z172 | 36501 | 6315 | -5.78 | 0.28354 | 75339 | 41312 | -1.82 | 0.27821 |
| G3UZY2;P97493 | 41364 | 9927 | -4.17 | 0.29512 | 57548 | 4312 | -13.35 | 0.08128 |
| G3X9L6;Q9DCX2;B1ASE2;D3Z1B2 | 127102 | 67873 | -1.87 | 0.13291 | 78139 | 101829 | 1.30 | 0.42855 |
| G3X9V0;P97372;E0CZ90 | 17413 | 19684 | 1.13 | 0.29065 | 74620 | 7983 | -9.35 | 0.02288 |
| G3X9V2;E9Q8Z5;E9Q8Z6;E9Q8Z8;P30999;E9Q8Z4;E9Q8Z9;D3Z2H2;E9Q903;E9Q901;E9Q904;<br>E9Q986;D3Z7H6;E9Q906;E9Q907;E9Q905;D3Z2H7 | 34454 | 3483 | -9.89 | 0.28229 | 76166 | 21882 | -3.48 | 0.07729 |
| G5E846;G3X981;P15331;REV__A2TJV2;REV__Q6P542;REV__P81122;Q9D4H4 | 492900 | 403175 | -1.22 | 0.31897 | 552010 | 495478 | -1.11 | 0.42867 |
| G5E884;O88643;S4R2K7 | 6947 | 6107 | -1.14 | 0.57544 | 74958 | 20502 | -3.66 | 0.08601 |
| G5E895;D3Z494;S4R2G9 | 12352 | 13816 | 1.12 | 0.25732 | 29708 | 20976 | -1.42 | 0.58969 |

| | | | | | | | | |
|---|---|---|---|---|---|---|---|---|
| G5E8R1 | 17856 | 13732 | -1.30 | 0.06342 | 76704 | 26295 | -2.92 | 0.08433 |
| G5E8T9;Q99KB8;E9PYA3;E9Q2H8;D3YWI0;D3YUX8 | 10572 | 10019 | -1.06 | 0.67265 | 75482 | 21734 | -3.47 | 0.08401 |
| G5E924;Q8R081;G3UY38 | 33364 | 29661 | -1.12 | 0.40578 | 49102 | 18807 | -2.61 | 0.16101 |
| H3BKH6;Q9R0P3;H3BK43;H3BLJ9;H3BJL6;H3BJP2;H3BJC6;H3BL99 | 53797 | 46644 | -1.15 | 0.15501 | 41473 | 22107 | -1.88 | 0.07300 |
| H3BKM0;Q9DBG3;H3BIY9;H3BJ06;Q5SWR0 | 165246 | 150375 | -1.10 | 0.06739 | 114744 | 95589 | -1.20 | 0.55102 |
| H7BX95;Q6PDM2;F7AI47;F6QXN3 | 17766 | 15580 | -1.14 | 0.52563 | 58462 | 5550 | -10.53 | 0.07852 |
| J3QK48;J3QP56;P97823;D3YUG4;D3Z269;D3Z111;J3QQ63 | 13554 | 13008 | -1.04 | 0.53147 | 58239 | 23435 | -2.49 | 0.27492 |
| J3QMM7;K3W4M4;Q9CZ42;J3QN06;J3QPU6 | 37149 | 6065 | -6.13 | 0.26801 | 75507 | 21322 | -3.54 | 0.08253 |
| K3W4R2;Q6URW6 | 15427 | 41524 | 2.69 | 0.28465 | 60475 | 100823 | 1.67 | 0.21138 |
| K4DI76;J3QJW3;Q80TJ1 | 22728 | 25439 | 1.12 | 0.49344 | 61839 | 29882 | -2.07 | 0.27990 |
| O08529 | 36169 | 37257 | 1.03 | 0.81766 | 29094 | 23094 | -1.26 | 0.40287 |
| O08539;Q6P1B9 | 13259 | 11762 | -1.13 | 0.59966 | 39070 | 28589 | -1.37 | 0.66226 |
| O08553 | 2842417 | 2546255 | -1.12 | 0.29039 | 1400167 | 1414582 | 1.01 | 0.96208 |
| O08599;Z4YL99;REV__P45377 | 64508 | 56059 | -1.15 | 0.21021 | 166620 | 109296 | -1.52 | 0.01623 |
| O08749 | 94000 | 93926 | -1.00 | 0.99431 | 54808 | 49755 | -1.10 | 0.78678 |
| O08795 | 35441 | 6248 | -5.67 | 0.30423 | 75102 | 20870 | -3.60 | 0.08554 |
| O08797;Q8VHQ1;Q8BMT0;Q80UK5;O08806;F6V5V4 | 9403 | 17885 | 1.90 | 0.01814 | 74960 | 19666 | -3.81 | 0.08359 |
| O08807;B1AZS9 | 43351 | 47800 | 1.10 | 0.01011 | 82361 | 38454 | -2.14 | 0.03654 |
| O08917;G3UYU4;G3UZZ5;G3UWW8;G3XA73 | 95105 | 56614 | -1.68 | 0.36413 | 69708 | 76280 | 1.09 | 0.86803 |
| O08997 | 43482 | 16544 | -2.63 | 0.29856 | 97519 | 6312 | -15.45 | 0.00001 |
| O09061 | 30476 | 33223 | 1.09 | 0.35190 | 61462 | 10214 | -6.02 | 0.07255 |
| O35215;G3UZN1 | 41122 | 51234 | 1.25 | 0.03400 | 42641 | 12662 | -3.37 | 0.15486 |
| O35350 | 6272 | 6615 | 1.05 | 0.79057 | 75429 | 22572 | -3.34 | 0.08711 |
| O35381;D3YYE1;D3Z7M9;F6UFG6;Q64G17 | 21335 | 23200 | 1.09 | 0.73927 | 59507 | 26141 | -2.28 | 0.28049 |
| O35526;D6RFB9 | 8252 | 8064 | -1.02 | 0.92538 | 39833 | 24756 | -1.61 | 0.57687 |
| O35593 | 67518 | 33106 | -2.04 | 0.41161 | 57286 | 24533 | -2.34 | 0.32212 |
| O35639 | 30705 | 45960 | 1.50 | 0.14468 | 76145 | 9557 | -7.97 | 0.01830 |
| O35685 | 12986 | 14838 | 1.14 | 0.18781 | 75453 | 20714 | -3.64 | 0.08140 |
| O35864;A0A087WQA8;A0A087WQ60 | 9925 | 15150 | 1.53 | 0.05804 | 75699 | 21362 | -3.54 | 0.08069 |
| O35887;G3V004;G3UWV3;G3UXA8;G3UWR0;G3UXA3;G3UY49;G3UYI3 | 42879 | 18006 | -2.38 | 0.34502 | 78203 | 11428 | -6.84 | 0.01332 |
| O54724;Q8CB77 | 46408 | 67634 | 1.46 | 0.19480 | 37670 | 78092 | 2.07 | 0.04184 |
| O54962 | 5224 | 7666 | 1.47 | 0.01221 | 59707 | 28203 | -2.12 | 0.31591 |
| O54983 | 18543 | 19824 | 1.07 | 0.72584 | 61651 | 9696 | -6.36 | 0.06996 |
| O54984 | 39255 | 7764 | -5.06 | 0.30414 | 57595 | 56071 | -1.03 | 0.96048 |
| O55026 | 33234 | 62558 | 1.88 | 0.49682 | 91451 | 113758 | 1.24 | 0.60166 |

| | | | | | | | | |
|---|---|---|---|---|---|---|---|---|
| O55042 | 26008 | 17945 | -1.45 | 0.23423 | 77092 | 25718 | -3.00 | 0.07944 |
| O55125;Q5SVG6 | 63192 | 31954 | -1.98 | 0.45159 | 61487 | 22331 | -2.75 | 0.20835 |
| O55222;F6Q5Z1;D3YZA5 | 34690 | 6250 | -5.55 | 0.31904 | 67671 | 51087 | -1.32 | 0.46563 |
| O55234 | 9223 | 7887 | -1.17 | 0.43589 | 75436 | 21537 | -3.50 | 0.08393 |
| O70209 | 68649 | 11612 | -5.91 | 0.11773 | 60334 | 25037 | -2.41 | 0.19642 |
| O70318;F7BUB8;F7CR30;F6S4K9 | 28951 | 25183 | -1.15 | 0.69992 | 64518 | 46955 | -1.37 | 0.30816 |
| O70400;S4R1V0 | 10909 | 15845 | 1.45 | 0.14492 | 59088 | 24082 | -2.45 | 0.26571 |
| O70435;E0CX62;E0CZ34;F8WH02;E0CYL6 | 27557 | 25028 | -1.10 | 0.15563 | 76957 | 25957 | -2.96 | 0.08080 |
| O70591;F8WJ30 | 35085 | 5033 | -6.97 | 0.29264 | 97519 | 20320 | -4.80 | 0.00344 |
| O88342 | 157813 | 161699 | 1.02 | 0.91068 | 70310 | 101520 | 1.44 | 0.18631 |
| O88398;D3YXP4;D3Z3N6 | 14318 | 15035 | 1.05 | 0.86177 | 76189 | 20377 | -3.74 | 0.07318 |
| O88531;B1B0P8;B1B0P9 | 26990 | 27971 | 1.04 | 0.80502 | 47097 | 17604 | -2.68 | 0.17155 |
| O88543;F6YCA7;D3Z036 | 38107 | 6810 | -5.60 | 0.26029 | 35235 | 21690 | -1.62 | 0.63519 |
| O88544;F6QTS1;D3YV99;D3Z1R9 | 12821 | 17055 | 1.33 | 0.00321 | 57271 | 23517 | -2.44 | 0.29449 |
| O88569 | 177477 | 175484 | -1.01 | 0.88197 | 83914 | 108380 | 1.29 | 0.24448 |
| O88696 | 8934 | 8366 | -1.07 | 0.65283 | 75159 | 54392 | -1.38 | 0.46190 |
| O88712 | 8396 | 10372 | 1.24 | 0.39328 | 58453 | 22830 | -2.56 | 0.26492 |
| O88844;A0A087WPT4;A0A087WRS9;D3YVY3;A0A087WRM4 | 202997 | 179702 | -1.13 | 0.37692 | 69870 | 94684 | 1.36 | 0.26686 |
| O88851 | 16042 | 13650 | -1.18 | 0.07165 | 75824 | 23786 | -3.19 | 0.08656 |
| O88935 | 110071 | 106097 | -1.04 | 0.46928 | 91683 | 90510 | -1.01 | 0.95032 |
| O88958;D3Z0R5 | 38960 | 8010 | -4.86 | 0.26329 | 75068 | 39639 | -1.89 | 0.26336 |
| O89023 | 45766 | 15737 | -2.91 | 0.29531 | 71915 | 20234 | -3.55 | 0.02270 |
| O89053;G3UYK8;D3YW57 | 12112 | 13407 | 1.11 | 0.65845 | 76465 | 25201 | -3.03 | 0.08361 |
| P00493 | 85316 | 79075 | -1.08 | 0.56881 | 74452 | 44751 | -1.66 | 0.07811 |
| P01831 | 65796 | 4350 | -15.13 | 0.09196 | 90470 | 63620 | -1.42 | 0.08919 |
| P03995 | 240524 | 160781 | -1.50 | 0.28397 | 430542 | 559022 | 1.30 | 0.41297 |
| P04117;P24526;O08716 | 117878 | 191753 | 1.63 | 0.01786 | 81951 | 42852 | -1.91 | 0.06766 |
| P05063 | 100428 | 86833 | -1.16 | 0.46901 | 47205 | 18759 | -2.52 | 0.21520 |
| P05064;A6ZI44;D3YWI1;Q9CPQ9;A6ZI46;A6ZI47;D3Z510;D3YV98;Q91Y97 | 1066323 | 929170 | -1.15 | 0.20633 | 322257 | 414735 | 1.29 | 0.54608 |
| P05201;F7ALS6 | 183993 | 188255 | 1.02 | 0.78749 | 94000 | 68287 | -1.38 | 0.10382 |
| P05202 | 315377 | 262078 | -1.20 | 0.58519 | 90321 | 180655 | 2.00 | 0.10703 |
| P05213 | 72367 | 58088 | -1.25 | 0.25348 | 178535 | 108819 | -1.64 | 0.37150 |
| P06151;G5E8N5;D3Z736;D3YZQ9;D3YVR7;D3YZE4;P00342 | 595154 | 492260 | -1.21 | 0.35817 | 382297 | 346862 | -1.10 | 0.46171 |
| P06745;F6SAC3;CON__Q3ZBD7 | 803742 | 814846 | 1.01 | 0.93467 | 1013126 | 439295 | -2.31 | 0.05507 |

| | | | | | | | | |
|---|---|---|---|---|---|---|---|---|
| P06801;Q3TQP6 | 73999 | 73356 | -1.01 | 0.91665 | 50232 | 37654 | -1.33 | 0.11573 |
| P06837;P60761 | 18411 | 13980 | -1.32 | 0.49991 | 79768 | 33354 | -2.39 | 0.07206 |
| P07091;D3YUT9 | 135282 | 111614 | -1.21 | 0.08533 | 182011 | 112075 | -1.62 | 0.30526 |
| P07356;B0V2N5;B0V2N7;B0V2N8;REV__F6WNG1;REV__Q8BND4 | 520311 | 551077 | 1.06 | 0.67966 | 259093 | 236657 | -1.09 | 0.68221 |
| P07724 | 62259 | 61374 | -1.01 | 0.90727 | 67328 | 37164 | -1.81 | 0.24399 |
| P07901;B7ZC50;A2A6A2;B7ZC49;H3BLM8 | 311583 | 272792 | -1.14 | 0.19804 | 96009 | 122665 | 1.28 | 0.29273 |
| P08003 | 42475 | 40384 | -1.05 | 0.67647 | 80000 | 30746 | -2.60 | 0.06163 |
| P08030 | 16298 | 17886 | 1.10 | 0.33366 | 33140 | 28666 | -1.16 | 0.81853 |
| P08113;F7C312 | 229959 | 222019 | -1.04 | 0.62077 | 108379 | 117609 | 1.09 | 0.59421 |
| P08207 | 44646 | 44621 | -1.00 | 0.99734 | 71814 | 24129 | -2.98 | 0.04675 |
| P08228 | 340996 | 298403 | -1.14 | 0.44848 | 35434 | 102800 | 2.90 | 0.03055 |
| P08249 | 824482 | 875187 | 1.06 | 0.62322 | 362758 | 361454 | -1.00 | 0.99156 |
| P08551 | 15822 | 14491 | -1.09 | 0.78094 | 69743 | 17055 | -4.09 | 0.03474 |
| P08553;D3YZ35 | 44133 | 12123 | -3.64 | 0.28317 | 64716 | 12517 | -5.17 | 0.05696 |
| P08752;P50149;A2AE32;P20612;Q3V3I2;F6QPU5 | 56519 | 54839 | -1.03 | 0.85959 | 180099 | 110339 | -1.63 | 0.17967 |
| P09103;E9Q8G8 | 120422 | 117157 | -1.03 | 0.84627 | 79751 | 64312 | -1.24 | 0.38351 |
| P09405 | 39947 | 51166 | 1.28 | 0.27163 | 56770 | 31327 | -1.81 | 0.21358 |
| P09411;S4R2M7;P09041 | 749945 | 689617 | -1.09 | 0.71592 | 417787 | 396319 | -1.05 | 0.60255 |
| P09528 | 76514 | 72689 | -1.05 | 0.64837 | 84639 | 46369 | -1.83 | 0.02872 |
| P09671 | 79214 | 77228 | -1.03 | 0.77760 | 31740 | 21314 | -1.49 | 0.18036 |
| P10107 | 58114 | 70213 | 1.21 | 0.36069 | 85693 | 61992 | -1.38 | 0.47403 |
| P10126;D3YZ68;D3Z3I8 | 249867 | 233586 | -1.07 | 0.30165 | 299053 | 353086 | 1.18 | 0.14823 |
| P10518 | 14586 | 14751 | 1.01 | 0.97098 | 75270 | 54742 | -1.38 | 0.46284 |
| P10605 | 128279 | 119220 | -1.08 | 0.59067 | 91085 | 49802 | -1.83 | 0.16168 |
| P10630;Q8BTU6;E9Q561;D6RJ60;D6RJ11 | 64689 | 70063 | 1.08 | 0.54703 | 65340 | 25974 | -2.52 | 0.13118 |
| P10639 | 65401 | 58769 | -1.11 | 0.31496 | 80600 | 24328 | -3.31 | 0.01737 |
| P10649;A2AE89;F6WHQ7 | 261395 | 309339 | 1.18 | 0.06407 | 229180 | 135835 | -1.69 | 0.09622 |
| P10833 | 4989 | 5269 | 1.06 | 0.74687 | 62566 | 30428 | -2.06 | 0.26631 |
| P10922 | 33858 | 5921 | -5.72 | 0.29388 | 84851 | 44659 | -1.90 | 0.02704 |
| P11031 | 7295 | 9204 | 1.26 | 0.14886 | 57089 | 54778 | -1.04 | 0.94065 |
| P11352 | 11899 | 26310 | 2.21 | 0.01674 | 74799 | 8616 | -8.68 | 0.02297 |
| P11404 | 20088 | 21123 | 1.05 | 0.81906 | 76171 | 20943 | -3.64 | 0.07478 |
| P11499;E9Q3D6;E9PX27;D3Z1R1;E9Q0C3 | 807267 | 779098 | -1.04 | 0.67975 | 224838 | 331764 | 1.48 | 0.19867 |
| P11531;A2A9Z1;A2A9Z2 | 34401 | 38618 | 1.12 | 0.91825 | 38322 | 12643 | -3.03 | 0.28251 |
| P11983;F2Z483 | 34567 | 24474 | -1.41 | 0.00679 | 64651 | 18491 | -3.50 | 0.08055 |

| | | | | | | | | |
|---|---|---|---|---|---|---|---|---|
| P12382 | 10855 | 8466 | -1.28 | 0.34600 | 60173 | 7127 | -8.44 | 0.07042 |
| P12815 | 41588 | 21318 | -1.95 | 0.39803 | 58771 | 8477 | -6.93 | 0.08915 |
| P12970;F6VBB8;V9GX35;F6SSI5 | 3823 | 3503 | -1.09 | 0.84799 | 58956 | 10946 | -5.39 | 0.10130 |
| P13020;CON__Q3SX14 | 179552 | 189334 | 1.05 | 0.16563 | 102136 | 136300 | 1.33 | 0.14445 |
| P13595 | 11977 | 4605 | -2.60 | 0.06222 | 77438 | 24041 | -3.22 | 0.07025 |
| P14069 | 321206 | 82227 | -3.91 | 0.00360 | 276805 | 151962 | -1.82 | 0.30090 |
| P14094 | 20726 | 20269 | -1.02 | 0.95932 | 147282 | 136965 | -1.08 | 0.64932 |
| P14115 | 4641 | 3924 | -1.18 | 0.23392 | 76000 | 8113 | -9.37 | 0.01718 |
| P14131 | 11273 | 8683 | -1.30 | 0.50811 | 61822 | 15678 | -3.94 | 0.09517 |
| P14148;F6XI62;F7CGG2;E9PZA7;E9QLK7;E9PWT1;Q80YV3 | 4583 | 3135 | -1.46 | 0.17746 | 63350 | 11758 | -5.39 | 0.06352 |
| P14152;B1ATQ3 | 383762 | 404523 | 1.05 | 0.54811 | 181666 | 217391 | 1.20 | 0.50981 |
| P14206 | 83015 | 72940 | -1.14 | 0.27134 | 75646 | 32494 | -2.33 | 0.04104 |
| P14211 | 76506 | 90084 | 1.18 | 0.34669 | 70974 | 52865 | -1.34 | 0.46992 |
| P14602;D3YZ06 | 24694 | 37945 | 1.54 | 0.01129 | 91494 | 102860 | 1.12 | 0.50111 |
| P14685;F7B7L8 | 36516 | 31183 | -1.17 | 0.87828 | 58153 | 21216 | -2.74 | 0.25466 |
| P14824 | 648878 | 794514 | 1.22 | 0.00760 | 274286 | 366654 | 1.34 | 0.40052 |
| P14869;E9Q070;S4R1N1;D3YVM5 | 19355 | 20361 | 1.05 | 0.83135 | 42384 | 15847 | -2.67 | 0.24094 |
| P14873 | 136453 | 132987 | -1.03 | 0.75310 | 131080 | 99614 | -1.32 | 0.33104 |
| P15532;Q5NC80 | 15952 | 15091 | -1.06 | 0.48906 | 49432 | 10300 | -4.80 | 0.09537 |
| P15626;D3YX76;Q8R5I6;F6Y363;A2AE91 | 23209 | 41541 | 1.79 | 0.04740 | 56511 | 62630 | 1.11 | 0.84491 |
| P16014 | 13153 | 14544 | 1.11 | 0.69597 | 75611 | 54422 | -1.39 | 0.44803 |
| P16045 | 125720 | 126694 | 1.01 | 0.92501 | 188149 | 133427 | -1.41 | 0.51839 |
| P16125;D3Z7F0;Q8BVP2 | 157753 | 165457 | 1.05 | 0.71264 | 105183 | 79590 | -1.32 | 0.35810 |
| P16330 | 63010 | 32288 | -1.95 | 0.45871 | 52189 | 6392 | -8.17 | 0.06897 |
| P16460;J3QNG0 | 487877 | 380830 | -1.28 | 0.24888 | 1507764 | 779759 | -1.93 | 0.08849 |
| P17156 | 33218 | 32664 | -1.02 | 0.85545 | 19140 | 22764 | 1.19 | 0.13442 |
| P17182;Q6PHC1;B0QZL1;B1ARR7;B1ARR6 | 1528809 | 1403670 | -1.09 | 0.30441 | 708734 | 795678 | 1.12 | 0.36512 |
| P17183;D3Z6E4;D3Z2S4;D3YVD3 | 362855 | 346628 | -1.05 | 0.68891 | 234877 | 164867 | -1.42 | 0.36007 |
| P17426;F6TPX8;F6VSP9;E9PWD0 | 33533 | 33775 | 1.01 | 0.93443 | 70192 | 19106 | -3.67 | 0.00781 |
| P17427 | 113929 | 102465 | -1.11 | 0.01479 | 162365 | 102088 | -1.59 | 0.05662 |
| P17563;Q63836;G3UYY2;D6RHN2;G3UWK0;G3UZT2 | 16474 | 24651 | 1.50 | 0.35763 | 75608 | 23655 | -3.20 | 0.10144 |
| P17742;F8VPN3;V9GXC1;V9GX31 | 1138570 | 1084483 | -1.05 | 0.30736 | 998216 | 617637 | -1.62 | 0.39512 |
| P17751;H7BXC3 | 360693 | 360807 | 1.00 | 0.99594 | 243727 | 190814 | -1.28 | 0.19437 |
| P18242;F6Y6L6;F8WIR1 | 69020 | 70744 | 1.02 | 0.66207 | 72220 | 31009 | -2.33 | 0.00581 |
| P18760;F8WGL3 | 397609 | 406281 | 1.02 | 0.67839 | 254601 | 214043 | -1.19 | 0.40790 |

| | | | | | | | | |
|---|---|---|---|---|---|---|---|---|
| P18872;D3Z2M7;F7BLT7;F6WC15;F6W1B2;A2AE31;S4R196 | 62846 | 59798 | -1.05 | 0.60534 | 137594 | 109252 | -1.26 | 0.22892 |
| P19096 | 52951 | 50169 | -1.06 | 0.82241 | 79069 | 42995 | -1.84 | 0.04801 |
| P19157;P46425;Q8VC73;F6RWR5 | 481567 | 527459 | 1.10 | 0.45953 | 162818 | 208400 | 1.28 | 0.51705 |
| P19324 | 39190 | 15926 | -2.46 | 0.39435 | 56879 | 59363 | 1.04 | 0.81011 |
| P20029 | 197862 | 166391 | -1.19 | 0.29251 | 67259 | 133860 | 1.99 | 0.03521 |
| P20065;Q9D2R9;A2AEH9 | 27367 | 17618 | -1.55 | 0.46692 | 77442 | 28758 | -2.69 | 0.08394 |
| P20108 | 59827 | 61724 | 1.03 | 0.67999 | 76759 | 33155 | -2.32 | 0.04607 |
| P20152;A2AKJ2;CON__Q8N1N4-2;CON__Q7RTT2 | 3855596 | 2871981 | -1.34 | 0.28776 | 4622468 | 4866326 | 1.05 | 0.80975 |
| P21279 | 63596 | 61380 | -1.04 | 0.95633 | 47442 | 37505 | -1.26 | 0.32880 |
| P21300 | 5153 | 40884 | 7.93 | 0.25991 | 75083 | 23501 | -3.19 | 0.10722 |
| P21661 | 38912 | 6375 | -6.10 | 0.25733 | 59615 | 24202 | -2.46 | 0.17386 |
| P21981;G3UXE8 | 57955 | 79461 | 1.37 | 0.06484 | 72277 | 45870 | -1.58 | 0.23788 |
| P22005;B1AZQ0 | 64412 | 62328 | -1.03 | 0.83769 | 177858 | 86491 | -2.06 | 0.02213 |
| P23116 | 4651 | 14362 | 3.09 | 0.00387 | 75164 | 54379 | -1.38 | 0.46159 |
| P23927 | 40015 | 12906 | -3.10 | 0.32047 | 72895 | 59123 | -1.23 | 0.54310 |
| P24369 | 10297 | 13564 | 1.32 | 0.20843 | 80480 | 21202 | -3.80 | 0.01393 |
| P24472;F6SC55;P13745 | 55906 | 61218 | 1.10 | 0.31987 | 42751 | 32960 | -1.30 | 0.57167 |
| P24527 | 53916 | 56504 | 1.05 | 0.68061 | 43879 | 20473 | -2.14 | 0.29500 |
| P24529;Q3UTB3;E9Q9G5;D3Z0B1;F6VPB5 | 43011 | 46337 | 1.08 | 0.63933 | 61759 | 22876 | -2.70 | 0.20787 |
| P24549;O35945 | 170513 | 148677 | -1.15 | 0.35609 | 88863 | 59166 | -1.50 | 0.11719 |
| P25444;D3YVC1;F6YTZ4;J3QMG5;D3YWJ3;E9Q1N8;D3Z536 | 37366 | 29383 | -1.27 | 0.82323 | 64831 | 11239 | -5.77 | 0.05313 |
| P26039;A2AIM2;F6S1V7;F6SX70 | 124769 | 136732 | 1.10 | 0.58807 | 321008 | 288919 | -1.11 | 0.73876 |
| P26040 | 10132 | 9102 | -1.11 | 0.78919 | 56198 | 5659 | -9.93 | 0.03938 |
| P26041 | 363489 | 305768 | -1.19 | 0.35956 | 139475 | 230053 | 1.65 | 0.10537 |
| P26043;Q7TSG6 | 16702 | 13284 | -1.26 | 0.48910 | 75226 | 21764 | -3.46 | 0.08699 |
| P26231 | 18953 | 24759 | 1.31 | 0.42546 | 34189 | 34570 | 1.01 | 0.97489 |
| P26443;F7CFA5;Q9D9C6 | 307320 | 341675 | 1.11 | 0.58447 | 85778 | 145276 | 1.69 | 0.09835 |
| P26516 | 36891 | 4888 | -7.55 | 0.25654 | 57840 | 42280 | -1.37 | 0.63691 |
| P26883;F6X9I3 | 43035 | 47782 | 1.11 | 0.34217 | 79708 | 34783 | -2.29 | 0.07646 |
| P27546;F7CK47;Q78TF3;F6XPV7;F6V4Z1 | 59363 | 64838 | 1.09 | 0.80864 | 68662 | 48439 | -1.42 | 0.32980 |
| P27659;A0A087WNS0;A0A087WQK0 | 9188 | 34671 | 3.77 | 0.34277 | 53542 | 24653 | -2.17 | 0.27127 |
| P27773;F6Q404 | 259065 | 229526 | -1.13 | 0.11382 | 156560 | 188467 | 1.20 | 0.37921 |
| P28271;Q811J3 | 18117 | 22697 | 1.25 | 0.18996 | 77260 | 40327 | -1.92 | 0.22243 |
| P28474;Q64437 | 39563 | 46090 | 1.16 | 0.08418 | 61591 | 30090 | -2.05 | 0.28243 |
| P28663;D6RHL2 | 6687 | 11616 | 1.74 | 0.06159 | 69964 | 29516 | -2.37 | 0.05115 |

| | | | | | | | | |
|---|---|---|---|---|---|---|---|---|
| P28738;P33175;A2ARD4 | 64964 | 1883 | -34.51 | 0.08863 | 56026 | 20506 | -2.73 | 0.28622 |
| P29341;Q9D4E6;F6ZAX1;A2A5N3;Q8C7D3 | 72130 | 59981 | -1.20 | 0.03437 | 65114 | 23594 | -2.76 | 0.10499 |
| P29595 | 30807 | 25660 | -1.20 | 0.16300 | 45958 | 11721 | -3.92 | 0.15826 |
| P29758 | 87119 | 103157 | 1.18 | 0.18515 | 79142 | 32603 | -2.43 | 0.02912 |
| P30275;A2ARP5;B0R0E9;Q6P8J7;B0R0E8;B0R0F0 | 40723 | 37138 | -1.10 | 0.31364 | 64607 | 20277 | -3.19 | 0.09275 |
| P30416;D6RDE2;F6S2D5;F7CAT1 | 30834 | 38576 | 1.25 | 0.41289 | 75670 | 24838 | -3.05 | 0.10492 |
| P31001 | 167809 | 240488 | 1.43 | 0.40704 | 598662 | 1219826 | 2.04 | 0.05306 |
| P31786;Q4VWZ5;D3Z563;M0QWU8 | 224646 | 247976 | 1.10 | 0.37487 | 196542 | 67321 | -2.92 | 0.17895 |
| P31938;E0CYY5;E9QJU1;E2QRQ0;REV__Q62028;REV__A0A0A0MQE3;Q9D413 | 15178 | 14374 | -1.06 | 0.78630 | 42527 | 31733 | -1.34 | 0.64625 |
| P32020 | 6730 | 6669 | -1.01 | 0.97962 | 74618 | 34694 | -2.15 | 0.19372 |
| P32067;A2AR07;D6RI87;F6SXM5 | 6909 | 7038 | 1.02 | 0.94573 | 57474 | 21897 | -2.62 | 0.27364 |
| P32648 | 129572 | 79969 | -1.62 | 0.07260 | 85917 | 61867 | -1.39 | 0.14155 |
| P32921 | 13262 | 15907 | 1.20 | 0.34191 | 56077 | 22435 | -2.50 | 0.23878 |
| P34022;H7BX22 | 10499 | 39337 | 3.75 | 0.26558 | 76424 | 6732 | -11.35 | 0.01433 |
| P34152 | 33726 | 5042 | -6.69 | 0.31972 | 74893 | 39989 | -1.87 | 0.26970 |
| P34884 | 88431 | 104509 | 1.18 | 0.14476 | 70164 | 45861 | -1.53 | 0.25558 |
| P35235 | 41308 | 13318 | -3.10 | 0.30013 | 58267 | 26925 | -2.16 | 0.31421 |
| P35278;Q8C266;A2A5F6;A2A5F5 | 11071 | 13045 | 1.18 | 0.15088 | 55131 | 11160 | -4.94 | 0.06704 |
| P35279;D3YV69 | 36736 | 40120 | 1.09 | 0.39904 | 46971 | 43462 | -1.08 | 0.88049 |
| P35282 | 35842 | 3681 | -9.74 | 0.27664 | 74975 | 21434 | -3.50 | 0.08866 |
| P35486;P35487 | 62887 | 60226 | -1.04 | 0.62716 | 85854 | 43469 | -1.98 | 0.01701 |
| P35564 | 2655 | 1785 | -1.49 | 0.26566 | 62886 | 87711 | 1.39 | 0.37922 |
| P35585;D3YZ71;Q9WVP1;S4R1Q4 | 34011 | 5946 | -5.72 | 0.32846 | 75501 | 40743 | -1.85 | 0.27038 |
| P35979;F8VQK7 | 34944 | 6294 | -5.55 | 0.27784 | 76336 | 6982 | -10.93 | 0.01488 |
| P35980;G3UZJ6;G3UZK4;G3UYV6;G3UX28 | 34856 | 4345 | -8.02 | 0.28691 | 41639 | 12416 | -3.35 | 0.16388 |
| P37804 | 288411 | 480604 | 1.67 | 0.11724 | 909486 | 1704629 | 1.87 | 0.16248 |
| P38647 | 90436 | 84450 | -1.07 | 0.17793 | 54335 | 44651 | -1.22 | 0.65111 |
| P39053;F6W8Z8;D6RH60;F6TH70 | 37873 | 28570 | -1.33 | 0.00673 | 87562 | 48071 | -1.82 | 0.08483 |
| P40124;B1ARS0 | 147447 | 129308 | -1.14 | 0.16201 | 238032 | 89573 | -2.66 | 0.12312 |
| P40142;E0CY51 | 257342 | 220055 | -1.17 | 0.44869 | 180150 | 135975 | -1.32 | 0.49382 |
| P40336 | 7222 | 8955 | 1.24 | 0.49951 | 75413 | 4998 | -15.09 | 0.01627 |
| P42125 | 20881 | 26225 | 1.26 | 0.41588 | 57260 | 20882 | -2.74 | 0.26750 |
| P42208;E9Q3V6;F6WYM0;D3YYB1;D3Z3C0;D3Z1S1;F6UKN5;D3YV76;D3YZU7;G3UYQ0 | 33546 | 30408 | -1.10 | 0.44155 | 67434 | 37462 | -1.80 | 0.15870 |

| | | | | | | | | |
|---|---|---|---|---|---|---|---|---|
| P42669 | 62289 | 42804 | -1.46 | 0.07284 | 47272 | 32385 | -1.46 | 0.42058 |
| P42932;H3BL49;H3BJB6;H3BLL1;H3BKG2;H3BKR8 | 62809 | 56738 | -1.11 | 0.17487 | 30019 | 29957 | -1.00 | 0.99250 |
| P43274 | 48988 | 77946 | 1.59 | 0.11583 | 57816 | 185202 | 3.20 | 0.02210 |
| P43277;I7HFT9;Q07133;P43275 | 36934 | 43038 | 1.17 | 0.87753 | 80606 | 30930 | -2.61 | 0.05784 |
| P45376;D3YVJ7 | 145434 | 127207 | -1.14 | 0.09993 | 57380 | 60575 | 1.06 | 0.85565 |
| P45591 | 34738 | 38729 | 1.11 | 0.23039 | 64184 | 41093 | -1.56 | 0.39164 |
| P45878 | 12057 | 13484 | 1.12 | 0.46321 | 76760 | 9277 | -8.27 | 0.01555 |
| P45952;D6RFD7 | 11255 | 12915 | 1.15 | 0.44873 | 42161 | 11110 | -3.79 | 0.14681 |
| P46096;D3Z7R4 | 43851 | 41453 | -1.06 | 0.75215 | 101983 | 78160 | -1.30 | 0.03567 |
| P46412;D3Z2Y7;D3Z7Y0;F2Z3Y2;P21765 | 5028 | 30180 | 6.00 | 0.28631 | 75208 | 35585 | -2.11 | 0.18729 |
| P46460;G3UX86;G3UX98 | 9033 | 12894 | 1.43 | 0.22664 | 62900 | 14566 | -4.32 | 0.02214 |
| P46638;G3UZD3;G3UZL4 | 28028 | 25021 | -1.12 | 0.53389 | 65628 | 17693 | -3.71 | 0.06799 |
| P46660 | 46658 | 35450 | -1.32 | 0.12503 | 84702 | 54093 | -1.57 | 0.06267 |
| P46935;V9GWV8;Q3V335;V9GXK3 | 65405 | 3375 | -19.38 | 0.09107 | 61152 | 9936 | -6.15 | 0.07471 |
| P47212 | 563660 | 575838 | 1.02 | 0.78461 | 251337 | 120752 | -2.08 | 0.10548 |
| P47738;D3YYF3 | 218187 | 276069 | 1.27 | 0.15428 | 105768 | 108884 | 1.03 | 0.92752 |
| P47754;D6RCW7 | 39952 | 40962 | 1.03 | 0.69584 | 68356 | 32510 | -2.10 | 0.12853 |
| P47791 | 17668 | 18639 | 1.05 | 0.68631 | 76063 | 7209 | -10.55 | 0.01597 |
| P47857 | 67242 | 4880 | -13.78 | 0.09784 | 76984 | 24266 | -3.17 | 0.07538 |
| P47867 | 87261 | 76846 | -1.14 | 0.35356 | 142839 | 65765 | -2.17 | 0.21667 |
| P47911;E9PUX4 | 6477 | 6281 | -1.03 | 0.93106 | 40024 | 21015 | -1.90 | 0.05556 |
| P47962;D3YYV8 | 16754 | 13553 | -1.24 | 0.45492 | 59201 | 26484 | -2.24 | 0.28887 |
| P47963 | 4749 | 32246 | 6.79 | 0.32114 | 58205 | 15353 | -3.79 | 0.08220 |
| P48036 | 1080314 | 1307451 | 1.21 | 0.01354 | 997247 | 554074 | -1.80 | 0.04217 |
| P48428 | 6441 | 5847 | -1.10 | 0.25380 | 74883 | 40129 | -1.87 | 0.27095 |
| P48678;D3YUF7 | 39543 | 32735 | -1.21 | 0.84405 | 122843 | 220293 | 1.79 | 0.26021 |
| P48722;E0CY23;F6SMW7;F6TFH3 | 13038 | 19965 | 1.53 | 0.21552 | 59789 | 9499 | -6.29 | 0.08506 |
| P48758 | 51187 | 57871 | 1.13 | 0.36430 | 66524 | 44996 | -1.48 | 0.38579 |
| P48774;E9PVM7;E9PV63;E9Q024;E9Q5L9;J3QQ61 | 32598 | 40219 | 1.23 | 0.12247 | 63310 | 24929 | -2.54 | 0.14480 |
| P48962;Q3V132 | 34284 | 77280 | 2.25 | 0.35504 | 519546 | 496562 | -1.05 | 0.78813 |
| P49442;E0CYQ4;E0CX64;H3BK26 | 8074 | 9789 | 1.21 | 0.35840 | 56650 | 54604 | -1.04 | 0.94787 |
| P49443;P36993 | 11932 | 15398 | 1.29 | 0.23196 | 76810 | 22465 | -3.42 | 0.07242 |
| P49722 | 33605 | 37541 | 1.12 | 0.02261 | 49388 | 12161 | -4.06 | 0.09271 |
| P49813;Q9JLH8 | 34781 | 4550 | -7.64 | 0.29167 | 75481 | 21567 | -3.50 | 0.08355 |
| P49817;D3Z148;H3BKG0;H3BLK5;D3Z0J2;P51637 | 4871 | 8221 | 1.69 | 0.21557 | 68221 | 49493 | -1.38 | 0.50014 |

| | | | | | | | | |
|---|---|---|---|---|---|---|---|---|
| P50114 | 186554 | 203091 | 1.09 | 0.43262 | 187989 | 79371 | -2.37 | 0.09912 |
| P50247;A2ALT5 | 85916 | 97314 | 1.13 | 0.17896 | 82147 | 19355 | -4.24 | 0.00830 |
| P50396;D6RI86;B7FAU8 | 435174 | 444038 | 1.02 | 0.87033 | 212805 | 194127 | -1.10 | 0.72885 |
| P50428;F6QGM0 | 8505 | 7368 | -1.15 | 0.23200 | 75301 | 35527 | -2.12 | 0.18519 |
| P50516;D3Z1B9;D3YWH3;D3YZ23 | 292320 | 274149 | -1.07 | 0.53162 | 166129 | 167321 | 1.01 | 0.98183 |
| P50518;Q9D593 | 49636 | 40515 | -1.23 | 0.19627 | 49671 | 36011 | -1.38 | 0.04076 |
| P50543 | 55191 | 65414 | 1.19 | 0.08273 | 68579 | 40694 | -1.69 | 0.43426 |
| P50544;B1AR28 | 63379 | 62826 | -1.01 | 0.98997 | 45875 | 34111 | -1.34 | 0.62631 |
| P50580;D3YVH7 | 33853 | 9064 | -3.73 | 0.38595 | 74922 | 21275 | -3.52 | 0.08878 |
| P51150;E0CZ42;D3YXV0;E9PYR6;Q9ESF1 | 54393 | 63367 | 1.16 | 0.13328 | 79457 | 44652 | -1.78 | 0.06214 |
| P51174 | 172369 | 190537 | 1.11 | 0.08748 | 77705 | 57183 | -1.36 | 0.29676 |
| P51410;D3Z629;D3YZT0;S4R1R7 | 33082 | 29704 | -1.11 | 0.92105 | 78252 | 10223 | -7.65 | 0.01161 |
| P51432 | 7442 | 9565 | 1.29 | 0.21114 | 75136 | 54525 | -1.38 | 0.46433 |
| P51855;Q3UEE2;A2AQN9;A2AQN7;H3BKH4 | 32249 | 4586 | -7.03 | 0.30675 | 74742 | 39771 | -1.88 | 0.27135 |
| P51863 | 96727 | 39294 | -2.46 | 0.19894 | 41616 | 32140 | -1.29 | 0.70062 |
| P51880;E9Q0H6 | 26087 | 26737 | 1.02 | 0.83082 | 75566 | 54760 | -1.38 | 0.45365 |
| P52480;E9Q509;G3X925;P53657 | 1320814 | 1361156 | 1.03 | 0.64343 | 776857 | 667445 | -1.16 | 0.20574 |
| P53810;J3QQ30;J3QPW1;F8WGG5 | 29791 | 28366 | -1.05 | 0.80605 | 18503 | 13660 | -1.35 | 0.08453 |
| P53994;Q3TEG7;P59279;G3UXQ7;G3V022 | 24908 | 25212 | 1.01 | 0.96318 | 88365 | 22115 | -4.00 | 0.00128 |
| P54071;D6RIL6 | 110722 | 128807 | 1.16 | 0.36681 | 67750 | 84709 | 1.25 | 0.35578 |
| P54728 | 19973 | 13949 | -1.43 | 0.11238 | 76217 | 21868 | -3.49 | 0.07682 |
| P54775 | 35456 | 32902 | -1.08 | 0.94602 | 56507 | 41275 | -1.37 | 0.64746 |
| P54869 | 34829 | 6450 | -5.40 | 0.31913 | 76133 | 22275 | -3.42 | 0.07867 |
| P54923 | 10336 | 11003 | 1.06 | 0.84006 | 75750 | 41212 | -1.84 | 0.26007 |
| P55264 | 12608 | 17647 | 1.40 | 0.28705 | 75680 | 41950 | -1.80 | 0.26709 |
| P56388 | 7663 | 6039 | -1.27 | 0.17124 | 79601 | 22116 | -3.60 | 0.06527 |
| P56480 | 1195895 | 1614127 | 1.35 | 0.32314 | 2155550 | 1091429 | -1.97 | 0.08900 |
| P56812;D3Z7Q5 | 12279 | 13929 | 1.13 | 0.39570 | 75446 | 22570 | -3.34 | 0.08712 |
| P57722;E9Q7D8;G3UYM5 | 37597 | 5863 | -6.41 | 0.27404 | 57846 | 40265 | -1.44 | 0.59593 |
| P57746 | 7693 | 33545 | 4.36 | 0.25985 | 75205 | 21812 | -3.45 | 0.08746 |
| P57759;F8WJI4;F8WIM7;D6RG87 | 15234 | 13936 | -1.09 | 0.41666 | 58499 | 5483 | -10.67 | 0.07792 |
| P57780;E9Q2W9;D3Z0L8;D3Z761 | 221456 | 225763 | 1.02 | 0.84876 | 128817 | 123425 | -1.04 | 0.74133 |
| P58044;G3XA48;H3BLF8;H3BLP1;H3BKD7 | 23166 | 21699 | -1.07 | 0.77058 | 60141 | 8752 | -6.87 | 0.07730 |
| P58252;G3UXK8 | 148546 | 147487 | -1.01 | 0.91087 | 131540 | 77166 | -1.70 | 0.06708 |
| P58389;A2AWE9;A2AWF0;B7ZDE0;D6RFC2;F6Z6I0 | 18528 | 23656 | 1.28 | 0.22534 | 27300 | 12960 | -2.11 | 0.00299 |

| | | | | | | | | |
|---|---|---|---|---|---|---|---|---|
| P59999;E9PWA7 | 24941 | 29075 | 1.17 | 0.03335 | 81219 | 27845 | -2.92 | 0.01980 |
| P60122;D3YW60;Q05CB6 | 34327 | 4041 | -8.49 | 0.29264 | 56562 | 20565 | -2.75 | 0.27696 |
| P60335 | 16797 | 14694 | -1.14 | 0.38229 | 62186 | 20462 | -3.04 | 0.06463 |
| P60487 | 8155 | 9628 | 1.18 | 0.57827 | 75488 | 54701 | -1.38 | 0.45540 |
| P60521 | 24223 | 19565 | -1.24 | 0.01124 | 47922 | 14255 | -3.36 | 0.13107 |
| P60710;E9Q1F2;E9Q5F4;G3UZ07;E9Q606;F8WGM8;E9Q2D1;F6WX90;E9Q3M9 | 1337006 | 1210998 | -1.10 | 0.22121 | 1806372 | 1145808 | -1.58 | 0.13227 |
| P60766 | 17578 | 15591 | -1.13 | 0.45133 | 49121 | 15717 | -3.13 | 0.08771 |
| P60843 | 4854 | 34559 | 7.12 | 0.27547 | 77651 | 6634 | -11.70 | 0.01063 |
| P60867 | 36395 | 5737 | -6.34 | 0.27758 | 76001 | 22298 | -3.41 | 0.08017 |
| P60879;B0R029;E9Q8A1;B0R030;Q9D3L3;O09044 | 37741 | 28579 | -1.32 | 0.24570 | 45419 | 34557 | -1.31 | 0.64888 |
| P61021 | 5363 | 5518 | 1.03 | 0.73406 | 58682 | 5355 | -10.96 | 0.07567 |
| P61027;D3YUS4 | 4879 | 5947 | 1.22 | 0.24925 | 30652 | 10590 | -2.89 | 0.01743 |
| P61089;A0A087WPV1;Q9CQ37 | 38152 | 26954 | -1.42 | 0.05860 | 69832 | 40377 | -1.73 | 0.10995 |
| P61148;D3Z6H1;D6RCX9;D3Z6E2;D3Z192 | 13928 | 12088 | -1.15 | 0.38258 | 75778 | 21470 | -3.53 | 0.08015 |
| P61161 | 22054 | 24110 | 1.09 | 0.22296 | 41813 | 16323 | -2.56 | 0.26648 |
| P61164 | 38865 | 8774 | -4.43 | 0.27330 | 76565 | 43096 | -1.78 | 0.26452 |
| P61202;A2AQE4 | 18238 | 20086 | 1.10 | 0.17996 | 59085 | 23281 | -2.54 | 0.25844 |
| P61264 | 35006 | 8369 | -4.18 | 0.34682 | 79214 | 28593 | -2.77 | 0.06415 |
| P61358;A2A4Q0 | 38671 | 4106 | -9.42 | 0.23291 | 79788 | 10880 | -7.33 | 0.00800 |
| P61458 | 35728 | 34942 | -1.02 | 0.86183 | 76409 | 10056 | -7.60 | 0.01791 |
| P61750;E9Q798;F6UFB9 | 35673 | 29173 | -1.22 | 0.85392 | 60136 | 39429 | -1.53 | 0.38191 |
| P61922;Q3TUE8;F7C9G3;F6RN86 | 106436 | 122319 | 1.15 | 0.19636 | 77465 | 57496 | -1.35 | 0.28807 |
| P61961;H7BWZ1;D3YW97 | 33999 | 3883 | -8.76 | 0.29669 | 78803 | 39796 | -1.98 | 0.22092 |
| P61971 | 26814 | 27992 | 1.04 | 0.68888 | 31677 | 8844 | -3.58 | 0.04156 |
| P61982 | 223653 | 229171 | 1.02 | 0.73636 | 209456 | 111531 | -1.88 | 0.11427 |
| P62137 | 5836 | 39503 | 6.77 | 0.29182 | 75032 | 35415 | -2.12 | 0.18973 |
| P62141 | 31679 | 38172 | 1.20 | 0.01215 | 56960 | 30928 | -1.84 | 0.17405 |
| P62192 | 9913 | 8187 | -1.21 | 0.49191 | 76554 | 24005 | -3.19 | 0.07908 |
| P62196;Q8K1K2 | 7742 | 6716 | -1.15 | 0.60297 | 76129 | 22019 | -3.46 | 0.07804 |
| P62204;Q3UKW2;Q9D6P8;G3UX57;P20801 | 828519 | 867319 | 1.05 | 0.38477 | 859623 | 382340 | -2.25 | 0.22125 |
| P62242 | 39645 | 6005 | -6.60 | 0.23967 | 64495 | 24123 | -2.67 | 0.12339 |
| P62245;F8WJ41;D3Z712;D3YVB4 | 34184 | 2555 | -13.38 | 0.29150 | 39491 | 10794 | -3.66 | 0.22699 |
| P62259;D6REF3;F6WA09 | 1023737 | 929370 | -1.10 | 0.41535 | 549997 | 444238 | -1.24 | 0.48471 |
| P62264;D3YVF4;D3Z7I1 | 36053 | 32839 | -1.10 | 0.93190 | 75689 | 23748 | -3.19 | 0.08786 |

| | | | | | | | | |
|---|---|---|---|---|---|---|---|---|
| P62301 | 5544 | 6556 | 1.18 | 0.51058 | 45575 | 18896 | -2.41 | 0.21950 |
| P62311 | 3504 | 3307 | -1.06 | 0.55914 | 74662 | 54035 | -1.38 | 0.47291 |
| P62315 | 38428 | 6076 | -6.32 | 0.26187 | 60314 | 9665 | -6.24 | 0.03108 |
| P62320 | 7880 | 7134 | -1.10 | 0.29960 | 42266 | 7753 | -5.45 | 0.14397 |
| P62334 | 8305 | 7467 | -1.11 | 0.65977 | 41077 | 6034 | -6.81 | 0.14517 |
| P62627;A2AVR9 | 12324 | 10656 | -1.16 | 0.16378 | 60728 | 8856 | -6.86 | 0.07289 |
| P62631 | 46392 | 38670 | -1.20 | 0.11543 | 142059 | 54281 | -2.62 | 0.06736 |
| P62702;Q3V1Z5;V9GWY0 | 65865 | 29228 | -2.25 | 0.36730 | 63688 | 32023 | -1.99 | 0.26883 |
| P62737;D3YZY0;D3Z2K3 | 157185 | 199877 | 1.27 | 0.49929 | 107045 | 254083 | 2.37 | 0.31412 |
| P62743 | 15962 | 14675 | -1.09 | 0.50750 | 62983 | 11950 | -5.27 | 0.06658 |
| P62748 | 24213 | 30134 | 1.24 | 0.36346 | 57148 | 21600 | -2.65 | 0.27666 |
| P62751;D3YTY6 | 65397 | 3931 | -16.64 | 0.09349 | 75514 | 21257 | -3.55 | 0.08227 |
| P62774 | 32171 | 33407 | 1.04 | 0.47695 | 78512 | 12941 | -6.07 | 0.01285 |
| P62806 | 14027 | 34291 | 2.44 | 0.11842 | 934471 | 1297580 | 1.39 | 0.19696 |
| P62814;Q91YH6 | 95118 | 87798 | -1.08 | 0.41296 | 85296 | 65554 | -1.30 | 0.33306 |
| P62823 | 30613 | 44577 | 1.46 | 0.07882 | 66228 | 39038 | -1.70 | 0.30207 |
| P62827;Q14AA6;Q61820 | 104372 | 83871 | -1.24 | 0.36445 | 80707 | 42336 | -1.91 | 0.04031 |
| P62843;D3YTQ9 | 68292 | 6704 | -10.19 | 0.08120 | 60572 | 28217 | -2.15 | 0.19128 |
| P62849 | 65553 | 3120 | -21.01 | 0.08883 | 44638 | 24201 | -1.84 | 0.44676 |
| P62852 | 10443 | 9365 | -1.12 | 0.57776 | 36330 | 14389 | -2.52 | 0.30937 |
| P62869;A6PWE0 | 6610 | 7585 | 1.15 | 0.45307 | 55947 | 22685 | -2.47 | 0.24844 |
| P62874;H3BKR2;H3BLF7 | 10152 | 33056 | 3.26 | 0.31061 | 80754 | 41732 | -1.94 | 0.03252 |
| P62880;E9QKR0;D3Z1M1;D3YZX3;D3Z1T4;E9PWM7;V9GWY1 | 32090 | 30224 | -1.06 | 0.58197 | 98540 | 89194 | -1.10 | 0.69247 |
| P62881 | 2909 | 3174 | 1.09 | 0.75031 | 97519 | 34296 | -2.84 | 0.01119 |
| P62900 | 6310 | 4299 | -1.47 | 0.05030 | 61770 | 25592 | -2.41 | 0.23167 |
| P62908;D3YV43 | 11445 | 8988 | -1.27 | 0.55106 | 60055 | 10328 | -5.81 | 0.08597 |
| P62962;Q5SX49;J3QMC2 | 172019 | 221905 | 1.29 | 0.00921 | 138850 | 167165 | 1.20 | 0.61724 |
| P62965;P22935 | 67779 | 3554 | -19.07 | 0.09497 | 75244 | 21017 | -3.58 | 0.08447 |
| P63005;Q5SW16 | 33463 | 29395 | -1.14 | 0.28026 | 78470 | 27607 | -2.84 | 0.06946 |
| P63011;D3YZP5 | 27189 | 35097 | 1.29 | 0.02213 | 65868 | 22697 | -2.90 | 0.10026 |
| P63017;Q504P4;D3Z5E2;D3YW43 | 1338187 | 1306047 | -1.02 | 0.39167 | 573585 | 732927 | 1.28 | 0.14550 |
| P63028;D3YU75 | 61027 | 76949 | 1.26 | 0.08675 | 65631 | 31447 | -2.09 | 0.17626 |
| P63038;D3Z2F2;D3Z7J9 | 525859 | 525996 | 1.00 | 0.99666 | 360791 | 224532 | -1.61 | 0.04648 |
| P63040;D3YZ72 | 21234 | 29115 | 1.37 | 0.38851 | 59261 | 25082 | -2.36 | 0.27428 |
| P63046 | 3451 | 32408 | 9.39 | 0.21663 | 74861 | 20337 | -3.68 | 0.08657 |

| | | | | | | | | |
|---|---|---|---|---|---|---|---|---|
| P63073 | 34950 | 6234 | -5.61 | 0.27607 | 37298 | 4834 | -7.72 | 0.19037 |
| P63085;F6VEI7;E9PXX5;E9Q3I6;Q6P5G0;Q61532 | 19363 | 40623 | 2.10 | 0.39171 | 77633 | 27915 | -2.78 | 0.07938 |
| P63094;Q6R0H7;Z4YKV1;A2A610;Q66L47;Q8CGK7 | 37215 | 5515 | -6.75 | 0.31014 | 47445 | 15704 | -3.02 | 0.15613 |
| P63101;D3YXN6;D3YXF4;D3YW45;A0A087WQC5;A0A087WR62;A0A087WPS7;A0A087WPI2; REV__Q52KR3;Q9QXJ1 | 562016 | 549435 | -1.02 | 0.76217 | 440224 | 288453 | -1.53 | 0.27812 |
| P63158;D3YVC6;D3YZ18;P30681 | 39966 | 11016 | -3.63 | 0.28935 | 82883 | 40920 | -2.03 | 0.05997 |
| P63213 | 37169 | 62969 | 1.69 | 0.55947 | 60588 | 9466 | -6.40 | 0.07646 |
| P63239 | 31742 | 29031 | -1.09 | 0.59302 | 51195 | 53199 | 1.04 | 0.87361 |
| P63268 | 109998 | 122524 | 1.11 | 0.56588 | 399280 | 126311 | -3.16 | 0.13537 |
| P63276 | 39124 | 7796 | -5.02 | 0.25677 | 66548 | 23779 | -2.80 | 0.11569 |
| P63280;Q8CFZ0;G3UYP0;G3UWL6;G3UWJ1;G3UZL6 | 4410 | 4630 | 1.05 | 0.58202 | 74993 | 54292 | -1.38 | 0.46586 |
| P63321 | 5677 | 4694 | -1.21 | 0.38578 | 78640 | 16554 | -4.75 | 0.01599 |
| P63325;Q3UW83 | 10803 | 8976 | -1.20 | 0.29274 | 61274 | 15242 | -4.02 | 0.05695 |
| P63328;P48455 | 35629 | 34981 | -1.02 | 0.88926 | 59332 | 11561 | -5.13 | 0.03247 |
| P63330;P62715 | 27366 | 28776 | 1.05 | 0.45231 | 68583 | 31612 | -2.17 | 0.16673 |
| P67871;G3UZA4;G3UXG7;G3UZJ5;G3UZX4;G3UWU5 | 38449 | 34580 | -1.11 | 0.92128 | 97519 | 5892 | -16.55 | 0.00001 |
| P67984 | 7750 | 8660 | 1.12 | 0.22224 | 75788 | 22027 | -3.44 | 0.08157 |
| P68033 | 3406278 | 3234705 | -1.05 | 0.68746 | 4803441 | 6511095 | 1.36 | 0.05519 |
| P68037;D3YZS3 | 22608 | 14653 | -1.54 | 0.00774 | 60818 | 29550 | -2.06 | 0.30877 |
| P68040 | 33869 | 29921 | -1.13 | 0.64000 | 80124 | 19925 | -4.02 | 0.01366 |
| P68181 | 66673 | 34216 | -1.95 | 0.43847 | 56534 | 22306 | -2.53 | 0.29554 |
| P68369;P05214 | 2115939 | 1713937 | -1.23 | 0.20371 | 3054223 | 2578455 | -1.18 | 0.38961 |
| P68372 | 2399000 | 1765853 | -1.36 | 0.08974 | 4037695 | 3189564 | -1.27 | 0.17399 |
| P68510 | 131480 | 104362 | -1.26 | 0.03460 | 83666 | 51861 | -1.61 | 0.20886 |
| P70168 | 51300 | 57976 | 1.13 | 0.13338 | 90209 | 34418 | -2.62 | 0.01915 |
| P70195 | 40611 | 10604 | -3.83 | 0.26607 | 75457 | 24249 | -3.11 | 0.09228 |
| P70290;B7ZCL8;A2AN84;B7ZCL9;B7ZCM0;D6RFD5;B7ZCM1 | 33112 | 28040 | -1.18 | 0.88825 | 59906 | 22267 | -2.69 | 0.23367 |
| P70296;D3Z1V4;D6RHS6;E9QLE5;Q8VIN1 | 244084 | 235533 | -1.04 | 0.70052 | 123675 | 107937 | -1.15 | 0.60912 |
| P70333;J3QPH6 | 98447 | 3500 | -28.12 | 0.01364 | 58334 | 23979 | -2.43 | 0.29590 |
| P70349;B0R1E3 | 93865 | 79279 | -1.18 | 0.35035 | 68293 | 49194 | -1.39 | 0.43171 |
| P70372 | 9660 | 7377 | -1.31 | 0.14549 | 75149 | 22841 | -3.29 | 0.09117 |
| P70404 | 67763 | 5266 | -12.87 | 0.09509 | 58273 | 7781 | -7.49 | 0.09087 |
| P70460 | 33803 | 4633 | -7.30 | 0.31187 | 55962 | 40505 | -1.38 | 0.64723 |
| P70699;A2AFL5;F6R5R5;F6VEG4;A2AFL3 | 40321 | 47114 | 1.17 | 0.42502 | 42429 | 10202 | -4.16 | 0.16395 |

| | | | | | | | | |
|---|---|---|---|---|---|---|---|---|
| P80313 | 43520 | 35494 | -1.23 | 0.12220 | 37772 | 21810 | -1.73 | 0.18112 |
| P80314 | 60525 | 48896 | -1.24 | 0.02018 | 60537 | 32606 | -1.86 | 0.17139 |
| P80315;G5E839;G3UYW5 | 27349 | 19949 | -1.37 | 0.11662 | 46400 | 15066 | -3.08 | 0.15361 |
| P80316;E0CZA1 | 44394 | 38722 | -1.15 | 0.05992 | 81330 | 25038 | -3.25 | 0.02187 |
| P80317;E9QPA6;Q61390;B1AT05 | 49242 | 37946 | -1.30 | 0.02087 | 40941 | 28069 | -1.46 | 0.10982 |
| P80318;E9Q133;Q3U0I3;F6Q609;F6ZVG8 | 54161 | 45846 | -1.18 | 0.01267 | 80724 | 39772 | -2.03 | 0.03366 |
| P80560;Q3UU93;REV__Q91Z96 | 11436 | 11453 | 1.00 | 0.99680 | 55533 | 39606 | -1.40 | 0.47698 |
| P82347;A2ACH6 | 32357 | 32403 | 1.00 | 0.99900 | 77441 | 25911 | -2.99 | 0.07512 |
| P84078;P61205 | 142641 | 130067 | -1.10 | 0.44898 | 175440 | 100253 | -1.75 | 0.07152 |
| P84089;G3UW85 | 16758 | 13843 | -1.21 | 0.15645 | 76497 | 41901 | -1.83 | 0.24852 |
| P84104;A2A4X6 | 24131 | 21366 | -1.13 | 0.32881 | 48552 | 14599 | -3.33 | 0.12060 |
| P97315 | 27910 | 16877 | -1.65 | 0.21250 | 88568 | 78007 | -1.14 | 0.66219 |
| P97351 | 12801 | 11830 | -1.08 | 0.76807 | 63912 | 16868 | -3.79 | 0.08173 |
| P97379 | 7859 | 5617 | -1.40 | 0.15101 | 75977 | 40623 | -1.87 | 0.25088 |
| P97384;D3Z7U0 | 10140 | 23301 | 2.30 | 0.03744 | 76704 | 25910 | -2.96 | 0.09682 |
| P97427;Q6P1J1 | 263323 | 262435 | -1.00 | 0.97093 | 122818 | 120145 | -1.02 | 0.94907 |
| P97429;D3Z0S1;F7ANV6;S4R1F2 | 30440 | 41540 | 1.36 | 0.08032 | 38442 | 24296 | -1.58 | 0.60349 |
| P97807;H3BKG7 | 85218 | 88498 | 1.04 | 0.84420 | 63961 | 36185 | -1.77 | 0.15333 |
| P99024 | 167401 | 113692 | -1.47 | 0.03447 | 355715 | 224841 | -1.58 | 0.17189 |
| P99026 | 16160 | 9519 | -1.70 | 0.06894 | 54174 | 21670 | -2.50 | 0.26866 |
| P99027 | 34695 | 38074 | 1.10 | 0.54217 | 72765 | 20341 | -3.58 | 0.03004 |
| P99029;G3UZJ4;H3BJQ7 | 110957 | 114267 | 1.03 | 0.69907 | 113454 | 55203 | -2.06 | 0.16323 |
| Q00493 | 103623 | 94032 | -1.10 | 0.21914 | 77069 | 65997 | -1.17 | 0.66642 |
| Q00612;A3KG36;G3UWD6;P97324 | 12224 | 17334 | 1.42 | 0.11784 | 59259 | 20822 | -2.85 | 0.23242 |
| Q00915 | 37034 | 51739 | 1.40 | 0.04358 | 66877 | 11541 | -5.79 | 0.03867 |
| Q01853 | 471527 | 441009 | -1.07 | 0.46504 | 268755 | 190843 | -1.41 | 0.00200 |
| Q02053;P31254 | 220438 | 231465 | 1.05 | 0.63720 | 134523 | 117156 | -1.15 | 0.15646 |
| Q02248;E9Q6A9;F7CRC6;D3YUH4;F7BAC9;F6QZ47;D3Z5Q1;E9PW26;D3Z7S6 | 33766 | 25951 | -1.30 | 0.15935 | 60515 | 32481 | -1.86 | 0.06780 |
| Q02819;H3BK79;D3Z7D7;D3Z1N1 | 19876 | 19034 | -1.04 | 0.79447 | 64292 | 26505 | -2.43 | 0.19629 |
| Q03265;D3Z6F5;D6RJ16;REV__P54729 | 290219 | 334468 | 1.15 | 0.48396 | 600371 | 624255 | 1.04 | 0.81901 |
| Q03517;Q4W8U9 | 674063 | 591322 | -1.14 | 0.18854 | 204406 | 272805 | 1.33 | 0.33338 |
| Q04447 | 956011 | 1055343 | 1.10 | 0.44907 | 1341781 | 902711 | -1.49 | 0.25449 |
| Q04736 | 93870 | 2160 | -43.46 | 0.01284 | 55890 | 19791 | -2.82 | 0.28153 |
| Q05816 | 27772 | 30163 | 1.09 | 0.45995 | 77190 | 11621 | -6.64 | 0.01641 |

| | | | | | | | | |
|---|---|---|---|---|---|---|---|---|
| Q06138 | 42156 | 15114 | -2.79 | 0.30509 | 59404 | 23949 | -2.48 | 0.25838 |
| Q07076 | 27309 | 32127 | 1.18 | 0.16862 | 78780 | 10506 | -7.50 | 0.01030 |
| Q07417;F6RAZ3 | 67494 | 5431 | -12.43 | 0.09815 | 56684 | 40313 | -1.41 | 0.62578 |
| Q08091 | 19322 | 61918 | 3.20 | 0.12394 | 114214 | 579083 | 5.07 | 0.00376 |
| Q08331 | 410321 | 370882 | -1.11 | 0.37330 | 686480 | 336364 | -2.04 | 0.04725 |
| Q0VGU4;A0A087WQ93 | 47201 | 19132 | -2.47 | 0.26650 | 64869 | 19756 | -3.28 | 0.07648 |
| Q11011;E9Q039;F6QYF8;E9Q6F4;F7ANF4;F6V7K3 | 78489 | 82571 | 1.05 | 0.59269 | 81727 | 22501 | -3.63 | 0.01364 |
| Q11136;G3UXC5 | 38195 | 7681 | -4.97 | 0.32268 | 56657 | 3431 | -16.51 | 0.08536 |
| Q3TA40;Q8R0A7;Q8BQB5;D3YUS2 | 34379 | 4628 | -7.43 | 0.30102 | 74923 | 20638 | -3.63 | 0.08680 |
| Q3THE2 | 34794 | 4370 | -7.96 | 0.28866 | 58631 | 31842 | -1.84 | 0.37534 |
| Q3THS6;Q91X83 | 7813 | 7627 | -1.02 | 0.73582 | 75513 | 35804 | -2.11 | 0.18268 |
| Q3TJD7;Q8BVJ7;B8JJB2;B8JJB3;B8JJB1 | 68133 | 48827 | -1.40 | 0.67355 | 58555 | 24283 | -2.41 | 0.23260 |
| Q3TLP8;P63001;P60764;A2AC13;Q05144;G3UZM2;D3Z3L1;F2Z463;D3YX61;Q8R527;Q9ER71 | 35017 | 33152 | -1.06 | 0.79475 | 68996 | 47063 | -1.47 | 0.34185 |
| Q3TML0;Q922R8 | 21579 | 24088 | 1.12 | 0.68968 | 42346 | 11685 | -3.62 | 0.18505 |
| Q3TMU8;O35098;D3Z360 | 52207 | 46737 | -1.12 | 0.57295 | 49526 | 28749 | -1.72 | 0.35785 |
| Q3TPJ8;A2BFF9;A2BFF5;O88487;A2BFF8;D3Z258;Q3TYJ3;D3Z0M6;O88485 | 5153 | 4234 | -1.22 | 0.45251 | 75748 | 22274 | -3.40 | 0.08283 |
| Q3TUE1;Q91WJ8 | 11686 | 14060 | 1.20 | 0.43267 | 74905 | 40636 | -1.84 | 0.27436 |
| Q3TVK3;Q9Z2W0;Q8BPW9;A0A087WS31;A0A087WSE6;A0A087WRC1;A0A087WNX3;<br>A0A087WSD3;A0A087WSU0 | 14551 | 13288 | -1.10 | 0.42262 | 57343 | 23171 | -2.47 | 0.28926 |
| Q3TWV4;P84091 | 34096 | 25312 | -1.35 | 0.15859 | 62797 | 31169 | -2.01 | 0.00336 |
| Q3TXS7;D6RGR5;J3QN38 | 36023 | 6621 | -5.44 | 0.29771 | 37581 | 22432 | -1.68 | 0.58593 |
| Q3U0V1 | 5765 | 4865 | -1.18 | 0.27944 | 74931 | 39905 | -1.88 | 0.26825 |
| Q3U1J4 | 14803 | 17745 | 1.20 | 0.19016 | 57671 | 22681 | -2.54 | 0.27783 |
| Q3U2G2;Q61316 | 131580 | 115349 | -1.14 | 0.50210 | 58532 | 43075 | -1.36 | 0.50297 |
| Q3U4W8;P56399;D3YYA5;D3Z4K7 | 47571 | 57508 | 1.21 | 0.34211 | 101956 | 39729 | -2.57 | 0.01535 |
| Q3U5Q7;REV__Q9WVQ0 | 9090 | 12456 | 1.37 | 0.33161 | 74648 | 54160 | -1.38 | 0.47497 |
| Q3U741;Q501J6 | 6823 | 30148 | 4.42 | 0.32031 | 63915 | 11726 | -5.45 | 0.05957 |
| Q3U7R1 | 6701 | 16785 | 2.50 | 0.14586 | 64379 | 37340 | -1.72 | 0.07429 |
| Q3UDC3;O88746 | 35114 | 30025 | -1.17 | 0.88521 | 75488 | 21516 | -3.51 | 0.08334 |
| Q3UDE2;F2Z423 | 68920 | 8860 | -7.78 | 0.10078 | 75765 | 20211 | -3.75 | 0.07688 |
| Q3UF75;Q9EPC1;D3Z4U2;F8WHJ2;Q8K013 | 69030 | 10159 | -6.79 | 0.10644 | 30953 | 31153 | 1.01 | 0.98262 |
| Q3UGB5;Q9JII5;D3Z4J1 | 9137 | 7211 | -1.27 | 0.13067 | 57184 | 54448 | -1.05 | 0.93000 |
| Q3UGR5;D3YZI3;D3YUN7 | 23714 | 20586 | -1.15 | 0.49081 | 60359 | 8914 | -6.77 | 0.07608 |

| | | | | | | | | |
|---|---|---|---|---|---|---|---|---|
| Q3UHB1 | 35970 | 7277 | -4.94 | 0.30903 | 57438 | 54640 | -1.05 | 0.92790 |
| Q3ULJ0;D3Z0L6 | 8321 | 7169 | -1.16 | 0.62095 | 58306 | 41392 | -1.41 | 0.60399 |
| Q3UM45;F6TGJ2;A0A087WRA7 | 22610 | 22920 | 1.01 | 0.93329 | 38586 | 7568 | -5.10 | 0.20753 |
| Q3UPL0;S4R2A9;S4R192;S4R256;S4R1T5 | 8759 | 9268 | 1.06 | 0.70968 | 76211 | 41014 | -1.86 | 0.24853 |
| Q3UW53;E9PYV4;D3Z233;D3YYZ9 | 62052 | 3039 | -20.42 | 0.09518 | 39567 | 49190 | 1.24 | 0.31542 |
| Q3UW66;Q99J99 | 8373 | 10701 | 1.28 | 0.22313 | 56954 | 54009 | -1.05 | 0.92534 |
| Q3UW96;S4R2N3;Q99MP3;S4R2C5 | 37731 | 47062 | 1.25 | 0.07587 | 52238 | 13300 | -3.93 | 0.10351 |
| Q3UYC0;E0CYP3;Q149T7 | 38854 | 8836 | -4.40 | 0.27447 | 58888 | 38253 | -1.54 | 0.43508 |
| Q3V117;Q91V92;Q3TS02 | 194776 | 159013 | -1.22 | 0.09630 | 135185 | 126020 | -1.07 | 0.74726 |
| Q4KMM3;E9Q0A7;B1H3M0 | 12396 | 15300 | 1.23 | 0.53232 | 75894 | 21890 | -3.47 | 0.08019 |
| Q4VA93;P20444;Q2NKI4;P63318 | 35893 | 31810 | -1.13 | 0.90637 | 65357 | 13912 | -4.70 | 0.05491 |
| Q543K9;P23492;Q9D8C9;Q8QZW3 | 26014 | 42926 | 1.65 | 0.00144 | 57819 | 42867 | -1.35 | 0.64317 |
| Q5EBP8;P49312 | 71152 | 69540 | -1.02 | 0.76307 | 64373 | 41422 | -1.55 | 0.20771 |
| Q5M8N0;F7C0H6 | 9847 | 9035 | -1.09 | 0.69278 | 75457 | 56243 | -1.34 | 0.48657 |
| Q5RKN9;P47753 | 6416 | 6505 | 1.01 | 0.92487 | 75658 | 54868 | -1.38 | 0.45209 |
| Q5SQB0;Q61937;Q9DAY9;Q5SQB5;E9Q5T3 | 19769 | 19099 | -1.04 | 0.26292 | 68874 | 16914 | -4.07 | 0.04156 |
| Q5SRX1;Q5SXA5;Q5SXA4;F6RBX1;F6ZDJ1 | 14149 | 16838 | 1.19 | 0.56031 | 27479 | 11832 | -2.32 | 0.08725 |
| Q5SVG5;Q5SVG4;O35643 | 11139 | 12080 | 1.08 | 0.79503 | 76474 | 21241 | -3.60 | 0.07272 |
| Q5SVI9;Q5SVJ1;Q5SVI3;Q5SVI0;Q5SVI2;Q68EG2;Q5SVI1;Q5SVJ0;P28652 | 35773 | 38564 | 1.08 | 0.94546 | 61288 | 8980 | -6.82 | 0.06811 |
| Q5SW88;Q5SW87;Q5SW86 | 21935 | 25162 | 1.15 | 0.08761 | 53379 | 13860 | -3.85 | 0.06827 |
| Q5SXR6;Q68FD5;F6Z1R4 | 754715 | 646110 | -1.17 | 0.21565 | 799086 | 554350 | -1.44 | 0.05587 |
| Q5UE59;E9Q7C9;Q7TNF4;Q8CD76;O88447;D3YXZ3;Q91YS4;O88448;F6UYN4;Q9DBS5 | 37559 | 5426 | -6.92 | 0.25175 | 58003 | 42550 | -1.36 | 0.63261 |
| Q5XJF6;P53026;D6RE43 | 10631 | 7248 | -1.47 | 0.13967 | 76882 | 12054 | -6.38 | 0.01819 |
| Q60597;Z4YJV4;Q5SVY0;Q5SVY1 | 77628 | 87776 | 1.13 | 0.60326 | 112225 | 85141 | -1.32 | 0.07286 |
| Q60598;Q921L6 | 10523 | 10465 | -1.01 | 0.99077 | 75981 | 55320 | -1.37 | 0.44741 |
| Q60605;Q8CI43 | 155952 | 322740 | 2.07 | 0.29822 | 620523 | 601563 | -1.03 | 0.93858 |
| Q60631;B1AT92;B1AT95 | 11057 | 12042 | 1.09 | 0.67274 | 57461 | 41333 | -1.39 | 0.62424 |
| Q60634;Q5SS83 | 64361 | 64564 | 1.00 | 0.99655 | 97000 | 88283 | -1.10 | 0.64721 |
| Q60692 | 12384 | 9629 | -1.29 | 0.04564 | 75961 | 22309 | -3.40 | 0.08057 |
| Q60737;A2ANR6 | 11281 | 39311 | 3.48 | 0.27591 | 61300 | 24877 | -2.46 | 0.23361 |
| Q60817;P70670 | 25884 | 17793 | -1.45 | 0.21010 | 77751 | 20954 | -3.71 | 0.04410 |
| Q60854;F8WIV2;K7E6F1;E9Q108;E9Q0P9;E9Q3Y1;E9Q6X2;E9PYY0;E9PZQ9;E9Q4R2; E9Q5Q5;Q3UWK8 | 168966 | 165042 | -1.02 | 0.85289 | 149995 | 77970 | -1.92 | 0.08242 |

| | | | | | | | | |
|---|---|---|---|---|---|---|---|---|
| Q60864 | 129705 | 122409 | -1.06 | 0.24974 | 79458 | 41669 | -1.91 | 0.04948 |
| Q60865;F6YLI0 | 36197 | 3204 | -11.30 | 0.24800 | 55921 | 19923 | -2.81 | 0.28229 |
| Q61029 | 5424 | 10558 | 1.95 | 0.17272 | 75323 | 5796 | -13.00 | 0.01731 |
| Q61081 | 12077 | 12154 | 1.01 | 0.96726 | 58095 | 21410 | -2.71 | 0.25762 |
| Q61166 | 82106 | 73713 | -1.11 | 0.17429 | 67979 | 32833 | -2.07 | 0.06122 |
| Q61171;D3Z4A4 | 128713 | 119486 | -1.08 | 0.26863 | 112413 | 78101 | -1.44 | 0.02913 |
| Q61205;D3Z7E6;D3Z2X5;Q8CA83 | 11960 | 16759 | 1.40 | 0.22084 | 75657 | 3531 | -21.43 | 0.01409 |
| Q61206 | 70035 | 67819 | -1.03 | 0.75006 | 50023 | 26405 | -1.89 | 0.01791 |
| Q61425 | 27849 | 27736 | -1.00 | 0.98802 | 48160 | 19980 | -2.41 | 0.19475 |
| Q61548;E9Q9A3;E9QQ05;E9QLK9;Q3TWS4;A0A087WSI9 | 39185 | 40541 | 1.03 | 0.84577 | 43740 | 25146 | -1.74 | 0.18459 |
| Q61553;F7BDR1;D3Z1X1;D3YWW3 | 43514 | 46535 | 1.07 | 0.70375 | 65463 | 21094 | -3.10 | 0.02511 |
| Q61598 | 399964 | 413781 | 1.03 | 0.74662 | 304946 | 226635 | -1.35 | 0.29519 |
| Q61644 | 11819 | 34300 | 2.90 | 0.31455 | 75635 | 36214 | -2.09 | 0.18272 |
| Q61696;P17879 | 20807 | 25089 | 1.21 | 0.26398 | 62254 | 38785 | -1.61 | 0.36119 |
| Q61699;E9Q0U7;D3Z027;D3Z3I9 | 19889 | 18438 | -1.08 | 0.86527 | 60046 | 14055 | -4.27 | 0.10635 |
| Q61753;F6ZSB7 | 32835 | 32398 | -1.01 | 0.91696 | 76316 | 24541 | -3.11 | 0.08420 |
| Q61768;E9QAK5 | 7008 | 3881 | -1.81 | 0.37095 | 76685 | 17976 | -4.27 | 0.04704 |
| Q61792;A2A6H0;A2A6G9;A2A6G6;A2A6G7;A2A6G8;A2A6G5;E9Q0N6;A2A6G0;A2A6G4 | 15136 | 10905 | -1.39 | 0.19572 | 76923 | 25321 | -3.04 | 0.07888 |
| Q62048;D3Z375 | 79405 | 79166 | -1.00 | 0.97813 | 83415 | 43145 | -1.93 | 0.02016 |
| Q62093 | 16489 | 16066 | -1.03 | 0.77480 | 76322 | 42108 | -1.81 | 0.25387 |
| Q62261;A0A0A0MQG2;D3YWH8;Q99MJ6 | 1060589 | 859004 | -1.23 | 0.27062 | 918306 | 959144 | 1.04 | 0.79249 |
| Q62348 | 9481 | 42807 | 4.51 | 0.28221 | 36474 | 22803 | -1.60 | 0.59936 |
| Q62418 | 38566 | 9563 | -4.03 | 0.29091 | 57008 | 21368 | -2.67 | 0.27685 |
| Q62426 | 13649 | 17649 | 1.29 | 0.26426 | 76264 | 25717 | -2.97 | 0.08793 |
| Q62433;E9Q5I8;E9Q3F9;E9PVF3;F6VLR8;E9Q514;E9Q0J8;E9PZC7;E9Q7G8;E9Q147;E9Q7V2 | 24528 | 35482 | 1.45 | 0.00540 | 69435 | 27786 | -2.50 | 0.09373 |
| Q62446 | 35414 | 7082 | -5.00 | 0.31727 | 57016 | 40799 | -1.40 | 0.62605 |
| Q62465 | 219043 | 182243 | -1.20 | 0.35334 | 277815 | 245905 | -1.13 | 0.37027 |
| Q63810;Q63811 | 11327 | 13854 | 1.22 | 0.34987 | 60926 | 21950 | -2.78 | 0.21435 |
| Q63844;D3Z3G6;D3Z6D8 | 70357 | 64149 | -1.10 | 0.52661 | 59108 | 36976 | -1.60 | 0.26330 |
| Q63918 | 38180 | 10737 | -3.56 | 0.31782 | 76057 | 13546 | -5.61 | 0.02424 |
| Q64010 | 12676 | 16330 | 1.29 | 0.33393 | 74816 | 42716 | -1.75 | 0.29367 |
| Q64237 | 66394 | 58145 | -1.14 | 0.84655 | 46291 | 25142 | -1.84 | 0.42515 |
| Q64332 | 108734 | 102191 | -1.06 | 0.17748 | 85455 | 67911 | -1.26 | 0.22278 |

| | | | | | | | | |
|---|---|---|---|---|---|---|---|---|
| Q64433 | 273142 | 305495 | 1.12 | 0.21242 | 100843 | 86494 | -1.17 | 0.37894 |
| Q64471;D3Z3X5;D3Z5W7 | 36727 | 8389 | -4.38 | 0.31018 | 76010 | 23750 | -3.20 | 0.08428 |
| Q64674 | 6342 | 7058 | 1.11 | 0.17969 | 97519 | 20640 | -4.72 | 0.00342 |
| Q64727 | 308503 | 389703 | 1.26 | 0.06519 | 351769 | 347227 | -1.01 | 0.94349 |
| Q64737;D6RCG1 | 4751 | 8839 | 1.86 | 0.00525 | 76739 | 42402 | -1.81 | 0.24730 |
| Q6GT24;O08709;D3Z0Y2;Q8BG37 | 282825 | 290675 | 1.03 | 0.63399 | 144211 | 128931 | -1.12 | 0.53630 |
| Q6IRU2 | 59281 | 71096 | 1.20 | 0.14431 | 57904 | 27194 | -2.13 | 0.05488 |
| Q6IRU5;F7BHJ0 | 35386 | 28019 | -1.26 | 0.83611 | 74978 | 21530 | -3.48 | 0.08892 |
| Q6NZD2;Q9WV80;D3YWH1;REV__A2AJB1 | 21470 | 17854 | -1.20 | 0.29657 | 63449 | 11829 | -5.36 | 0.06266 |
| Q6P069 | 31134 | 27726 | -1.12 | 0.51509 | 58321 | 24293 | -2.40 | 0.28316 |
| Q6P1F6;Q6ZWR4;G3UZP6;G3UXS9;Q8BG02 | 12799 | 10263 | -1.25 | 0.50257 | 21965 | 9803 | -2.24 | 0.02177 |
| Q6P5E4;G3UYG7;G3UZC1;G3UZU8;G3UY73;G3UXP5;E9Q4X2;G3UY35 | 65184 | 8067 | -8.08 | 0.11517 | 32844 | 21266 | -1.54 | 0.66780 |
| Q6P5F9;F6YA11;A2AKT6 | 34526 | 3798 | -9.09 | 0.28536 | 53881 | 20623 | -2.61 | 0.26341 |
| Q6PE15;F6X5P5 | 9054 | 13190 | 1.46 | 0.05641 | 74701 | 58890 | -1.27 | 0.57183 |
| Q6PER3;D3Z6G3;D3YUY6 | 27142 | 19996 | -1.36 | 0.10900 | 48841 | 16766 | -2.91 | 0.14827 |
| Q6WVG3 | 48967 | 54057 | 1.10 | 0.37657 | 70286 | 41040 | -1.71 | 0.13414 |
| Q6ZQ38;D3YWC5 | 67379 | 65210 | -1.03 | 0.86437 | 55762 | 48879 | -1.14 | 0.55337 |
| Q6ZWN5;F7CJS8;D3YWH9;Q9CXW7;D3Z673 | 34345 | 37600 | 1.09 | 0.93719 | 78791 | 10833 | -7.27 | 0.01082 |
| Q6ZWQ5;Q3TGS7;Q3V2H3;O70493 | 4021 | 6720 | 1.67 | 0.10597 | 75637 | 21312 | -3.55 | 0.08116 |
| Q6ZWX6 | 3638 | 3234 | -1.12 | 0.57391 | 74873 | 21148 | -3.54 | 0.08889 |
| Q6ZWY9;Q64525;Q64478;P10854;G3X9D5 | 55938 | 150788 | 2.70 | 0.08018 | 2072305 | 1154162 | -1.80 | 0.27294 |
| Q6ZWZ6;P63323 | 17683 | 10810 | -1.64 | 0.10490 | 39896 | 26513 | -1.50 | 0.61486 |
| Q6ZWZ7;Q9CPR4;B2RY53 | 37987 | 4040 | -9.40 | 0.27663 | 76672 | 9964 | -7.69 | 0.01651 |
| Q70IV5 | 67909 | 48628 | -1.40 | 0.65667 | 19882 | 97235 | 4.89 | 0.01194 |
| Q76MZ3;G3UWL2;H3BLQ0;Q3TTF6;H3BKU1;G3UWS4;Q7TNP2;G3UXQ1;H3BIV7;H3BLE7;H3BJ83;H3BK50 | 117937 | 112339 | -1.05 | 0.56402 | 99257 | 77617 | -1.28 | 0.10571 |
| Q78PY7;Q3TJ56;E9Q3E9 | 8298 | 11202 | 1.35 | 0.33920 | 58323 | 19155 | -3.04 | 0.20692 |
| Q78ZA7 | 17459 | 15910 | -1.10 | 0.76971 | 38787 | 25157 | -1.54 | 0.59223 |
| Q7TMG8;O55126 | 127102 | 9788 | -12.98 | 0.00000 | 40925 | 13186 | -3.10 | 0.22936 |
| Q7TMK9;G3UZ48;G3UZI2;G3V018;G3UXJ6;G3UWM1;G3UXU5;S4R1E9;G3XA76 | 38727 | 10432 | -3.71 | 0.30205 | 37041 | 5493 | -6.74 | 0.20305 |
| Q7TMM9;G3UZR1 | 461372 | 320313 | -1.44 | 0.12608 | 733185 | 593145 | -1.24 | 0.54042 |
| Q7TQD2 | 38678 | 34173 | -1.13 | 0.10675 | 133453 | 32603 | -4.09 | 0.12261 |
| Q7TQF7;S4R2J8;S4R1B8;S4R270;D3Z6Q9 | 19591 | 28793 | 1.47 | 0.14551 | 56448 | 26908 | -2.10 | 0.24825 |
| Q7TSJ2;D3Z6W1 | 96223 | 62493 | -1.54 | 0.16200 | 100139 | 66005 | -1.52 | 0.03232 |

| | | | | | | | | |
|---|---|---|---|---|---|---|---|---|
| Q7TSV4 | 33841 | 3845 | -8.80 | 0.29932 | 35195 | 35494 | 1.01 | 0.99166 |
| Q80TB8;D3Z434 | 178307 | 142279 | -1.25 | 0.17863 | 152984 | 115389 | -1.33 | 0.39043 |
| Q80TL4;S4R1R0;F6SBE4;S4R193;S4R202 | 7576 | 31512 | 4.16 | 0.30057 | 75731 | 21895 | -3.46 | 0.08189 |
| Q80UW2 | 14380 | 15214 | 1.06 | 0.70498 | 74889 | 36715 | -2.04 | 0.20003 |
| Q80VP1;D3Z4V3;D3Z550 | 67312 | 3750 | -17.95 | 0.09258 | 55976 | 35059 | -1.60 | 0.52148 |
| Q80X50 | 33044 | 2584 | -12.79 | 0.29633 | 74879 | 19821 | -3.78 | 0.08487 |
| Q80X90 | 41672 | 48402 | 1.16 | 0.50986 | 73522 | 32806 | -2.24 | 0.10579 |
| Q80XR6;Q20BD0;Q99020 | 38495 | 34666 | -1.11 | 0.30366 | 46632 | 17411 | -2.68 | 0.18589 |
| Q8BFW7 | 65727 | 1867 | -35.20 | 0.09776 | 74584 | 32151 | -2.32 | 0.15158 |
| Q8BG13;O89086;S4R2M6 | 23320 | 21913 | -1.06 | 0.67174 | 61218 | 26414 | -2.32 | 0.24916 |
| Q8BG32;G3UYH2;G3UZ33;G3UWW7;G3UXL5;G3UYI4;G3UYL3;G3UZ28;G3UYL8;G3UX15;G3UX67;G3UWV7 | 67493 | 6007 | -11.24 | 0.08476 | 75707 | 22673 | -3.34 | 0.08438 |
| Q8BGB7 | 39212 | 36559 | -1.07 | 0.93610 | 58941 | 22096 | -2.67 | 0.24906 |
| Q8BGC4 | 12386 | 15102 | 1.22 | 0.18775 | 44580 | 27479 | -1.62 | 0.51444 |
| Q8BGF0;D3Z518;D3Z5E6 | 9168 | 9859 | 1.08 | 0.83130 | 77598 | 22200 | -3.50 | 0.06459 |
| Q8BGJ5;Q922I7;Q8CB58;P17225;E9QMW9;F7DCW4;E9Q0W3;E9Q279;F7C521;F7AXP1 | 13058 | 12149 | -1.07 | 0.78073 | 51211 | 15688 | -3.26 | 0.10127 |
| Q8BGQ7 | 5843 | 7930 | 1.36 | 0.40139 | 75915 | 54977 | -1.38 | 0.44523 |
| Q8BH44;G3UW48;G3UWZ2;G3UYG8 | 6469 | 5371 | -1.20 | 0.03218 | 59845 | 22753 | -2.63 | 0.23957 |
| Q8BH64;D3Z7U7 | 37708 | 10509 | -3.59 | 0.32688 | 57720 | 81999 | 1.42 | 0.45531 |
| Q8BH66 | 64332 | 102914 | 1.60 | 0.40678 | 26858 | 17843 | -1.51 | 0.23477 |
| Q8BH80;Q9QY76 | 9091 | 8996 | -1.01 | 0.96992 | 58457 | 43990 | -1.33 | 0.65274 |
| Q8BH95;F6T930 | 16539 | 17213 | 1.04 | 0.83673 | 61641 | 26868 | -2.29 | 0.24550 |
| Q8BHN3 | 57416 | 58659 | 1.02 | 0.77201 | 64917 | 18377 | -3.53 | 0.08203 |
| Q8BIJ6;E9PWN2;E9PWN3 | 12604 | 18763 | 1.49 | 0.15874 | 75325 | 21618 | -3.48 | 0.08536 |
| Q8BJY1;F7BA91 | 3841 | 4882 | 1.27 | 0.28002 | 76751 | 23680 | -3.24 | 0.07615 |
| Q8BK64;A2AF82;A2AF81;Q8N9S3 | 39045 | 10803 | -3.61 | 0.30107 | 58489 | 23781 | -2.46 | 0.27455 |
| Q8BKC5 | 12566 | 16672 | 1.33 | 0.26512 | 76779 | 24938 | -3.08 | 0.07936 |
| Q8BL97 | 9152 | 7684 | -1.19 | 0.40438 | 76115 | 22245 | -3.42 | 0.07878 |
| Q8BMF4 | 29770 | 19849 | -1.50 | 0.05909 | 73504 | 37524 | -1.96 | 0.04182 |
| Q8BMS1 | 26537 | 31554 | 1.19 | 0.54409 | 275660 | 136613 | -2.02 | 0.12124 |
| Q8BP47 | 8913 | 11473 | 1.29 | 0.21765 | 58042 | 19077 | -3.04 | 0.21069 |
| Q8BQV2;Q03059;REV__Q91VW5;E9Q3P4;Q5SX79 | 15897 | 13977 | -1.14 | 0.63318 | 77712 | 38051 | -2.04 | 0.15290 |
| Q8BT60;Q3UYN2;Q1RLL3;Q9Z140;Q0VE82;Q9DC53;Q8JZW4 | 50013 | 51903 | 1.04 | 0.62329 | 67074 | 25656 | -2.61 | 0.09949 |
| Q8BVE3 | 21849 | 21253 | -1.03 | 0.91161 | 49605 | 19172 | -2.59 | 0.14850 |

| | | | | | | | | |
|---|---|---|---|---|---|---|---|---|
| Q8BVI4;D3YWR7;D3Z1A1;D3Z099 | 52297 | 56322 | 1.08 | 0.22171 | 77810 | 33759 | -2.30 | 0.03528 |
| Q8BVQ5 | 11753 | 11326 | -1.04 | 0.86048 | 56846 | 40431 | -1.41 | 0.62902 |
| Q8BVQ9;P46471 | 6153 | 8195 | 1.33 | 0.28040 | 57412 | 21636 | -2.65 | 0.27208 |
| Q8BWF0 | 38240 | 9668 | -3.96 | 0.30233 | 75123 | 39453 | -1.90 | 0.26086 |
| Q8BWG8;J3QNU6;E0CY53;E0CYB1;F7DF62;Q5F2D9;Q91YI4 | 15620 | 19035 | 1.22 | 0.26743 | 58705 | 41899 | -1.40 | 0.60255 |
| Q8BWT1 | 18496 | 40471 | 2.19 | 0.10483 | 48669 | 27814 | -1.75 | 0.31855 |
| Q8BZF8 | 37445 | 66975 | 1.79 | 0.19795 | 60820 | 80636 | 1.33 | 0.52781 |
| Q8C0M9 | 8898 | 8734 | -1.02 | 0.93563 | 58962 | 23217 | -2.54 | 0.25937 |
| Q8C0P5;B1AVH5 | 33488 | 2456 | -13.64 | 0.28604 | 75038 | 34884 | -2.15 | 0.18679 |
| Q8C166;B7ZCP4;B7ZCP6;E9Q806;B7ZCP7;A3KGK7;B7ZCP5;V9GWY2;<br>F6R587;V9GX86;<br>B7ZCP8;V9GXM6 | 19792 | 25354 | 1.28 | 0.12876 | 46666 | 25851 | -1.81 | 0.42946 |
| Q8C1A5 | 23521 | 20258 | -1.16 | 0.25066 | 57202 | 21723 | -2.63 | 0.27706 |
| Q8C1B7 | 18200 | 14468 | -1.26 | 0.29114 | 42542 | 34054 | -1.25 | 0.70513 |
| Q8C253;P16110 | 93801 | 8161 | -11.49 | 0.01753 | 75710 | 4810 | -15.74 | 0.01502 |
| Q8C2Q7;O35737 | 32514 | 30578 | -1.06 | 0.51032 | 57666 | 24595 | -2.34 | 0.08813 |
| Q8C483;P26638;A2AFS0;A2AFS1 | 15535 | 16488 | 1.06 | 0.84284 | 35935 | 26490 | -1.36 | 0.72192 |
| Q8C522 | 64054 | 30028 | -2.13 | 0.38081 | 75935 | 7679 | -9.89 | 0.01703 |
| Q8C605;Q9WUA3;D3YUA3;F6YL81 | 35424 | 34294 | -1.03 | 0.81005 | 72188 | 34078 | -2.12 | 0.08038 |
| Q8C7J6;Q50H33 | 5690 | 4838 | -1.18 | 0.24872 | 79871 | 5281 | -15.12 | 0.01089 |
| Q8C845;Q9D8Y0;Q9D4J1 | 8263 | 36423 | 4.41 | 0.28905 | 59253 | 22898 | -2.59 | 0.25131 |
| Q8CAQ8;E9Q800;E9QAY6;Q3TZK4;E9PVS5 | 36453 | 75672 | 2.08 | 0.41790 | 49551 | 34171 | -1.45 | 0.50472 |
| Q8CAY6;G3XA25;Q80X81;F2Z459 | 31317 | 32480 | 1.04 | 0.79763 | 79405 | 10231 | -7.76 | 0.00877 |
| Q8CBB7;P22892 | 4224 | 6559 | 1.55 | 0.02827 | 75812 | 34964 | -2.17 | 0.17288 |
| Q8CDN6 | 20009 | 23180 | 1.16 | 0.61599 | 62888 | 8034 | -7.83 | 0.05304 |
| Q8CG76 | 5818 | 36982 | 6.36 | 0.16880 | 74687 | 39855 | -1.87 | 0.27317 |
| Q8CGC7 | 6931 | 4948 | -1.40 | 0.37544 | 60083 | 5131 | -11.71 | 0.06343 |
| Q8CHP8 | 6880 | 9964 | 1.45 | 0.01306 | 57921 | 22885 | -2.53 | 0.27535 |
| Q8CI51;Q9CRA2;F8WJI6;E9Q8P5 | 6394 | 7415 | 1.16 | 0.12403 | 76297 | 6617 | -11.53 | 0.01487 |
| Q8CI94 | 114837 | 111145 | -1.03 | 0.87940 | 108474 | 153334 | 1.41 | 0.08306 |
| Q8CIB5;A6X941 | 15433 | 25904 | 1.68 | 0.00602 | 82975 | 50346 | -1.65 | 0.20697 |
| Q8CIN4;A3KGC4 | 14064 | 9761 | -1.44 | 0.16475 | 75818 | 20793 | -3.65 | 0.07808 |
| Q8JZP2;D3Z620 | 66288 | 2091 | -31.71 | 0.09405 | 77957 | 21734 | -3.59 | 0.06058 |
| Q8JZQ9 | 36187 | 35279 | -1.03 | 0.98052 | 75835 | 20212 | -3.75 | 0.07621 |
| Q8K0T0 | 11527 | 38247 | 3.32 | 0.30620 | 51560 | 25800 | -2.00 | 0.31303 |

| | | | | | | | | |
|---|---|---|---|---|---|---|---|---|
| Q8K0U4 | 67869 | 5293 | -12.82 | 0.09439 | 45915 | 25491 | -1.80 | 0.42661 |
| Q8K183;D3Z7R1 | 68736 | 69212 | 1.01 | 0.93217 | 27671 | 34072 | 1.23 | 0.29550 |
| Q8K1M3;P12367 | 26059 | 25576 | -1.02 | 0.89818 | 36562 | 18633 | -1.96 | 0.16989 |
| Q8K2B3 | 28224 | 35972 | 1.27 | 0.36034 | 51292 | 32734 | -1.57 | 0.35192 |
| Q8K310;A0A087WSU2;A0A087WSP7;A0A087WQZ1;A0A087WQD6;A0A087WQ54; A0A087WQP4;A0A087WNP3;A0A087WPU5;A0A087WPQ5;A0A087WNW7 | 35314 | 8158 | -4.33 | 0.33729 | 48417 | 8067 | -6.00 | 0.08653 |
| Q8K3H0 | 6314 | 5832 | -1.08 | 0.77695 | 50653 | 39630 | -1.28 | 0.71189 |
| Q8K4Z3 | 24919 | 33511 | 1.34 | 0.00376 | 59648 | 9461 | -6.30 | 0.08555 |
| Q8QZS1;E0CX19 | 26759 | 27562 | 1.03 | 0.82726 | 60549 | 10179 | -5.95 | 0.08068 |
| Q8QZT1 | 108069 | 105898 | -1.02 | 0.85461 | 49992 | 65630 | 1.31 | 0.25443 |
| Q8QZY1 | 36554 | 8447 | -4.33 | 0.31494 | 75949 | 41081 | -1.85 | 0.25467 |
| Q8R016;E9PY26;E9Q6V3;E9PZH4;E9QA53 | 12252 | 16644 | 1.36 | 0.22869 | 79235 | 60463 | -1.31 | 0.51235 |
| Q8R086 | 10278 | 8455 | -1.22 | 0.13434 | 75954 | 55358 | -1.37 | 0.44877 |
| Q8R146 | 11443 | 16540 | 1.45 | 0.17798 | 75102 | 40214 | -1.87 | 0.26684 |
| Q8R164 | 5247 | 6204 | 1.18 | 0.51895 | 75162 | 43607 | -1.72 | 0.31978 |
| Q8R1F1;A2ARS6 | 66917 | 32482 | -2.06 | 0.43117 | 56606 | 23378 | -2.42 | 0.32238 |
| Q8R1G2;E0CXH4;E0CXT6 | 15130 | 12324 | -1.23 | 0.41588 | 78776 | 42391 | -1.86 | 0.24563 |
| Q8R1I2;V9GX72 | 42178 | 48004 | 1.14 | 0.70371 | 67458 | 22369 | -3.02 | 0.07444 |
| Q8R317 | 15466 | 17609 | 1.14 | 0.40711 | 76157 | 42648 | -1.79 | 0.26194 |
| Q8R3B1 | 13501 | 21839 | 1.62 | 0.04781 | 57852 | 23085 | -2.51 | 0.27870 |
| Q8R3P0;V9GXG0;B0QZP3;D6RJ20 | 8004 | 8058 | 1.01 | 0.95936 | 75357 | 40364 | -1.87 | 0.26238 |
| Q8R4N0 | 7029 | 15295 | 2.18 | 0.21648 | 75264 | 60959 | -1.23 | 0.61491 |
| Q8R570;B2FDF6;I7HPB1;I7HFU4;J3JS29 | 93620 | 1441 | -64.96 | 0.01301 | 56535 | 20987 | -2.69 | 0.28168 |
| Q8R5C5;E0CZD4;E0CYB4 | 23680 | 20716 | -1.14 | 0.34383 | 57134 | 19975 | -2.86 | 0.11834 |
| Q8R5L1;O35658 | 77421 | 77691 | 1.00 | 0.97889 | 92490 | 38182 | -2.42 | 0.16169 |
| Q8VCT3;E9PYF1 | 2995 | 35100 | 11.72 | 0.23990 | 74954 | 34064 | -2.20 | 0.18409 |
| Q8VCT4;P23953;D3Z5G7;Q8VCC2;D3Z298;Q91WU0;H3BL34;E9PYP1;Q64176 | 19609 | 60287 | 3.07 | 0.22257 | 41333 | 42622 | 1.03 | 0.96425 |
| Q8VD37;A2AIY5 | 13585 | 13359 | -1.02 | 0.94789 | 60887 | 25384 | -2.40 | 0.24636 |
| Q8VDD5;Q8BXF2;F2Z494 | 34000 | 48726 | 1.43 | 0.72089 | 82697 | 148693 | 1.80 | 0.17989 |
| Q8VDK1;D3YY53;D3Z2Y2;D3Z3I3 | 37008 | 8190 | -4.52 | 0.30136 | 75256 | 40991 | -1.84 | 0.26923 |
| Q8VDM4;J3KMQ2 | 13089 | 10482 | -1.25 | 0.21384 | 34564 | 7887 | -4.38 | 0.10122 |
| Q8VDN2;Q91WH7;E9QNX7;Q64436 | 33189 | 23810 | -1.39 | 0.21313 | 505229 | 553981 | 1.10 | 0.69363 |
| Q8VDQ1;Q3TXN1;D6RGL6 | 8114 | 12844 | 1.58 | 0.04520 | 35335 | 40763 | 1.15 | 0.85767 |

| | | | | | | | | |
|---|---|---|---|---|---|---|---|---|
| Q8VEH3;F6QKK2 | 36985 | 31653 | -1.17 | 0.87726 | 43264 | 33707 | -1.28 | 0.69391 |
| Q8VEK3;G3XA10 | 16088 | 10176 | -1.58 | 0.11112 | 85996 | 30741 | -2.80 | 0.00545 |
| Q8VHM5;F7B5B5;A2AW41;V9GWW3;A2AW40 | 5121 | 6957 | 1.36 | 0.27348 | 57290 | 21739 | -2.64 | 0.27533 |
| Q8VIJ6 | 9701 | 8554 | -1.13 | 0.80812 | 65057 | 13180 | -4.94 | 0.05946 |
| Q91V41;A2AL34 | 13696 | 20586 | 1.50 | 0.06879 | 60553 | 14702 | -4.12 | 0.10500 |
| Q91V64 | 16637 | 18401 | 1.11 | 0.35628 | 59753 | 8598 | -6.95 | 0.02741 |
| Q91V76 | 6957 | 8444 | 1.21 | 0.10035 | 57448 | 21773 | -2.64 | 0.27278 |
| Q91V77;P56565;D3YUT6 | 50883 | 51938 | 1.02 | 0.93333 | 68964 | 17096 | -4.03 | 0.04211 |
| Q91VA7;V9GXV0 | 15663 | 16190 | 1.03 | 0.87467 | 45909 | 15438 | -2.97 | 0.16637 |
| Q91VB8;P01942;A7M7S6;P06467 | 246226 | 370651 | 1.51 | 0.08667 | 80196 | 73916 | -1.08 | 0.79448 |
| Q91VD9;A0A087WSU3;A0A087WQR0;A0A087WR47;A0A087WP77 | 95400 | 68545 | -1.39 | 0.55690 | 44505 | 29482 | -1.51 | 0.48523 |
| Q91VI7 | 13538 | 16411 | 1.21 | 0.25801 | 76764 | 24016 | -3.20 | 0.07696 |
| Q91VM9;D3Z636;G8JL76;D3Z096 | 13560 | 19669 | 1.45 | 0.42832 | 57008 | 40144 | -1.42 | 0.61490 |
| Q91VR5 | 39348 | 8051 | -4.89 | 0.25431 | 62101 | 28775 | -2.16 | 0.25592 |
| Q91VR7;J3QN16 | 29200 | 25110 | -1.16 | 0.11541 | 80903 | 14350 | -5.64 | 0.00768 |
| Q91VW3;I7HPY0 | 15185 | 16668 | 1.10 | 0.55597 | 77361 | 25054 | -3.09 | 0.07352 |
| Q91WD9 | 55496 | 54420 | -1.02 | 0.89074 | 109884 | 31277 | -3.51 | 0.02824 |
| Q91WK2 | 34671 | 3655 | -9.48 | 0.28063 | 75109 | 40508 | -1.85 | 0.26885 |
| Q91WU5 | 6869 | 8876 | 1.29 | 0.27402 | 75799 | 56012 | -1.35 | 0.46254 |
| Q91X97;D3YVA2;D3Z2Z8;D3Z1M0 | 52603 | 51513 | -1.02 | 0.86701 | 61228 | 26752 | -2.29 | 0.08633 |
| Q91XH5;Q64105;G3UXX3;G3UZ79 | 61000 | 62890 | 1.03 | 0.66642 | 64799 | 26719 | -2.43 | 0.13361 |
| Q91YI0;E0CXM2;E0CY49;F7D439;E0CYV3 | 128805 | 122658 | -1.05 | 0.71202 | 86621 | 60902 | -1.42 | 0.02558 |
| Q91YR1;D3Z2H0 | 35824 | 8608 | -4.16 | 0.33286 | 75359 | 4311 | -17.48 | 0.01576 |
| Q91ZA3;H3BL62;H3BK61;D3YWM4;D3YZC0;H3BKW6 | 5915 | 19446 | 3.29 | 0.04811 | 75583 | 60542 | -1.25 | 0.59740 |
| Q91ZJ5 | 14518 | 14818 | 1.02 | 0.79699 | 76267 | 42914 | -1.78 | 0.26117 |
| Q920E5 | 77315 | 79482 | 1.03 | 0.70173 | 79524 | 33985 | -2.34 | 0.12470 |
| Q921F2;A0A087WR97;A0A087WRZ5;Q8R0B4;Q8BLD4;Q6VYI5;Q6VYI4; A0A087WQA5; A0A087WQX8;A0A087WNY6;A0A087WS74;A0A087WRP4;H3BJV1;A0A087WSE4; A0A087WSH7;A0A087WS17;A0A087WP57;A0A087WSC6 | 12091 | 13954 | 1.15 | 0.12099 | 51843 | 11227 | -4.62 | 0.07915 |
| Q921H8;H3BKL5;H3BJZ9;H3BKA1;H3BJC1 | 34299 | 32571 | -1.05 | 0.96377 | 75286 | 21424 | -3.51 | 0.08520 |
| Q921M7 | 18951 | 26252 | 1.39 | 0.19329 | 62609 | 7098 | -8.82 | 0.05169 |
| Q922B2;Q8BJY7 | 6652 | 7085 | 1.07 | 0.76565 | 79218 | 26123 | -3.03 | 0.05859 |
| Q924B0;O55023;Q80ZJ2;D3Z703 | 24873 | 21786 | -1.14 | 0.21068 | 61236 | 23675 | -2.59 | 0.22374 |
| Q924M7;F6Q3K8 | 35450 | 5809 | -6.10 | 0.29691 | 74800 | 39491 | -1.89 | 0.26803 |

| | | | | | | | | |
|---|---|---|---|---|---|---|---|---|
| Q93092 | 119316 | 107371 | -1.11 | 0.39339 | 46795 | 23065 | -2.03 | 0.29493 |
| Q99JA0;P70160 | 9879 | 10038 | 1.02 | 0.91047 | 56920 | 74819 | 1.31 | 0.51952 |
| Q99JF8;F6RB63;A2BI12 | 34469 | 4816 | -7.16 | 0.30144 | 39859 | 6580 | -6.06 | 0.16978 |
| Q99JI6 | 13365 | 13196 | -1.01 | 0.94915 | 97318 | 47105 | -2.07 | 0.13239 |
| Q99JY0;D3YXU1 | 4540 | 9568 | 2.11 | 0.01757 | 70021 | 52684 | -1.33 | 0.37353 |
| Q99JY9;A0A087WRA1;A0A087WP86;A0A087WS98;A0A087WQ14;Q641P0;A0A087WQ83 | 35378 | 40009 | 1.13 | 0.20444 | 66539 | 37614 | -1.77 | 0.36200 |
| Q99K48;B1AXT0 | 34875 | 40517 | 1.16 | 0.88942 | 76054 | 10224 | -7.44 | 0.01935 |
| Q99K70;B1AWT2;B1AWT3;Q7TT45;B1AWT4 | 2494 | 2887 | 1.16 | 0.53376 | 74815 | 39571 | -1.89 | 0.26829 |
| Q99K85;Q3U6K9;E9Q6P1 | 11337 | 16516 | 1.46 | 0.15027 | 74721 | 19498 | -3.83 | 0.08562 |
| Q99KC8;F6TIL5 | 68514 | 60696 | -1.13 | 0.15496 | 49189 | 31923 | -1.54 | 0.35838 |
| Q99KE1 | 28340 | 30376 | 1.07 | 0.79127 | 77865 | 34984 | -2.23 | 0.13812 |
| Q99KI0 | 370340 | 361936 | -1.02 | 0.82951 | 214917 | 157085 | -1.37 | 0.31992 |
| Q99KJ8 | 39903 | 30057 | -1.33 | 0.00436 | 80234 | 13371 | -6.00 | 0.00910 |
| Q99L13 | 30273 | 36960 | 1.22 | 0.26569 | 40838 | 12673 | -3.22 | 0.22577 |
| Q99L88 | 35985 | 63945 | 1.78 | 0.50595 | 62548 | 11540 | -5.42 | 0.06945 |
| Q99LB4;P24452;D3YTL5;D3YU77;D3YZN3;D3Z014;D3Z4K5 | 19572 | 27109 | 1.39 | 0.22499 | 66614 | 14105 | -4.72 | 0.04976 |
| Q99LB6;E0CYU5 | 34581 | 4409 | -7.84 | 0.29320 | 74813 | 34530 | -2.17 | 0.18913 |
| Q99LC3;A0A087WR38 | 127102 | 38582 | -3.29 | 0.02426 | 34840 | 22889 | -1.52 | 0.25937 |
| Q99LC5 | 49206 | 52647 | 1.07 | 0.69893 | 112757 | 47238 | -2.39 | 0.08632 |
| Q99LD8;O08972;G3UZR0 | 51818 | 44894 | -1.15 | 0.24205 | 78982 | 25812 | -3.06 | 0.06207 |
| Q99LF4 | 8037 | 9018 | 1.12 | 0.52007 | 77933 | 26200 | -2.97 | 0.07065 |
| Q99LT0 | 6721 | 34345 | 5.11 | 0.30834 | 39938 | 74819 | 1.87 | 0.08262 |
| Q99LX0;A2A813;A2A815;A2A817;A2A816 | 98216 | 85535 | -1.15 | 0.11896 | 121683 | 55997 | -2.17 | 0.13293 |
| Q99M71 | 34347 | 29126 | -1.18 | 0.19265 | 63480 | 25328 | -2.51 | 0.19978 |
| Q99MK8;Q7TS64;E9PW16;F6QY34;E9Q419;F7AEX1;F6Y9P3 | 32726 | 68450 | 2.09 | 0.43747 | 57766 | 5094 | -11.34 | 0.08275 |
| Q99MN1;Q8R2P8 | 10940 | 15609 | 1.43 | 0.32285 | 76652 | 5287 | -14.50 | 0.01248 |
| Q99MN9;E9Q1J7;D3YZC1;A0A087WQV1 | 11046 | 21425 | 1.94 | 0.08733 | 58253 | 42351 | -1.38 | 0.62228 |
| Q99N15;A2AFQ2;O08756 | 67464 | 6212 | -10.86 | 0.08607 | 32077 | 21953 | -1.46 | 0.00312 |
| Q99P58 | 67157 | 3614 | -18.58 | 0.09326 | 37289 | 6190 | -6.02 | 0.16527 |
| Q99P72;Q8BHF5 | 11482 | 10851 | -1.06 | 0.65910 | 45300 | 18358 | -2.47 | 0.23136 |
| Q99PT1 | 251973 | 259054 | 1.03 | 0.80037 | 104541 | 100509 | -1.04 | 0.82389 |
| Q9CPS5;Q9CX56 | 62982 | 4575 | -13.77 | 0.09374 | 75159 | 17570 | -4.28 | 0.05862 |
| Q9CPU0 | 64094 | 73282 | 1.14 | 0.02565 | 72573 | 31529 | -2.30 | 0.06333 |
| Q9CPV4;E9Q197;F6ZTG3;F7BB55;E9Q055;E9Q2R6 | 42096 | 53413 | 1.27 | 0.01735 | 81289 | 19645 | -4.14 | 0.01056 |

| | | | | | | | | |
|---|---|---|---|---|---|---|---|---|
| Q9CPW4;Q3UA72;E9Q2K4 | 11133 | 10350 | -1.08 | 0.64113 | 76174 | 41916 | -1.82 | 0.25581 |
| Q9CPX4;P29391;P49945;H3BKD3 | 38352 | 43552 | 1.14 | 0.20250 | 70401 | 31860 | -2.21 | 0.09028 |
| Q9CPY7 | 16163 | 16436 | 1.02 | 0.89995 | 52376 | 21751 | -2.41 | 0.30555 |
| Q9CQ19;D3Z249 | 183465 | 307403 | 1.68 | 0.34972 | 623617 | 668329 | 1.07 | 0.83786 |
| Q9CQ60;Q8CBG6;D3Z4X1;F6X8L5 | 13009 | 18872 | 1.45 | 0.33581 | 37804 | 11330 | -3.34 | 0.09674 |
| Q9CQ62 | 14086 | 12870 | -1.09 | 0.64086 | 63977 | 26873 | -2.38 | 0.20472 |
| Q9CQ65 | 36795 | 8236 | -4.47 | 0.30718 | 75756 | 23926 | -3.17 | 0.08793 |
| Q9CQ92;G3X9U9;D3YZ32 | 95731 | 2519 | -38.00 | 0.01328 | 46337 | 10712 | -4.33 | 0.05946 |
| Q9CQA3 | 35486 | 41863 | 1.18 | 0.87403 | 66012 | 22027 | -3.00 | 0.08375 |
| Q9CQD1 | 4477 | 4832 | 1.08 | 0.66997 | 57500 | 35495 | -1.62 | 0.49045 |
| Q9CQF3 | 36487 | 6217 | -5.87 | 0.28286 | 74939 | 40023 | -1.87 | 0.26895 |
| Q9CQI3;D3YY93;D3Z2F6;D3YY16;Q9ERL7 | 20271 | 18174 | -1.12 | 0.53108 | 59449 | 37769 | -1.57 | 0.47644 |
| Q9CQI6 | 38563 | 33963 | -1.14 | 0.29629 | 45627 | 16646 | -2.74 | 0.19360 |
| Q9CQR2 | 13314 | 9626 | -1.38 | 0.03557 | 75519 | 36127 | -2.09 | 0.18424 |
| Q9CQR4 | 37059 | 10861 | -3.41 | 0.34388 | 97519 | 19963 | -4.88 | 0.00100 |
| Q9CQV6;M0QWC2 | 42933 | 36502 | -1.18 | 0.84178 | 60753 | 26634 | -2.28 | 0.27811 |
| Q9CQV8;A2A5N1 | 134512 | 144734 | 1.08 | 0.44818 | 112539 | 70063 | -1.61 | 0.12007 |
| Q9CQW1 | 3398 | 28394 | 8.36 | 0.29929 | 75433 | 35361 | -2.13 | 0.18185 |
| Q9CQZ1 | 36933 | 8322 | -4.44 | 0.30497 | 63055 | 56198 | -1.12 | 0.74173 |
| Q9CR16;A2BGI9;Q99KR7 | 18619 | 19701 | 1.06 | 0.80505 | 77141 | 22459 | -3.43 | 0.06922 |
| Q9CR51 | 8624 | 36922 | 4.28 | 0.28469 | 76115 | 7416 | -10.26 | 0.01608 |
| Q9CR57 | 3843 | 2482 | -1.55 | 0.02573 | 58196 | 7195 | -8.09 | 0.08850 |
| Q9CR86 | 34941 | 4076 | -8.57 | 0.28143 | 74871 | 40007 | -1.87 | 0.27030 |
| Q9CRB6 | 503613 | 530697 | 1.05 | 0.67629 | 329059 | 318154 | -1.03 | 0.91897 |
| Q9CRC9;D3YWR1;G3UXF5;D6RCJ1 | 7892 | 10919 | 1.38 | 0.20483 | 75463 | 40549 | -1.86 | 0.26140 |
| Q9CS42 | 12582 | 12137 | -1.04 | 0.82284 | 36985 | 40733 | 1.10 | 0.86253 |
| Q9CVB6;D3YXG6;A0A087WRT2 | 25461 | 31057 | 1.22 | 0.35994 | 51839 | 15984 | -3.24 | 0.13269 |
| Q9CWF2 | 27201 | 17524 | -1.55 | 0.06152 | 81182 | 36843 | -2.20 | 0.06601 |
| Q9CWJ9;REV__B2KFW1 | 45108 | 43818 | -1.03 | 0.84542 | 65340 | 19411 | -3.37 | 0.08442 |
| Q9CWK8 | 64814 | 2326 | -27.87 | 0.09162 | 74811 | 20049 | -3.73 | 0.08625 |
| Q9CWS0;D3YU15 | 77040 | 67324 | -1.14 | 0.63096 | 77472 | 27163 | -2.85 | 0.07891 |
| Q9CX80 | 23345 | 22480 | -1.04 | 0.85476 | 75871 | 4675 | -16.23 | 0.01436 |
| Q9CXW3 | 10209 | 10909 | 1.07 | 0.76379 | 57477 | 24172 | -2.38 | 0.31431 |
| Q9CXY6 | 10381 | 11724 | 1.13 | 0.17878 | 56906 | 20714 | -2.75 | 0.27220 |
| Q9CY64;A2ASB1;A2ASB8;A2ASB7 | 11681 | 18195 | 1.56 | 0.00651 | 57404 | 34809 | -1.65 | 0.48253 |

| | | | | | | | | |
|---|---|---|---|---|---|---|---|---|
| Q9CZ13 | 10982 | 60467 | 5.51 | 0.14153 | 81489 | 53117 | -1.53 | 0.09357 |
| Q9CZ30;B1AYJ9 | 16865 | 23269 | 1.38 | 0.04151 | 53931 | 24998 | -2.16 | 0.31330 |
| Q9CZ44;A2AT02 | 18905 | 22756 | 1.20 | 0.50046 | 77457 | 21733 | -3.56 | 0.06469 |
| Q9CZC8;D3YXW0 | 44219 | 36124 | -1.22 | 0.15156 | 71789 | 20914 | -3.43 | 0.03476 |
| Q9CZD3 | 12871 | 9845 | -1.31 | 0.53835 | 75214 | 39849 | -1.89 | 0.26173 |
| Q9CZS1;G3UYH1 | 18307 | 48561 | 2.65 | 0.19704 | 55932 | 42848 | -1.31 | 0.69242 |
| Q9CZT8;A2A7Z6 | 8174 | 8427 | 1.03 | 0.80308 | 39463 | 40794 | 1.03 | 0.96437 |
| Q9CZU6 | 204860 | 250901 | 1.22 | 0.18784 | 233374 | 148026 | -1.58 | 0.04477 |
| Q9CZW5 | 32612 | 66055 | 2.03 | 0.45588 | 77405 | 6084 | -12.72 | 0.01095 |
| Q9D031;Q01730;A2AUR7;B1AYQ0;E0CXG5;E0CXH9 | 23617 | 45078 | 1.91 | 0.13024 | 221211 | 66614 | -3.32 | 0.16590 |
| Q9D051 | 107938 | 116171 | 1.08 | 0.56546 | 87288 | 65886 | -1.32 | 0.18544 |
| Q9D0E1;B8JK33;B8JK32;F6W322;F7C9U3 | 35592 | 8249 | -4.31 | 0.33331 | 61863 | 16778 | -3.69 | 0.10238 |
| Q9D0F9;A2CEK3 | 81778 | 97495 | 1.19 | 0.04578 | 53227 | 35883 | -1.48 | 0.30738 |
| Q9D0K2;Q3UJQ9 | 61352 | 73257 | 1.19 | 0.30643 | 52612 | 32435 | -1.62 | 0.17427 |
| Q9D0M5;D6RIN4;Q80ZS7 | 30386 | 25520 | -1.19 | 0.42096 | 78609 | 33365 | -2.36 | 0.08529 |
| Q9D0S9 | 66161 | 4435 | -14.92 | 0.08926 | 57844 | 44008 | -1.31 | 0.67972 |
| Q9D172 | 17954 | 15310 | -1.17 | 0.25172 | 77172 | 24240 | -3.18 | 0.07331 |
| Q9D1A2 | 69804 | 68855 | -1.01 | 0.92314 | 62006 | 41606 | -1.49 | 0.34050 |
| Q9D1G1 | 34158 | 40811 | 1.19 | 0.06934 | 49107 | 28542 | -1.72 | 0.32420 |
| Q9D1H6;A2AJX3;REV__Q6NS46 | 5875 | 39656 | 6.75 | 0.32806 | 75121 | 21201 | -3.54 | 0.08639 |
| Q9D1K2;F7B2B4 | 17085 | 15480 | -1.10 | 0.71584 | 59739 | 55733 | -1.07 | 0.87969 |
| Q9D1Q6 | 39915 | 11997 | -3.33 | 0.30164 | 56168 | 36192 | -1.55 | 0.54930 |
| Q9D1X0 | 20139 | 18558 | -1.09 | 0.52163 | 77950 | 24097 | -3.23 | 0.06588 |
| Q9D2G2 | 15814 | 18196 | 1.15 | 0.69056 | 58955 | 17149 | -3.44 | 0.14637 |
| Q9D394;D6RET4;D3Z4D2;Q8BR30;Q8R4C2;Q8BIJ7 | 10016 | 8456 | -1.18 | 0.26085 | 60278 | 24847 | -2.43 | 0.25250 |
| Q9D3D9;D3Z7S4 | 17820 | 12221 | -1.46 | 0.27190 | 61553 | 19710 | -3.12 | 0.12566 |
| Q9D6F9 | 30230 | 20052 | -1.51 | 0.03378 | 76477 | 38910 | -1.97 | 0.01360 |
| Q9D6R2 | 71584 | 75214 | 1.05 | 0.69020 | 107330 | 62031 | -1.73 | 0.08830 |
| Q9D819 | 57288 | 76192 | 1.33 | 0.05659 | 65217 | 20794 | -3.14 | 0.10881 |
| Q9D8B3 | 3797 | 32269 | 8.50 | 0.30544 | 74906 | 40227 | -1.86 | 0.27119 |
| Q9D8E6 | 4475 | 4881 | 1.09 | 0.76884 | 63910 | 18155 | -3.52 | 0.09284 |
| Q9D8N0 | 54207 | 48737 | -1.11 | 0.10113 | 72899 | 28730 | -2.54 | 0.05128 |
| Q9D8S4 | 15100 | 8789 | -1.72 | 0.05629 | 57042 | 41093 | -1.39 | 0.63068 |
| Q9D8U8 | 33982 | 4342 | -7.83 | 0.30425 | 76217 | 20472 | -3.72 | 0.07314 |
| Q9D8W5;B1AT36;Q3TRH2 | 62490 | 2337 | -26.74 | 0.08838 | 56065 | 40799 | -1.37 | 0.65477 |

| | | | | | | | | |
|---|---|---|---|---|---|---|---|---|
| Q9DAK9 | 17621 | 16046 | -1.10 | 0.13891 | 59325 | 37050 | -1.60 | 0.46905 |
| Q9DAR7;Q3TBW9;D6RFQ0 | 9165 | 10320 | 1.13 | 0.68083 | 75700 | 54421 | -1.39 | 0.44523 |
| Q9DB20;F7D3P8;F6XVM5 | 99289 | 38908 | -2.55 | 0.17912 | 121318 | 82430 | -1.47 | 0.10569 |
| Q9DB77 | 96357 | 59652 | -1.62 | 0.35857 | 68352 | 93412 | 1.37 | 0.19887 |
| Q9DBB8 | 14565 | 15751 | 1.08 | 0.66681 | 75387 | 22423 | -3.36 | 0.08719 |
| Q9DBC7;D3Z068;D3Z0V6;P12849;D3YTM5;D3Z4L4;A2AI69 | 36126 | 9848 | -3.67 | 0.34725 | 23517 | 8087 | -2.91 | 0.04713 |
| Q9DBF1;G3UYR8;G3UY72;E9Q1H3;E9Q1G1 | 56750 | 59062 | 1.04 | 0.63136 | 65635 | 26518 | -2.48 | 0.17527 |
| Q9DBJ1;O70250 | 320486 | 337369 | 1.05 | 0.38310 | 253482 | 154864 | -1.64 | 0.24124 |
| Q9DBP5 | 28787 | 27890 | -1.03 | 0.89548 | 45992 | 16276 | -2.83 | 0.17766 |
| Q9DC61;A2AIW9 | 5678 | 5505 | -1.03 | 0.85787 | 53251 | 40522 | -1.31 | 0.67757 |
| Q9DCC5;P23198;D3Z1A9;D3Z313 | 8497 | 10829 | 1.27 | 0.12997 | 76654 | 22706 | -3.38 | 0.07465 |
| Q9DCD0 | 56349 | 50703 | -1.11 | 0.65380 | 51130 | 16715 | -3.06 | 0.03036 |
| Q9DCH4 | 5593 | 6596 | 1.18 | 0.59164 | 54891 | 20735 | -2.65 | 0.24576 |
| Q9DCL9;D3Z6P1;D6RCU8 | 5676 | 33891 | 5.97 | 0.30134 | 57206 | 5634 | -10.15 | 0.09059 |
| Q9DCN2;F2Z456;A0A0A0MQM3 | 66414 | 73294 | 1.10 | 0.89125 | 79662 | 51817 | -1.54 | 0.29318 |
| Q9DCT8 | 25264 | 16933 | -1.49 | 0.26558 | 84063 | 38862 | -2.16 | 0.04115 |
| Q9DCW4 | 24753 | 25746 | 1.04 | 0.67104 | 78084 | 16963 | -4.60 | 0.01894 |
| Q9DCZ1;F6VY18 | 6474 | 41387 | 6.39 | 0.30641 | 57301 | 41346 | -1.39 | 0.59298 |
| Q9EQ20 | 33172 | 47388 | 1.43 | 0.02017 | 80118 | 30625 | -2.62 | 0.06007 |
| Q9EQF6;Q3SYJ1 | 267667 | 218274 | -1.23 | 0.23949 | 90679 | 81728 | -1.11 | 0.77161 |
| Q9EQH2 | 2957 | 6898 | 2.33 | 0.22487 | 97519 | 39385 | -2.48 | 0.02721 |
| Q9EQH3 | 27111 | 44425 | 1.64 | 0.10492 | 44226 | 31654 | -1.40 | 0.58599 |
| Q9ER00 | 9160 | 10099 | 1.10 | 0.56913 | 76935 | 22949 | -3.35 | 0.07235 |
| Q9ER97;Q3USR6;D3YW07 | 15182 | 12560 | -1.21 | 0.56851 | 75707 | 22083 | -3.43 | 0.08270 |
| Q9ERD7 | 236779 | 160883 | -1.47 | 0.08116 | 417621 | 287299 | -1.45 | 0.07341 |
| Q9ERE7;D3YVR4;F6SWV4 | 36004 | 4555 | -7.90 | 0.28602 | 74883 | 39844 | -1.88 | 0.26884 |
| Q9ES97 | 34312 | 29342 | -1.17 | 0.88345 | 66034 | 34509 | -1.91 | 0.23852 |
| Q9JHI5 | 32697 | 38870 | 1.19 | 0.38856 | 60557 | 32450 | -1.87 | 0.35069 |
| Q9JHU4;F6ZX84 | 65723 | 28617 | -2.30 | 0.15923 | 181186 | 174768 | -1.04 | 0.90556 |
| Q9JI75 | 9720 | 14520 | 1.49 | 0.22426 | 74925 | 35948 | -2.08 | 0.19492 |
| Q9JII6;B1AXW3 | 86134 | 99331 | 1.15 | 0.13065 | 52488 | 22294 | -2.35 | 0.14233 |
| Q9JJU8 | 12146 | 10226 | -1.19 | 0.05536 | 57974 | 7716 | -7.51 | 0.09329 |
| Q9JJV2;D3YWS3 | 70224 | 65098 | -1.08 | 0.45821 | 75413 | 37638 | -2.00 | 0.03651 |
| Q9JKB1;P58321 | 2971 | 4406 | 1.48 | 0.35729 | 97519 | 39765 | -2.45 | 0.02695 |
| Q9JKF1 | 19923 | 25288 | 1.27 | 0.24221 | 31824 | 30272 | -1.05 | 0.83832 |

| | | | | | | | | |
|---|---|---|---|---|---|---|---|---|
| Q9JKK7 | 26029 | 27291 | 1.05 | 0.85475 | 46302 | 22544 | -2.05 | 0.08966 |
| Q9JKR6;E0CYZ2;F6TRP3 | 22873 | 21164 | -1.08 | 0.63952 | 44688 | 31578 | -1.42 | 0.59061 |
| Q9JLJ2;Q3U367 | 37084 | 38251 | 1.03 | 0.59215 | 60546 | 28370 | -2.13 | 0.28790 |
| Q9JLM8;Q80VB6;H7BX36;Q9CXL6;Q6PGI2;O88809 | 4404 | 5091 | 1.16 | 0.75612 | 59114 | 21735 | -2.72 | 0.24289 |
| Q9JLV1 | 17928 | 16991 | -1.06 | 0.50148 | 76901 | 24622 | -3.12 | 0.07745 |
| Q9JLZ3;F6RT60;F6R307;E9Q6L3;E9QMT1 | 2228 | 66040 | 29.65 | 0.09595 | 52490 | 40205 | -1.31 | 0.67339 |
| Q9JM14;A2A9X5 | 13483 | 17292 | 1.28 | 0.22436 | 41928 | 56188 | 1.34 | 0.60549 |
| Q9JM76;H7BWZ3;D3Z2F7;D3Z2F8 | 17421 | 18116 | 1.04 | 0.70415 | 67193 | 30203 | -2.22 | 0.18146 |
| Q9JMA1;E9PYI8 | 19003 | 17772 | -1.07 | 0.57124 | 54894 | 25440 | -2.16 | 0.26049 |
| Q9JME5;E9Q1S6 | 2617 | 68309 | 26.10 | 0.09921 | 75917 | 20155 | -3.77 | 0.07528 |
| Q9JMG7;Q3UMU9 | 35797 | 31283 | -1.14 | 0.89182 | 43445 | 21128 | -2.06 | 0.37912 |
| Q9JMH6 | 20498 | 27595 | 1.35 | 0.16623 | 58938 | 25422 | -2.32 | 0.28249 |
| Q9QUI0;Q9CR99 | 22455 | 16926 | -1.33 | 0.34535 | 58776 | 30111 | -1.95 | 0.17752 |
| Q9QUM9;E0CXB1;E0CYT2 | 36269 | 32986 | -1.10 | 0.08058 | 78589 | 12514 | -6.28 | 0.01241 |
| Q9QUR6 | 13636 | 23807 | 1.75 | 0.03410 | 74991 | 35166 | -2.13 | 0.18917 |
| Q9QWR8 | 5864 | 8204 | 1.40 | 0.17304 | 75874 | 22125 | -3.43 | 0.08094 |
| Q9QX60;Q504N4 | 6657 | 14248 | 2.14 | 0.01038 | 75365 | 54912 | -1.37 | 0.46203 |
| Q9QXS1;E9Q3W4;F6R059;E9Q9J6;E9PW24 | 411266 | 362226 | -1.14 | 0.35068 | 309837 | 362071 | 1.17 | 0.41842 |
| Q9QXS6;F7CPL2 | 8495 | 7834 | -1.08 | 0.55917 | 59146 | 23447 | -2.52 | 0.25806 |
| Q9QXV0 | 294715 | 273581 | -1.08 | 0.61446 | 85507 | 93442 | 1.09 | 0.75645 |
| Q9QYB5 | 34865 | 36926 | 1.06 | 0.74068 | 72412 | 29453 | -2.46 | 0.06350 |
| Q9QYB8 | 18251 | 15543 | -1.17 | 0.09099 | 78613 | 25729 | -3.06 | 0.06334 |
| Q9QYC0;F8WHZ9;F8WGR0;E9Q1K3;F6RDR0;D3Z0T1;F6V4G5 | 74195 | 66188 | -1.12 | 0.05466 | 87108 | 64381 | -1.35 | 0.40675 |
| Q9QYG0 | 42543 | 48041 | 1.13 | 0.18128 | 62061 | 24252 | -2.56 | 0.05226 |
| Q9QYJ0 | 65434 | 37811 | -1.73 | 0.54211 | 54523 | 27033 | -2.02 | 0.34521 |
| Q9QYR9 | 23408 | 25393 | 1.08 | 0.62693 | 75941 | 24176 | -3.14 | 0.08642 |
| Q9QZ06;Q8C5G6;A9JEI5;F7AT44 | 10558 | 13313 | 1.26 | 0.06055 | 62370 | 10406 | -5.99 | 0.06516 |
| Q9QZE7 | 7090 | 7170 | 1.01 | 0.91582 | 97519 | 40465 | -2.41 | 0.02647 |
| Q9QZQ8 | 95370 | 59311 | -1.61 | 0.38241 | 56855 | 54141 | -1.05 | 0.87072 |
| Q9R0P5 | 168358 | 217308 | 1.29 | 0.01583 | 208139 | 208873 | 1.00 | 0.99128 |
| Q9R0P9 | 733414 | 629315 | -1.17 | 0.17759 | 342511 | 326657 | -1.05 | 0.75811 |
| Q9R0Q6;D3YVI5 | 36750 | 9846 | -3.73 | 0.33423 | 58352 | 22205 | -2.63 | 0.26080 |
| Q9R0Q7;D3Z7C6 | 35498 | 7137 | -4.97 | 0.31616 | 54417 | 55441 | 1.02 | 0.97077 |
| Q9R0X4;Q32MW3 | 39794 | 12602 | -3.16 | 0.31420 | 60158 | 9741 | -6.18 | 0.08325 |
| Q9R0Y5;Z4YN97 | 45761 | 47072 | 1.03 | 0.89242 | 73858 | 33108 | -2.23 | 0.00734 |

| | | | | | | | | |
|---|---|---|---|---|---|---|---|---|
| Q9R111;D3YU09 | 14628 | 16298 | 1.11 | 0.69721 | 58089 | 22120 | -2.63 | 0.26469 |
| Q9R1P0;E9PW69;E9Q0X0 | 28721 | 26006 | -1.10 | 0.20197 | 77489 | 11752 | -6.59 | 0.01523 |
| Q9R1P1;D3YUM8 | 21125 | 23857 | 1.13 | 0.33781 | 45170 | 24896 | -1.81 | 0.44085 |
| Q9R1P3 | 26540 | 25506 | -1.04 | 0.28875 | 55031 | 28453 | -1.93 | 0.30809 |
| Q9R1P4 | 30915 | 34550 | 1.12 | 0.45543 | 78892 | 45936 | -1.72 | 0.22352 |
| Q9R1Q8 | 16796 | 17313 | 1.03 | 0.82756 | 61277 | 10863 | -5.64 | 0.07650 |
| Q9R1T2 | 36873 | 34117 | -1.08 | 0.94089 | 56247 | 20908 | -2.69 | 0.28624 |
| Q9R1T4;A2A3W1 | 15584 | 12218 | -1.28 | 0.01920 | 79682 | 18173 | -4.38 | 0.01364 |
| Q9WTI7 | 127102 | 31751 | -4.00 | 0.01009 | 35046 | 27672 | -1.27 | 0.70380 |
| Q9WTL7 | 12820 | 11656 | -1.10 | 0.31526 | 20014 | 74819 | 3.74 | 0.00035 |
| Q9WTP6;F7BP55 | 33745 | 10362 | -3.26 | 0.37559 | 97519 | 19735 | -4.94 | 0.00348 |
| Q9WTP7;F6RP11 | 15650 | 20720 | 1.32 | 0.04297 | 56591 | 24282 | -2.33 | 0.31643 |
| Q9WTQ5 | 144292 | 110000 | -1.31 | 0.28329 | 58865 | 62393 | 1.06 | 0.89582 |
| Q9WTX5;E9PUV4 | 23660 | 25289 | 1.07 | 0.61291 | 25875 | 10516 | -2.46 | 0.06407 |
| Q9WU78 | 18162 | 21736 | 1.20 | 0.49901 | 25462 | 12154 | -2.09 | 0.10334 |
| Q9WUK2 | 13215 | 11652 | -1.13 | 0.49080 | 33154 | 7796 | -4.25 | 0.08292 |
| Q9WUL7 | 11172 | 9086 | -1.23 | 0.30428 | 60054 | 7117 | -8.44 | 0.07122 |
| Q9WUM4;E9PX03;E9PZJ0;E9PVJ1 | 37930 | 39843 | 1.05 | 0.62318 | 82313 | 64092 | -1.28 | 0.27144 |
| Q9WUM5 | 14007 | 13102 | -1.07 | 0.43061 | 58649 | 13090 | -4.48 | 0.11724 |
| Q9WUR9;A2ARF6;F6TEU8;A2ARF5 | 33641 | 3609 | -9.32 | 0.30022 | 75053 | 39765 | -1.89 | 0.26457 |
| Q9WV02;A2AFI4;A2AFI3;Q91VM5;S4R1F6 | 10357 | 12827 | 1.24 | 0.71980 | 75838 | 21166 | -3.58 | 0.07868 |
| Q9WV32;F6VVE6;D3Z6S0;F6THG2 | 33584 | 32544 | -1.03 | 0.97828 | 75019 | 20448 | -3.67 | 0.08520 |
| Q9WV54;D3Z015;D3Z505 | 62570 | 41003 | -1.53 | 0.63453 | 44778 | 11202 | -4.00 | 0.14511 |
| Q9WV55 | 36376 | 8213 | -4.43 | 0.31591 | 43217 | 25190 | -1.72 | 0.50409 |
| Q9WV92;A2A842 | 50795 | 50539 | -1.01 | 0.98843 | 33284 | 23785 | -1.40 | 0.47163 |
| Q9WVA4 | 378358 | 368280 | -1.03 | 0.79953 | 149965 | 154717 | 1.03 | 0.91531 |
| Q9WVE8 | 8225 | 6190 | -1.33 | 0.06371 | 75219 | 21741 | -3.46 | 0.08689 |
| Q9WVJ2;E9Q5I9;E9Q0U1;F6ZQQ3;F6PXS6;E9PY93 | 35713 | 6508 | -5.49 | 0.30298 | 35589 | 21858 | -1.63 | 0.60173 |
| Q9WVL0 | 4676 | 5426 | 1.16 | 0.31981 | 56244 | 40063 | -1.40 | 0.63233 |
| Q9Z0E0 | 33598 | 57737 | 1.72 | 0.55040 | 56980 | 20548 | -2.77 | 0.26925 |
| Q9Z0E6;E9Q2N7;Q01514;Q8CFB4 | 16856 | 30997 | 1.84 | 0.06591 | 67196 | 40564 | -1.66 | 0.35033 |
| Q9Z0F7 | 265862 | 195405 | -1.36 | 0.20722 | 71403 | 83190 | 1.17 | 0.76081 |
| Q9Z0P4;G3X9P1;Q8BR92 | 18100 | 11710 | -1.55 | 0.14329 | 77565 | 29680 | -2.61 | 0.08613 |
| Q9Z0S1;D3Z0E6;D3Z5X0 | 26563 | 27585 | 1.04 | 0.77110 | 77593 | 28014 | -2.77 | 0.07961 |
| Q9Z1A1;B8JJG8;B8JJG6;F6QJV5;B8JJG7;B8JJG9;B8JJG3 | 9448 | 10833 | 1.15 | 0.59713 | 79651 | 40470 | -1.97 | 0.20825 |

| | | | | | | | | |
|---|---|---|---|---|---|---|---|---|
| Q9Z1G3 | 12739 | 10531 | -1.21 | 0.57640 | 35002 | 21695 | -1.61 | 0.61370 |
| Q9Z1J3;A6QRH3;F7CZD1;F6TXD3 | 33483 | 28074 | -1.19 | 0.88058 | 56139 | 40103 | -1.40 | 0.63568 |
| Q9Z1N5;Q8VDW0;G3UXI6;D6RHT5 | 55794 | 51409 | -1.09 | 0.55747 | 68536 | 31140 | -2.20 | 0.11718 |
| Q9Z1Q5 | 19118 | 24228 | 1.27 | 0.44207 | 79632 | 15122 | -5.27 | 0.01169 |
| Q9Z1Q9;G3UY93;G3UZ22;G3UYW2 | 35586 | 5693 | -6.25 | 0.29266 | 74939 | 39811 | -1.88 | 0.26740 |
| Q9Z1S5 | 36785 | 5237 | -7.02 | 0.28061 | 57941 | 7563 | -7.66 | 0.09319 |
| Q9Z1Z0 | 64549 | 3586 | -18.00 | 0.09929 | 76464 | 21160 | -3.61 | 0.07253 |
| Q9Z1Z2 | 13095 | 17336 | 1.32 | 0.07773 | 77205 | 22633 | -3.41 | 0.06919 |
| Q9Z204 | 34856 | 41514 | 1.19 | 0.87538 | 61864 | 13702 | -4.52 | 0.08468 |
| Q9Z2I9 | 28068 | 34124 | 1.22 | 0.19427 | 66946 | 23218 | -2.88 | 0.08165 |
| Q9Z2M7 | 33936 | 3793 | -8.95 | 0.29668 | 74670 | 20288 | -3.68 | 0.08850 |
| Q9Z2Q6 | 19795 | 18098 | -1.09 | 0.59796 | 74151 | 39823 | -1.86 | 0.13664 |
| Q9Z2U0;Q9CWH6 | 53017 | 55925 | 1.05 | 0.53249 | 44234 | 35643 | -1.24 | 0.71748 |
| Q9Z2U1;D3YX79 | 35306 | 37777 | 1.07 | 0.27863 | 65663 | 12568 | -5.22 | 0.04870 |
| Q9Z2X1;J3QM80;J3QMT0 | 8103 | 7313 | -1.11 | 0.56123 | 75605 | 42026 | -1.80 | 0.26937 |
| REV__E9Q0M7;REV__Q6DFW5 | 12097 | 19060 | 1.58 | 0.00029 | 76024 | 24878 | -3.06 | 0.08767 |
| REV__Q9D451 | 201220 | 202732 | 1.01 | 0.91259 | 120152 | 82661 | -1.45 | 0.05192 |
| REV__Q9EP53 | 38857 | 44130 | 1.14 | 0.89959 | 73139 | 29707 | -2.46 | 0.06390 |
| S4R1F9;S4R249;S4R285;Q8C8R3;S4R2D0;S4R2H6;S4R2F3;S4R1J9;S4R1V6;S4R2T7; S4R2B2;S4R2D6;S4R1I1;S4R1M4;S4R1B7;S4R1U2;S4R241;Q8JZQ2 | 238455 | 192760 | -1.24 | 0.05507 | 161377 | 225113 | 1.39 | 0.24711 |
| S4R1S2;G5E8K2;G5E8K3;S4R2S8;G5E8K5;S4R2K9;S4R236;S4R1X7;S4R2C1;S4R187; S4R2F5;S4R2J6;S4R165;G3X971;A0A087WNU5;S4R278;S4R208;S4R229;S4R1U4; Q3UVY0;S4R162;A0A087WRP9 | 30154 | 30430 | 1.01 | 0.96682 | 45915 | 55643 | 1.21 | 0.49185 |

**Supplementary Table 2. Protein expression profile in the myenteric plexus (MP) of the small intestine (SI) and large intestine (LI) in pre-symptomatic (ps)A30P and wild type (WT) mice detected by mass spectroscopy.** Total protein was isolated from the MP of the SI and the LI in psA30P and WT mice. Proteins were separated by high-performance liquid chromotography and analyzed by mass spectroscopy. In total, 1,044 proteins were expressed in the MP from the SI and the LI in psA30P and WT mice. Data represent n = 4 independent experiments. * $p \leq 0.05$, ** $p \leq 0.01$, and *** $p \leq 0.001$ using Student's t test.

### Supplementary Table 3

| Official miRNA Symbol | Accession | Target Sequence |
|---|---|---|
| mmu-let-7a | MIMAT0000521 | UGAGGUAGUAGGUUGUAUAGUU |
| mmu-let-7b | MIMAT0000522 | UGAGGUAGUAGGUUGUGUGGUU |
| mmu-let-7c | MIMAT0000523 | UGAGGUAGUAGGUUGUAUGGUU |
| mmu-let-7d | MIMAT0000383 | AGAGGUAGUAGGUUGCAUAGUU |
| mmu-let-7e | MIMAT0000524 | UGAGGUAGGAGGUUGUAUAGUU |
| mmu-let-7f | MIMAT0000525 | UGAGGUAGUAGAUUGUAUAGUU |
| mmu-let-7g | MIMAT0000121 | UGAGGUAGUAGUUUGUACAGUU |
| mmu-let-7i | MIMAT0000122 | UGAGGUAGUAGUUUGUGCUGUU |
| mmu-miR-1 | MIMAT0000123 | UGGAAUGUAAAGAAGUAUGUAU |
| mmu-miR-7a | MIMAT0000677 | UGGAAGACUAGUGAUUUUGUUGU |
| mmu-miR-7b | MIMAT0000678 | UGGAAGACUUGUGAUUUUGUUGU |
| mmu-miR-9 | MIMAT0000142 | UCUUUGGUUAUCUAGCUGUAUGA |
| mmu-miR-10a | MIMAT0000648 | UACCCUGUAGAUCCGAAUUUGUG |
| mmu-miR-10b | MIMAT0000208 | UACCCUGUAGAACCGAAUUUGUG |
| mmu-miR-15a | MIMAT0000526 | UAGCAGCACAUAAUGGUUUGUG |
| mmu-miR-15b | MIMAT0000124 | UAGCAGCACAUCAUGGUUUACA |
| mmu-miR-16 | MIMAT0000527 | UAGCAGCACGUAAAUAUUGGCG |
| mmu-miR-17 [*1] | MIMAT0000649 | CAAAGUGCUUACAGUGCAGGUAG |
| mmu-miR-18a | MIMAT0000528 | UAAGGUGCAUCUAGUGCAGAUAG |
| mmu-miR-18b | MIMAT0004858 | UAAGGUGCAUCUAGUGCUGUUAG |
| mmu-miR-19a | MIMAT0000651 | UGUGCAAAUCUAUGCAAAACUGA |
| mmu-miR-19b | MIMAT0000513 | UGUGCAAAUCCAUGCAAAACUGA |
| mmu-miR-20a [*2] | MIMAT0000529 | UAAAGUGCUUAUAGUGCAGGUAG |
| mmu-miR-20b [*2] | MIMAT0003187 | CAAAGUGCUCAUAGUGCAGGUAG |
| mmu-miR-21 | MIMAT0000530 | UAGCUUAUCAGACUGAUGUUGA |
| mmu-miR-22 | MIMAT0000531 | AAGCUGCCAGUUGAAGAACUGU |
| mmu-miR-23a | MIMAT0000532 | AUCACAUUGCCAGGGAUUUCC |
| mmu-miR-23b | MIMAT0000125 | AUCACAUUGCCAGGGAUUACC |
| mmu-miR-24 | MIMAT0000219 | UGGCUCAGUUCAGCAGGAACAG |
| mmu-miR-25 | MIMAT0000652 | CAUUGCACUUGUCUCGGUCUGA |
| mmu-miR-26a | MIMAT0000533 | UUCAAGUAAUCCAGGAUAGGCU |
| mmu-miR-26b | MIMAT0000534 | UUCAAGUAAUUCAGGAUAGGU |
| mmu-miR-27a | MIMAT0000537 | UUCACAGUGGCUAAGUUCCGC |
| mmu-miR-27b | MIMAT0000126 | UUCACAGUGGCUAAGUUCUGC |
| mmu-miR-28 | MIMAT0000653 | AAGGAGCUCACAGUCUAUUGAG |
| mmu-miR-29a | MIMAT0000535 | UAGCACCAUCUGAAAUCGGUUA |
| mmu-miR-29b | MIMAT0000127 | UAGCACCAUUUGAAAUCAGUGUU |
| mmu-miR-29c | MIMAT0000536 | UAGCACCAUUUGAAAUCGGUUA |
| mmu-miR-30a | MIMAT0000128 | UGUAAACAUCCUCGACUGGAAG |
| mmu-miR-30b | MIMAT0000130 | UGUAAACAUCCUACACUCAGCU |
| mmu-miR-30c | MIMAT0000514 | UGUAAACAUCCUACACUCUCAGC |
| mmu-miR-30d | MIMAT0000515 | UGUAAACAUCCCCGACUGGAAG |
| mmu-miR-30e | MIMAT0000248 | UGUAAACAUCCUUGACUGGAAG |
| mmu-miR-31 | MIMAT0000538 | AGGCAAGAUGCUGGCAUAGCUG |
| mmu-miR-32 | MIMAT0000654 | UAUUGCACAUUACUAAGUUGCA |
| mmu-miR-33 | MIMAT0000667 | GUGCAUUGUAGUUGCAUUGCA |
| mmu-miR-34a | MIMAT0000542 | UGGCAGUGUCUUAGCUGGUUGU |
| mmu-miR-34b-3p | MIMAT0004581 | AAUCACUAACUCCACUGCCAUC |
| mmu-miR-34b-5p | MIMAT0000382 | AGGCAGUGUAAUUAGCUGAUUGU |
| mmu-miR-34c | MIMAT0000381 | AGGCAGUGUAGUUAGCUGAUUGC |
| mmu-miR-92a | MIMAT0000539 | UAUUGCACUUGUCCCGGCCUG |
| mmu-miR-92b | MIMAT0004899 | UAUUGCACUCGUCCCGGCCUCC |

| | | |
|---|---|---|
| mmu-miR-93 | MIMAT0000540 | CAAAGUGCUGUUCGUGCAGGUAG |
| mmu-miR-96 | MIMAT0000541 | UUUGGCACUAGCACAUUUUUGCU |
| mmu-miR-98 | MIMAT0000545 | UGAGGUAGUAAGUUGUAUUGUU |
| mmu-miR-99a | MIMAT0000131 | AACCCGUAGAUCCGAUCUUGUG |
| mmu-miR-99b | MIMAT0000132 | CACCCGUAGAACCGACCUUGCG |
| mmu-miR-100 | MIMAT0000655 | AACCCGUAGAUCCGAACUUGUG |
| mmu-miR-101a | MIMAT0000133 | UACAGUACUGUGAUAACUGAA |
| mmu-miR-101b | MIMAT0000616 | UACAGUACUGUGAUAGCUGAA |
| mmu-miR-103 | MIMAT0000546 | AGCAGCAUUGUACAGGGCUAUGA |
| mmu-miR-105 | MIMAT0004856 | CCAAGUGCUCAGAUGCUUGUGGU |
| mmu-miR-106a *1 | MIMAT0000385 | CAAAGUGCUAACAGUGCAGGUAG |
| mmu-miR-106b | MIMAT0000386 | UAAAGUGCUGACAGUGCAGAU |
| mmu-miR-107 | MIMAT0000647 | AGCAGCAUUGUACAGGGCUAUCA |
| mmu-miR-122 | MIMAT0000246 | UGGAGUGUGACAAUGGUGUUUG |
| mmu-miR-124 | MIMAT0000134 | UAAGGCACGCGGUGAAUGCC |
| mmu-miR-125a-3p | MIMAT0004528 | ACAGGUGAGGUUCUUGGGAGCC |
| mmu-miR-125a-5p | MIMAT0000135 | UCCCUGAGACCCUUUAACCUGUGA |
| mmu-miR-125b-3p | MIMAT0004669 | ACGGGUUAGGCUCUUGGGAGCU |
| mmu-miR-125b-5p | MIMAT0000136 | UCCCUGAGACCCUAACUUGUGA |
| mmu-miR-126-3p | MIMAT0000138 | UCGUACCGUGAGUAAUAAUGCG |
| mmu-miR-126-5p | MIMAT0000137 | CAUUAUUACUUUUGGUACGCG |
| mmu-miR-127 | MIMAT0000139 | UCGGAUCCGUCUGAGCUUGGCU |
| mmu-miR-128 | MIMAT0000140 | UCACAGUGAACCGGUCUCUUU |
| mmu-miR-129-3p | MIMAT0000544 | AAGCCCUUACCCCAAAAAGCAU |
| mmu-miR-129-5p | MIMAT0000209 | CUUUUUGCGGUCUGGGCUUGC |
| mmu-miR-130a | MIMAT0000141 | CAGUGCAAUGUUAAAAGGGCAU |
| mmu-miR-130b | MIMAT0000387 | CAGUGCAAUGAUGAAAGGGCAU |
| mmu-miR-132 | MIMAT0000144 | UAACAGUCUACAGCCAUGGUCG |
| mmu-miR-133a | MIMAT0000145 | UUUGGUCCCCUUCAACCAGCUG |
| mmu-miR-133b | MIMAT0000769 | UUUGGUCCCCUUCAACCAGCUA |
| mmu-miR-134 | MIMAT0000146 | UGUGACUGGUUGACCAGAGGGG |
| mmu-miR-135a | MIMAT0000147 | UAUGGCUUUUUAUUCCUAUGUGA |
| mmu-miR-135b | MIMAT0000612 | UAUGGCUUUUCAUUCCUAUGUGA |
| mmu-miR-136 | MIMAT0000148 | ACUCCAUUUGUUUUGAUGAUGG |
| mmu-miR-137 | MIMAT0000149 | UUAUUGCUUAAGAAUACGCGUAG |
| mmu-miR-138 | MIMAT0000150 | AGCUGGUGUUGUGAAUCAGGCCG |
| mmu-miR-139-3p | MIMAT0004662 | UGGAGACGCGGCCCUGUUGGAG |
| mmu-miR-139-5p | MIMAT0000656 | UCUACAGUGCACGUGUCUCCAG |
| mmu-miR-140 | MIMAT0000151 | CAGUGGUUUUACCCUAUGGUAG |
| mmu-miR-141 | MIMAT0000153 | UAACACUGUCUGGUAAAGAUGG |
| mmu-miR-142-3p | MIMAT0000155 | UGUAGUGUUUCCUACUUUAUGGA |
| mmu-miR-142-5p | MIMAT0000154 | CAUAAAGUAGAAAGCACUACU |
| mmu-miR-143 | MIMAT0000247 | UGAGAUGAAGCACUGUAGCUC |
| mmu-miR-144 | MIMAT0000156 | UACAGUAUAGAUGAUGUACU |
| mmu-miR-145 | MIMAT0000157 | GUCCAGUUUUCCCAGGAAUCCCU |
| mmu-miR-146a | MIMAT0000158 | UGAGAACUGAAUUCCAUGGGUU |
| mmu-miR-146b | MIMAT0003475 | UGAGAACUGAAUUCCAUAGGCU |
| mmu-miR-147 | MIMAT0004857 | GUGUGCGGAAAUGCUUCUGCUA |
| mmu-miR-148a | MIMAT0000516 | UCAGUGCACUACAGAACUUUGU |
| mmu-miR-148b | MIMAT0000580 | UCAGUGCAUCACAGAACUUUGU |
| mmu-miR-149 | MIMAT0000159 | UCUGGCUCCGUGUCUUCACUCCC |
| mmu-miR-150 | MIMAT0000160 | UCUCCCAACCCUUGUACCAGUG |
| mmu-miR-151-3p | MIMAT0000161 | CUAGACUGAGGCUCCUUGAGG |
| mmu-miR-151-5p | MIMAT0004536 | UCGAGGAGCUCACAGUCUAGU |
| mmu-miR-152 | MIMAT0000162 | UCAGUGCAUGACAGAACUUGG |

| | | |
|---|---|---|
| mmu-miR-153 | MIMAT0000163 | UUGCAUAGUCACAAAAGUGAUC |
| mmu-miR-154 | MIMAT0000164 | UAGGUUAUCCGUGUUGCCUUCG |
| mmu-miR-155 | MIMAT0000165 | UUAAUGCUAAUUGUGAUAGGGGU |
| mmu-miR-181a | MIMAT0000210 | AACAUUCAACGCUGUCGGUGAGU |
| mmu-miR-181b *3 | MIMAT0000673 | AACAUUCAUUGCUGUCGGUGGGU |
| mmu-miR-181c | MIMAT0000674 | AACAUUCAACCUGUCGGUGAGU |
| mmu-miR-181d *3 | MIMAT0004324 | AACAUUCAUUGUUGUCGGUGGGU |
| mmu-miR-182 | MIMAT0000211 | UUUGGCAAUGGUAGAACUCACACCG |
| mmu-miR-183 | MIMAT0000212 | UAUGGCACUGGUAGAAUUCACU |
| mmu-miR-184 | MIMAT0000213 | UGGACGGAGAACUGAUAAGGGU |
| mmu-miR-185 | MIMAT0000214 | UGGAGAGAAAGGCAGUUCCUGA |
| mmu-miR-186 | MIMAT0000215 | CAAAGAAUUCUCCUUUUGGGCU |
| mmu-miR-187 | MIMAT0000216 | UCGUGUCUUGUGUUGCAGCCGG |
| mmu-miR-188-3p | MIMAT0004541 | CUCCCACAUGCAGGGUUUGCA |
| mmu-miR-188-5p | MIMAT0000217 | CAUCCCUUGCAUGGUGGAGGG |
| mmu-miR-190 | MIMAT0000220 | UGAUAUGUUUGAUAUAUUAGGU |
| mmu-miR-190b | MIMAT0004852 | UGAUAUGUUUGAUAUUGGGUU |
| mmu-miR-191 | MIMAT0000221 | CAACGGAAUCCCAAAAGCAGCUG |
| mmu-miR-192 | MIMAT0000517 | CUGACCUAUGAAUUGACAGCC |
| mmu-miR-193 | MIMAT0000223 | AACUGGCCUACAAAGUCCCAGU |
| mmu-miR-193b | MIMAT0004859 | AACUGGCCCACAAAGUCCCGCU |
| mmu-miR-194 | MIMAT0000224 | UGUAACAGCAACUCCAUGUGGA |
| mmu-miR-195 | MIMAT0000225 | UAGCAGCACAGAAAUAUUGGC |
| mmu-miR-196a | MIMAT0000518 | UAGGUAGUUUCAUGUUGUUGGG |
| mmu-miR-196b | MIMAT0001081 | UAGGUAGUUUCCUGUUGUUGGG |
| mmu-miR-199a-3p | MIMAT0000230 | ACAGUAGUCUGCACAUUGGUUA |
| mmu-miR-199a-5p | MIMAT0000229 | CCCAGUGUUCAGACUACCUGUUC |
| mmu-miR-200a | MIMAT0000519 | UAACACUGUCUGGUAACGAUGU |
| mmu-miR-200b | MIMAT0000233 | UAAUACUGCCUGGUAAUGAUGA |
| mmu-miR-200c | MIMAT0000657 | UAAUACUGCCGGGUAAUGAUGGA |
| mmu-miR-201 | MIMAT0000234 | UACUCAGUAAGGCAUUGUUCUU |
| mmu-miR-202-3p | MIMAT0000235 | AGAGGUAUAGCGCAUGGGAAGA |
| mmu-miR-202-5p | MIMAT0004546 | UUCCUAUGCAUAUACUUCUUU |
| mmu-miR-203 | MIMAT0000236 | GUGAAAUGUUUAGGACCACUAG |
| mmu-miR-204 | MIMAT0000237 | UUCCCUUUGUCAUCCUAUGCCU |
| mmu-miR-205 | MIMAT0000238 | UCCUUCAUUCCACCGGAGUCUG |
| mmu-miR-206 | MIMAT0000239 | UGGAAUGUAAGGAAGUGUGUGG |
| mmu-miR-207 | MIMAT0000240 | GCUUCUCCUGGCUCUCCUCCCUC |
| mmu-miR-208a | MIMAT0000520 | AUAAGACGAGCAAAAAGCUUGU |
| mmu-miR-208b | MIMAT0004939 | AUAAGACGAACAAAAGGUUUGU |
| mmu-miR-210 | MIMAT0000658 | CUGUGCGUGUGACAGCGGCUGA |
| mmu-miR-211 | MIMAT0000668 | UUCCCUUUGUCAUCCUUUGCCU |
| mmu-miR-212 | MIMAT0000659 | UAACAGUCUCCAGUCACGGCCA |
| mmu-miR-214 | MIMAT0000661 | ACAGCAGGCACAGACAGGCAGU |
| mmu-miR-216a | MIMAT0000662 | UAAUCUCAGCUGGCAACUGUGA |
| mmu-miR-216b | MIMAT0003729 | AAAUCUCUGCAGGCAAAUGUGA |
| mmu-miR-217 | MIMAT0000679 | UACUGCAUCAGGAACUGACUGGA |
| mmu-miR-218 | MIMAT0000663 | UUGUGCUUGAUCUAACCAUGU |
| mmu-miR-219 | MIMAT0000664 | UGAUUGUCCAAACGCAAUUCU |
| mmu-miR-220 | MIMAT0004863 | CCACCACAGUGUCAGACACUU |
| mmu-miR-221 | MIMAT0000669 | AGCUACAUUGUCUGCUGGGUUUC |
| mmu-miR-222 | MIMAT0000670 | AGCUACAUCUGGCUACUGGGU |
| mmu-miR-223 | MIMAT0000665 | UGUCAGUUUGUCAAAUACCCCA |
| mmu-miR-224 | MIMAT0000671 | UAAGUCACUAGUGGUUCCGUU |
| mmu-miR-290-3p | MIMAT0004572 | AAAGUGCCGCCUAGUUUUAAGCCC |

| | | |
|---|---|---|
| mmu-miR-290-5p | MIMAT0000366 | ACUCAAACUAUGGGGGCACUUU |
| mmu-miR-291a-3p | MIMAT0000368 | AAAGUGCUUCCACUUUGUGUGC |
| mmu-miR-291a-5p | MIMAT0000367 | CAUCAAAGUGGAGGCCCUCUCU |
| mmu-miR-291b-3p | MIMAT0003190 | AAAGUGCAUCCAUUUUGUUUGU |
| mmu-miR-291b-5p | MIMAT0003189 | GAUCAAAGUGGAGGCCCUCUCC |
| mmu-miR-292-3p | MIMAT0000370 | AAAGUGCCGCCAGGUUUUGAGUGU |
| mmu-miR-292-5p | MIMAT0000369 | ACUCAAACUGGGGGCUCUUUUG |
| mmu-miR-293 | MIMAT0000371 | AGUGCCGCAGAGUUUGUAGUGU |
| mmu-miR-294 | MIMAT0000372 | AAAGUGCUUCCCUUUUGUGUGU |
| mmu-miR-295 | MIMAT0000373 | AAAGUGCUACUACUUUUGAGUCU |
| mmu-miR-296-3p | MIMAT0004576 | GAGGGUUGGGUGGAGGCUCUCC |
| mmu-miR-296-5p | MIMAT0000374 | AGGGCCCCCCCUCAAUCCUGU |
| mmu-miR-297a *4 | MIMAT0000375 | AUGUAUGUGUGCAUGUGCAUGU |
| mmu-miR-297b-3p | MIMAT0004827 | UAUACAUACACACAUACCCAUA |
| mmu-miR-297b-5p | MIMAT0003480 | AUGUAUGUGUGCAUGAACAUGU |
| mmu-miR-297c | MIMAT0004865 | AUGUAUGUGUGCAUGUACAUGU |
| mmu-miR-298 | MIMAT0000376 | GGCAGAGGAGGGCUGUUCUUCCC |
| mmu-miR-299 | MIMAT0004577 | UAUGUGGGACGGUAAACCGCUU |
| mmu-miR-300 | MIMAT0000378 | UAUGCAAGGGCAAGCUCUCUUC |
| mmu-miR-301a | MIMAT0000379 | CAGUGCAAUAGUAUUGUCAAAGC |
| mmu-miR-301b | MIMAT0004186 | CAGUGCAAUGGUAUUGUCAAAGC |
| mmu-miR-302a | MIMAT0000380 | UAAGUGCUUCCAUGUUUUGGUGA |
| mmu-miR-302b | MIMAT0003374 | UAAGUGCUUCCAUGUUUUAGUAG |
| mmu-miR-302c | MIMAT0003376 | AAGUGCUUCCAUGUUUCAGUGG |
| mmu-miR-302d | MIMAT0003377 | UAAGUGCUUCCAUGUUUGAGUGU |
| mmu-miR-320 | MIMAT0000666 | AAAAGCUGGGUUGAGAGGGCGA |
| mmu-miR-322 | MIMAT0000548 | CAGCAGCAAUUCAUGUUUUGGA |
| mmu-miR-323-3p | MIMAT0000551 | CACAUUACACGGUCGACCUCU |
| mmu-miR-323-5p | MIMAT0004638 | AGGUGGUCCGUGGCGCGUUCGC |
| mmu-miR-324-3p | MIMAT0000556 | CCACUGCCCCAGGUGCUGCU |
| mmu-miR-324-5p | MIMAT0000555 | CGCAUCCCCUAGGGCAUUGGUGU |
| mmu-miR-325 | MIMAT0004640 | UUUAUUGAGCACCUCCUAUCAA |
| mmu-miR-326 | MIMAT0000559 | CCUCUGGGCCCUUCCUCCAGU |
| mmu-miR-327 | MIMAT0004867 | ACUUGAGGGGCAUGAGGAU |
| mmu-miR-328 | MIMAT0000565 | CUGGCCCUCUCUGCCCUUCCGU |
| mmu-miR-329 | MIMAT0000567 | AACACACCCAGCUAACCUUUUU |
| mmu-miR-330 | MIMAT0004642 | UCUCUGGGCCUGUGUCUUAGGC |
| mmu-miR-331-3p | MIMAT0000571 | GCCCCUGGGCCUAUCCUAGAA |
| mmu-miR-331-5p | MIMAT0004643 | CUAGGUAUGGUCCCAGGGAUCC |
| mmu-miR-335-3p | MIMAT0004704 | UUUUUCAUUAUUGCUCCUGACC |
| mmu-miR-335-5p | MIMAT0000766 | UCAAGAGCAAUAACGAAAAAUGU |
| mmu-miR-337-3p | MIMAT0000578 | UUCAGCUCCUAUAUGAUGCCU |
| mmu-miR-337-5p | MIMAT0004644 | GAACGGCGUCAUGCAGGAGUU |
| mmu-miR-338-3p | MIMAT0000582 | UCCAGCAUCAGUGAUUUUGUUG |
| mmu-miR-338-5p | MIMAT0004647 | AACAAUAUCCUGGUGCUGAGUG |
| mmu-miR-339-3p | MIMAT0004649 | UGAGCGCCUCGGCGACAGAGCCG |
| mmu-miR-339-5p | MIMAT0000584 | UCCCUGUCCUCCAGGAGCUCACG |
| mmu-miR-340-3p | MIMAT0000586 | UCCGUCUCAGUUACUUUAUAGC |
| mmu-miR-340-5p | MIMAT0004651 | UUAUAAAGCAAUGAGACUGAUU |
| mmu-miR-341 | MIMAT0000588 | UCGGUCGAUCGGUCGGUCGGU |
| mmu-miR-342-3p | MIMAT0000590 | UCUCACACAGAAAUCGCACCCGU |
| mmu-miR-342-5p | MIMAT0004653 | AGGGGUGCUAUCUGUGAUUGAG |
| mmu-miR-343 | MIMAT0004868 | UCUCCCUUCAUGUGCCCAGA |
| mmu-miR-344 | MIMAT0000593 | UGAUCUAGCCAAAGCCUGACUGU |
| mmu-miR-345-3p | MIMAT0004656 | CCUGAACUAGGGGUCUGGAGAC |

| | | |
|---|---|---|
| mmu-miR-345-5p | MIMAT0000595 | GCUGACCCCUAGUCCAGUGCUU |
| mmu-miR-346 | MIMAT0000597 | UGUCUGCCCGAGUGCCUGCCUCU |
| mmu-miR-350 | MIMAT0000605 | UUCACAAAGCCCAUACACUUUC |
| mmu-miR-351 | MIMAT0000609 | UCCCUGAGGAGCCCUUUGAGCCUG |
| mmu-miR-361 | MIMAT0000704 | UUAUCAGAAUCUCCAGGGGUAC |
| mmu-miR-362-3p | MIMAT0004684 | AACACACCUGUUCAAGGAUUCA |
| mmu-miR-362-5p | MIMAT0000706 | AAUCCUUGGAACCUAGGUGUGAAU |
| mmu-miR-363 | MIMAT0000708 | AAUUGCACGGUAUCCAUCUGUA |
| mmu-miR-365 | MIMAT0000711 | UAAUGCCCCUAAAAAUCCUUAU |
| mmu-miR-367 | MIMAT0003181 | AAUUGCACUUUAGCAAUGGUGA |
| mmu-miR-369-3p | MIMAT0003186 | AAUAAUACAUGGUUGAUCUUU |
| mmu-miR-369-5p | MIMAT0003185 | AGAUCGACCGUGUUAUAUUCGC |
| mmu-miR-370 | MIMAT0001095 | GCCUGCUGGGGUGGAACCUGGU |
| mmu-miR-374 | MIMAT0003727 | AUAUAAUACAACCUGCUAAGUG |
| mmu-miR-375 | MIMAT0000739 | UUUGUUCGUUCGGCUCGCGUGA |
| mmu-miR-376a | MIMAT0000740 | AUCGUAGAGGAAAAUCCACGU |
| mmu-miR-376b | MIMAT0001092 | AUCAUAGAGGAACAUCCACUU |
| mmu-miR-376c | MIMAT0003183 | AACAUAGAGGAAAUUUCACGU |
| mmu-miR-377 | MIMAT0000741 | AUCACACAAAGGCAACUUUUGU |
| mmu-miR-378 | MIMAT0003151 | ACUGGACUUGGAGUCAGAAGG |
| mmu-miR-379 | MIMAT0000743 | UGGUAGACUAUGGAACGUAGG |
| mmu-miR-380-3p | MIMAT0000745 | UAUGUAGUAUGGUCCACAUCUU |
| mmu-miR-380-5p | MIMAT0000744 | AUGGUUGACCAUAGAACAUGCG |
| mmu-miR-381 | MIMAT0000746 | UAUACAAGGGCAAGCUCUCUGU |
| mmu-miR-382 | MIMAT0000747 | GAAGUUGUUCGUGGUGGAUUCG |
| mmu-miR-383 | MIMAT0000748 | AGAUCAGAAGGUGACUGUGGCU |
| mmu-miR-384-3p | MIMAT0001076 | AUUCCUAGAAAUUGUUCACAAU |
| mmu-miR-384-5p | MIMAT0004745 | UGUAAACAAUUCCUAGGCAAUGU |
| mmu-miR-409-3p | MIMAT0001090 | GAAUGUUGCUCGGUGAACCCCU |
| mmu-miR-409-5p | MIMAT0004746 | AGGUUACCCGAGCAACUUUGCAU |
| mmu-miR-410 | MIMAT0001091 | AAUAUAACACAGAUGGCCUGU |
| mmu-miR-411 | MIMAT0004747 | UAGUAGACCGUAUAGCGUACG |
| mmu-miR-412 | MIMAT0001094 | UUCACCUGGUCCACUAGCCG |
| mmu-miR-421 | MIMAT0004869 | AUCAACAGACAUUAAUUGGGCGC |
| mmu-miR-423-3p | MIMAT0003454 | AGCUCGGUCUGAGGCCCCUCAGU |
| mmu-miR-423-5p | MIMAT0004825 | UGAGGGGCAGAGAGCGAGACUUU |
| mmu-miR-425 | MIMAT0004750 | AAUGACACGAUCACUCCCGUUGA |
| mmu-miR-429 | MIMAT0001537 | UAAUACUGUCUGGUAAUGCCGU |
| mmu-miR-431 | MIMAT0001418 | UGUCUUGCAGGCCGUCAUGCA |
| mmu-miR-432 | MIMAT0012771 | UCUUGGAGUAGAUCAGUGGGCAG |
| mmu-miR-433 | MIMAT0001420 | AUCAUGAUGGGCUCCUCGGUGU |
| mmu-miR-434-3p | MIMAT0001422 | UUUGAACCAUCACUCGACUCCU |
| mmu-miR-434-5p | MIMAT0001421 | GCUCGACUCAUGGUUUGAACCA |
| mmu-miR-448 | MIMAT0001533 | UUGCAUAUGUAGGAUGUCCCAU |
| mmu-miR-449a | MIMAT0001542 | UGGCAGUGUAUUGUUAGCUGGU |
| mmu-miR-449b | MIMAT0005447 | AGGCAGUGUUGUUAGCUGGC |
| mmu-miR-449c | MIMAT0003460 | AGGCAGUGCAUUGCUAGCUGG |
| mmu-miR-450a-3p | MIMAT0004789 | AUUGGGGAUGCUUUGCAUUCAU |
| mmu-miR-450a-5p | MIMAT0001546 | UUUUGCGAUGUGUUCCUAAUAU |
| mmu-miR-450b-3p | MIMAT0003512 | AUUGGGAACAUUUUGCAUGCAU |
| mmu-miR-450b-5p | MIMAT0003511 | UUUUGCAGUAUGUUCCUGAAUA |
| mmu-miR-451 | MIMAT0001632 | AAACCGUUACCAUUACUGAGUU |
| mmu-miR-452 | MIMAT0001637 | UGUUUGCAGAGGAAACUGAGAC |
| mmu-miR-453 | MIMAT0004870 | AGGUUGCCUCAUAGUGAGCUUGCA |
| mmu-miR-455 | MIMAT0003742 | GCAGUCCACGGGCAUAUACAC |
| mmu-miR-463 | MIMAT0004758 | UGAUAGACACCAUAUAAGGUAG |

| | | |
|---|---|---|
| mmu-miR-464 | MIMAT0002105 | UACCAAGUUUAUUCUGUGAGAUA |
| mmu-miR-465a-3p | MIMAT0004217 | GAUCAGGGCCUUUCUAAGUAGA |
| mmu-miR-465a-5p | MIMAT0002106 | UAUUUAGAAUGGCACUGAUGUGA |
| mmu-miR-465b-5p | MIMAT0004871 | UAUUUAGAAUGGUGCUGAUCUG |
| mmu-miR-465c-5p | MIMAT0004873 | UAUUUAGAAUGGCGCUGAUCUG |
| mmu-miR-466a-3p *5 | MIMAT0002107 | UAUACAUACACGCACACAUAAGA |
| mmu-miR-466a-5p *6 | MIMAT0004759 | UAUGUGUGUGUACAUGUACAUA |
| mmu-miR-466b-3-3p *5 | MIMAT0005453 | AAUACAUACACGCACACAUAAGA |
| mmu-miR-466c-5p | MIMAT0004877 | GAUGUGUGUGUGCAUGUACAUA |
| mmu-miR-466d-3p | MIMAT0004931 | UAUACAUACACGCACACAUAG |
| mmu-miR-466d-5p | MIMAT0004930 | UGUGUGUGCGUACAUGUACAUG |
| mmu-miR-466e-5p *6 | MIMAT0004879 | GAUGUGUGUGUACAUGUACAUA |
| mmu-miR-466f-5p | MIMAT0004881 | UACGUGUGUGUGCAUGUGCAUG |
| mmu-miR-466f *4 | MIMAT0005844 | ACGUGUGUGUGCAUGUGCAUGU |
| mmu-miR-466g | MIMAT0004883 | AUACAGACACAUGCACACACA |
| mmu-miR-466h | MIMAT0004884 | UGUGUGCAUGUGCUUGUGUGUA |
| mmu-miR-466i | MIMAT0005834 | AUACACACACACAUACACACUA |
| mmu-miR-466j | MIMAT0005848 | UGUGUGCAUGUGCAUGUGUGUAA |
| mmu-miR-466k | MIMAT0005845 | UGUGUGUGUACAUGUACAUGUGA |
| mmu-miR-466l | MIMAT0005830 | UAUAAAUACAUGCACACAUAUU |
| mmu-miR-467a | MIMAT0003409 | UAAGUGCCUGCAUGUAUAUGCG |
| mmu-miR-467b | MIMAT0005448 | GUAAGUGCCUGCAUGUAUAUG |
| mmu-miR-467c | MIMAT0004885 | UAAGUGCGUGCAUGUAUAUGUG |
| mmu-miR-467d | MIMAT0004886 | UAAGUGCGCGCAUGUAUAUGCG |
| mmu-miR-467e | MIMAT0005293 | AUAAGUGUGAGCAUGUAUAUGU |
| mmu-miR-467f | MIMAT0005846 | AUAUACACACACACACCUACA |
| mmu-miR-467g | MIMAT0005854 | UAUACAUACACACACAUAUAU |
| mmu-miR-467h *7 | MIMAT0005855 | AUAAGUGUGUGCAUGUAUAUGU |
| mmu-miR-468 | MIMAT0002109 | UAUGACUGAUGUGCGUGUGUCUG |
| mmu-miR-469 | MIMAT0002110 | UGCCUCUUUCAUUGAUCUUGGUGUCC |
| mmu-miR-470 | MIMAT0002111 | UUCUUGGACUGGCACUGGUGAGU |
| mmu-miR-471 | MIMAT0002112 | UACGUAGUAUAGUGCUUUUCAC |
| mmu-miR-483 | MIMAT0004782 | AAGACGGGAGAAGAGAAGGGAG |
| mmu-miR-484 | MIMAT0003127 | UCAGGCUCAGUCCCCUCCCGAU |
| mmu-miR-485 | MIMAT0003128 | AGAGGCUGGCCGUGAUGAAUUC |
| mmu-miR-486 | MIMAT0003130 | UCCUGUACUGAGCUGCCCCGAG |
| mmu-miR-487b | MIMAT0003184 | AAUCGUACAGGGUCAUCCACUU |
| mmu-miR-488 | MIMAT0003450 | UUGAAAGGCUGUUUCUUGGUC |
| mmu-miR-489 | MIMAT0003112 | AAUGACACCACAUAUAUGGCAGC |
| mmu-miR-490 | MIMAT0003780 | CAACCUGGAGGACUCCAUGCUG |
| mmu-miR-491 | MIMAT0003486 | AGUGGGGAACCCUUCCAUGAGG |
| mmu-miR-493 | MIMAT0004888 | UGAAGGUCCUACUGUGUGCCAGG |
| mmu-miR-494 | MIMAT0003182 | UGAAACAUACACGGGAAACCUC |
| mmu-miR-495 | MIMAT0003456 | AAACAAACAUGGUGCACUUCUU |
| mmu-miR-496 | MIMAT0003738 | UGAGUAUUACAUGGCCAAUCUC |
| mmu-miR-497 | MIMAT0003453 | CAGCAGCACACUGUGGUUUGUA |
| mmu-miR-499 | MIMAT0003482 | UUAAGACUUGCAGUGAUGUUU |
| mmu-miR-500 | MIMAT0003507 | AAUGCACCUGGGCAAGGGUUCA |
| mmu-miR-501-3p | MIMAT0003509 | AAUGCACCCGGGCAAGGAUUUG |
| mmu-miR-501-5p | MIMAT0003508 | AAUCCUUUGUCCCUGGGUGAAA |
| mmu-miR-503 | MIMAT0003188 | UAGCAGCGGGAACAGUACUGCAG |
| mmu-miR-504 | MIMAT0004889 | AGACCCUGGUCUGCACUCUAUC |
| mmu-miR-505 | MIMAT0003513 | CGUCAACACUUGCUGGUUUUCU |
| mmu-miR-509-3p | MIMAT0004891 | UGAUUGACAUUUCUGUAAUGG |
| mmu-miR-509-5p | MIMAT0004890 | UACUCCAGAAUGUGGCAAUCAU |

| | | |
|---|---|---|
| mmu-miR-511 | MIMAT0004940 | AUGCCUUUUGCUCUGCACUCA |
| mmu-miR-532-3p | MIMAT0004781 | CCUCCCACACCCAAGGCUUGCA |
| mmu-miR-532-5p | MIMAT0002889 | CAUGCCUUGAGUGUAGGACCGU |
| mmu-miR-539 | MIMAT0003169 | GGAGAAAUUAUCCUUGGUGUGU |
| mmu-miR-540-3p | MIMAT0003167 | AGGUCAGAGGUCGAUCCUGG |
| mmu-miR-540-5p | MIMAT0004786 | CAAGGGUCACCCUCUGACUCUGU |
| mmu-miR-541 | MIMAT0003170 | AAGGGAUUCUGAUGUUGGUCACACU |
| mmu-miR-542-3p | MIMAT0003172 | UGUGACAGAUUGAUAACUGAAA |
| mmu-miR-542-5p | MIMAT0003171 | CUCGGGGAUCAUCAUGUCACGA |
| mmu-miR-543 | MIMAT0003168 | AAACAUUCGCGGUGCACUUCUU |
| mmu-miR-544 | MIMAT0004941 | AUUCUGCAUUUUUAGCAAGCUC |
| mmu-miR-546 | MIMAT0003166 | AUGGUGGCACGGAGUC |
| mmu-miR-547 | MIMAT0003173 | CUUGGUACAUCUUUGAGUGAG |
| mmu-miR-551b | MIMAT0003890 | GCGACCCAUACUUGGUUUCAG |
| mmu-miR-568 | MIMAT0004892 | AUGUAUAAAUGUAUACACAC |
| mmu-miR-574-3p | MIMAT0004894 | CACGCUCAUGCACACACCCACA |
| mmu-miR-574-5p | MIMAT0004893 | UGAGUGUGUGUGUGUGAGUGUGU |
| mmu-miR-582-3p | MIMAT0005292 | CCUGUUGAACAACUGAACCCAA |
| mmu-miR-582-5p | MIMAT0005291 | UACAGUUGUUCAACCAGUUACU |
| mmu-miR-590-3p | MIMAT0004896 | UAAUUUUAUGUAUAAGCUAGU |
| mmu-miR-590-5p | MIMAT0004895 | GAGCUUAUUCAUAAAAGUGCAG |
| mmu-miR-592 | MIMAT0003730 | AUUGUGUCAAUAUGCGAUGAUGU |
| mmu-miR-599 | MIMAT0012772 | UUGUGUCAGUUUAUCAAAC |
| mmu-miR-615-3p | MIMAT0003783 | UCCGAGCCUGGGUCUCCCUCUU |
| mmu-miR-615-5p | MIMAT0004837 | GGGGGUCCCCGGUGCUCGGAUC |
| mmu-miR-652 | MIMAT0003711 | AAUGGCGCCACUAGGGUUGUG |
| mmu-miR-653 | MIMAT0004943 | GUGUUGAAACAAUCUCUACUG |
| mmu-miR-654-3p | MIMAT0004898 | UAUGUCUGCUGACCAUCACCUU |
| mmu-miR-654-5p | MIMAT0004897 | UGGUAAGCUGCAGAACAUGUGU |
| mmu-miR-664 | MIMAT0012774 | UAUUCAUUUACUCCCCAGCCUA |
| mmu-miR-665 | MIMAT0003733 | ACCAGGAGGCUGAGGUCCCU |
| mmu-miR-666-3p | MIMAT0004823 | GGCUGCAGCGUGAUCGCCUGCU |
| mmu-miR-666-5p | MIMAT0003737 | AGCGGGCACAGCUGUGAGAGCC |
| mmu-miR-667 | MIMAT0003734 | UGACACCUGCCACCCAGCCCAAG |
| mmu-miR-668 | MIMAT0003732 | UGUCACUCGGCUCGGCCCACUACC |
| mmu-miR-669a | MIMAT0003477 | AGUUGUGUGUGCAUGUUCAUGU |
| mmu-miR-669b *4 | MIMAT0003476 | AGUUUUGUGUGCAUGUGCAUGU |
| mmu-miR-669d *7 | MIMAT0005833 | ACUUGUGUGUGCAUGUAUAUGU |
| mmu-miR-669e | MIMAT0005853 | UGUCUUGUGUGUGCAUGUUCAU |
| mmu-miR-669f | MIMAT0005839 | CAUAUACAUACACACACACGUAU |
| mmu-miR-669g | MIMAT0005832 | UGCAUUGUAUGUGUUGACAUGAU |
| mmu-miR-669h-5p | MIMAT0005841 | AUGCAUGGGUGUAUAGUUGAGUGC |
| mmu-miR-669i | MIMAT0005840 | UGCAUAUACACACAUGCAUAC |
| mmu-miR-669j | MIMAT0005838 | UGCAUAUACUCACAUGCAAACA |
| mmu-miR-669l *7 | MIMAT0009418 | AGUUGUGUGUGCAUGUAUAUGU |
| mmu-miR-669m | MIMAT0009419 | AUAUACAUCCACACAAACAUAU |
| mmu-miR-669n *8 | MIMAT0009427 | AUUUGUGUGUGGAUGUGUGU |
| mmu-miR-669o | MIMAT0009421 | UAGUUGUGUGUGCAUGUUUAUGU |
| mmu-miR-670 | MIMAT0003736 | AUCCCUGAGUGUAUGUGGUGAA |
| mmu-miR-671-3p | MIMAT0004821 | UCCGGUUCUCAGGGCUCCACC |
| mmu-miR-671-5p | MIMAT0003731 | AGGAAGCCCUGGAGGGGCUGGAG |
| mmu-miR-672 | MIMAT0003735 | UGAGGUUGGUGUACUGUGUGUGA |
| mmu-miR-673-3p | MIMAT0004824 | UCCGGGGCUGAGUUCUGUGCACC |
| mmu-miR-673-5p | MIMAT0003739 | CUCACAGCUCUGGUCCUUGGAG |
| mmu-miR-674 | MIMAT0003740 | GCACUGAGAUGGGAGUGGUGUA |

| | | |
|---|---|---|
| mmu-miR-675-3p | MIMAT0003726 | CUGUAUGCCCUAACCGCUCAGU |
| mmu-miR-675-5p | MIMAT0003725 | UGGUGCGGAAAGGGCCCACAGU |
| mmu-miR-676 | MIMAT0003782 | CCGUCCUGAGGUUGUUGAGCU |
| mmu-miR-677 | MIMAT0003451 | UUCAGUGAUGAUUAGCUUCUGA |
| mmu-miR-678 | MIMAT0003452 | GUCUCGGUGCAAGGACUGGAGG |
| mmu-miR-679 | MIMAT0003455 | GGACUGUGAGGUGACUCUUGGU |
| mmu-miR-680 | MIMAT0003457 | GGGCAUCUGCUGACAUGGGGG |
| mmu-miR-681 | MIMAT0003458 | CAGCCUCGCUGGCAGGCAGCU |
| mmu-miR-682 | MIMAT0003459 | CUGCAGUCACAGUGAAGUCUG |
| mmu-miR-683 | MIMAT0003461 | CCUGCUGUAAGCUGUGUCCUC |
| mmu-miR-684 | MIMAT0003462 | AGUUUUCCCUUCAAGUCAA |
| mmu-miR-686 | MIMAT0003464 | AUUGCUUCCCAGACGGUGAAGA |
| mmu-miR-687 | MIMAT0003466 | CUAUCCUGGAAUGCAGCAAUGA |
| mmu-miR-688 | MIMAT0003467 | UCGCAGGCGACUACUUAUUC |
| mmu-miR-689 | MIMAT0003468 | CGUCCCCGCUCGGCGGGGUCC |
| mmu-miR-690 | MIMAT0003469 | AAAGGCUAGGCUCACAACCAAA |
| mmu-miR-691 | MIMAT0003470 | AUUCCUGAAGAGAGGCAGAAAA |
| mmu-miR-692 | MIMAT0003471 | AUCUCUUUGAGCGCCUCACUC |
| mmu-miR-693-3p | MIMAT0004189 | GCAGCUUUCAGAUGUGGCUGUAA |
| mmu-miR-693-5p | MIMAT0003472 | CAGCCACAUCCGAAAGUUUUC |
| mmu-miR-694 | MIMAT0003474 | CUGAAAAUGUUGCCUGAAG |
| mmu-miR-695 | MIMAT0003481 | AGAUUGGGCAUAGGUGACUGAA |
| mmu-miR-696 | MIMAT0003483 | GCGUGUGCUUGCUGUGGG |
| mmu-miR-697 | MIMAT0003487 | AACAUCCUGGUCCUGUGGAGA |
| mmu-miR-698 | MIMAT0003488 | CAUUCUCGUUUCCUUCCCU |
| mmu-miR-700 | MIMAT0003490 | CACGCGGGAACCGAGUCCACC |
| mmu-miR-701 | MIMAT0003491 | UUAGCCGCUGAAAUAGAUGGA |
| mmu-miR-702 | MIMAT0003492 | UGCCCACCCUUUACCCCGCUC |
| mmu-miR-703 | MIMAT0003493 | AAAACCUUCAGAAGGAAAGAA |
| mmu-miR-704 | MIMAT0003494 | AGACAUGUGCUCUGCUCCUAG |
| mmu-miR-706 | MIMAT0003496 | AGAGAAACCCUGUCUCAAAAAA |
| mmu-miR-707 | MIMAT0003497 | CAGUCAUGCCGCUUGCCUACG |
| mmu-miR-708 | MIMAT0004828 | AAGGAGCUUACAAUCUAGCUGGG |
| mmu-miR-709 | MIMAT0003499 | GGAGGCAGAGGCAGGAGGA |
| mmu-miR-710 | MIMAT0003500 | CCAAGUCUUGGGGAGAGUUGAG |
| mmu-miR-711 | MIMAT0003501 | GGGACCCGGGGAGAGAUGUAAG |
| mmu-miR-712 | MIMAT0003502 | CUCCUUCACCCGGGCGGUACC |
| mmu-miR-713 | MIMAT0003504 | UGCACUGAAGGCACACAGC |
| mmu-miR-714 | MIMAT0003505 | CGACGAGGGCCGGUCGGUCGC |
| mmu-miR-715 | MIMAT0003506 | CUCCGUGCACACCCCCGCGUG |
| mmu-miR-717 | MIMAT0003510 | CUCAGACAGAGAUACCUUCUCU |
| mmu-miR-718 | MIMAT0003514 | CUUCCGCCCGGCCGGGUGUCG |
| mmu-miR-719 | MIMAT0003465 | AUCUCGGCUACAGAAAAAUGUU |
| mmu-miR-720 | MIMAT0003484 | AUCUCGCUGGGGCCUCCA |
| mmu-miR-741 | MIMAT0004236 | UGAGAGAUGCCAUUCUAUGUAGA |
| mmu-miR-742 | MIMAT0004237 | GAAAGCCACCAUGCUGGGUAAA |
| mmu-miR-743a | MIMAT0004238 | GAAAGACACCAAGCUGAGUAGA |
| mmu-miR-743b-3p | MIMAT0004840 | GAAAGACAUCAUGCUGAAUAGA |
| mmu-miR-743b-5p | MIMAT0004839 | UGUUCAGACUGGUGUCCAUCA |
| mmu-miR-744 | MIMAT0004187 | UGCGGGGCUAGGGCUAACAGCA |
| mmu-miR-758 | MIMAT0003889 | UUUGUGACCUGGUCCACUA |
| mmu-miR-759 | MIMAT0003897 | GCAGAGUGCAAACAAUUUUGAC |
| mmu-miR-760 | MIMAT0003898 | CGGCUCUGGGUCUGUGGGGA |
| mmu-miR-761 | MIMAT0003893 | GCAGCAGGGUGAAACUGACACA |
| mmu-miR-762 | MIMAT0003892 | GGGGCUGGGGCCGGGACAGAGC |
| mmu-miR-763 | MIMAT0003896 | CCAGCUGGGAAGAACCAGUGGC |

| | | |
|---|---|---|
| mmu-miR-764-3p | MIMAT0003895 | AGGAGGCCAUAGUGGCAACUGU |
| mmu-miR-764-5p | MIMAT0003894 | GGUGCUCACAUGUCCUCCU |
| mmu-miR-767 | MIMAT0012773 | UGCACCAUGGUUGUCUGAGCA |
| mmu-miR-770-3p | MIMAT0003891 | CGUGGGCCUGACGUGGAGCUGG |
| mmu-miR-770-5p | MIMAT0004822 | AGCACCACGUGUCUGGGCCACG |
| mmu-miR-802 | MIMAT0004188 | UCAGUAACAAAGAUUCAUCCUU |
| mmu-miR-804 | MIMAT0004210 | UGUGAGUUGUUCCUCACCUGGA |
| mmu-miR-871 | MIMAT0004841 | UAUUCAGAUUAGUGCCAGUCAUG |
| mmu-miR-872 | MIMAT0004934 | AAGGUUACUUGUUAGUUCAGG |
| mmu-miR-873 | MIMAT0004936 | GCAGGAACUUGUGAGUCUCCU |
| mmu-miR-874 | MIMAT0004853 | CUGCCCUGGCCCGAGGGACCGA |
| mmu-miR-875-3p | MIMAT0004938 | CCUGAAAAUACUGAGGCUAUG |
| mmu-miR-875-5p | MIMAT0004937 | UAUACCUCAGUUUUAUCAGGUG |
| mmu-miR-876-3p | MIMAT0004855 | UAGUGGUUUACAAAGUAAUUCA |
| mmu-miR-876-5p | MIMAT0004854 | UGGAUUUCUCUGUGAAUCACUA |
| mmu-miR-877 | MIMAT0004861 | GUAGAGGAGAUGGCGCAGGG |
| mmu-miR-878-3p | MIMAT0004933 | GCAUGACACCACACUGGGUAGA |
| mmu-miR-878-5p | MIMAT0004932 | UAUCUAGUUGGAUGUCAAGACA |
| mmu-miR-879 | MIMAT0004842 | AGAGGCUUAUAGCUCUAAGCC |
| mmu-miR-880 | MIMAT0004844 | UACUCCAUCCUCUCUGAGUAGA |
| mmu-miR-881 | MIMAT0004846 | AACUGUGUCUUUUCUGAAUAGA |
| mmu-miR-882 | MIMAT0004847 | AGGAGAGAGUUAGCGCAUUAGU |
| mmu-miR-883a-3p | MIMAT0004849 | UAACUGCAACAGCUCUCAGUAU |
| mmu-miR-883a-5p | MIMAT0004848 | UGCUGAGAGAAGUAGCAGUUAC |
| mmu-miR-883b-3p | MIMAT0004851 | UAACUGCAACAUCUCUCAGUAU |
| mmu-miR-883b-5p | MIMAT0004850 | UACUGAGAAUGGGUAGCAGUCA |
| mmu-miR-1186 | MIMAT0005836 | GAGUGCUGGAAUUAAAGGCAUG |
| mmu-miR-1186b | MIMAT0015644 | UGGGAUUAAAGGCAUGCACCAC |
| mmu-miR-1187 | MIMAT0005837 | UAUGUGUGUGUGUAUGUGUGUAA |
| mmu-miR-1188 | MIMAT0005843 | UGGUGUGAGGUUGGGCCAGGA |
| mmu-miR-1190 | MIMAT0005847 | UCAGCUGAGGUUCCCCUCUGUC |
| mmu-miR-1191 | MIMAT0005849 | CAGUCUUACUAUGUAGCCCUA |
| mmu-miR-1192 | MIMAT0005850 | AAACAAACAAACAGACCAAAUU |
| mmu-miR-1193 | MIMAT0005851 | UAGGUCACCCGUUUUACUAUC |
| mmu-miR-1194 | MIMAT0005852 | GAAUGAGUAACUGCUAGAUCCU |
| mmu-miR-1195 | MIMAT0005856 | UGAGUUCGAGGCCAGCCUGCUCA |
| mmu-miR-1196 | MIMAT0005857 | AAAUCUACCUGCCUCUGCCU |
| mmu-miR-1197 | MIMAT0005858 | UAGGACACAUGGUCUACUUCU |
| mmu-miR-1198 | MIMAT0005859 | UAUGUGUUCCUGGCUGGCUUGG |
| mmu-miR-1199 | MIMAT0005860 | UCUGAGUCCCGGUCGCGCGG |
| mmu-miR-1224 | MIMAT0005460 | GUGAGGACUGGGGAGGUGGAG |
| mmu-miR-1274a | MIMAT0009445 | UCAGGUCCCUGUUCAGGCGCCA |
| mmu-miR-1306 | MIMAT0009411 | ACGUUGGCUCUGGUGGUGAUG |
| mmu-miR-1839-3p | MIMAT0009457 | AGACCUACUUAUCUACCAACAGC |
| mmu-miR-1839-5p | MIMAT0009456 | AAGGUAGAUAGAACAGGUCUUG |
| mmu-miR-1892 | MIMAT0007871 | AUUUGGGGACGGGAGGGAGGAU |
| mmu-miR-1893 | MIMAT0007879 | GGCGCGGGCGCUGGACGCCUCG |
| mmu-miR-1894-3p | MIMAT0007878 | GCAAGGGAGAGGGUGAAGGGAG |
| mmu-miR-1894-5p | MIMAT0007877 | CUCUCCCCUACCACCUGCCUCU |
| mmu-miR-1895 | MIMAT0007867 | CCCCCGAGGAGGACGAGGAGGA |
| mmu-miR-1896 | MIMAT0007873 | CUCUCUGAUGGUGGGUGAGGAG |
| mmu-miR-1897-3p | MIMAT0007865 | UCAACUCGUUCUGUCCGGUGAG |
| mmu-miR-1898 | MIMAT0007875 | AGGUCAAGGUUCACAGGGGAUC |
| mmu-miR-1899 | MIMAT0007869 | AGCGAUGGCCGAAUCUGCUUCC |
| mmu-miR-1900 | MIMAT0007870 | GGCCGCCCUCUCUGGUCCUUCA |
| mmu-miR-1901 | MIMAT0007880 | CCGCUCGUACUCCCGGGGGUCC |

| | | |
|---|---|---|
| mmu-miR-1902 | MIMAT0007863 | AGAGGUGCAGUAGGCAUGACUU |
| mmu-miR-1903 | MIMAT0007868 | CCUUCUUCUUCUUCCUGAGACA |
| mmu-miR-1904 | MIMAT0007874 | GUUCUGCUCCUCUGGAGGGAGG |
| mmu-miR-1905 | MIMAT0007866 | CACCAGUCCCACCACGCGGUAG |
| mmu-miR-1906 | MIMAT0007872 | UGCAGCAGCCUGAGGCAGGGCU |
| mmu-miR-1907 | MIMAT0007876 | GAGCAGCAGAGGAUCUGGAGGU |
| mmu-miR-1927 | MIMAT0009390 | GACCUCUGGAUGUUAGGGACUGA |
| mmu-miR-1928 | MIMAT0009391 | AGCUACAUUGCCAGCUC |
| mmu-miR-1929 | MIMAT0009392 | UUCUAGGACUUUAUAGAGCAGAG |
| mmu-miR-1930 | MIMAT0009393 | ACCUCCAUAGUACCUGCAGCGU |
| mmu-miR-1931 | MIMAT0009394 | AUGCAAGGGCUGGUGCGAUGGC |
| mmu-miR-1932 | MIMAT0009395 | GUUGCGGACAGCGCUAGGUCGG |
| mmu-miR-1933-3p | MIMAT0009397 | CCAGGACCAUCAGUGUGACUAU |
| mmu-miR-1933-5p | MIMAT0009396 | AGUCAUGGUGUUCGGUCUUAGUUU |
| mmu-miR-1934 | MIMAT0009398 | UCUGGUCCCCUGCUUCGUCCUCU |
| mmu-miR-1935 | MIMAT0009399 | AGGCAGAGGCUGGCGGAUCUCU |
| mmu-miR-1936 | MIMAT0009400 | UAACUGACCUGCUGUGAACUGGC |
| mmu-miR-1937a [*9] | MIMAT0009401 | AAUCCCGGACGAGCCCCCA |
| mmu-miR-1937b [*9] | MIMAT0009414 | AUCCCGGACGAGCCCCCA |
| mmu-miR-1937c | MIMAT0009429 | AUCCCGGAAGAGCCCCCA |
| mmu-miR-1938 | MIMAT0009402 | CGGUGGGACUUGUAGUUCGGUC |
| mmu-miR-1939 | MIMAT0009403 | UCGAUUCCCUGCCAAUGCAC |
| mmu-miR-1940 | MIMAT0009404 | AUGGAGGACUGAGAAGGUGGAGCAGUU |
| mmu-miR-1941-3p | MIMAT0009406 | CAUCUUAGCAGUAUCUCCCAU |
| mmu-miR-1941-5p | MIMAT0009405 | AGGGAGAUGCUGGUACAGAGGCUU |
| mmu-miR-1942 | MIMAT0009407 | UCAGAUGUCUUCAUCUGGUUG |
| mmu-miR-1943 | MIMAT0009408 | AAGGGAGGAUCUGGGCACCUGGA |
| mmu-miR-1944 | MIMAT0009409 | CUCUGUGCUGAAUGUCAAGUUCUGAUU |
| mmu-miR-1945 | MIMAT0009410 | UCUUCGCGGGUACUGUCGGGAC |
| mmu-miR-1946a | MIMAT0009412 | AGCCGGGCAGUGGUGGCACACACUUUU |
| mmu-miR-1946b | MIMAT0009443 | GCCGGGCAGUGGUGGCACAUGCUUUU |
| mmu-miR-1947 | MIMAT0009413 | AGGACGAGCUAGCUGAGUGCUG |
| mmu-miR-1948 | MIMAT0009415 | UUUAGGCAGAGCACUCGUACAG |
| mmu-miR-1949 | MIMAT0009416 | CUAUACCAGGAUGUCAGCAUAGUU |
| mmu-miR-1950 | MIMAT0009417 | UCUGCAUCUAAGGAUAUGGUCA |
| mmu-miR-1951 | MIMAT0009422 | GUAGUGGAGACUGGUGUGGCUA |
| mmu-miR-1952 | MIMAT0009423 | UCUCCACCCUCCUUCUG |
| mmu-miR-1953 | MIMAT0009424 | UGGGAAAGUUCUCAGGCUUCUG |
| mmu-miR-1954 | MIMAT0009425 | ACUGCAGAGUGAGACCCUGUU |
| mmu-miR-1955 | MIMAT0009426 | AGUCCCAGGAUGCACUGCAGCUUUU |
| mmu-miR-1956 | MIMAT0009428 | AGUCCAGGGCUGAGUCAGCGGA |
| mmu-miR-1957 | MIMAT0009430 | CAGUGGUAGAGCAUAUGAC |
| mmu-miR-1958 | MIMAT0009431 | UAGGAAAGUGGAAGCAGUAAGU |
| mmu-miR-1959 | MIMAT0009432 | GGGGAUGUAGCUCAGUGGAG |
| mmu-miR-1960 | MIMAT0009433 | CCAGUGCUGUUAGAAGAGGGCU |
| mmu-miR-1961 | MIMAT0009434 | UGAGGUAGUAGUUAGAA |
| mmu-miR-1962 | MIMAT0009435 | AGAGGCUGGCACUGGGACACAU |
| mmu-miR-1963 | MIMAT0009436 | UGGGACGAGAUCAUGAGGCCUUC |
| mmu-miR-1964 | MIMAT0009437 | CCGACUUCUGGGCUCCGGCUUU |
| mmu-miR-1965 | MIMAT0009438 | AAGCCGGGCCGUAGUGGCGCA |
| mmu-miR-1966 | MIMAT0009439 | AAGGGAGCUGGCUCAGGAGAGAGUC |
| mmu-miR-1967 | MIMAT0009440 | UGAGGAUCCUGGGGAGAAGAUGC |
| mmu-miR-1968 | MIMAT0009441 | UGCAGCUGUUAAGGAUGGUGGACU |
| mmu-miR-1969 | MIMAT0009442 | AAGAUGGAGACUUUAACAUGGGU |
| mmu-miR-1970 | MIMAT0009444 | UGUGUCACUGGGGAUAGGCUUUG |

| | | |
|---|---|---|
| mmu-miR-1971 | MIMAT0009446 | GUAAAGGCUGGGCUGAGA |
| mmu-miR-1981 | MIMAT0009458 | GUAAAGGCUGGGCUUAGACGUGGC |
| mmu-miR-1982 | MIMAT0009460 | TCTCACCCTATGTTCTCCCACAG |
| mmu-miR-1983 | MIMAT0009455 | CUCACCUGGAGCAUGUUUUCU |
| mmu-miR-2132 | MIMAT0011208 | GGCGGGUGUUGACGCGAUG |
| mmu-miR-2133 | MIMAT0011209 | GUCCCGCGGGGCCCGAAGCGUU |
| mmu-miR-2134 | MIMAT0011210 | GUCUUGGGAAACGGGGUGC |
| mmu-miR-2135 | MIMAT0011211 | AGAGGUCUUGGGGCCGAAAC |
| mmu-miR-2136 | MIMAT0011212 | CUGGGUGUUGACUGAGAUGUG |
| mmu-miR-2137 | MIMAT0011213 | GCCGGCGGGAGCCCCAGGGAG |
| mmu-miR-2138 | MIMAT0011214 | AAGGGAACGGGCUUGGCGGAAU |
| mmu-miR-2139 | MIMAT0011215 | AGCUGCGCUGCUCCUGGUAACUGC |
| mmu-miR-2140 | MIMAT0011216 | AGGUGCAGAUCUUGGUGGU |
| mmu-miR-2141 | MIMAT0011217 | AGGAGGUGUCAGAAAAGUU |
| mmu-miR-2145 | MIMAT0011221 | AGCAGGGUCGGGCCUGGUU |
| mmu-miR-2146 | MIMAT0011222 | GUGGAGAAGGGUUCCAUGUG |
| mmu-miR-2182 | MIMAT0011286 | ACGCCACAUUUCCCACGCCGCG |
| mmu-miR-2183 | MIMAT0011287 | UUGAACCCCUGACCUCCU |
| mmu-miR-2861 | MIMAT0013803 | GGGGCCUGGCGGCGGGCGG |
| mmu-miR-3072 | MIMAT0014853 | UGCCCCCUCCAGGAAGCCUUCU |
| mmu-miR-3099 | MIMAT0014816 | UAGGCUAGAGAGAGGUUGGGGA |
| mmu-miR-3470a *10 | MIMAT0015640 | UCACUUUGUAGACCAGGCUGG |
| mmu-miR-3470b *10 | MIMAT0015641 | UCACUCUGUAGACCAGGCUGG |
| mmu-miR-3471 | MIMAT0015642 | UGAGAUCCAACUGUAAGGCAUU |
| mmu-miR-3472 | MIMAT0015643 | UAAUAGCCAGAAGCUGGAAGGAACC |
| mmu-miR-3473 | MIMAT0015645 | UGGAGAGAUGGCUCAGCC |
| mmu-miR-3474 | MIMAT0015646 | CCCUGGGAGGAGACGUGGAUUC |
| mmu-miR-3475 | MIMAT0015219 | UCUGGAGGCACAUGGUUUGAA |
| **Mouse Viral miRNA** | | |
| mcmv-miR-m01-1 | MIMAT0005533 | AGAGGAGAAUAACGUCGAACGG |
| mcmv-miR-m01-2 | MIMAT0005534 | GAAGAGAAUCGGGUUGGAACGGU |
| mcmv-miR-m01-3 | MIMAT0005536 | CGGUGAAGCGACUGUUGCCUCGA |
| mcmv-miR-m01-4 | MIMAT0005538 | UCCUAUGCUAACACGUGCGCGUG |
| mcmv-miR-m21-1 | MIMAT0005540 | AUAGGGGACACGUUCAAGCCG |
| mcmv-miR-m22-1 | MIMAT0005541 | UUCCCCGUCCGUACCGAGGCCA |
| mcmv-miR-M23-1-3p | MIMAT0005543 | CUCCUGCGUCGGCCCGAGGCC |
| mcmv-miR-M23-1-5p | MIMAT0005542 | CUCGGUACGGACGGGGAACCGU |
| mcmv-miR-M23-2 | MIMAT0005545 | AUGGGGGCCUCGGUCAAGCGG |
| mcmv-miR-M44-1 | MIMAT0005546 | UAUCUUUUUCCAGAGCCGCGGU |
| mcmv-miR-M55-1 | MIMAT0005547 | UGGUGAUCGGCGUGCUAGCCGU |
| mcmv-miR-m59-1 | MIMAT0005548 | UUAGCAGUGCCUCGACCGUCAG |
| mcmv-miR-m59-2 | MIMAT0005549 | CCCGAAGAGCCCUCACAGAGCC |
| mcmv-miR-M87-1 | MIMAT0005550 | AGGCAGCCGUCGGCAGCGGCAGC |
| mcmv-miR-m88-1 | MIMAT0005552 | CAGAAGUCGAUGUCGGGGUCU |
| mcmv-miR-M95-1-3p | MIMAT0005554 | AGCGACGUCGGACCGCGACGGC |
| mcmv-miR-M95-1-5p | MIMAT0005553 | GGUCGUGGGCUUGUGUCGCUUG |
| mcmv-miR-m107-1-3p | MIMAT0005556 | UGCUCGCGUCGAGUGACCGCUC |
| mcmv-miR-m107-1-5p | MIMAT0005555 | CGGUCACUCGUCUCGAGUCACC |
| mcmv-miR-m108-1 | MIMAT0005558 | UUUCUGACGGUGGCUCGUGUCG |
| mcmv-miR-m108-2-3p | MIMAT0005561 | GUGACUCGAGACGAGUGACCGGU |
| mcmv-miR-m108-2-5p.1 | MIMAT0005560 | GCGGUCACUCGACGCGAGCACCG |
| mcmv-miR-m108-2-5p.2 | MIMAT0005559 | UCACUCGUCGCGAGCGGUCAC |
| mghv-miR-M1-1 | MIMAT0001564 | UAGAAAUGGCCGUACUUCCUUU |
| mghv-miR-M1-2 | MIMAT0001565 | CAGACCCCCUCUCCCCCUCUUU |
| mghv-miR-M1-3 | MIMAT0001566 | GAGGUGAGCAGGAGUUGCGCUU |

| | | |
|---|---|---|
| mghv-miR-M1-4 | MIMAT0001567 | UCGAGGAGCACGUGUUAUUCUA |
| mghv-miR-M1-5 | MIMAT0001568 | AGAGUUGAGAUCGGGUCGUCUC |
| mghv-miR-M1-6 | MIMAT0001569 | UGAAACUGUGUGAGGUGGUUUU |
| mghv-miR-M1-7-3p | MIMAT0001571 | GAUAUCGCGCCCACCUUUAUU |
| mghv-miR-M1-7-5p | MIMAT0001570 | AAAGGUGGAGGUGCGGUAACCU |
| mghv-miR-M1-8 | MIMAT0001572 | AGCACUCACUGGGGGUUUGGUC |
| mghv-miR-M1-9 | MIMAT0001573 | UCACAUUUGCCUGGACCUUUUU |
| **Non-Mammalian Spike In miRNA probes** | | |
| ath-miR159a | MIMAT0000177 | UUUGGAUUGAAGGGAGCUCUA |
| cel-miR-248 | MIMAT0000304 | AUACACGUGCACGGAUAACGCUCA |
| osa-miR414 | MIMAT0001330 | UCAUCCUCAUCAUCAUCGUCC |
| | - | |
| Actb | NM_007393.1 | CAGGTCATCACTATTGGCAACGAGCGGTTCCGATGCCCTGAGGCTCTTTTCCAGCCTTCCTTCTTGGGTATGGAATCCTGTGGCATCCATGAAACTACAT |
| B2m | NM_009735.3 | CTGAACTGCTACGTAACACAGTTCCACCCGCCTCACATTGAAATCCAAATGCTGAAGAACGGGAAAAAAATTCCTAAAGTAGAGATGTCAGATATGTCCT |
| Gapdh | NM_008084.1 | ATGTGTCCGTCGTGGATCTGACGTGCCGCCTGGAGAAACCTGCCAAGTATGATGACATCAAGAAGGTGGTGAAGCAGGCATCTGAGGGCCCACTGAAGGG |
| Rpl19 | NM_009078.1 | GAAGAGGCTTGCCTCTAGTGTCCTCCGCTGCGGGAAAAAGAAGGTCTGGTTGGATCCCAATGAGACCAATGAAATCGCCAATGCCAACTCCCGTCAGCAG |
| © 2010 NanoString Technologies | | Rev 20101108 LBL-C0068-01 |

**Supplementary Table 3. nCounter Mouse v1.5 miRNA Gene List.** The nCounter Mouse v1.5 miRNA panel, used to examine miRNA expression in the mesencephalon and myenteric plexus of the large intestine in pre-symptomatic (ps)A30P and wild type (WT) mice, involves 578 mouse miRNAs, 33 viral mouse miRNAs, and seven non-mammalian spike in miRNA probes.

#### Supplementary Table 4

| miRNA | Counts WT | Counts psA30P | Fold Change | $Log_2$ Fold Change | *p*-value |
|---|---|---|---|---|---|
| mmu-miR-350 | 673 | 979 | 1.46 | 0.55 | 0.01316 |
| mmu-miR-22 | 3607 | 5583 | 1.55 | 0.63 | 0.01584 |
| mmu-miR-340-3p | 121 | 195 | 1.61 | 0.69 | 0.01623 |
| mmu-miR-15a | 524 | 1047 | 2.00 | 1.00 | 0.01889 |
| mmu-miR-125a-3p | 102 | 156 | 1.53 | 0.61 | 0.02587 |
| mmu-miR-374 | 99 | 152 | 1.54 | 0.62 | 0.03256 |
| mmu-miR-140 | 271 | 421 | 1.55 | 0.63 | 0.04988 |
| mmu-miR-152 | 205 | 312 | 1.53 | 0.61 | 0.05024 |
| mmu-miR-30c | 6356 | 8798 | 1.38 | 0.46 | 0.05345 |
| mmu-miR-28 | 100 | 140 | 1.41 | 0.50 | 0.05780 |
| mmu-miR-218 | 3190 | 4544 | 1.42 | 0.51 | 0.05830 |
| mmu-miR-99a | 2166 | 3194 | 1.47 | 0.56 | 0.06424 |
| mmu-miR-532-5p | 332 | 410 | 1.24 | 0.31 | 0.06681 |
| mmu-miR-338-5p | 74 | 115 | 1.56 | 0.64 | 0.06760 |
| mmu-miR-27a | 1592 | 2285 | 1.44 | 0.53 | 0.06884 |
| mmu-miR-126-3p | 4496 | 7026 | 1.56 | 0.64 | 0.07071 |
| mmu-miR-199a-3p | 95 | 165 | 1.73 | 0.79 | 0.07212 |
| mmu-miR-1983 | 74 | 109 | 1.47 | 0.56 | 0.07357 |
| mmu-miR-451 | 1565 | 2154 | 1.38 | 0.46 | 0.07604 |
| mmu-miR-25 | 447 | 641 | 1.44 | 0.53 | 0.07725 |
| mmu-miR-133a | 156 | 227 | 1.45 | 0.54 | 0.07808 |
| mmu-miR-872 | 179 | 312 | 1.75 | 0.81 | 0.07860 |
| mmu-miR-411 | 203 | 301 | 1.48 | 0.57 | 0.08255 |
| mmu-miR-337-3p | 73 | 120 | 1.63 | 0.70 | 0.08661 |
| mmu-miR-322 | 95 | 139 | 1.46 | 0.55 | 0.08795 |
| mmu-miR-126-5p | 395 | 603 | 1.53 | 0.61 | 0.09366 |
| mmu-miR-223 | 62 | 103 | 1.67 | 0.74 | 0.09765 |
| mmu-miR-338-3p | 2051 | 3626 | 1.77 | 0.82 | 0.09905 |
| mmu-miR-499 | 55 | 103 | 1.86 | 0.90 | 0.09974 |
| mmu-miR-151-3p | 95 | 136 | 1.43 | 0.52 | 0.10693 |
| mmu-miR-190 | 146 | 246 | 1.69 | 0.76 | 0.10883 |
| mmu-miR-19b | 58 | 108 | 1.86 | 0.90 | 0.10898 |
| mmu-miR-19a | 84 | 143 | 1.70 | 0.77 | 0.10939 |
| mmu-miR-27b | 96 | 166 | 1.73 | 0.79 | 0.11356 |
| mmu-miR-29c | 10702 | 15122 | 1.41 | 0.50 | 0.11683 |
| mmu-miR-323-3p | 167 | 226 | 1.35 | 0.43 | 0.11895 |
| mmu-miR-574-3p | 97 | 156 | 1.60 | 0.68 | 0.11937 |
| mmu-miR-15b | 435 | 602 | 1.38 | 0.46 | 0.11964 |
| mmu-miR-137 | 365 | 516 | 1.41 | 0.50 | 0.14262 |
| mmu-miR-100 | 1511 | 2282 | 1.51 | 0.59 | 0.14265 |
| mmu-miR-181c | 348 | 472 | 1.36 | 0.44 | 0.14714 |
| mmu-miR-96 | 97 | 169 | 1.73 | 0.79 | 0.14889 |

| | | | | | |
|---|---|---|---|---|---|
| mmu-miR-326 | 656 | 807 | 1.23 | 0.30 | 0.15348 |
| mmu-miR-206 | 263 | 375 | 1.43 | 0.52 | 0.15349 |
| mmu-miR-148b | 387 | 529 | 1.37 | 0.45 | 0.15569 |
| mmu-miR-210 | 62 | 104 | 1.69 | 0.76 | 0.16507 |
| mmu-miR-7a | 181 | 298 | 1.64 | 0.71 | 0.17117 |
| mmu-miR-148a | 656 | 896 | 1.37 | 0.45 | 0.17204 |
| mmu-miR-20a / miR-20b | 343 | 457 | 1.33 | 0.41 | 0.17367 |
| mmu-miR-434-5p | 320 | 448 | 1.40 | 0.49 | 0.17394 |
| mmu-miR-125b-3p | 273 | 343 | 1.25 | 0.32 | 0.17808 |
| mmu-miR-21 | 888 | 1404 | 1.58 | 0.66 | 0.17826 |
| mmu-miR-181a | 8682 | 11886 | 1.37 | 0.45 | 0.17925 |
| mmu-miR-142-3p | 137 | 188 | 1.38 | 0.46 | 0.18149 |
| mmu-miR-432 | 69 | 120 | 1.75 | 0.81 | 0.18365 |
| mmu-miR-134 | 104 | 135 | 1.29 | 0.37 | 0.18422 |
| mmu-miR-301a | 450 | 652 | 1.45 | 0.54 | 0.18767 |
| mmu-miR-9 | 19164 | 24260 | 1.27 | 0.34 | 0.18850 |
| mmu-miR-29b | 12160 | 16020 | 1.32 | 0.40 | 0.19996 |
| mmu-miR-369-5p | 75 | 109 | 1.44 | 0.53 | 0.20214 |
| mmu-miR-670 | 69 | 109 | 1.59 | 0.67 | 0.20469 |
| mmu-miR-300 | 1052 | 1310 | 1.25 | 0.32 | 0.20550 |
| mmu-miR-93 | 94 | 143 | 1.52 | 0.60 | 0.20582 |
| mmu-miR-34b-3p | 93 | 129 | 1.40 | 0.49 | 0.20586 |
| mmu-miR-29a | 15229 | 19724 | 1.30 | 0.38 | 0.20749 |
| mmu-miR-136 | 1952 | 2940 | 1.51 | 0.59 | 0.20798 |
| mmu-miR-340-5p | 1198 | 1595 | 1.33 | 0.41 | 0.20876 |
| mmu-miR-106a / miR-17 | 319 | 392 | 1.23 | 0.30 | 0.21363 |
| mmu-miR-26b | 578 | 815 | 1.41 | 0.50 | 0.21516 |
| mmu-miR-592 | 182 | 272 | 1.49 | 0.58 | 0.21705 |
| mmu-miR-467f | 83 | 148 | 1.78 | 0.83 | 0.22034 |
| mmu-miR-1 | 176 | 271 | 1.54 | 0.62 | 0.22187 |
| mmu-miR-130a | 588 | 835 | 1.42 | 0.51 | 0.22596 |
| mmu-miR-539 | 128 | 182 | 1.43 | 0.52 | 0.23449 |
| mmu-miR-151-5p | 2022 | 2546 | 1.26 | 0.33 | 0.23861 |
| mmu-miR-421 | 88 | 135 | 1.53 | 0.61 | 0.23868 |
| mmu-miR-876-3p | 127 | 186 | 1.46 | 0.55 | 0.23916 |
| mmu-miR-101b | 186 | 322 | 1.73 | 0.79 | 0.23917 |
| mmu-miR-582-5p | 145 | 250 | 1.72 | 0.78 | 0.23969 |
| mmu-miR-98 | 381 | 514 | 1.35 | 0.43 | 0.24639 |
| mmu-miR-376c | 581 | 843 | 1.45 | 0.54 | 0.24725 |
| mmu-miR-191 | 422 | 495 | 1.17 | 0.23 | 0.25393 |
| mmu-miR-181b / miR-181d | 102 | 147 | 1.45 | 0.54 | 0.25769 |
| mmu-miR-466a-3p /<br>miR-466b-3-3p | 83 | 112 | 1.34 | 0.42 | 0.26061 |
| mmu-miR-203 | 70 | 105 | 1.49 | 0.58 | 0.27198 |
| mmu-miR-139-5p | 172 | 229 | 1.33 | 0.41 | 0.27214 |
| mmu-miR-106b | 213 | 265 | 1.24 | 0.31 | 0.27765 |

| | | | | | |
|---|---|---|---|---|---|
| mmu-miR-34b-5p | 115 | 161 | 1.39 | 0.48 | 0.28248 |
| mmu-miR-135b | 111 | 159 | 1.43 | 0.52 | 0.29285 |
| mmu-miR-212 | 143 | 194 | 1.35 | 0.43 | 0.29522 |
| mmu-miR-674 | 110 | 147 | 1.34 | 0.42 | 0.29821 |
| mmu-miR-410 | 988 | 1228 | 1.24 | 0.31 | 0.29846 |
| mmu-miR-187 | 80 | 104 | 1.31 | 0.39 | 0.30628 |
| mmu-miR-365 | 218 | 308 | 1.41 | 0.50 | 0.30700 |
| mmu-miR-431 | 94 | 121 | 1.29 | 0.37 | 0.31113 |
| mmu-miR-551b | 117 | 148 | 1.26 | 0.33 | 0.31172 |
| mmu-miR-770-3p | 114 | 151 | 1.32 | 0.40 | 0.31508 |
| mmu-miR-30e | 344 | 441 | 1.28 | 0.36 | 0.31571 |
| mmu-miR-208a | 98 | 144 | 1.47 | 0.56 | 0.31858 |
| mmu-miR-409-5p | 93 | 117 | 1.26 | 0.33 | 0.33324 |
| mmu-miR-873 | 82 | 106 | 1.28 | 0.36 | 0.33333 |
| mmu-miR-200b | 165 | 264 | 1.60 | 0.68 | 0.33667 |
| mmu-miR-101a | 142 | 234 | 1.65 | 0.72 | 0.33678 |
| mmu-miR-146a | 352 | 476 | 1.35 | 0.43 | 0.34063 |
| mmu-miR-130b | 227 | 335 | 1.47 | 0.56 | 0.34944 |
| mmu-miR-466d-3p | 84 | 112 | 1.34 | 0.42 | 0.36539 |
| mmu-miR-192 | 85 | 110 | 1.29 | 0.37 | 0.37412 |
| mmu-miR-1196 | 118 | 168 | 1.42 | 0.51 | 0.38255 |
| mmu-miR-23a | 1428 | 1693 | 1.19 | 0.25 | 0.38380 |
| mmu-miR-153 | 429 | 600 | 1.40 | 0.49 | 0.38642 |
| mmu-miR-1956 | 82 | 106 | 1.29 | 0.37 | 0.40072 |
| mmu-miR-99b | 858 | 1056 | 1.23 | 0.30 | 0.40076 |
| mmu-miR-1839-5p | 194 | 239 | 1.23 | 0.30 | 0.40224 |
| mmu-miR-194 | 221 | 273 | 1.24 | 0.31 | 0.40613 |
| mmu-miR-690 | 165 | 222 | 1.35 | 0.43 | 0.40672 |
| mmu-miR-346 | 79 | 106 | 1.34 | 0.42 | 0.42482 |
| mmu-miR-146b | 108 | 132 | 1.22 | 0.29 | 0.42674 |
| mmu-miR-382 | 1616 | 2014 | 1.25 | 0.32 | 0.42899 |
| mmu-miR-221 | 105 | 142 | 1.35 | 0.43 | 0.43261 |
| mmu-let-7d | 10645 | 12291 | 1.15 | 0.20 | 0.43411 |
| mmu-let-7i | 4503 | 5582 | 1.24 | 0.31 | 0.43536 |
| mmu-miR-149 | 820 | 942 | 1.15 | 0.20 | 0.43630 |
| mmu-miR-486 | 209 | 245 | 1.17 | 0.23 | 0.43655 |
| mmu-miR-664 | 100 | 138 | 1.38 | 0.46 | 0.43815 |
| mmu-miR-16 | 8085 | 10487 | 1.30 | 0.38 | 0.44497 |
| mmu-miR-124 | 45528 | 57412 | 1.26 | 0.33 | 0.45081 |
| mmu-miR-541 | 89 | 122 | 1.37 | 0.45 | 0.45311 |
| mmu-miR-125a-5p | 7366 | 8148 | 1.11 | 0.15 | 0.45439 |
| mmu-miR-202-5p | 73 | 109 | 1.49 | 0.58 | 0.45869 |
| mmu-miR-384-3p | 425 | 544 | 1.28 | 0.36 | 0.45968 |
| mmu-miR-301b | 161 | 202 | 1.26 | 0.33 | 0.46041 |
| mmu-miR-154 | 544 | 680 | 1.25 | 0.32 | 0.46210 |
| mmu-miR-378 | 329 | 385 | 1.17 | 0.23 | 0.46268 |

| | | | | | |
|---|---|---|---|---|---|
| mmu-miR-423-3p | 105 | 129 | 1.23 | 0.30 | 0.46479 |
| mmu-miR-466g | 145 | 249 | 1.72 | 0.78 | 0.47419 |
| mmu-miR-425 | 196 | 224 | 1.14 | 0.19 | 0.48473 |
| mmu-miR-669f | 166 | 213 | 1.28 | 0.36 | 0.48530 |
| mmu-miR-33 | 104 | 142 | 1.37 | 0.45 | 0.48818 |
| mmu-miR-652 | 149 | 166 | 1.11 | 0.15 | 0.49147 |
| mmu-miR-487b | 665 | 759 | 1.14 | 0.19 | 0.49249 |
| mmu-miR-381 | 630 | 757 | 1.20 | 0.26 | 0.49374 |
| mmu-miR-150 | 1493 | 1688 | 1.13 | 0.18 | 0.49385 |
| mmu-miR-32 | 88 | 110 | 1.26 | 0.33 | 0.51087 |
| mmu-miR-377 | 326 | 394 | 1.21 | 0.28 | 0.51729 |
| mmu-miR-142-5p | 116 | 157 | 1.35 | 0.43 | 0.53565 |
| mmu-miR-543 | 719 | 371 | -1.94 | 0.96 | 0.53731 |
| mmu-miR-193b | 82 | 103 | 1.25 | 0.32 | 0.53809 |
| mmu-miR-335-5p | 479 | 580 | 1.21 | 0.28 | 0.53935 |
| mmu-miR-1198 | 163 | 221 | 1.36 | 0.44 | 0.54269 |
| mmu-miR-133b | 101 | 127 | 1.26 | 0.33 | 0.54385 |
| mmu-miR-497 | 106 | 138 | 1.30 | 0.38 | 0.55213 |
| mmu-miR-376a | 813 | 1040 | 1.28 | 0.36 | 0.55590 |
| mmu-miR-30b | 1733 | 2233 | 1.29 | 0.37 | 0.55747 |
| mmu-miR-219 | 4223 | 5490 | 1.30 | 0.38 | 0.56024 |
| mmu-miR-129-3p | 8206 | 6988 | -1.17 | 0.23 | 0.56047 |
| mmu-miR-345-5p | 157 | 177 | 1.13 | 0.18 | 0.56409 |
| mmu-miR-331-3p | 251 | 289 | 1.15 | 0.20 | 0.58737 |
| mmu-miR-2183 | 82 | 121 | 1.47 | 0.56 | 0.59274 |
| mmu-miR-135a | 4440 | 5215 | 1.17 | 0.23 | 0.59348 |
| mmu-miR-23b | 2080 | 2346 | 1.13 | 0.18 | 0.59713 |
| mmu-miR-145 | 2281 | 2665 | 1.17 | 0.23 | 0.60597 |
| mmu-miR-342-3p | 1843 | 2052 | 1.11 | 0.15 | 0.61224 |
| mmu-miR-423-5p | 93 | 107 | 1.15 | 0.20 | 0.61393 |
| mmu-miR-92b | 69 | 102 | 1.47 | 0.56 | 0.61636 |
| mmu-miR-329 | 635 | 698 | 1.10 | 0.14 | 0.61976 |
| mmu-miR-344 | 126 | 154 | 1.22 | 0.29 | 0.64813 |
| mmu-miR-369-3p | 1240 | 1440 | 1.16 | 0.21 | 0.65068 |
| mmu-miR-204 | 11512 | 13042 | 1.13 | 0.18 | 0.65652 |
| mmu-miR-143 | 577 | 647 | 1.12 | 0.16 | 0.65710 |
| mmu-miR-26a | 317 | 423 | 1.33 | 0.41 | 0.66187 |
| mmu-miR-24 | 284 | 334 | 1.18 | 0.24 | 0.67777 |
| mmu-miR-296-5p | 135 | 153 | 1.13 | 0.18 | 0.71110 |
| mmu-miR-376b | 2074 | 1932 | -1.07 | 0.10 | 0.72245 |
| mmu-miR-107 | 574 | 621 | 1.08 | 0.11 | 0.73248 |
| mmu-miR-132 | 4698 | 5017 | 1.07 | 0.10 | 0.74539 |
| mmu-miR-500 | 458 | 589 | 1.29 | 0.37 | 0.74556 |
| mmu-miR-128 | 1183 | 1379 | 1.17 | 0.23 | 0.77012 |
| mmu-miR-222 | 107 | 121 | 1.13 | 0.18 | 0.77355 |
| mmu-miR-30a | 2111 | 2288 | 1.08 | 0.11 | 0.77884 |

| | | | | | |
|---|---|---|---|---|---|
| mmu-miR-138 | 686 | 740 | 1.08 | 0.11 | 0.78293 |
| mmu-miR-324-5p | 264 | 290 | 1.10 | 0.14 | 0.79144 |
| mmu-miR-720 | 16605 | 19622 | 1.18 | 0.24 | 0.79653 |
| mmu-miR-384-5p | 589 | 630 | 1.07 | 0.10 | 0.79950 |
| mmu-miR-185 | 588 | 624 | 1.06 | 0.08 | 0.80460 |
| mmu-miR-1937a / miR-1937b | 834 | 1013 | 1.21 | 0.28 | 0.81960 |
| mmu-let-7f | 3481 | 3767 | 1.08 | 0.11 | 0.81985 |
| mmu-let-7a | 7764 | 8330 | 1.07 | 0.10 | 0.82310 |
| mmu-miR-483 | 44 | 48 | 1.10 | 0.14 | 0.83639 |
| mmu-miR-103 | 1436 | 1531 | 1.07 | 0.10 | 0.84132 |
| mmu-miR-195 | 436 | 466 | 1.07 | 0.10 | 0.84165 |
| mmu-miR-484 | 278 | 306 | 1.10 | 0.14 | 0.84282 |
| mmu-miR-144 | 308 | 335 | 1.09 | 0.12 | 0.84700 |
| mmu-miR-1224 | 104 | 111 | 1.06 | 0.08 | 0.86342 |
| mmu-miR-667 | 342 | 347 | 1.01 | 0.01 | 0.86382 |
| mmu-let-7e | 3244 | 3158 | -1.03 | 0.04 | 0.88527 |
| mmu-miR-1937c | 309 | 342 | 1.10 | 0.14 | 0.89026 |
| mmu-miR-34a | 625 | 657 | 1.05 | 0.07 | 0.92008 |
| mmu-miR-1944 | 1398 | 1363 | -1.03 | 0.04 | 0.93002 |
| mmu-miR-129-5p | 347 | 338 | -1.03 | 0.04 | 0.93559 |
| mmu-miR-127 | 2016 | 2061 | 1.02 | 0.03 | 0.94280 |
| mmu-let-7g | 20173 | 20767 | 1.03 | 0.04 | 0.94863 |
| mmu-miR-433 | 1064 | 1073 | 1.01 | 0.01 | 0.95642 |
| mmu-miR-125b-5p | 41770 | 42352 | 1.01 | 0.01 | 0.95913 |
| mmu-let-7c | 16307 | 16520 | 1.01 | 0.01 | 0.96525 |
| mmu-let-7b | 7149 | 7053 | -1.01 | 0.01 | 0.96911 |
| mmu-miR-361 | 411 | 417 | 1.01 | 0.01 | 0.97477 |
| mmu-miR-434-3p | 3266 | 3247 | -1.01 | 0.01 | 0.98172 |
| mmu-miR-328 | 472 | 476 | 1.01 | 0.01 | 0.98848 |
| mmu-miR-495 | 2023 | 2030 | 1.00 | 0.00 | 0.99062 |
| mmu-miR-30d | 2009 | 2009 | 1.00 | 0.00 | 0.99933 |

**Supplementary Table 4. Expressed miRNAs in the mesencephalon of pre-symptomatic (ps)A30P and wild type (WT) mice**. In total, 210 different miRNAs were expressed in the mesencephalon, and eight miRNAs showed a significantly different expression profile in psA30P mice compared with in WT mice. Data represent n = 5 (WT) and n = 6 (psA30P) independent experiments. * $p \le 0.05$, ** $p \le 0.01$, and *** $p \le 0.001$ using Student's t test.

#### Supplementary Table 5

| miRNA | Counts WT | Counts psA30P | Fold Change | $Log_2$ Fold Change | *p*-value |
|---|---|---|---|---|---|
| mmu-miR-338-3p | 221 | 574 | 2.60 | 1.38 | 0.00199 |
| mmu-miR-301a | 104 | 267 | 2.58 | 1.37 | 0.00210 |
| mmu-miR-674 | 86 | 118 | 1.37 | 0.45 | 0.00232 |
| mmu-miR-126-3p | 785 | 2647 | 3.37 | 1.75 | 0.00364 |
| mmu-miR-210 | 104 | 195 | 1.87 | 0.90 | 0.00537 |
| mmu-miR-146a | 258 | 484 | 1.88 | 0.91 | 0.00564 |
| mmu-miR-132 | 5071 | 6971 | 1.37 | 0.45 | 0.00585 |
| mmu-miR-377 | 97 | 223 | 2.31 | 1.21 | 0.00700 |
| mmu-miR-1937c | 73 | 152 | 2.08 | 1.06 | 0.00709 |
| mmu-let-7d | 9011 | 13316 | 1.48 | 0.57 | 0.00837 |
| mmu-miR-30c | 2923 | 5078 | 1.74 | 0.80 | 0.00865 |
| mmu-miR-136 | 312 | 735 | 2.35 | 1.23 | 0.00920 |
| mmu-miR-19a | 45 | 118 | 2.63 | 1.40 | 0.01196 |
| mmu-miR-10a | 2218 | 3172 | 1.43 | 0.52 | 0.01248 |
| mmu-miR-434-5p | 202 | 328 | 1.62 | 0.70 | 0.01263 |
| mmu-miR-486 | 79 | 103 | 1.30 | 0.38 | 0.01648 |
| mmu-miR-431 | 219 | 317 | 1.45 | 0.54 | 0.01665 |
| mmu-miR-126-5p | 48 | 172 | 3.54 | 1.82 | 0.01715 |
| mmu-miR-99a | 3491 | 4995 | 1.43 | 0.52 | 0.01836 |
| mmu-miR-574-3p | 151 | 230 | 1.53 | 0.61 | 0.01904 |
| mmu-miR-135a | 621 | 1096 | 1.77 | 0.82 | 0.01948 |
| mmu-miR-342-3p | 1200 | 1742 | 1.45 | 0.54 | 0.01965 |
| mmu-miR-350 | 296 | 551 | 1.86 | 0.90 | 0.02116 |
| mmu-miR-130a | 631 | 1354 | 2.14 | 1.10 | 0.02281 |
| mmu-miR-106a / miR-17 | 520 | 805 | 1.55 | 0.63 | 0.02338 |
| mmu-miR-543 | 449 | 648 | 1.44 | 0.53 | 0.02436 |
| mmu-miR-421 | 92 | 142 | 1.53 | 0.61 | 0.02527 |
| mmu-miR-149 | 1133 | 1498 | 1.32 | 0.40 | 0.02704 |
| mmu-miR-26b | 596 | 998 | 1.67 | 0.74 | 0.02754 |
| mmu-miR-20a / miR-20b | 599 | 956 | 1.60 | 0.68 | 0.02794 |
| mmu-miR-196b | 49 | 117 | 2.41 | 1.27 | 0.02950 |
| mmu-miR-27a | 6603 | 9802 | 1.48 | 0.57 | 0.03074 |
| mmu-miR-181a | 3332 | 5377 | 1.61 | 0.69 | 0.03112 |
| mmu-miR-140 | 92 | 175 | 1.91 | 0.93 | 0.03386 |
| mmu-miR-148a | 698 | 1255 | 1.80 | 0.85 | 0.03516 |
| mmu-miR-23a | 12425 | 17150 | 1.38 | 0.46 | 0.03922 |
| mmu-miR-22 | 5273 | 8584 | 1.63 | 0.70 | 0.04043 |
| mmu-let-7i | 5033 | 6759 | 1.34 | 0.42 | 0.04059 |
| mmu-miR-326 | 463 | 584 | 1.26 | 0.33 | 0.04252 |
| mmu-miR-125a-5p | 6293 | 8180 | 1.30 | 0.38 | 0.04637 |
| mmu-miR-720 | 4260 | 8337 | 1.96 | 0.97 | 0.04712 |

| | | | | | |
|---|---|---|---|---|---|
| mmu-miR-495 | 1075 | 1594 | 1.48 | 0.57 | 0.04756 |
| mmu-miR-532-5p | 177 | 243 | 1.37 | 0.45 | 0.04768 |
| mmu-miR-376c | 957 | 1239 | 1.29 | 0.37 | 0.04833 |
| mmu-miR-676 | 172 | 218 | 1.27 | 0.34 | 0.05002 |
| mmu-miR-29c | 3301 | 5850 | 1.77 | 0.82 | 0.05102 |
| mmu-miR-199a-3p | 111 | 271 | 2.44 | 1.29 | 0.05519 |
| mmu-miR-194 | 73 | 124 | 1.69 | 0.76 | 0.05610 |
| mmu-miR-434-3p | 1080 | 1606 | 1.49 | 0.58 | 0.05613 |
| mmu-miR-504 | 79 | 102 | 1.29 | 0.37 | 0.05626 |
| mmu-miR-15b | 459 | 701 | 1.53 | 0.61 | 0.05855 |
| mmu-miR-331-3p | 136 | 194 | 1.43 | 0.52 | 0.05873 |
| mmu-miR-148b | 155 | 270 | 1.75 | 0.81 | 0.06032 |
| mmu-miR-15a | 221 | 434 | 1.96 | 0.97 | 0.06035 |
| mmu-miR-21 | 449 | 1007 | 2.25 | 1.17 | 0.06049 |
| mmu-miR-152 | 384 | 662 | 1.73 | 0.79 | 0.06119 |
| mmu-miR-29b | 3932 | 6589 | 1.68 | 0.75 | 0.06214 |
| mmu-miR-381 | 606 | 1003 | 1.66 | 0.73 | 0.06224 |
| mmu-miR-484 | 93 | 141 | 1.52 | 0.60 | 0.06281 |
| mmu-miR-323-3p | 301 | 431 | 1.43 | 0.52 | 0.06549 |
| mmu-miR-181c | 159 | 213 | 1.34 | 0.42 | 0.06604 |
| mmu-miR-154 | 445 | 679 | 1.53 | 0.61 | 0.06636 |
| mmu-miR-376a | 269 | 553 | 2.06 | 1.04 | 0.06662 |
| mmu-miR-93 | 73 | 108 | 1.49 | 0.58 | 0.07060 |
| mmu-miR-9 | 2522 | 3691 | 1.46 | 0.55 | 0.07217 |
| mmu-miR-23b | 7016 | 10462 | 1.49 | 0.58 | 0.07341 |
| mmu-miR-99b | 763 | 1208 | 1.58 | 0.66 | 0.07389 |
| mmu-let-7e | 3120 | 4156 | 1.33 | 0.41 | 0.07530 |
| mmu-miR-199a-5p | 45 | 116 | 2.56 | 1.36 | 0.07743 |
| mmu-miR-203 | 106 | 157 | 1.48 | 0.57 | 0.08089 |
| mmu-miR-30a | 702 | 1244 | 1.77 | 0.82 | 0.08823 |
| mmu-miR-2134 | 80 | 226 | 2.82 | 1.50 | 0.09092 |
| mmu-miR-151-5p | 715 | 966 | 1.35 | 0.43 | 0.09134 |
| mmu-miR-100 | 3108 | 4220 | 1.36 | 0.44 | 0.09350 |
| mmu-miR-1937a / miR-1937b | 218 | 538 | 2.47 | 1.30 | 0.09423 |
| mmu-miR-301b | 69 | 109 | 1.59 | 0.67 | 0.09554 |
| mmu-miR-329 | 996 | 1295 | 1.30 | 0.38 | 0.09935 |
| mmu-miR-337-3p | 85 | 114 | 1.34 | 0.42 | 0.10064 |
| mmu-miR-382 | 2240 | 2966 | 1.32 | 0.40 | 0.10212 |
| mmu-miR-300 | 771 | 1067 | 1.38 | 0.46 | 0.10232 |
| mmu-miR-30d | 940 | 1398 | 1.49 | 0.58 | 0.10293 |
| mmu-miR-212 | 97 | 135 | 1.38 | 0.46 | 0.10454 |
| mmu-miR-345-5p | 83 | 113 | 1.37 | 0.45 | 0.10556 |
| mmu-miR-134 | 197 | 238 | 1.21 | 0.28 | 0.10771 |
| mmu-miR-124 | 1449 | 2224 | 1.54 | 0.62 | 0.11141 |
| mmu-miR-204 | 3402 | 4467 | 1.31 | 0.39 | 0.11291 |
| mmu-miR-29a | 12727 | 19261 | 1.51 | 0.59 | 0.11569 |

| | | | | | |
|---|---|---|---|---|---|
| mmu-miR-16 | 5113 | 7662 | 1.50 | 0.58 | 0.12082 |
| mmu-miR-365 | 199 | 342 | 1.72 | 0.78 | 0.12083 |
| mmu-miR-1839-5p | 177 | 230 | 1.30 | 0.38 | 0.12127 |
| mmu-miR-145 | 7294 | 15333 | 2.10 | 1.07 | 0.12216 |
| mmu-miR-411 | 192 | 255 | 1.33 | 0.41 | 0.12371 |
| mmu-miR-497 | 68 | 103 | 1.51 | 0.59 | 0.12815 |
| mmu-miR-10b | 265 | 406 | 1.53 | 0.61 | 0.13019 |
| mmu-miR-876-3p | 76 | 107 | 1.41 | 0.50 | 0.13344 |
| mmu-miR-27b | 289 | 467 | 1.61 | 0.69 | 0.13538 |
| mmu-miR-30e | 112 | 171 | 1.52 | 0.60 | 0.13589 |
| mmu-miR-1944 | 1962 | 2716 | 1.38 | 0.46 | 0.13877 |
| mmu-miR-384-3p | 139 | 213 | 1.53 | 0.61 | 0.14205 |
| mmu-miR-7a | 648 | 867 | 1.34 | 0.42 | 0.14567 |
| mmu-miR-28 | 96 | 138 | 1.43 | 0.52 | 0.15141 |
| mmu-miR-409-5p | 138 | 191 | 1.38 | 0.46 | 0.15145 |
| mmu-miR-872 | 225 | 293 | 1.30 | 0.38 | 0.15237 |
| mmu-miR-125b-3p | 271 | 330 | 1.22 | 0.29 | 0.15331 |
| mmu-miR-218 | 1095 | 1813 | 1.66 | 0.73 | 0.15364 |
| mmu-miR-25 | 560 | 721 | 1.29 | 0.37 | 0.15403 |
| mmu-miR-107 | 263 | 375 | 1.43 | 0.52 | 0.15527 |
| mmu-miR-410 | 1210 | 2016 | 1.67 | 0.74 | 0.15801 |
| mmu-let-7c | 27579 | 41392 | 1.50 | 0.58 | 0.15960 |
| mmu-miR-369-3p | 697 | 1017 | 1.46 | 0.55 | 0.16227 |
| mmu-let-7a | 7018 | 10924 | 1.56 | 0.64 | 0.16373 |
| mmu-miR-340-5p | 231 | 341 | 1.48 | 0.57 | 0.16395 |
| mmu-let-7f | 3124 | 5060 | 1.62 | 0.70 | 0.16434 |
| mmu-miR-125a-3p | 93 | 116 | 1.25 | 0.32 | 0.16445 |
| mmu-miR-127 | 1562 | 2407 | 1.54 | 0.62 | 0.16779 |
| mmu-miR-24 | 1239 | 1970 | 1.59 | 0.67 | 0.17124 |
| mmu-miR-139-5p | 116 | 133 | 1.15 | 0.20 | 0.17272 |
| mmu-miR-30b | 432 | 897 | 2.08 | 1.06 | 0.17427 |
| mmu-miR-2146 | 91 | 144 | 1.58 | 0.66 | 0.18055 |
| mmu-miR-369-5p | 83 | 100 | 1.21 | 0.28 | 0.18471 |
| mmu-miR-539 | 263 | 322 | 1.22 | 0.29 | 0.19182 |
| mmu-miR-433 | 982 | 1095 | 1.11 | 0.15 | 0.19247 |
| mmu-miR-487b | 872 | 1169 | 1.34 | 0.42 | 0.19277 |
| mmu-miR-335-5p | 2410 | 3007 | 1.25 | 0.32 | 0.19775 |
| mmu-miR-374 | 109 | 155 | 1.41 | 0.50 | 0.21242 |
| mmu-let-7b | 14036 | 17314 | 1.23 | 0.30 | 0.21804 |
| mmu-miR-103 | 956 | 1321 | 1.38 | 0.46 | 0.21899 |
| mmu-miR-106b | 87 | 139 | 1.60 | 0.68 | 0.22686 |
| mmu-miR-582-5p | 56 | 100 | 1.77 | 0.82 | 0.23251 |
| mmu-miR-130b | 274 | 451 | 1.64 | 0.71 | 0.23405 |
| mmu-miR-324-5p | 129 | 183 | 1.43 | 0.52 | 0.23659 |
| mmu-miR-137 | 666 | 1338 | 2.01 | 1.01 | 0.24042 |
| mmu-miR-2140 | 97 | 141 | 1.46 | 0.55 | 0.24467 |

| | | | | | |
|---|---|---|---|---|---|
| mmu-let-7g | 10721 | 13550 | 1.26 | 0.33 | 0.24821 |
| mmu-miR-138 | 408 | 536 | 1.32 | 0.40 | 0.25034 |
| mmu-miR-384-5p | 193 | 267 | 1.38 | 0.46 | 0.25607 |
| mmu-miR-98 | 534 | 749 | 1.40 | 0.49 | 0.27979 |
| mmu-miR-195 | 151 | 219 | 1.45 | 0.54 | 0.28870 |
| mmu-miR-1224 | 166 | 223 | 1.35 | 0.43 | 0.29073 |
| mmu-miR-361 | 280 | 439 | 1.57 | 0.65 | 0.29458 |
| mmu-miR-26a | 226 | 448 | 1.98 | 0.99 | 0.29629 |
| mmu-miR-652 | 109 | 133 | 1.22 | 0.29 | 0.29969 |
| mmu-miR-128 | 172 | 275 | 1.60 | 0.68 | 0.31060 |
| mmu-miR-143 | 1886 | 3275 | 1.74 | 0.80 | 0.31681 |
| mmu-miR-7b | 235 | 350 | 1.49 | 0.58 | 0.32218 |
| mmu-miR-101b | 96 | 138 | 1.44 | 0.53 | 0.32611 |
| mmu-miR-129-5p | 434 | 550 | 1.27 | 0.34 | 0.34346 |
| mmu-miR-2141 | 497 | 687 | 1.38 | 0.46 | 0.35028 |
| mmu-miR-378 | 168 | 240 | 1.42 | 0.51 | 0.36447 |
| mmu-miR-142-5p | 122 | 104 | -1.18 | 0.24 | 0.40759 |
| mmu-miR-376b | 1231 | 1524 | 1.24 | 0.31 | 0.41857 |
| mmu-miR-1902 | 214 | 391 | 1.82 | 0.86 | 0.42886 |
| mmu-miR-335-3p | 122 | 149 | 1.22 | 0.29 | 0.43472 |
| mmu-miR-200b | 148 | 170 | 1.14 | 0.19 | 0.43737 |
| mmu-miR-125b-5p | 38351 | 45294 | 1.18 | 0.24 | 0.43815 |
| mmu-miR-541 | 125 | 159 | 1.28 | 0.36 | 0.44637 |
| mmu-miR-2183 | 91 | 116 | 1.27 | 0.34 | 0.46379 |
| mmu-miR-425 | 101 | 122 | 1.21 | 0.28 | 0.48958 |
| mmu-miR-191 | 344 | 386 | 1.12 | 0.16 | 0.55111 |
| mmu-miR-129-3p | 4912 | 5455 | 1.11 | 0.15 | 0.60846 |
| mmu-miR-1196 | 150 | 133 | -1.13 | 0.18 | 0.61216 |
| mmu-miR-185 | 306 | 342 | 1.12 | 0.16 | 0.61622 |
| mmu-miR-423-5p | 113 | 119 | 1.06 | 0.08 | 0.68469 |
| mmu-miR-328 | 277 | 328 | 1.19 | 0.25 | 0.69059 |
| mmu-miR-667 | 315 | 327 | 1.04 | 0.06 | 0.73456 |
| mmu-miR-466g | 179 | 184 | 1.02 | 0.03 | 0.90796 |

**Supplementary Table 5. Expressed miRNAs in the myenteric plexus (MP) of the large intestine (LI) in pre-symptomatic (ps)A30P and wild type (WT) mice**. In total, 166 different miRNAs were expressed in the MP of the LI, while 45 miRNAs showed a significantly different expression profile in psA30P mice compared with in WT mice. Data represent n = 5 (WT) and n = 6 (psA30P) independent experiments. * $p \leq 0.05$, ** $p \leq 0.01$, and *** $p \leq 0.001$ using Student's t test.

#### Supplementary Table 6

**a**

##### Motility Recordings

| Parameter | Cohen's d small intestine |
|---|---|
| contraction number | 1.55 |
| mean interval (s) | -0.78 |
| velocity (mm/s) | 2.12 |

| Parameter | Cohen's d large intestine |
|---|---|
| contraction number | 1.63 |
| mean interval (s) | -1.28 |
| velocity (mm/s) | 2.44 |

**b**

##### Proteomics

| Protein Name | Cohen's d small intestine |
|---|---|
| Gbas | 20.72 |
| Atp11b | -4.60 |
| Dnase1 | 3.87 |
| Pnp | -3.40 |
| Cops4 | -2.90 |
| S100a6 | 2.83 |
| Apoa1bp | -2.81 |
| Eif3a | -2.79 |
| Dctn2 | 2.72 |
| Rps19 | 2.70 |
| Gart | -2.62 |
| Ndrg1 | -2.60 |
| Fermt2 | -2.54 |
| Blvra | -2.50 |
| Dnm1 | 2.48 |
| Tcp1 | 2.48 |
| Anxa6 | -2.41 |
| Ube2l3 | 2.40 |
| Clta | 2.37 |
| Pfn 1 | -2.31 |
| Myo1c | 2.27 |
| Prdx4 | -2.26 |
| Dguok | -2.25 |
| Gabarapl2 | 2.21 |
| Hspb1 | -2.21 |
| Ppp1cb | -2.17 |
| Banf1 | -2.17 |
| Cct3 | 2.15 |
| Yes 1 | 2.14 |
| Snap47 | 2.14 |
| Pgp | -2.13 |
| Fis 1 | 2.13 |
| Anxa 5 | -2.12 |
| Hnrnph2 | 2.11 |

| Protein Name | Cohen's d large intestine |
|---|---|
| csnK2b | 9.25 |
| Atox1 | 9.07 |
| Lypla2 | -4.45 |
| Acot13 | 3.65 |
| Rab2A | 3.48 |
| Vcp | 3.19 |
| Ptpa | 2.94 |
| Hsd17b10 | 2.92 |
| Ap2m1 | 2.87 |
| Srm | 2.86 |
| Pfdn2 | 2.86 |
| Ak2 | 2.85 |
| Cnn1 | -2.81 |
| Vamp2 | 2.61 |
| Ctsd | 2.56 |
| Ak1 | 2.43 |
| Ap2a1 | 2.40 |
| Rpl27 | 2.39 |
| Ahcy | 2.37 |
| Acat2 | 2.34 |
| Dctn2 | 2.32 |
| Anxa7 | 2.26 |
| Glod4 | 2.24 |
| Eif4a | 2.24 |
| Rps9 | 2.23 |
| Kctd12b | 2.23 |
| Tomm70a | 2.22 |
| Gnb5 | 2.21 |
| Clic1 | 2.19 |
| Synm | -2.18 |
| Psma6 | 2.16 |
| Kars | 2.16 |
| Mtpn | 2.14 |
| Ube2v1 | 2.14 |

| | |
|---|---|
| Ap2a1 | 2.07 |
| Dstn | -2.04 |
| Gpx1 | -2.01 |
| Glod4 | -1.99 |
| Lgals3 | 1.99 |
| Hadhb | -1.99 |
| Fabp4 | -1.98 |
| Serpinb9; serine | -1.97 |
| sept6 | 1.94 |
| Aldh6a1 | -1.92 |
| Cct2 | 1.92 |
| sept7 | 1.91 |
| Cct6A | 1.90 |
| Rab 3A | -1.88 |
| Psma2 | -1.87 |
| Ndufa10 | 1.83 |
| Glo1 | -1.81 |
| Rpl14 | 1.80 |
| Ap1g1 | -1.76 |
| Elavl4 | 1.71 |
| Coro2b | 1.70 |
| Arpc4 | -1.68 |
| Tubb4a | 1.68 |
| Ddt | -1.67 |
| Prep | -1.67 |
| Pabpc1 | 1.67 |
| Tubb 5 | 1.67 |
| Ywhah | 1.67 |
| Rps21 | 1.65 |
| Anxa 11 | -1.63 |
| Ola1 | -1.58 |
| Ak3 | -1.57 |
| Rbp1 | -1.56 |
| Ptgr2 | -1.54 |
| Psmb6 | 1.54 |
| Pgm2 | -1.54 |
| Gstm2 | -1.52 |
| Plcd1 | -1.52 |
| Pcca | -1.52 |
| Uba52 | -1.50 |

| | |
|---|---|
| Napg | 2.13 |
| Calu | 2.12 |
| Hnrnpa3 | 2.12 |
| Tubb4a | 2.11 |
| Npepps | 2.11 |
| Sept6 | 2.11 |
| Rack1 | 2.11 |
| Flna | -2.11 |
| Ppib | 2.10 |
| Pafah1b3 | 2.10 |
| Ranbp1 | 2.09 |
| Cygb | 2.09 |
| Pdlim5 | 2.07 |
| Lgals3 | 2.06 |
| Psma4 | 2.06 |
| Spata5 | 2.05 |
| Usp 5 | 2.05 |
| Fkbp2 | 2.05 |
| Rps5 | 2.05 |
| Twf1 | 2.04 |
| Gsr | 2.03 |
| Rala | 2.03 |
| Atp6v1g1 | 2.03 |
| Tpm1 | 2.03 |
| Stxbp1 | 2.03 |
| Vps26a | 2.02 |
| Fabp5 | 2.02 |
| Rpl17 | 2.02 |
| Pdha2 | 2.00 |
| Endod1 | 2.00 |
| H1f0 | 2.00 |
| Tmpo | 1.99 |
| Txn1 | 1.99 |
| Rab3 | 1.99 |
| Pafah1b2 | 1.98 |
| Pcbd1 | 1.98 |
| Rpl10a | 1.97 |
| Anxa3 | 1.97 |
| Tpm2 | -1.95 |
| Etfb | 1.95 |
| Kpnb1 | 1.95 |
| Nono | 1.94 |
| Tac1 | 1.93 |
| Arpc4 | 1.93 |
| Pea15a | 1.92 |
| Ppp2r2a | 1.88 |
| Mapre2 | 1.88 |
| Cct5 | 1.88 |
| Hist1h1e | -1.88 |
| Pank | 1.88 |
| Nsf | 1.88 |
| tripeptidyl peptidase I | 1.86 |
| Psme2 | 1.86 |
| Gpx1 | 1.86 |
| Sdpr | 1.83 |
| Sept7 | 1.83 |

| | |
|---|---|
| Fscn1 | 1.82 |
| Asl | 1.81 |
| Tsnax | 1.79 |
| Uchl3 | 1.78 |
| H1f0 | 1.78 |
| Erap1 | 1.78 |
| Isoc1 | 1.77 |
| Scgn | 1.76 |
| Ptprn | 1.76 |
| Fth1 | 1.75 |
| Oat | 1.75 |
| Prdx2 | 1.75 |
| Rplp2 | 1.73 |
| Pam | 1.73 |
| Sptan1 | 1.73 |
| Pgd | 1.73 |
| Sod1 | -1.72 |
| Snrpd1 | 1.72 |
| Map6 | 1.70 |
| Ppp3cc | 1.70 |
| Gnb1 | 1.69 |
| Mylk | -1.68 |
| Mapt | 1.68 |
| Cct3 | 1.68 |
| Nefl | 1.66 |
| Scrn1 | 1.66 |
| Hspa5 | -1.66 |
| Qdpr | 1.66 |
| Syt1 | 1.65 |
| Eef1b2 | 1.64 |
| Pfn2 | 1.64 |
| Prdx4 | 1.64 |
| Rbp1 | 1.62 |
| Ezr | 1.61 |
| Ran | 1.60 |
| Atp6v1e2 | 1.59 |
| Crip2 | 1.59 |
| Npm1 | 1.58 |
| Dlat | 1.58 |
| Tceb3 | -1.58 |
| S100a1 | 1.58 |
| Anxa5 | 1.58 |
| Naca | 1.56 |
| Cs | 1.55 |
| Prdx3 | 1.54 |
| Eef1b2 | 1.53 |
| S100a10 | 1.53 |
| Kif5b | 1.53 |
| Prkar1a | 1.53 |
| Calb2 | 1.52 |
| Fasn | 1.52 |
| Dst | 1.51 |
| Hnrnpd | 1.51 |
| Psma5 | 1.51 |
| Stip1 | 1.50 |
| Capg | 1.50 |

| | |
|---|---|
| Dhx9 | 1.49 |

#### Immunostaining Whole Mount

**c**

| Parameter | Cohen's d large intestine |
|---|---|
| Nefl expression/picture section (%) | 3.59 |
| Vamp2 expression/picture section (%) | 2.47 |
| Calb2 expression/picture section (%) | 3.11 |
| ganglionic area/picture section (%) | 2.97 |

#### Immunocytochemistry

**d**

| Parameter | Cohen's d myenteric plexus cells |
|---|---|
| ratio Nefl/PGP9.5 | 1.76 |
| ratio Calb2/PGP9.5 | -2.54 |
| Vamp2 expression/picture section (%) | 3.01 |
| PGP9.5$^{+}$ cells/picture section | 3.23 |
| Nefl$^{+}$ cells/picture section (%) | 3.21 |
| Calb2$^{+}$ cells/picture section (%) | -1.90 |
| Live-Dead-Assay | 4.70 |

**e**

#### miRNA Expression Profile

| miRNA name | Cohen's d mesencephalon |
|---|---|
| mmu-miR-350 | -1.77 |
| mmu-miR-22 | -1.78 |
| mmu-miR-340-3p | -2.03 |
| mmu-miR-15a | -1.53 |
| mmu-miR-125a-3p | -1.55 |
| mmu-miR-374 | -1.81 |
| mmu-miR-140 | -1.19 |
| mmu-miR-152 | -1.78 |

| miRNA name | Cohen's d myenteric plexus large intestine |
|---|---|
| mmu-miR-338-3p | -2.33 |
| mmu-miR-301a | -1.82 |
| mmu-miR-674 | -2.53 |
| mmu-miR-126-3p | -1.39 |
| mmu-miR-210 | -2.12 |
| mmu-miR-146a | -1.87 |
| mmu-miR-132 | -2.03 |
| mmu-miR-377 | -1.55 |
| mmu-miR-1937c | -1.70 |
| mmu-let-7d | -1.71 |
| mmu-miR-30c | -1.80 |
| mmu-miR-136 | -1.73 |

| | |
|---|---|
| mmu-miR-19a | -1.31 |
| mmu-miR-10a | -2.07 |
| mmu-miR-434-5p | -2.09 |
| mmu-miR-486 | -1.74 |
| mmu-miR-431 | -1.69 |
| mmu-miR-126-5p | -1.10 |
| mmu-miR-99a | -1.55 |
| mmu-miR-574-3p | -1.50 |
| mmu-miR-135a | -1.75 |
| mmu-miR-342-3p | -1.58 |
| mmu-miR-350 | -1.48 |
| mmu-miR-130a | -1.20 |
| mmu-miR-106a / miR-17 | -1.65 |
| mmu-miR-543 | -1.70 |
| mmu-miR-421 | -1.64 |
| mmu-miR-149 | -1.63 |
| mmu-miR-26b | -1.40 |
| mmu-miR-20a / miR-20b | -1.68 |
| mmu-miR-196b | -1.04 |
| mmu-miR-27a | -1.52 |
| mmu-miR-181a | -1.49 |
| mmu-miR-140 | -1.16 |
| mmu-miR-148a | -1.30 |
| mmu-miR-23a | -1.38 |
| mmu-miR-22 | -1.28 |
| mmu-let-7i | -1.48 |
| mmu-miR-326 | -1.85 |
| mmu-miR-125a-5p | -1.59 |
| mmu-miR-720 | -1.49 |
| mmu-miR-495 | -1.55 |
| mmu-miR-532-5p | -1.37 |
| mmu-miR-376c | -1.30 |
| mmu-miR-676 | -1.33 |

**Supplementary Table 6. List of Cohen's d effect size[132] calculations.** When measuring if two groups have similar standard deviations (SDs) and are the same size, Cohen's d gives the appropriate effect size[132]. d is defined as the difference between the means of two groups divided by the SD of either group. Effect sizes are defined to have a small (d = 0.2), a medium (d = 0.5), or a large effect (d = 0.8 or d ≥ 0.8). (a) Cohen's d calculations from gut motility recordings with different parameters, e.g. contraction numbers, mean interval, and velocity for the small intestine and large intestine (LI) of pre-symptomatic (ps)A30P and wild type (WT) mice. (b) Cohen's d effect sizes were estimated for significantly changed psA30P vs. WT proteomic data. (c) Cohen's d values from quantifications of neurofilament light chain (Nefl), vesicle-associated membrane protein 2 (Vamp2), and calbindin 2 (Calb2) expression as well as percentage of ganglionic areas per picture section in whole mounts from psA30P and WT mice. (d) Respective expression ratios (protein gene product (PGP) 9.5, Nefl, Calb2, Vamp2, live-dead-assay) and cell numbers (PGP9.5, Nefl, Calb2, live-dead-assay) from *in vitro* studies using myenteric plexus (MP) cells (Postnatal day 2) exposed to A30P α-synuclein (0.5 μM) for 5 days and corresponding controls. (e) Calculations of Cohen's d effect size for significantly dysregulated miRNAs in the mesencephalon and MP of the LI derived in psA30P and WT mice.

**Supplementary video 1:** Motility recordings during luminal perfusion of the small intestine in 2-month-old wild type mice.

**Supplementary video 2:** Motility recordings during luminal perfusion of the small intestine in 2-month-old pre-sysmptomatic A30P mice.

**Supplementary video 3:** Motility recordings during luminal perfusion of the large intestine in 2-month-old wild type mice.

**Supplementary video 4:** Motility recordings during luminal perfusion of the large intestine in 2-month-old pre-sysmptomatic A30P mice.
